# The human RNA-DNA interactome is cell type-specific and dynamic

**DOI:** 10.64898/2026.08.25.746868

**Authors:** Alice Lambolez, Pelin Sahlén, Wenjing Kang, Xufeng Shu, Jessica Severin, Rodrigo Pracana, Ilyes Abdelhamid, Bhavya Dhaka, Christophe Vroland, Valeria Ranzani, Benedetto Polimeni, Masaru Koido, Andrea Vandelli, Maria Mintseva, Marton Rohaly Medved, Kayoko Yasuzawa, Mitsuyoshi Murata, Diane Delobel, Wing Hin Yip, Hiromi Nishiyori-Sueki, Satoshi Takizawa, Tomoe Nobusada, Matthew Brown, Valeria Di Gioia, Yoshimi Inaba, Sachi Kato, Callum Parr, Kentaro Kaji, Tsugumi Kawashima, Tsukasa Kouno, Michihira Tagami, Kokoro Ozaki, Rebecca Vadalà, Federica Marasca, Elisabetta Cozzi, Robert Krautz, Christian Vaagensø, Tomohiro Yamazaki, Xiaoze Li Wang, Quentin Verron, Yuichi Ichikawa, Jen-Chien Chang, Matthew Valentine, Hjörleifur Einarsson, Jonathan Moody, Akira Hasegawa, Ziheng Liao, Kohei Tomizuka, Artemii Zhigulev, Marco Gaviraghi, Zekiye Altan, Stefania Policicchio, Yifan Yu, Claudia Latini, Rossana Piccioni, Chiara Medaglia, Amanda Piveta Schnepper, Laura Carpen, Eugenia Ricciardelli, Matteo Bonfanti, Roberta Bosotti, Camilla Callierotti, Giulia Maria Scotti, Yung-Li Chen, Vipin Kumar, Aslihan Karabacak, Xiangfu Zhong, Aonghus Naughton, Irene Farabella, Pauline Robbe, Nicola Pirastu, Giovanni Pascarella, Lokesh P. Tripathi, Vincenzo Lagani, Robert Lehmann, Jesper Nils Tegnér, David Gomez-Cabrero, Lorenzo Calviello, Yari Ciani, Eugenio Morelli, Ivano Legnini, Colin A. Semple, Alok Sharma, Alessandro Vitriolo, Yuki Ishikawa, Laura Broglia, Miguel Ángel Berrocal Rubio, Ilaria Nisoli, Stefania Giussani, Xiaolin Li, Ilaria Castiglioni, Matteo Jacopo Marzi, Imad Abugessaisa, Stefano Gustincich, Hideya Kawaji, Yasuhiro Murakawa, Yasushi Okazaki, Nicole Soranzo, Blagoje Soskic, Giuseppe Testa, Alberto Riva, Clelia Peano, Michiel Jan Laurens De Hoon, Oliver Harschnitz, Francesco Nicassio, Nicola Crosetto, Musa Mhlanga, Dafne Campigli Di Giammartino, Chung Chau Hon, Sören Lehmann, Robin Andersson, Noriko Saitoh, Gian Gaetano Tartaglia, Anthony Mathelier, Tetsuro Hirose, Andreas Lennartsson, Beatrice Bodega, Chikashi Terao, Rory Johnson, Remo Sanges, Christine Anne Wells, Charles-Henri Lecellier, Magda Bienko, Robert S. Young, Carlo Vittorio Cannistraci, Jay Woo Shin, Takeya Kasukawa, Chi Wai Yip, Masaki Kato, Hazuki Takahashi, Piero Carninci

## Abstract

More than twenty years ago, the FANTOM consortium uncovered that mammalian genomes are pervasively transcribed, revealing multitudes of RNAs with unknown functions. A subset of these transcripts has since then been linked to transcriptional control and to chromatin organization *via* their ability to interact with DNA, suggesting that chromatin-associated RNAs could be key players in genome regulation. Although recent technological advances now enable the mapping of genome-wide RNA-DNA contacts, a lack of analyses integrating these methods with other genomic features and across multiple cellular contexts hinders our comprehensive understanding of the principles underlying RNA-DNA interactions and of their biological importance. As part of the FANTOM6 project, we thus generated RNA-DNA interaction maps in 16 different human cell types, then combined these contacts with multiple layers of other genomic data to investigate how patterns of interaction between RNA and DNA relate to chromatin organization and function. We show that the RNA-DNA interactome is highly dynamic yet reproducibly organized in cell-type specific networks, constituted of a great diversity of interactions that vary in function of their distance, the nature of their sources and the chromatin state of their targets. In particular, we detected numerous regulatory elements that exhibit marked changes in activity when differentially bound by transcripts, implying that thousands of RNA-DNA interactions can play a mechanistic role in gene expression. This regulatory function correlates with RNA-protein interactions and significantly associates with cell type-relevant and disorder-related traits. In addition to providing essential resources for future research in RNA-mediated chromatin regulation, cellular biology and human diseases, our study thus establishes the RNA-DNA interactome as a new genome regulatory layer that defines and maintains cellular identity and behavior.

## Introduction

Since its launch in 2000, the Functional ANnoTation Of the Mammalian genome (FANTOM) initiative has substantially contributed to shaping our understanding of mammalian transcriptomes and their regulatory architecture (Abugessaisa *et al*., 2020). The FANTOM3 project notably revealed that more than 60% of the mammalian genome is transcribed, despite only 2% encoding proteins (Carninci *et al*., 2005). Subsequent advances in the completion of the human genome catalog have resulted in a greater number of gene models annotated as “non-coding” than “protein-coding” (Amaral *et al*., 2023). Transcripts produced by these loci are collectively referred to as non-coding RNAs (ncRNAs) and comprise: ribosomal RNA (rRNAs) and transfer RNAs (tRNAs), which participate in protein translation as part of the ribosomal complex; small nuclear RNAs (snRNAs) and small nucleolar RNAs (snoRNAs), which are involved in transcript maturation within the spliceosome and the small nucleolar ribonucleoprotein particles (snoRNPs), respectively (Z. Huang *et al*., 2022; Y. Su *et al*., 2024); and small regulatory RNAs such as microRNAs (miRNAs) and PIWI-interacting RNAs (piRNAs), which associate with proteins of the Argonaute family to post-transcriptionally regulate gene expression and transposable elements (Saliminejad *et al*., 2018; X. Wang *et al*., 2022). Another major class is constituted by long non-coding RNAs (lncRNAs), defined broadly as any ncRNA longer than 200 nucleotides (nt). LncRNAs include most of the tens of thousands of enhancer RNAs (eRNAs) identified by large-scale efforts such as the ENCODE and FANTOM5 projects (Andersson *et al*., 2014; The ENCODE Project Consortium, 2020) that are transcribed from distal regulatory elements and contribute to the transcriptional activation of their target loci (Q. Chen *et al*., 2023).

The functions and modes of action of only a limited number of human lncRNAs have been thoroughly characterized so far. Nevertheless, even this small subset exhibits a remarkable diversity, with roles in chromatin organization and remodeling, gene expression, RNA processing, and protein translation and modification (L.-L. Chen & Kim, 2024; Mattick *et al*., 2023; Qian *et al*., 2019; Statello *et al*., 2020; X. Zhang *et al*., 2019). A well-known example is *X-INACTIVE SPECIFIC TRANSCRIPT* (*XIST*), which orchestrates X chromosome inactivation in mammalian females. *XIST* silences one of the X chromosomes by recruiting POLYCOMB REPRESSIVE COMPLEX 1 and 2 (PRC1 and PRC2), which catalyze repressive histone H2A ubiquitination (H2Aub) and trimethylation of the 27^th^ lysine of histone H3 (H3K27me3), respectively (Almeida *et al*., 2017; Engreitz *et al*., 2013). At the same time, *XIST* also concentrates SPEN, a protein facilitating histone deacetylation and removal of RNA Polymerase II (Dossin *et al*., 2020). While *XIST* functions as a repressor, other lncRNAs were demonstrated to function as activators of transcription. For instance, *ZINC FINGER E-BOX BINDING HOMEOBOX 1 ANTISENSE RNA 1* (*ZEB1-AS1*) is an antisense transcript that promotes the expression of its corresponding sense gene, in part by recruiting MIXED LINEAGE LEUKEMIA 1 (MLL1), which deposits activating trimethylation on the 4^th^ lysine of histone H3 (H3K4me3) at the *ZEB1* promoter (W. Su *et al*., 2017).

Aside from transcriptional regulation, some lncRNAs have been associated with chromatin and nuclear organization, like *TELOMERIC REPEAT-CONTAINING RNA* (*TERRA*) which is transcribed from the subtelomeric regions, then forms R-loops with telomeres and participates in their maintenance together with the homology-directed repair machinery (Kyriacou & Lingner, 2024). Likewise, eRNAs containing Alu sequences can form duplexes with promoter-derived antisense RNAs, thereby modulating chromatin looping and contributing to enhancer-promoter interaction specificity (Liang *et al*., 2023). A similar structural function is also played by lncRNA *NUCLEAR ENRICHED ABUNDANT TRANSCRIPT 1* (*NEAT1*) in the formation of nuclear paraspeckles: these condensates, which participate in regulating gene expression, arise through phase separation driven by multivalent protein integration with *NEAT1* scaffolds (Ingram & Fox, 2024). Nuclear speckles, another type of subnuclear bodies that act as pre-mRNA processing hubs, are instead abundant in lncRNA *METASTASIS ASSOCIATED LUNG ADENOCARCINOMA TRANSCRIPT 1* (*MALAT1*; Arun *et al*., 2020).

Results from the FANTOM5 project (Hon *et al*., 2017) and pilot studies for the FANTOM6 project (Ramilowski *et al*., 2020; Yip *et al*., 2022) revealed that nearly 70% of lncRNA-producing loci are associated with functional traits, and that knocking down a subset of them produces reproducible molecular phenotypes. Furthermore, mutations or misregulation of chromatin-associated RNAs have been implicated in a wide variety of human diseases and notably in cancer, highlighting their crucial cellular roles (Arun *et al*., 2020; Ghafouri-Fard *et al*., 2023; Kroupa *et al*., 2022; Liang *et al*., 2023; Morelli *et al*., 2025; W. Wang *et al*., 2021; Z. Wang *et al*., 2020). For instance, high levels of *ZEB1-AS* transcription is associated with increased cell cycle progression and metastatic potential in hepatocarcinoma (Mu *et al*., 2021), while overexpression of *TERRA* leads to break-induced replication of telomeres, which contributes to the telomere instability seen in multiple types of cancer (Silva *et al*., 2021). These findings suggest that many ncRNAs are functionally relevant and prompt the question of whether the mechanisms elucidated for well-studied transcripts, such as *XIST*, *TERRA*, or *ZEB1-AS1*, represent isolated cases or reflect more general principles applicable to a broader class of RNAs. Unfortunately, these well-characterized examples represent only an infinitesimal fraction of the 88,000 lncRNA-producing loci present in the human genome (Yip *et al*., 2024).

In all the cases described above, the function of the transcript involves its interaction with chromatin, either directly or through its association with protein complexes. Consistent with this observation, more than a third of the lncRNAs detected by our pilot study in human dermal fibroblasts (HDF) were found to be chromatin-bound (Ramilowski *et al*., 2020), raising the possibility that RNA-mediated regulation relies on direct and/or indirect RNA-DNA interaction. Understanding the roles and mechanisms of ncRNAs at a larger scale thus implies comprehensively identifying all chromatin-associated transcripts and their target sites. Several technologies have been developed to survey RNA-DNA interactions with an “all-to-all” approach, including Global RNA Interactions with DNA by deep sequencing (GRID-seq; X. Li *et al*., 2017), Chromatin-Associated RNA sequencing (ChAR-seq, Bell *et al*., 2018), Mapping of RNA-Genome Interactions (MARGI and iMARGI, Sridhar *et al*., 2017; Yan *et al*., 2019), RNA ends on DNA capture (Red-C, Gavrilov *et al*., 2020) and RNA And DNA Interacting Complexes Ligated and sequenced (RADICL-seq, Bonetti *et al*., 2020). While studies using these technologies have laid the foundation for our current understanding of the RNA-DNA interactome, the typically low and cell type-specific expression of ncRNAs (Hon *et al*., 2017) necessitates the profiling of their interactions across a large range of cell types to fully capture their complexity. In particular, the few mammalian studies that have examined RNA-DNA contacts in a transitional context, namely during embryonic development (Bonetti *et al*., 2020; Limouse *et al*., 2023) and in endothelial cells exposed to diabetes-like stress (Calandrelli *et al*., 2020), revealed their substantial remodeling across these processes, which was not necessarily reflected by changes in the source RNAs expression. The RNA-DNA interactome thus needs to be studied as a dynamic system, notably *via* the use of time-series, and with regards to the broader chromatin context, *via* the integration of RNA-DNA contacts with an array of complementary genomic data.

As part of the FANTOM6 project, we addressed this need by performing RADICL-seq across 16 distinct human samples, which comprise multiple ENCODE cell lines, different types of cancer cells including primary samples, as well as several time-series recapitulating distinct pathways of cellular specialization. The present study provides a comprehensive overview of these datasets, through which we further demonstrate the cell type-specificity, diversity and temporal dynamics of the RNA-DNA interactome. To illustrate the multiple applications of the FANTOM6 collection, we integrated our RADICL-seq data with a wide variety of pre-existing and newly generated high-throughput sequencing data, such as cap-trap full-length cDNA sequencing (CFC-seq), high-throughput chromosome conformation capture (Hi-C), Cleavage Under Targets and Tagmentation (CUT&Tag) or Assay for Transposase-Accessible Chromatin using sequencing (ATAC-seq). This strategy allowed us to explore the modalities and effects of RNA-DNA interactions with respect to the three-dimensional (3D) organization of the chromatin as well as the transcriptional, epigenetic and RNA-RNA interaction landscapes. We show that the RNA-DNA interactome network is organized around thousands of pivotal transcripts that appear to modulate the transcriptional remodeling accompanying cell differentiation. Our analyses using RNA-protein binding predictions and publicly available ChIP-seq data suggest that such RNA-based regulation involves a wide array of RNA-binding proteins. Integrating our results with Gene Ontology (GO) and genome-wide association studies (GWAS) databases further revealed that chromatin-associated RNAs control pathways that are essential for cellular identity and human health.

Altogether, the FANTOM6 consortium hereby provides an unprecedentedly diverse resource of RNA-DNA contacts that can be easily embedded within the broader chromatin landscape thanks to an array of companion datasets. This study defines the general principles of RNA-DNA interactions shared across mammalian cell types and, in doing so, establishes the RNA-DNA interactome as a widespread and structured layer of gene regulation. As we profiled both extensively characterized cell lines and cell types relevant for the study of various human diseases, and given the implication of RNA-DNA interactions in an ever-rising number of pathways and disorders, we expect the FANTOM6 data collection to serve as a valuable resource for both transcriptomic and medical research.

## Results

### The FANTOM6 data collection provides an integrative cartography of the human RNA-DNA interactome

To broaden the public catalog of RNA-DNA interactions, we profiled whole-genome RNA-DNA contacts across 16 different human cellular contexts by RADICL-seq, a proximity ligation-based method (Bonetti *et al*., 2020), using at least two biological replicates per sample (Supplementary Table 1).

Half of these samples belong to three time-series designed to explore how RNA-DNA interactions evolve during cell specialization (Figure 1; Supplementary Table 1): a neural differentiation series (“Neuron series”) to study chromatin-associated transcript dynamics along neuronal commitment, and two white blood cell activation series, set respectively in macrophages (“THP-1 series”) and in T cells (“T cell series”), to investigate changes in the RNA-DNA interactome within the immune context. The neuron series was generated from human induced pluripotent stem cells (“iPSC” sample) that were differentiated into neural stem cells (“NSC” sample) and subsequently into cortical neurons (“Neuron” sample). In the THP-1 series, THP-1 monocytes were stimulated with phorbol 12-myristate 13-acetate (PMA) to induce their differentiation into macrophages, with sampling at early (“PMA 24h” sample) and late (“PMA 96h” sample) time points, alongside a dimethyl sulfoxide (DMSO)-treated control (“DMSO 96h” sample). The T cell series consists of naïve T cells obtained from donors (“T cell Naïve” sample) that were activated *in vitro* (“T cell Activated” sample; see Methods).

**Figure 1:**
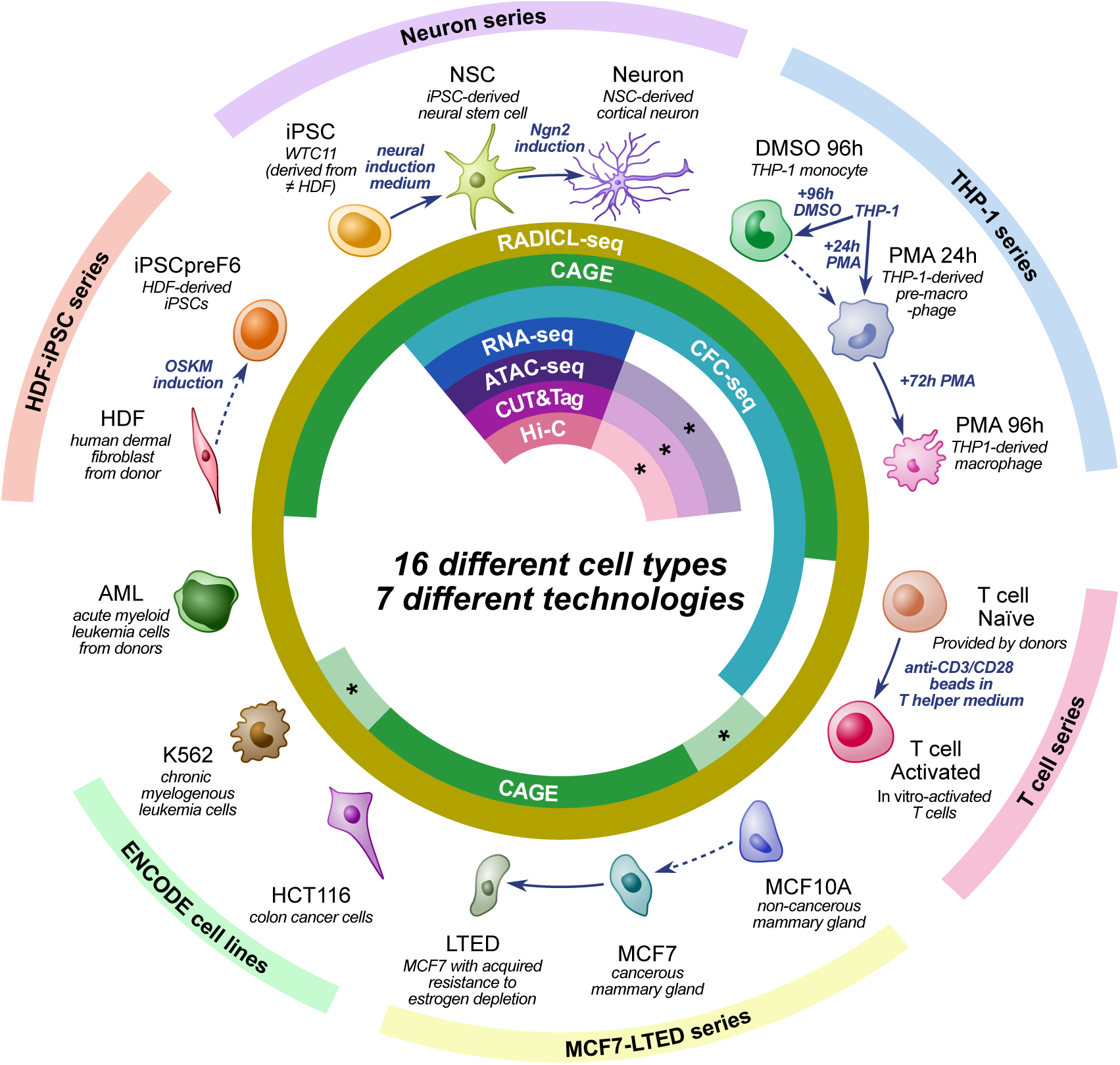
Description of the cell types and technologies used in this study. All datasets were generated by the FANTOM6 consortium except those marked with an asterisk *, which correspond to equivalent, publicly available data generated by other groups.

The other half of our samples comprises 6 different cell lines with established relevance to cancer research: healthy and malignant mammary gland cells (“MCF10A” and “MCF7” samples, respectively) as well as MCF7 cells with acquired resistance to estrogen depletion (“LTED” sample), colon cancer cells (“HCT116” sample), chronic myelogenous leukemia lymphoblasts (“K562” sample) and acute myeloid leukemia leukocytes which were obtained from patients (“AML” sample).

Finally, we also included RADICL-seq data from HDFs (“HDF” sample) as well as iPSCs derived from these cells (“iPSCpreF6” sample), part of which were previously published in our aforementioned pilot studies (Ramilowski *et al*., 2020; Yip *et al*., 2022).

Aside from these 16 samples, our FANTOM6 sample collection additionally features data generated in an iPSC-to-macrophage series, as listed in Supplementary Table 2. These samples have not been examined in the present study but are described in detail in Berrocal-Rubio *et al*. (2026).

With the aim of investigating these RNA-DNA contacts in their global cellular environment, we conjointly measured the expression landscape in our samples using cap analysis of gene expression (CAGE; Murata *et al*., 2014; Takahashi *et al*., 2021), CFC-seq (Delobel *et al*., 2025; Yip *et al*., 2024) and/or RNA-seq (Figure 1; Supplementary Tables 1 and 3).

To assess whether the RNA-DNA interactome is linked with the 3D conformation of the chromatin, we performed Hi-C combined with CUT&Tag (Kaya-Okur *et al*., 2019) for CCCTC-binding factor (CTCF) in the Neuron series. Similarly, to evaluate the epigenomic profile of RNA-targeted sites, we subjected these samples to single-nucleus ATAC-seq (snATAC-seq) and CUT&Tag for mono- and trimethylation of the 4^th^ of histone H3 (H3K4me1 and H3K4me3), as well as for acetylation and trimethylation of the 27^th^ lysine 27 of histone H3 (H3K27ac and H3K27me3; Figure 1; Supplementary Table 1). For the THP-1 series, we made use of publicly available datasets generated by digestion-ligation-only (DLO) Hi-C, ATAC-seq and chromatin immunoprecipitation followed by sequencing (ChIP-seq), which were previously published in Lin *et al*. (2022) and Phanstiel *et al*. (2017) (Figure 1; Supplementary Table 3).

The FANTOM6 data collection also provides data obtained by Psoralen Analysis of RNA Interactions and Structures with High Throughput and Resolution (PARIS; Lu *et al*., 2018), targeted chromosome conformation capture (Promoter Capture Hi-C; Sahlén *et al*., 2015) and genomic loci positioning by sequencing (GPSeq; Girelli *et al*., 2020) in the Neuron series (Supplementary Table 2). PARIS reveals intra- and intermolecular RNA-RNA contacts, while Promoter Capture Hi-C consists of high-resolution maps of promoter-anchored DNA-DNA interactions, and GPseq profiles the radial positioning of the genome. These data have not been used in the present study, but our Promoter Capture Hi-C and GPseq datasets are fully analyzed in Sahlén *et al*. (2025) and Kang *et al*. (2025), respectively.

Altogether, these complementary data enable the study of RNA-DNA interactions in conjunction with the transcriptomic and chromatin contexts of our samples.

Given that many RNAs originating from unannotated loci were previously found to interact with the chromatin (Limouse *et al*., 2023) and in order to maximize the accuracy and completeness of our analyses, we used a custom human genome annotation, hereinafter referred to as SACAGE (<u>SA</u>LA+<u>CA</u>T+<u>GE</u>NCODE). The SACAGE annotation combines (a) new Start-site Aware Long-read Assembler (SALA)-derived gene models obtained from CFC-seq data in our Neuron and THP-1 series, as described in Yip *et al*. (2024), (b) the CAGE-associated transcriptome (CAT) gene models defined from FANTOM5 promoter and enhancer data (Hon *et al*., 2017), and (c) the standard gene models from GENCODE version 39. SACAGE contains 65% more gene models than GENCODE version 39 alone, including 48% derived from enhancers and 11% from promoters (Figure 2), allowing a uniquely comprehensive analysis of the RNA-DNA interactome.

**Figure 2:**
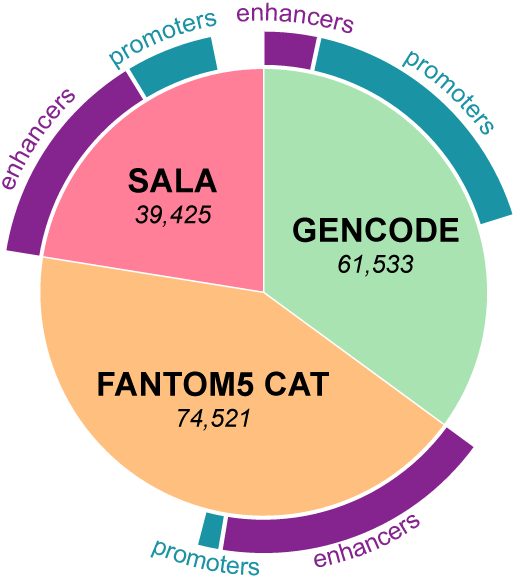
Distribution of the gene models present in the SACAGE annotation, in function of whether they originate from SALA-processed CFC-seq data, the FANTOM5 CAT annotation, or the GENCODE version 39 annotation. The proportion of regulatory elements (promoters or enhancers) among these models is indicated in purple and blue, respectively.

All the FANTOM6 data used in this study as well as the SACAGE annotation will be made available for interactive exploration on our Zenbu browser (Severin *et al*., 2014; Figure 3; https://fantom.gsc.riken.jp/zenbu/).

**Figure 3:**
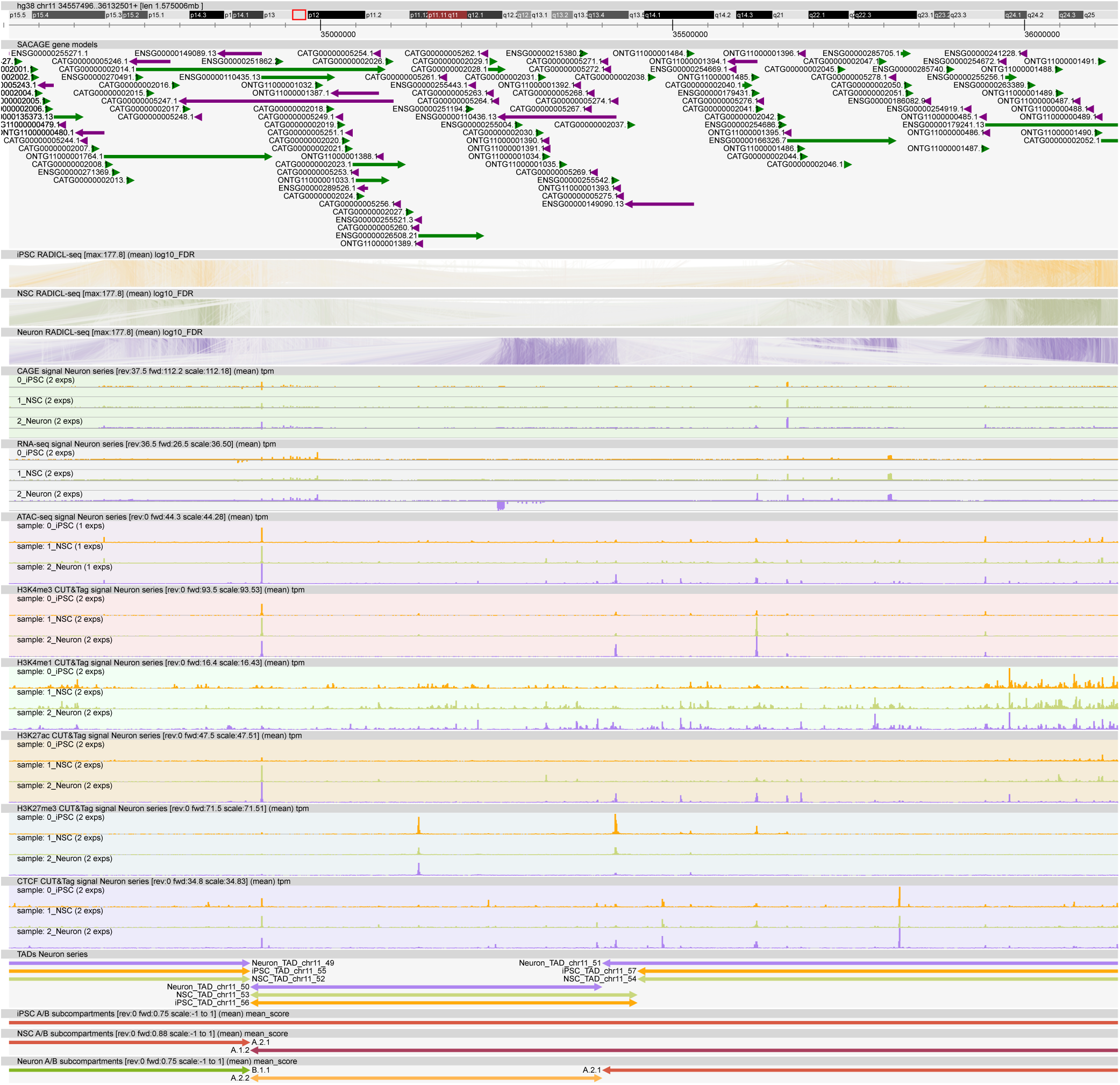
Screenshot of the Zenbu view dedicated to the FANTOM6 data collection, here showing the second half of the 11p13 region (chr11:34,557,496-36,132,501) in the three samples of the Neuron series.

### RNA-DNA interactions are cell type-specific, dynamic and diverse

After filtering out multimapped reads and non-significant interactions (Methods; Pracana *et al*., in preparation), the RADICL-seq data obtained from our 16 samples allows the investigation of human RNA-DNA interactions in specific cellular contexts. To globally compare the RNA-DNA contacts across our cell types, we calculated the Pearson correlation between all our RADICL-seq biological replicates based on their log-transformed interaction counts, then reduced it to two dimensions using multidimensional scaling (MDS; Figure 4, Methods). Not only did we observe a clear separation of the samples, but biologically similar cell types also consistently projected close to each other, with blood cells (THP-1 series, T cell series, AML and K562 samples) on one side and neural cells (NSC and Neuron samples) on the other side, separated by epithelial cells (MCF7-LTED series and HCT116 sample) in the center while the iPSC samples (iPSC and iPSCpreF6 samples) clustered at the top. As this broad biological grouping substantially agrees with an unsupervised clustering of our samples, significantly exceeding the null expectation (adjusted Rand index = 0.40 *versus* 0.11±0.07, *p*-value < 0.01; Methods), these results confirm that the RNA-DNA interaction landscape is reproducibly cell type-specific.

**Figure 4:**
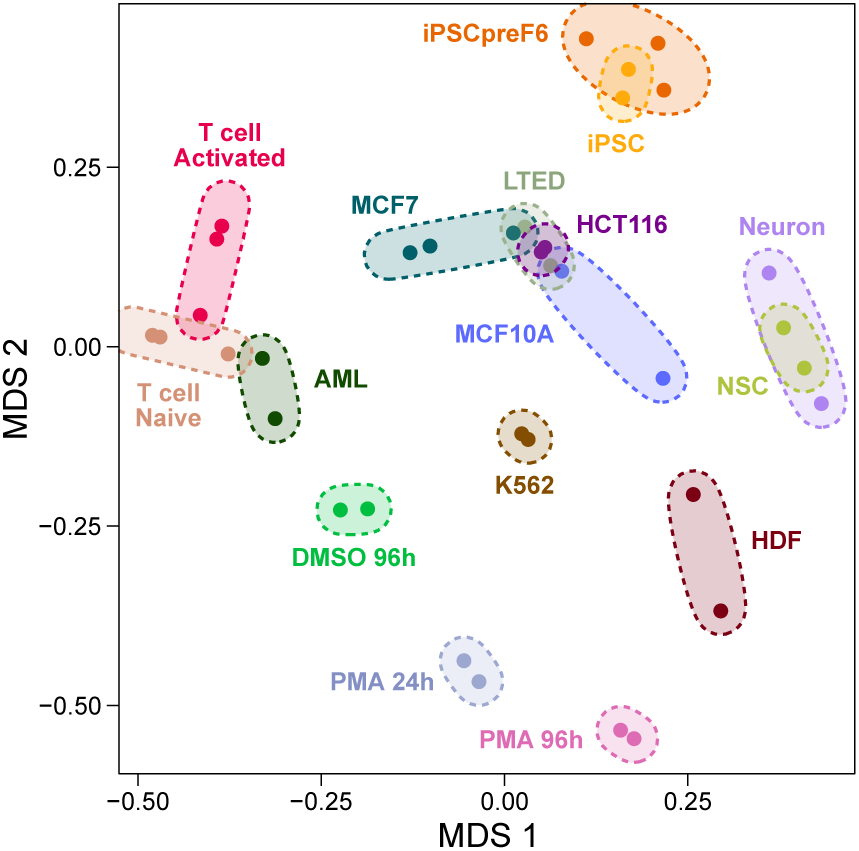
MDS plot of the Pearson correlation between all RADICL-seq replicates used in this study, calculated from the log-transformed CPMs of their transcript-isoform-to-25-kb-window interaction. For better visualization, points corresponding to the same sample are circled by colored ellipses.

In particular, when taken separately, most chromatin-interacting transcripts and RNA-targeted 25-kb genomic bins are shared by all or nearly all cell types analyzed in this study (Figure 5). However, most RNA/25-kb bin contacts are unique to each sample, indicating that the cell type-specificity of the RNA-DNA interactome is primarily defined by its source-target combinations, rather than by the overall set of chromatin-associated RNAs or of target DNA regions.

**Figure 5:**
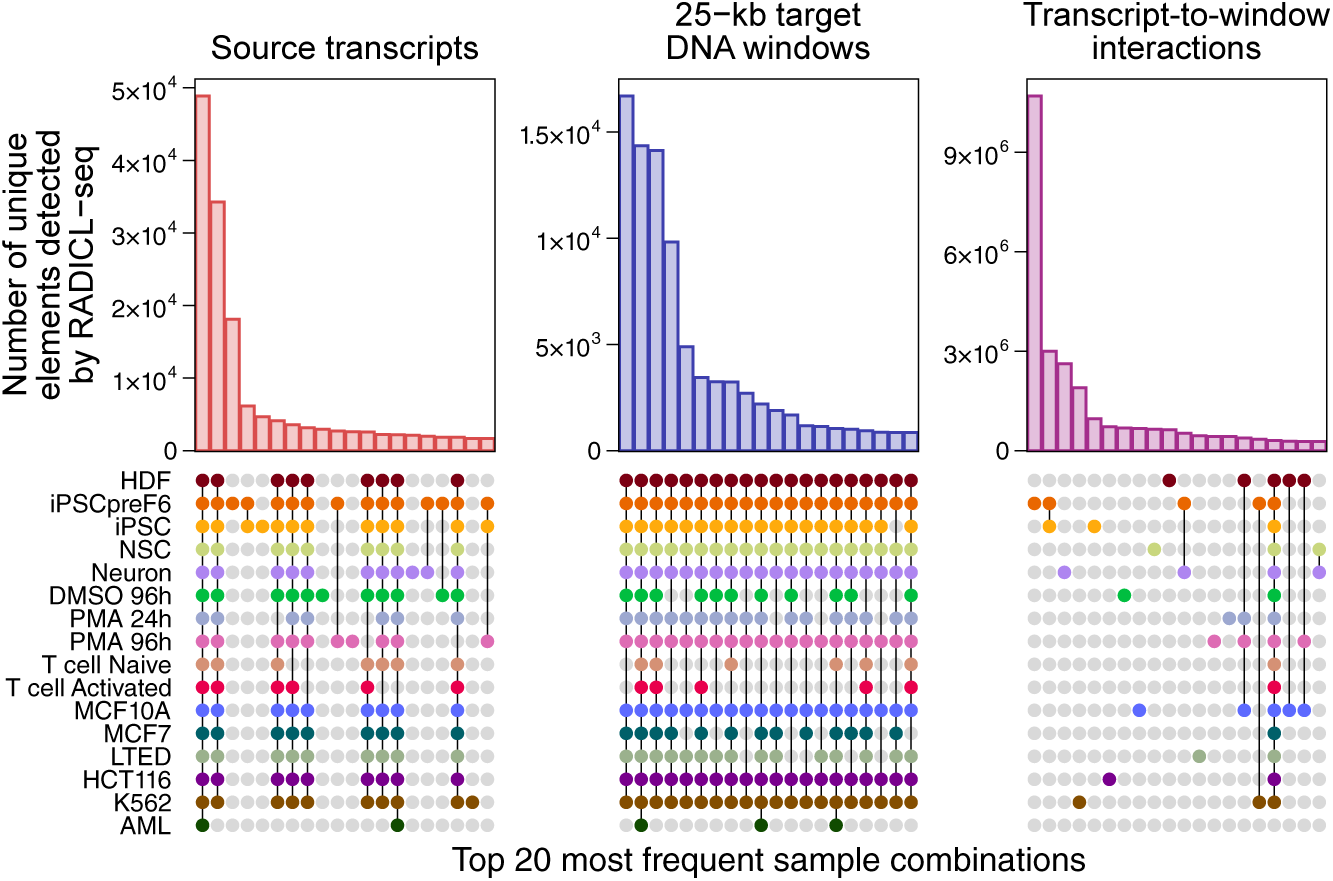
Distribution of unique DNA-interacting transcript isoforms, unique 25-kb target DNA windows and unique pairs of isoform/target window, in function of the samples in which they were detected by RADICL-seq after pooling biological replicates. For better visibility, only the top 20 most frequent sample combinations are shown.

To better understand the organization of this interaction landscape, we next modeled RNA-DNA contacts in the Neuron and THP-1 series as networks in which chromatin-bound RNAs and DNA-targeted sites are represented as nodes (Figure 6; Supplementary Table 4; Methods; Boguñá *et al*., 2021; Cannistraci *et al*., 2013; Cannistraci & Muscoloni, 2022; Muscoloni & Cannistraci, 2018; Muscoloni *et al*., 2017).

**Figure 6:**
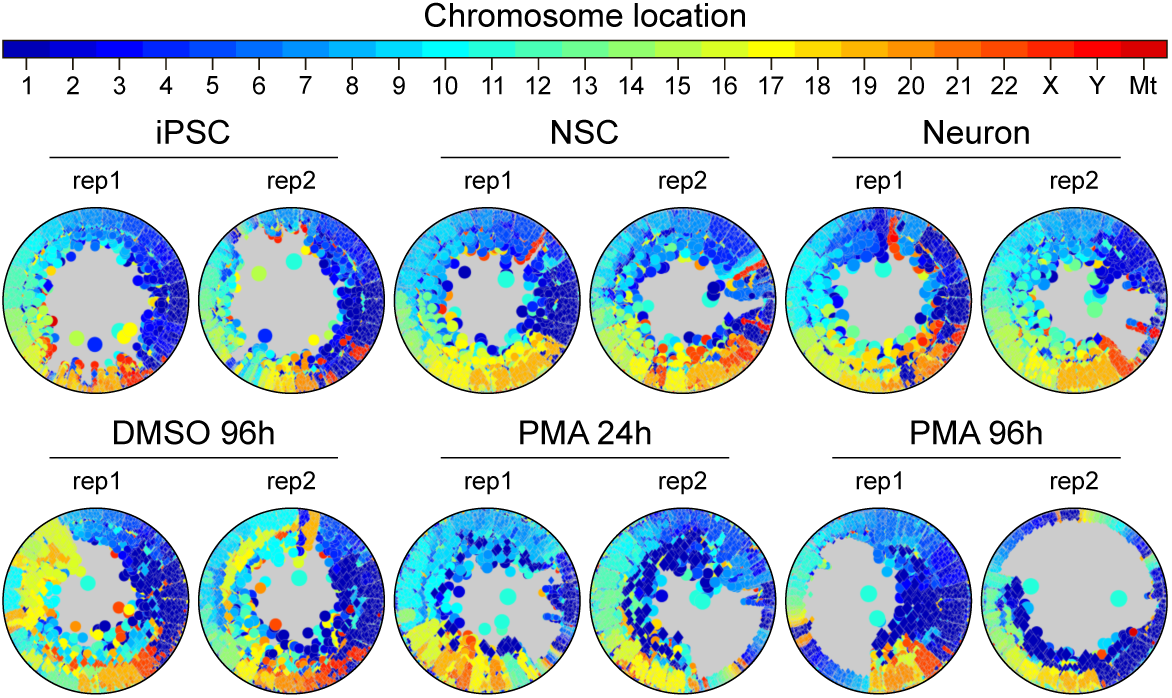
Hyperbolic network embedding representations in 2D disk spaces of the RNA-DNA interactome from each RADICL-seq replicate of the Neuron and THP-1 series. Source genes of interacting RNAs are shown as circles, and targeted 25-kb DNA windows are shown as diamonds. Both are colored in function of the chromosome on which they are located. Node sizes reflect their connection number in the entire network.

The networks obtained across all 6 samples are scale-free with (*γ*)—the power law exponent of their degree distribution)—lower than 3 (Figure 7; Supplementary Table 4; Voitalov *et al*., 2019), indicating that a small subset of nodes concentrates a large number of interactions, which is typical of biological networks with functional relevance (Almaas *et al*., 2007; Barabási & Albert, 1999). RNA nodes are significantly enriched among the most central nodes when embedding these networks within two-dimensional disks (Figures 6 and 8; Krioukov *et al*., 2010; Muscoloni *et al*., 2017; Y. Zhao *et al*., 2023; Methods), suggesting that RNAs play a greater role than their target DNA regions in governing the network architectures. In this two-dimensional representation, nodes originating from the same chromosome tend to cluster together (Figures 6 and 7; Supplementary Table 4), consistent with the influence of chromosome territories on the interactome first reported by X. Li *et al*. (2017). This chromosomal organization becomes more pronounced upon neuronal differentiation, as reflected by the higher angular separability index (ASI; Muscoloni & Cannistraci, 2019; Methods) in NSC compared to iPSC (Figure 7; Supplementary Table 4). The ultra-small-world connectivity of these networks ((*γ*)∼2 and characteristic path length (CPL)∼3; Figure 7; Supplementary Table 4; Methods) reveals their great efficiency in connecting different RNA and DNA nodes with each other (Boguñá *et al*., 2021; Boguñá & Krioukov, 2009; Cannistraci & Muscoloni, 2022). This connectivity notably increases during neuronal differentiation and macrophage activation, with a shorter CPL in NSC and Neuron than in iPSC and, although not significantly, in the two PMA samples compared to DMSO 96h (Figure 7; Supplementary Table 4).

**Figure 7:**
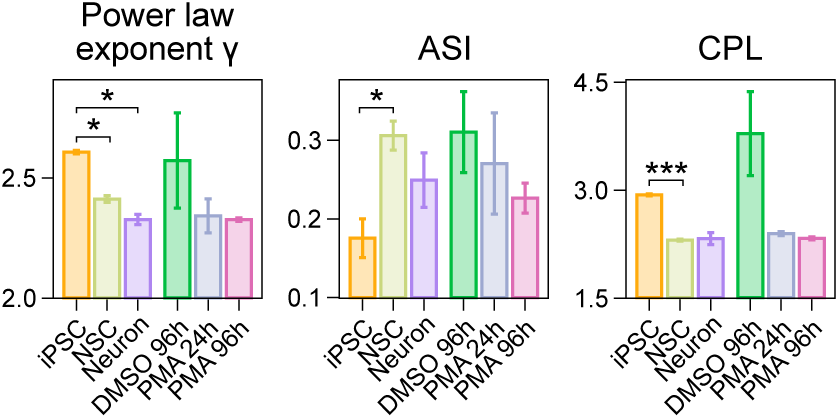
Power law exponent (γ) (left), angular separability index (ASI; middle) and characteristic path length (CPL; right) in RNA-DNA interaction networks created in each sample of the Neuron and THP-1 series (Figure 6), averaged across replicates for each sample. Error bars indicate the standard error of the mean for each type of measure. Stars indicate significant differences samples (Student’s t-test; *p value < 0.05; ***p value < 0.001). ASI values are all significantly positive (p value < 0.05).

**Figure 8:**
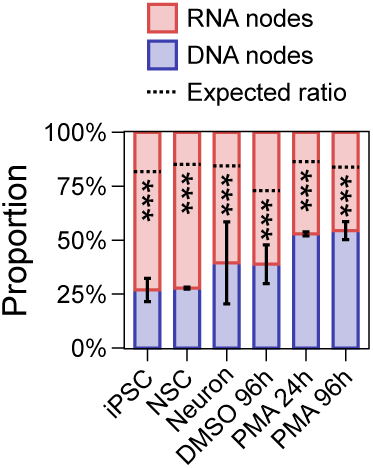
Proportion of RNA and DNA nodes among the 5,000^th^ nodes with the lowest radial coordinates in the RNA-DNA interaction networks created in each sample of the Neuron and THP-1 series (Figure 6). Error bars indicate the standard deviation between the two replicates. Black dotted lines indicate the expected RNA/DNA node ratio, as determined by random permutation test, and stars indicate significant differences between this expectation and the observation (***p value < 0.001).

Altogether, this analysis demonstrates that the RNA-DNA interactome is not random but hierarchically organized by a subset of central RNAs, further shaped by chromosome territories and subjected to a global reconfiguration during cell specialization.

The influence of chromosome territories on the RNA-DNA interactome suggests that the genomic distance between source RNAs and their targets conditions their interaction. The extent of this conditioning appears to vary between chromatin-bound transcripts, as observed when clustering all RNAs detected in our 16 samples based on the span of their contacting region relative to their source locus (Figure 9). We detected a group of relatively long, highly expressed and highly interacting transcripts that primarily interact with regions very close to their transcription site, including a large proportion of self interactions (clusters 1 and 2; Figures 9 to 13). Conversely, the second-biggest group of RNAs disclosed by this approach consists of shorter, less expressed and, overall, less frequently interacting transcripts that mainly interact with more distal regions (cluster 5; Figures 9 to 13).

**Figure 9:**
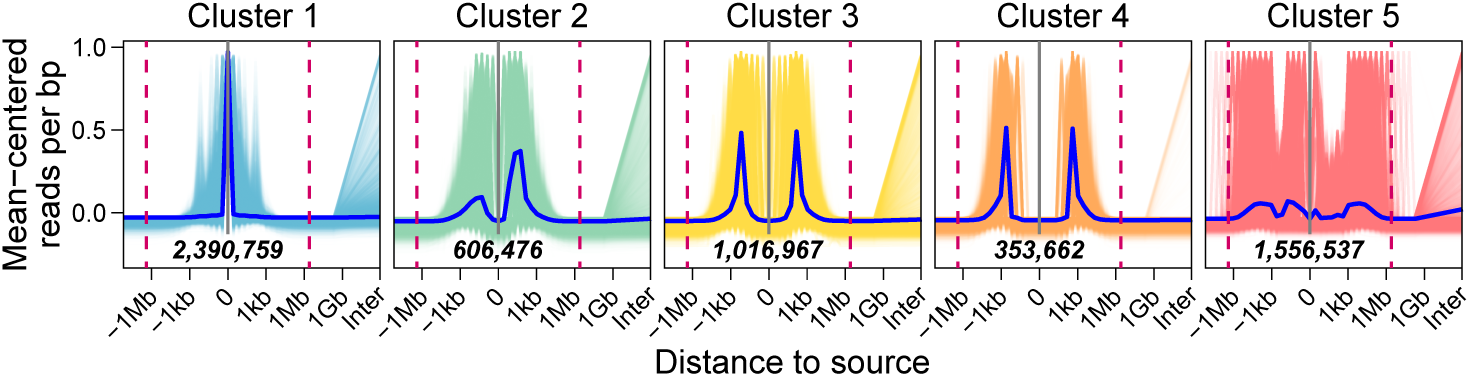
RNA-DNA interaction landscapes around the RNA-producing locus in all RADICL-seq replicates of the FANTOM6 collection. Each line shows the mean-centered density of interaction across the genome of a given transcript isoform in a given replicate. K-means clustering was used to group all landscapes in function of their shape; bold-italic numbers indicate the number of transcript-isoforms-in-replicate within each cluster, while the dark blue lines display the average interaction density. The purple dashed lines correspond to a source-target distance of ±2.5 Mb.

**Figure 10:**
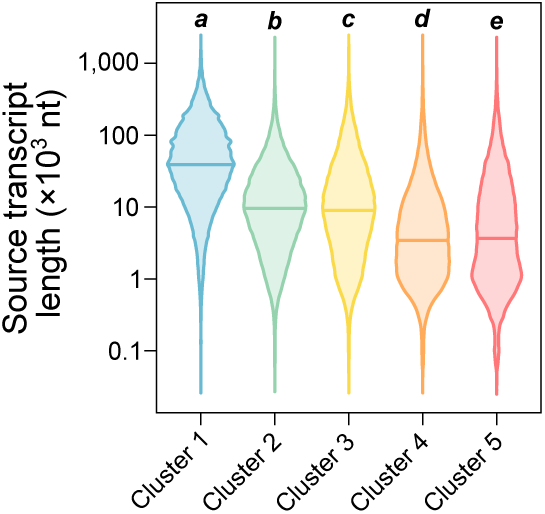
Distribution of the length of the coding region (introns included) for all DNA-interacting transcript isoforms detected in each RADICL-seq replicate, separated in function of the clusters defined in Figure 9. The bold italic letters indicate significantly different groups of clusters, defined by one-way ANOVA with post-hoc Tukey test.

**Figure 11:**
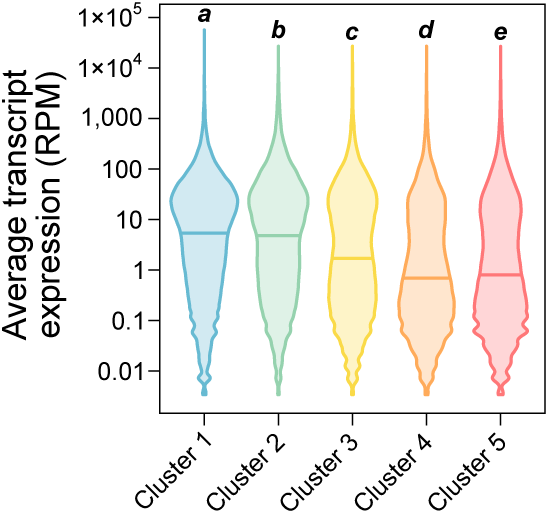
Distribution of the average CAGE reads coverage (RPM) on the source locus of all DNA-interacting transcript isoforms detected in each RADICL-seq replicate, separated in function of the clusters defined in Figure 9. The bold italic letters indicate significantly different groups of clusters, defined by one-way ANOVA with post-hoc Tukey test.

**Figure 12:**
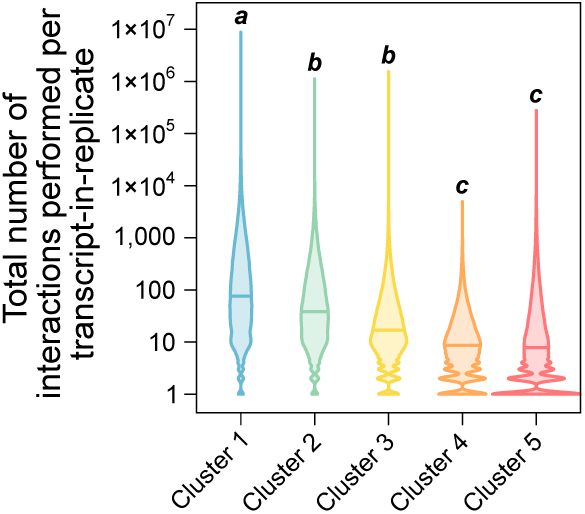
Distribution of the total number of interactions performed by all DNA-interacting transcript isoforms detected in each RADICL-seq replicate, separated in function of the clusters defined in Figure 9. The bold italic letters indicate significantly different groups of clusters, defined by one-way ANOVA with post-hoc Tukey test.

**Figure 13:**
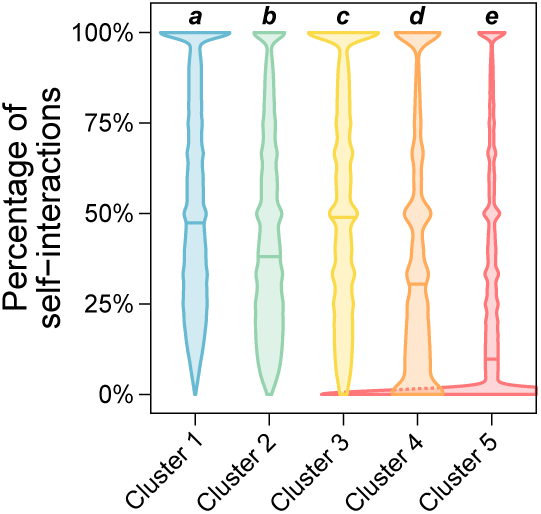
Distribution of the percentage of self interactions performed by all DNA-interacting transcript isoforms detected in each RADICL-seq replicate, separated in function of the clusters defined in Figure 9. The bold italic letters indicate significantly different groups of clusters, defined by one-way ANOVA with post-hoc Tukey test. For better visibility, the violin corresponding to Cluster 5 is cropped on the right side.

Comparing our RADICL-seq data with Hi-C in the Neuron and THP-1 series reveals that, on the one hand, RNAs predominantly interact with DNA targets located within the same topologically associated domain (TAD) as their source locus (Figures 14 and 15), consistent with previous reports (Bonetti *et al*., 2020; Calandrelli *et al*., 2023; X. Li *et al*., 2017; Limouse *et al*., 2023). On the other hand, RNAs that escape this general rule and perform the most inter-TAD intrachromosomal interactions tend to cluster with those that perform the most interchromosomal interactions (cluster 5; Figures 9 and 16). Therefore, we henceforth consider that any intrachromosomal RNA-DNA contact spanning a distance greater than 2.5 Mb, *i.e.* a reasonable capping size for TADs (Figure 17; J. Xu *et al*., 2024), is equivalent to an inter-chromosomal RNA-DNA contact. Both are hereinafter referred to as *trans* interactions and, accordingly, we call *cis* interaction any intrachromosomal RNA-DNA contact occurring within 2.5 Mb from the RNA’s source locus.

**Figure 14:**
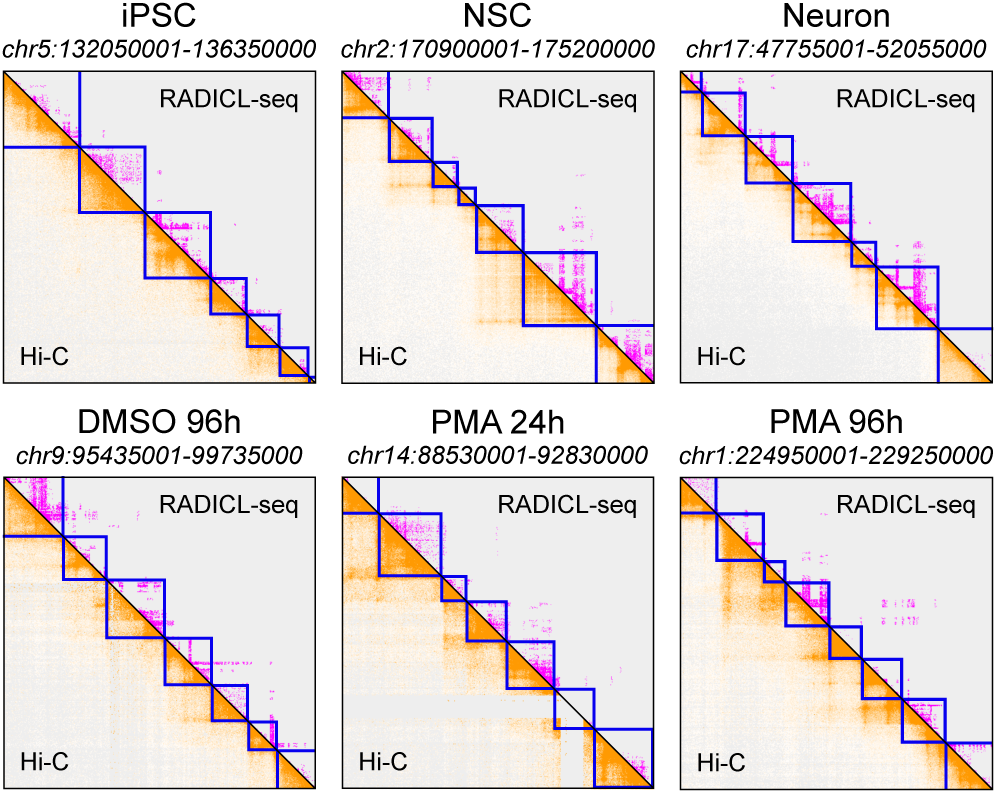
Juicebox screenshots showing DNA-DNA (bottom left, orange) and RNA-DNA (top right, pink) contact matrices, along with TADs detected by Hi-C (blue squares), in each sample of the Neuron and THP-1 series. For RADICL-seq data, biological replicates were pooled together. A different representative region is shown for each sample.

**Figure 15:**
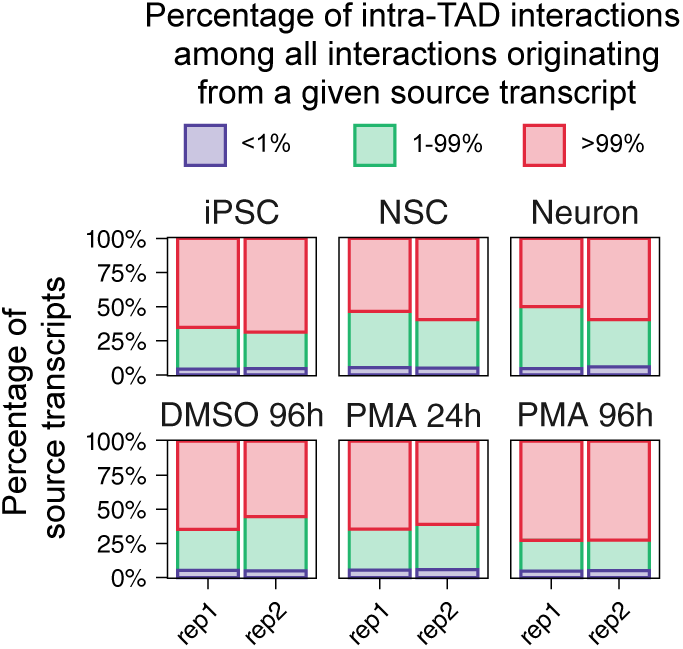
Proportion of source transcript isoforms detected by RADICL-seq in function of whether they perform <1%, >99% or an intermediate percentage of their RNA-DNA interactions within the same TAD as their coding locus, in each replicate of the Neuron and THP-1 series.

**Figure 16:**
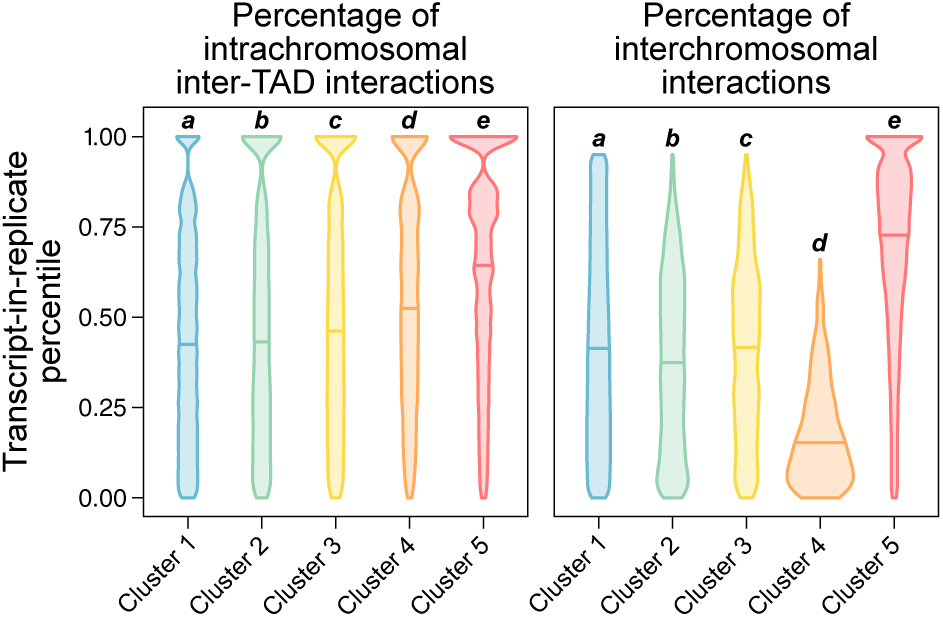
Distribution of the percentiles to which all DNA-interacting transcript isoforms belong in function of the percentage of intrachromosomal interactions that they perform beyond the TAD of their source locus (left) or the percentage of all interactions that they perform on another chromosome than that of their source locus (right). In the left panel, transcripts-in-replicate performing all their intrachromosomal interactions within their source TADs were excluded, and only data from RADICL-seq replicates of the Neuron and THP-1 series (i.e. samples with corresponding Hi-C data) are shown. In the right panel, transcripts-in-replicate performing only intrachromosomal interactions were excluded, and data from all RADICL-seq replicates in the FANTOM6 collection are shown. For both panels, transcripts-in-replicate are separated in function of the clusters defined in Figure 9, and the bold italic letters indicate significantly different groups of clusters, defined by one-way ANOVA with post-hoc Tukey test.

**Figure 17:**
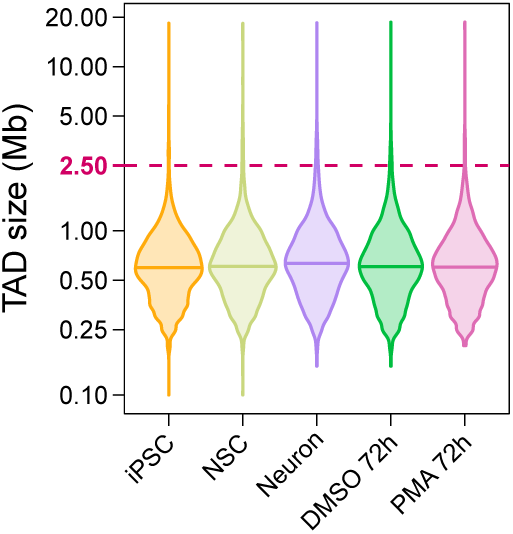
Size distribution of all TADs detected by Hi-C in each sample of the Neuron and THP-1 series.

Using these definitions, we found that *trans* contacts represent 42.67% of all interactions detected across all of our samples, with considerable variations between cell types (from 14.69% in T cell Activated to 78.58% in PMA 96h; Figure 18). Interestingly, we detected a clear rise in the share of *trans* contacts in THP-1 cells with the increasing duration of PMA exposure (DMSO 96h to PMA 96h) as well as during neuronal differentiation (iPSC to NSC to Neuron) and, in the opposite direction, during HDF reprogramming into iPSC (HDF to iPSCpreF6), suggesting that long-distance RNA-DNA interactions play a key role in cell specialization.

**Figure 18:**
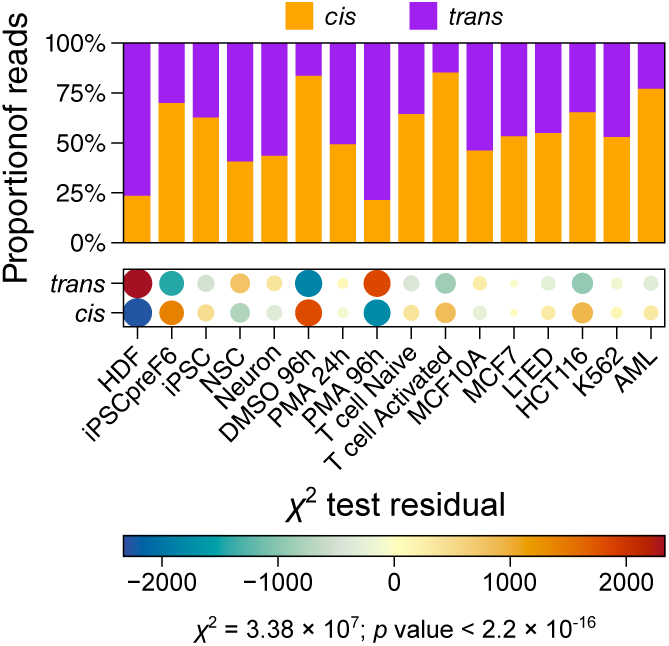
Percentage of reads corresponding to either cis (≤2.5 Mb between source and target loci) or trans (>2.5 Mb between source and target loci) RNA-DNA interactions in each RADICL-seq sample used in this study, after pooling replicates. The enrichment (positive residual) or depletion (negative residual) of each type of interaction in each sample, as determined by χ^2^ test, is indicated in the bottom balloon plot.

*MALAT1*-mediated contacts account for most *trans* interactions in all cell types (Figures 19 and 20). We also detected a significant contribution from *NEAT1* in several samples, particularly in HDF, in the THP-1 series and in MCF10A (Figures 19 and 20). Aside from these two lncRNAs, long-distance contacts are performed by diverse types of RNAs, and their contribution varies across cell types (Figure 21). For example, transcripts from snoRNA host genes are important sources of *trans* interactions in the iPSC and iPSCpreF6 samples, which is consistent with previous observations in embryonic cells (Limouse *et al*., 2023; Sridhar *et al*., 2017), but are less prevalent in the other samples (Figure 21). Similarly, *trans*-interacting RNAs derived from introns of protein-coding genes are particularly frequent in HDF and in Neuron compared to the other cell types (Figure 21). These observations thus highlight the value of surveying various cellular contexts to fully characterize the diversity of the RNA-DNA interaction landscape. By contrast, sources of short-distance contacts are remarkably uniform across samples (Figure 19).

**Figure 19:**
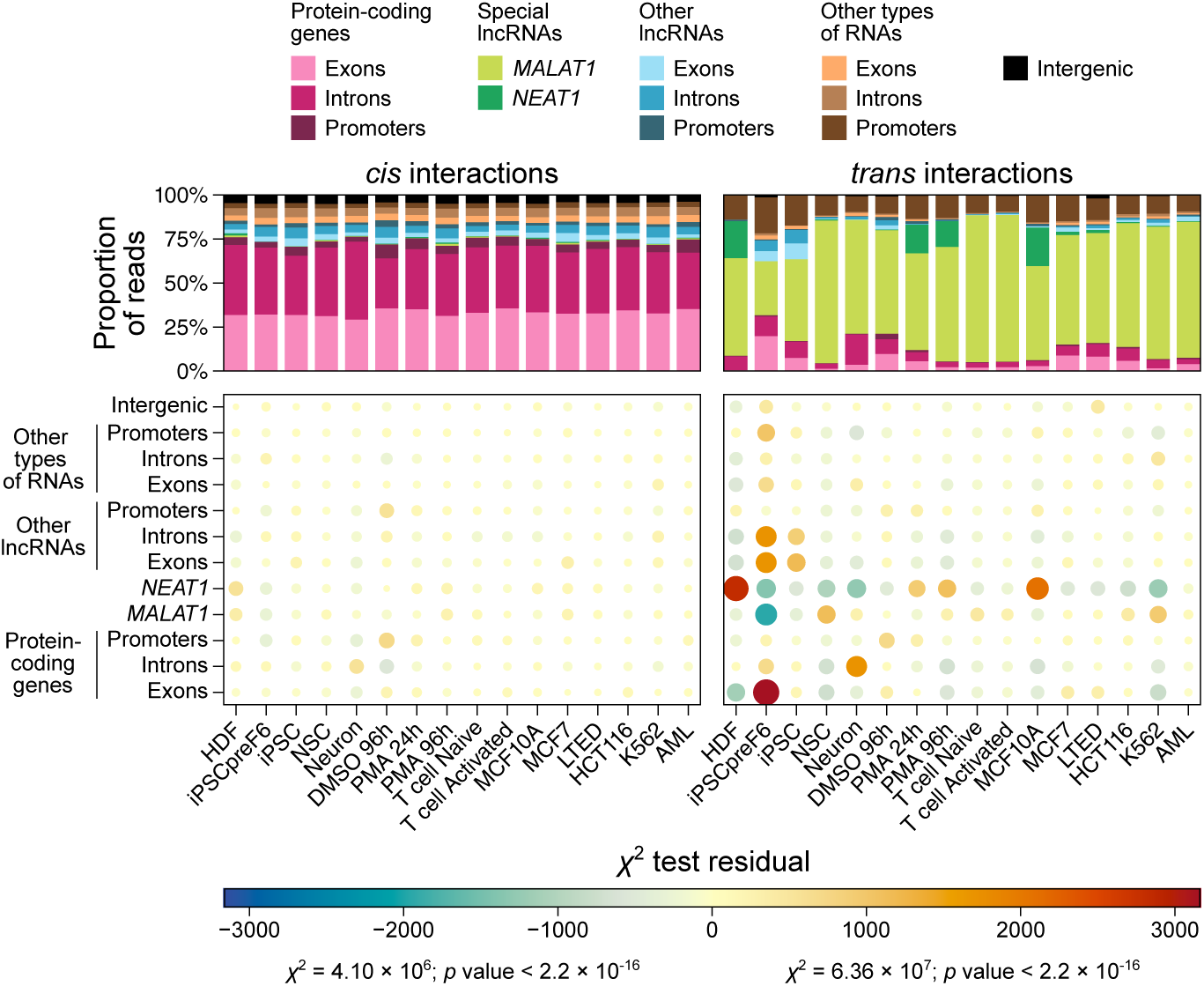
Percentage of reads relative to the type of locus to which the RNA fragment maps, in each RADICL-seq sample used in this study, after pooling replicates. Reads are separated in function of whether they correspond to a cis or trans RNA-DNA interaction. The enrichment (positive residual) or depletion (negative residual) of each type of source RNA for each type of interaction in each sample, as determined by χ^2^ test, is indicated in the bottom balloon plot.

**Figure 20:**
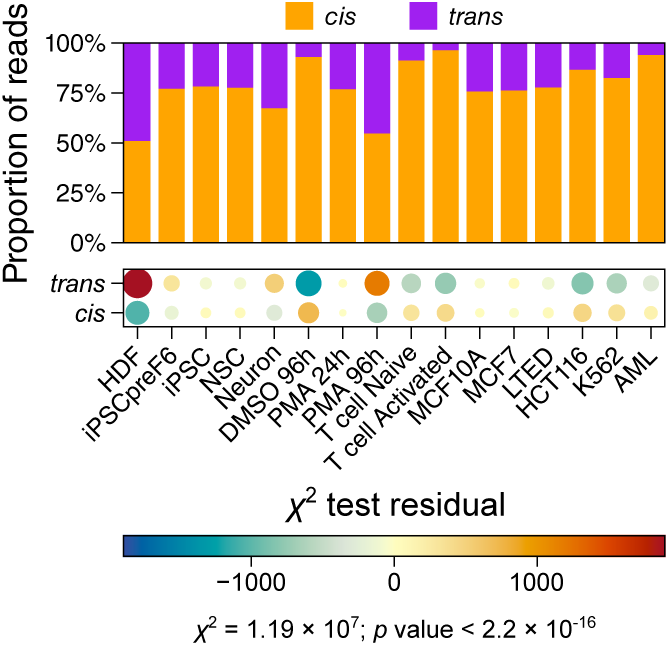
Percentage of reads corresponding to either cis (≤2.5 Mb between source and target loci) or trans (>2.5 Mb between source and target loci) RNA-DNA interactions in each RADICL-seq sample used in this study, after pooling replicates and removing all reads for which the RNA side maps to either MALAT1 or NEAT1. The enrichment (positive residual) or depletion (negative residual) of each type of interaction in each sample, as determined by χ^2^ test, is indicated in the bottom balloon plot.

**Figure 21:**
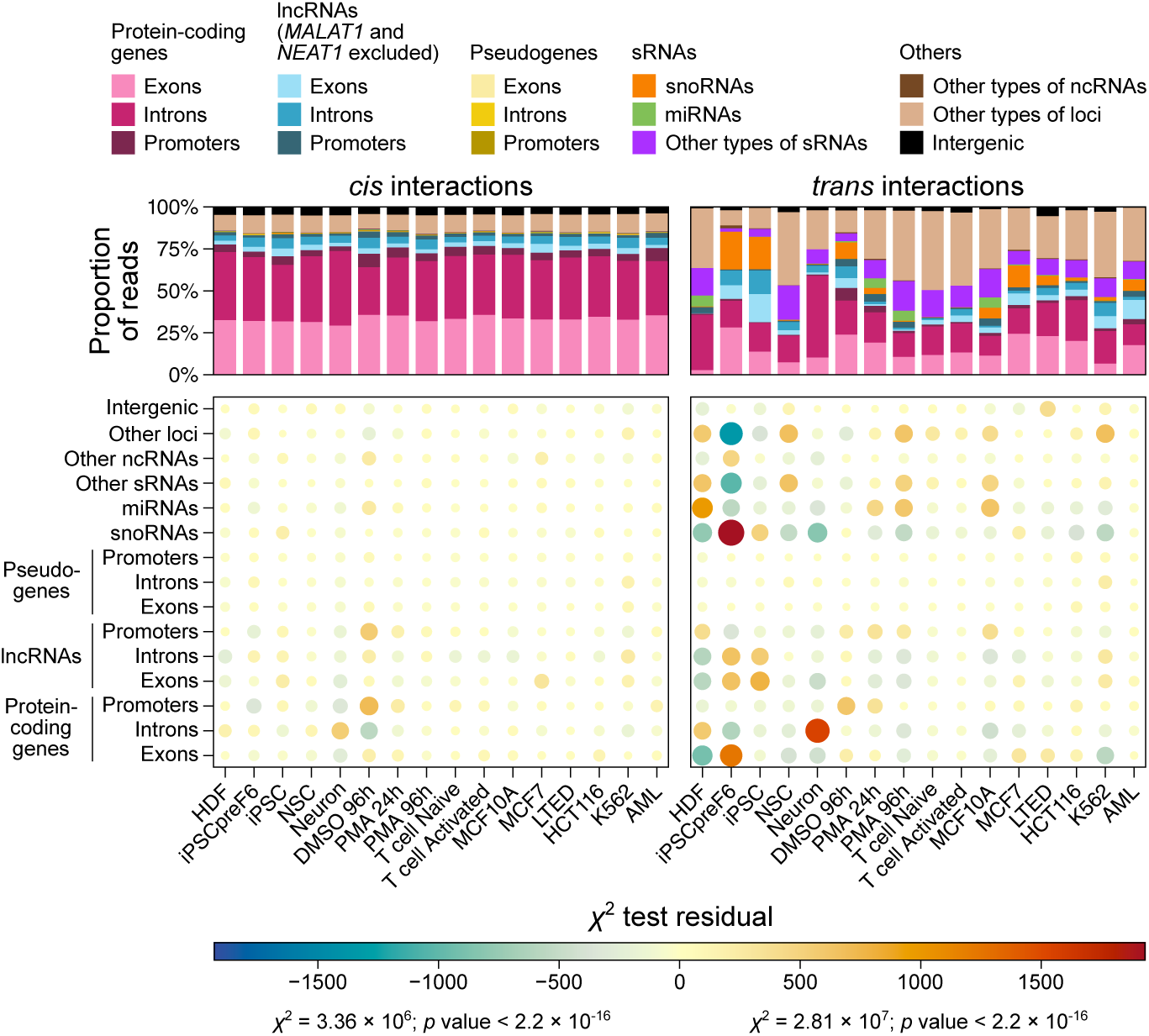
Percentage of reads relative to the type of locus to which the RNA fragment maps, in each RADICL-seq sample used in this study, after pooling replicates and removing all reads for which the RNA side maps to either MALAT1 or NEAT1. Reads are separated in function of whether they correspond to a cis or trans RNA-DNA interaction. The enrichment (positive residual) or depletion (negative residual) of each type of source RNA for each type of interaction in each sample, as determined by χ^2^ test, is indicated in the bottom balloon plot.

Nearly half of *cis* interactions involve RNAs derived from introns of protein-coding genes, and an additional 25% are performed by RNAs derived from their exonic counterparts, possibly reflecting contacts performed by nascent RNAs. Surprisingly, non-*MALAT1* or *NEAT1* lncRNAs generally represent a relatively small part of all chromatin-associated RNAs detected in our RADICL-seq collection, and they make up a higher percentage of sources of *trans* interactions than of *cis* interactions (Figure 21).

Overall, our results pertaining to the distance distribution of RNA-DNA contacts and to the nature of the RNAs performing these interactions are consistent with previous findings obtained by RADICL-seq and related technologies (Bell *et al*., 2018; Bonetti *et al*., 2020; Gavrilov *et al*., 2020; Jayne *et al*., 2024; L. Li *et al*., 2021; X. Li *et al*., 2017; Limouse *et al*., 2023; Tenorio *et al*., 2023; Yun *et al*., 2024), supporting the accuracy and reliability of our data collection. Furthermore, the fact that these concordant observations have been reported across different mammalian cell types (Bonetti *et al*., 2020; Jayne *et al*., 2024; X. Li *et al*., 2017; Limouse *et al*., 2023; Yun *et al*., 2024) and even different species (L. Li *et al*., 2021; X. Li *et al*., 2017; Tenorio *et al*., 2023) suggests that these features of the RNA-DNA interactome are broadly conserved.

### Parameters influencing RNA-DNA interactions at their source and target loci

RADICL-seq read counts may be affected by local variations in sequenceable DNA due to copy number variations (CNVs) and early DNA replication timing (RT), which are cell type-specific features (Koren *et al*., 2014, 2021). Consistently, significant interaction counts correlate with CNVs and RT in the highly proliferative K562 cell line (Figure 22), and a Support Vector Machine (SVM)-based model using CNVs and RT as predictive features can accurately and reproducibly predict K562 and MCF7 RADICL-seq counts, with RT being a more influential factor than CNVs (Figures 23 and 24). RT may thus shape the RNA-DNA interaction counts by locally altering the amount of recoverable DNA, especially in fast-cycling cancerous cells. Alternatively, RNA-DNA interactions may also play a role in DNA replication and the two phenomena may be intertwined, as suggested by previous studies (Ge & Lin, 2014; Heskett *et al*., 2022; Thayer *et al*., 2024). Given this possibility and the lack of available CNV data for several of our samples, we therefore chose not to correct for these factors in our further analyses, but caution against their influence in the interpretation of our data.

**Figure 22:**
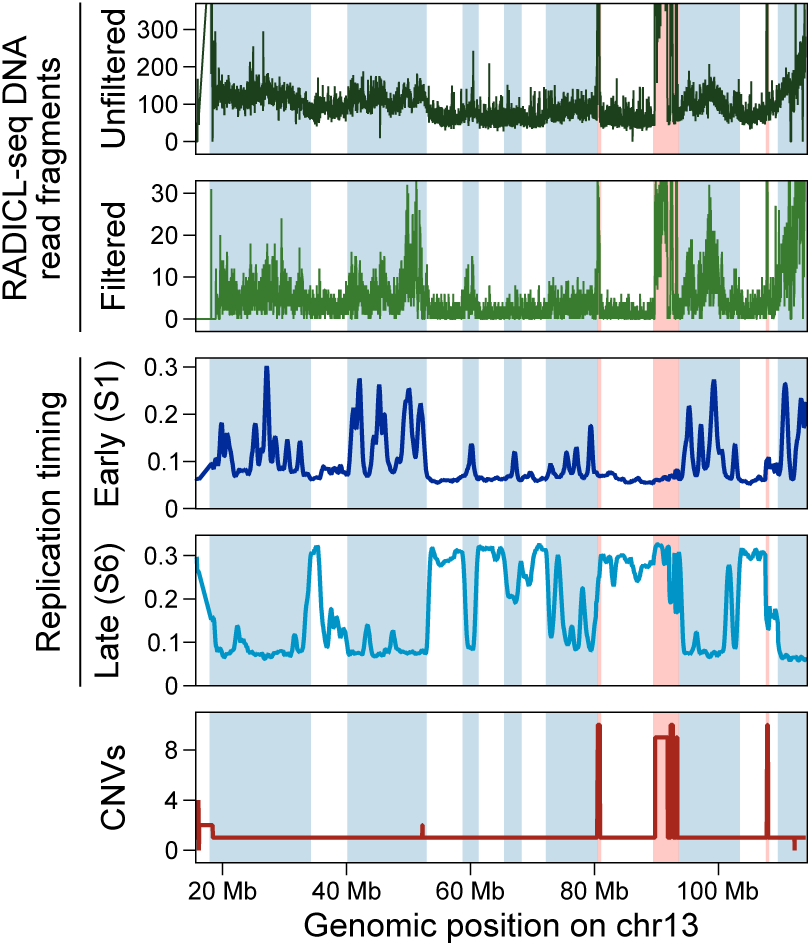
Correlation between raw and filtered RADICL-seq DNA signal, replication timing (RT) and copy number variations (CNVs) in chromosome 13 of K562 cells. RADICL-seq DNA read fragments (excluding those corresponding to self interactions) were counted by 1-kb bin. For RT, only two S phase fractions (S1 as early-replicating and S6 as late-replicating) are shown. For all tracks and for better clarity, only points corresponding to 5000 randomly selected bins in the chr13:20,000,000-110,000,000 region are plotted. Light blue and light red areas indicate DNA regions where RADICL-seq counts correlate with RT and CNVs, respectively.

**Figure 23:**
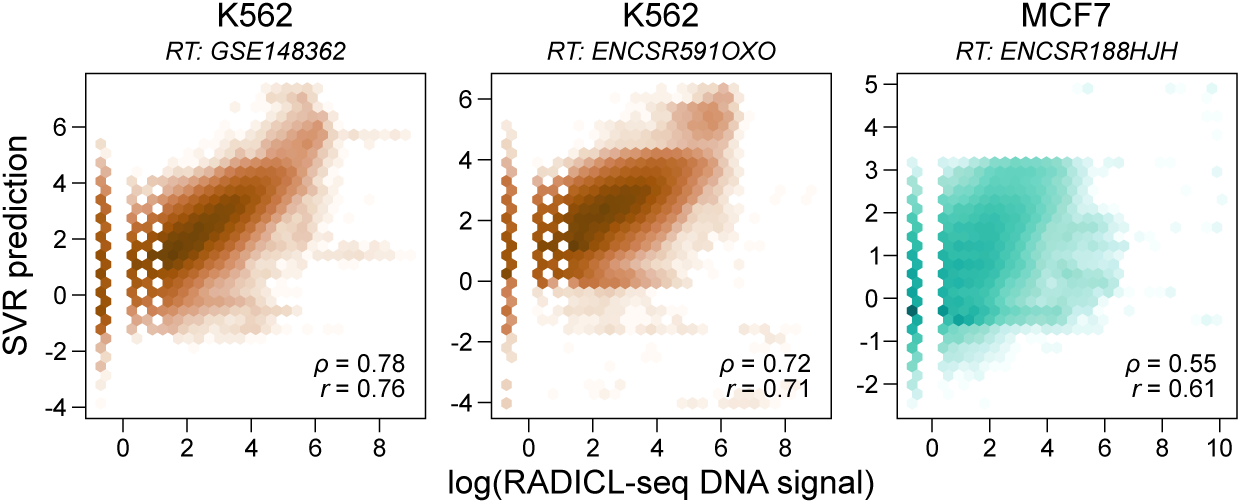
Prediction of log-transformed RADICL-seq DNA read fragment counts per 1-kb bin by a support vector machine regression (SVR) from RT and CNV data, in function of the real RADICL-seq signal. The analysis was performed using two different RT datasets in K562 cells (GSE148362, 6 S phases, and ENCSR591OXO, 4 S phases) and one RT dataset in MCF7 cells (ENCSR188HJH, 4 S phases). Spearman’s (ρ) and Pearson’s (r) correlation coefficients are indicated in the bottom right corner of each panel. These coefficients are lower in MCF7 cells due to the ENCSR188HJH dataset having only one replicate (36 nt), unlike the ENCSR591OXO dataset in K562 (36 and 49 nt).

**Figure 24:**
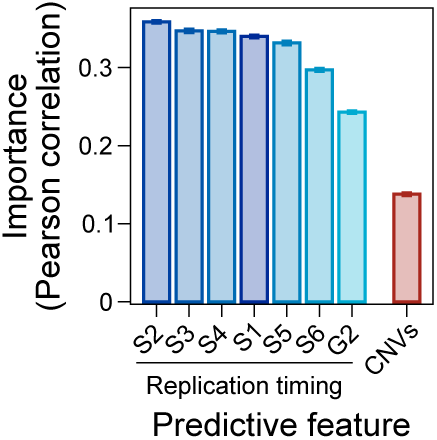
Mean feature importance of RT and CNV parameters used as predictive values for the SVR trained on K562 cells data with the GSE148362 RT dataset described in Figure 23. Permutation feature importance was evaluated on the model’s Pearson correlation coefficient. Error bars indicate the standard deviation between 50 iterations of permutation.

Another factor that could impact RNA-DNA interaction patterns is the presence of repeat elements. More than half of the human genome is repetitive (Hoyt *et al*., 2022) and repeat elements such as short interspersed nuclear elements (SINEs), long interspersed nuclear elements (LINEs), endogenous retrovirus/long terminal repeats (ERV/LTRs), DNA transposons, microsatellites and low-complexity regions (LCRs) are particularly present in a large proportion of protein-coding transcripts and lncRNAs (Figure 25). Moreover, many repeats show cell-specific transcription levels (Faulkner *et al*., 2009). Given this abundance, we next assessed whether these repeats may influence the RNA-DNA interaction landscape. When looking at the amount of RADICL-seq RNA read fragments encompassing repeat elements, we observed different profiles depending on the class of the repeat, independently of the cell type (Figures 26 and 27): while transcripts derived from LINEs or DNA transposons do not seem to interact differently with the chromatin than other randomly chosen RNAs, transcripts containing ERV/LTRs are globally depleted from RNAs detected by RADICL-seq. As this contradicts other studies reporting an enrichment in these elements among chromatin-associated transcripts (Bonetti *et al*., 2020; Limouse *et al*., 2023; Panariello *et al*., in preparation; W. Xu *et al*., 2021), it is possible that this depletion was artificially induced by our choice to filter out multimapped reads when processing RADICL-seq data (Methods; Pracana *et al*., in preparation). Likewise, this technical artefact could explain the drop in RADICL-seq RNA signal on SINEs compared to their flanking regions, which also occurs when looking at the RADICL-seq DNA signal (Figures 26 and 27). Conversely, the regions surrounding the start of LCRs and, to a lesser extent, the regions upstream of microsatellites tend to produce more DNA-interacting RNA fragments than expected by chance, a pattern that is not reproduced by the RADICL-seq DNA signal (Figures 26 and 27). The microsatellites and/or LCRs within a transcript may thus promote its ability to associate with the chromatin. However, this parameter does not appear to influence the distance of this association, as RNAs containing microsatellites or LCRs contact the DNA in *trans* virtually as often as RNAs lacking any repeat element (Figure 28). On the other hand, RNAs containing SINEs, ERV/LTRs and DNA transposons tend to interact with the chromatin primarily in *cis* (Figure 28), consistent with prior observations in mice (Bonetti *et al*., 2020). On the DNA side, repeat elements appear to be targeted essentially as frequently as any other loci (Figures 26 and 27) and the disparity in *cis*/*trans* ratio for interactions targeting repeated *versus* non-repeated regions is relatively minimal, with repeat elements being slightly more often targeted in *trans* only in some samples (PMA 24h, MCF7, HCT116 and K562; Figure 28). Together, these results strengthen previous conclusions (Bonetti *et al*., 2020) that certain types of repeated sequences within a transcript appear to affect either its frequency of interactions with the DNA or with the distance at which these interactions occur, whereas the presence of repeats in the DNA is unlikely to be a major determinant of RNA-targeted sites.

**Figure 25:**
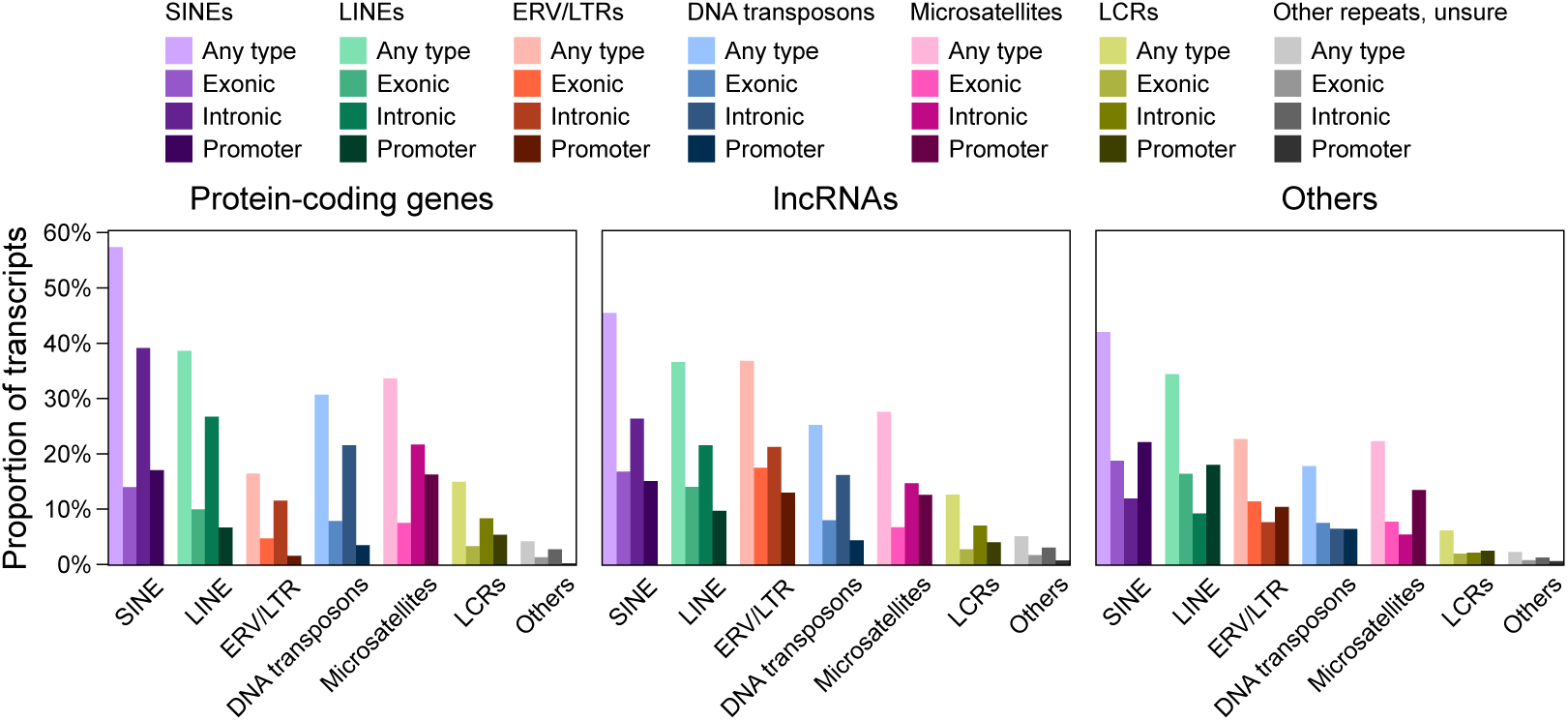
Percentage of SACAGE transcripts models that overlap at least one short interspersed nuclear element (SINE), long interspersed nuclear element (LINE), endogenous retrovirus/long terminal repeat (ERV/LTR), DNA transposon, microsatellite or low-complexity region (LCR). Transcripts are separated in function of whether they are derived from protein-coding genes, lncRNAs or other types of loci, and repeat elements are split in function of whether they are located within the promoter, exonic or intronic region of their overlapping transcript.

**Figure 26:**
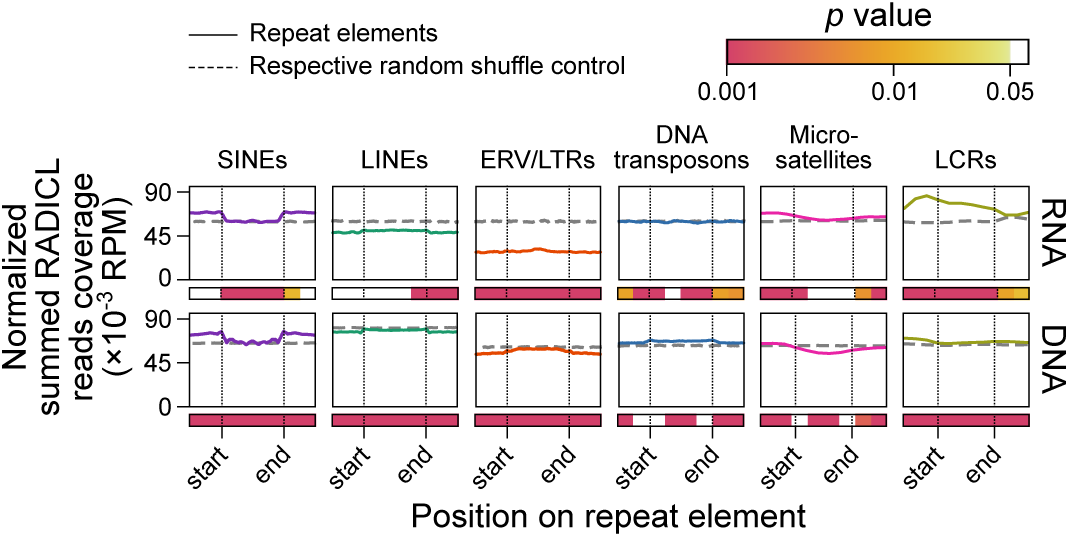
Distribution of RADICL-seq RNA (top) and DNA read fragments (bottom), pooled across all replicates used in this study, over different classes of repeat elements and their ± 50% surrounding region. All values for all repeat elements of a given class were summed together, after normalizing position values to the total length of each element. As a control, the RNA and DNA fragment coverage on equivalent, randomly shuffled sets of regions for each class of repeat elements is shown as grey dotted lines. The significance (p value, permutation test) of the difference between the real and control coverage per 25% bin is indicated by colored squares below each panel.

**Figure 27:**
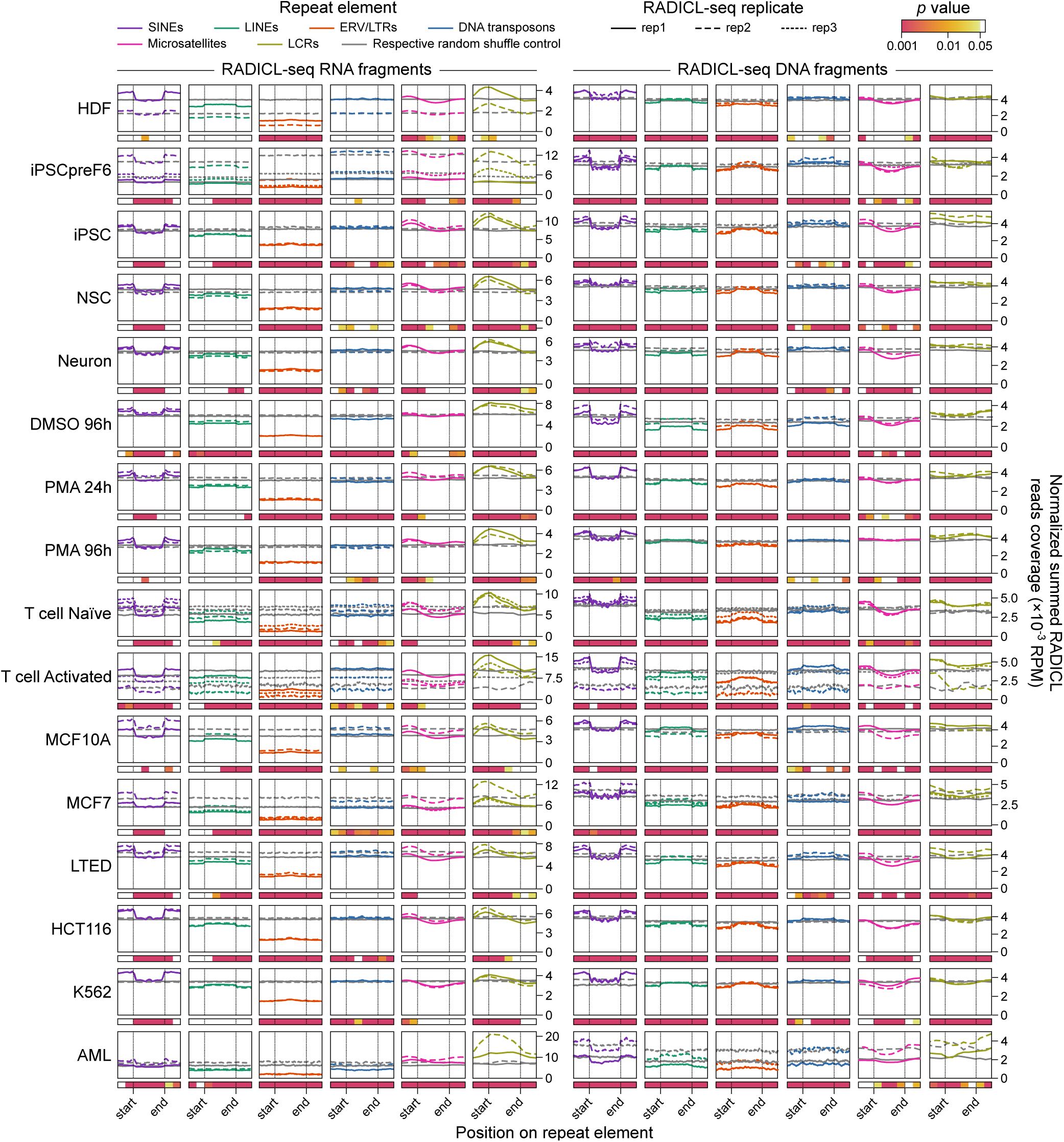
Distribution of RADICL-seq RNA (left) and DNA read fragments (right) over different classes of repeat elements and their ± 50% surrounding region, in each RADICL-seq replicate used in this study. Values for all repeat elements of a given class were summed together, after normalizing position values to the total length of each element. As a control, the RNA and DNA fragment coverage on equivalent, randomly shuffled sets of regions for each class of repeat elements is shown as grey dotted lines. The significance (p value, permutation test) of the difference between the real and control coverage per 25% bin is indicated by colored squares below each panel.

**Figure 28:**
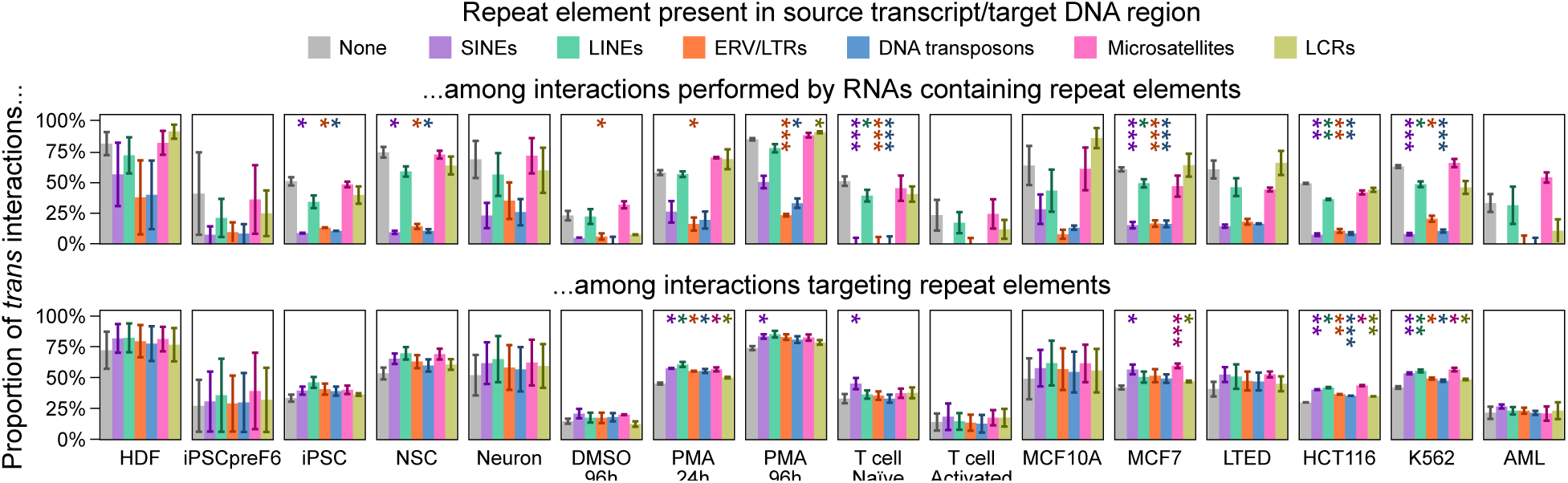
Percentage of trans interactions among all interactions performed by transcript isoforms containing either a SINE, LINE, ERV/LTR, DNA transposon, microsatellite, LCR, or no repeat element (top), and among all interactions targeting DNA regions containing either or none of those repeats (bottom), in each RADICL-seq sample used in this study. Error bars indicate the standard deviation between biological replicates. Stars indicate significant differences between transcripts or target regions containing a given type of repeat element versus those that do not contain any (Student’s t-test; *p value < 0.05; **p value < 0.01; ***p value < 0.001).

We next assessed whether other parameters than their sequence influence which loci are sources or targets of chromatin-associated RNAs. To this end, we first examined the distribution of RADICL-seq reads along the length of all chromosomes (Figure 29). We found that the extremities of chromosomes tend to have a higher coverage in both RNA and DNA fragments compared to their arms and centromeres, especially in cancerous and/or immune cell types (THP-1 and T cell series, LTED, HCT116 and K562 samples), suggesting that telomeres and pre-telomeric regions are prone to intense RNA-DNA interactions.

**Figure 29:**
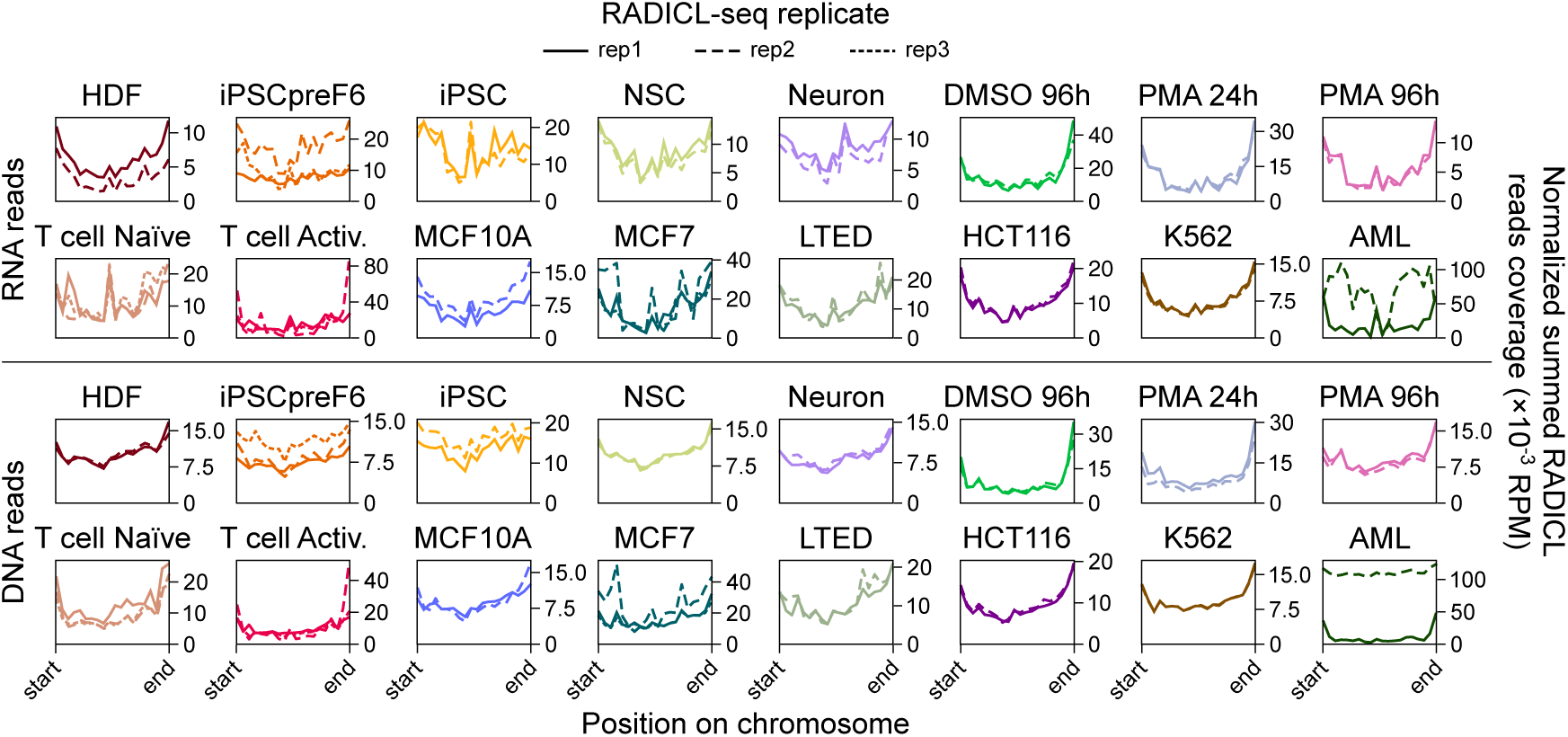
Distribution of RADICL-seq RNA (top) and DNA read fragments (bottom) across chromosomes, in each replicate used in this study. Values for all chromosomes were summed together, after normalizing position values to the total length of each chromosome.

When performing the same analysis on TADs and chromatin loops detected by Hi-C in the Neuron and THP-1 series, we observed that the borders of both types of chromatin structures are particularly prone to be targeted by RNAs (Figures 14 and 30), as previously reported (Bell *et al*., 2018; Bonetti *et al*., 2020; Shu *et al*., 2024; Zvezdin *et al*., 2025). However, unlike at the whole-chromosome scale, this enrichment in RADICL-seq DNA fragments does not correlate with that of RNA fragments, which are more abundant in the center of these regions.

**Figure 30:**
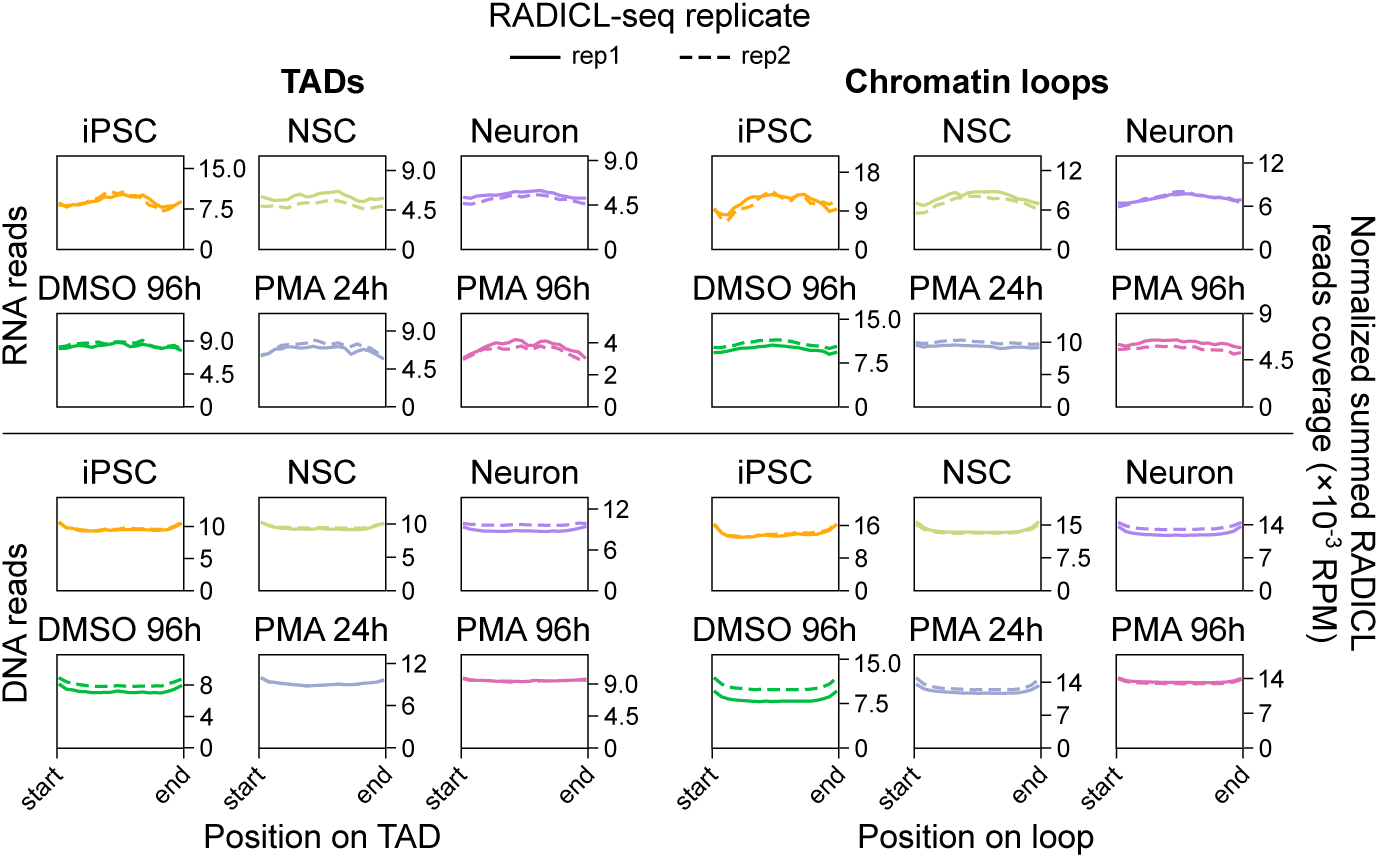
Distribution of RADICL-seq RNA (top) and DNA read fragments (bottom) across TADs (left) and chromatin loops (right), in each sample of the Neuron and THP-1 series. Values for all TADs and loops were summed together, after normalizing position values to the total length of each TAD or each loop. Loop regions were defined as beginning at the start of the first anchor and finishing at the end of the second anchor.

Aside from TADs and loops, the chromatin is also arranged into A/B compartments and subcompartments, which correlate with different levels of accessibility and transcriptional activity (Liu *et al*., 2021; Rao *et al*., 2014). We found that A subcompartments overall produce more DNA-interacting transcripts than B subcompartments, consistent with their global degree of expression, and are also more frequently targeted by RNAs (Figure 31), corroborating past results (Calandrelli *et al*., 2023). These levels gradually decrease as the index of both A and B subcompartment increases, *i.e.* as the chromatin state becomes less active. These results indicate that the RNA-DNA interactome is coupled to the 3D conformation of the chromatin across multiple organizational scales, confirming and generalizing previous observations from other cell types (Agrawal *et al*., 2024; Bonetti *et al*., 2020; Farabella *et al*., 2021; L. Li *et al*., 2021; X. Li *et al*., 2017; Limouse *et al*., 2023; Morf *et al*., 2019; Zvezdin *et al*., 2025).

**Figure 31:**
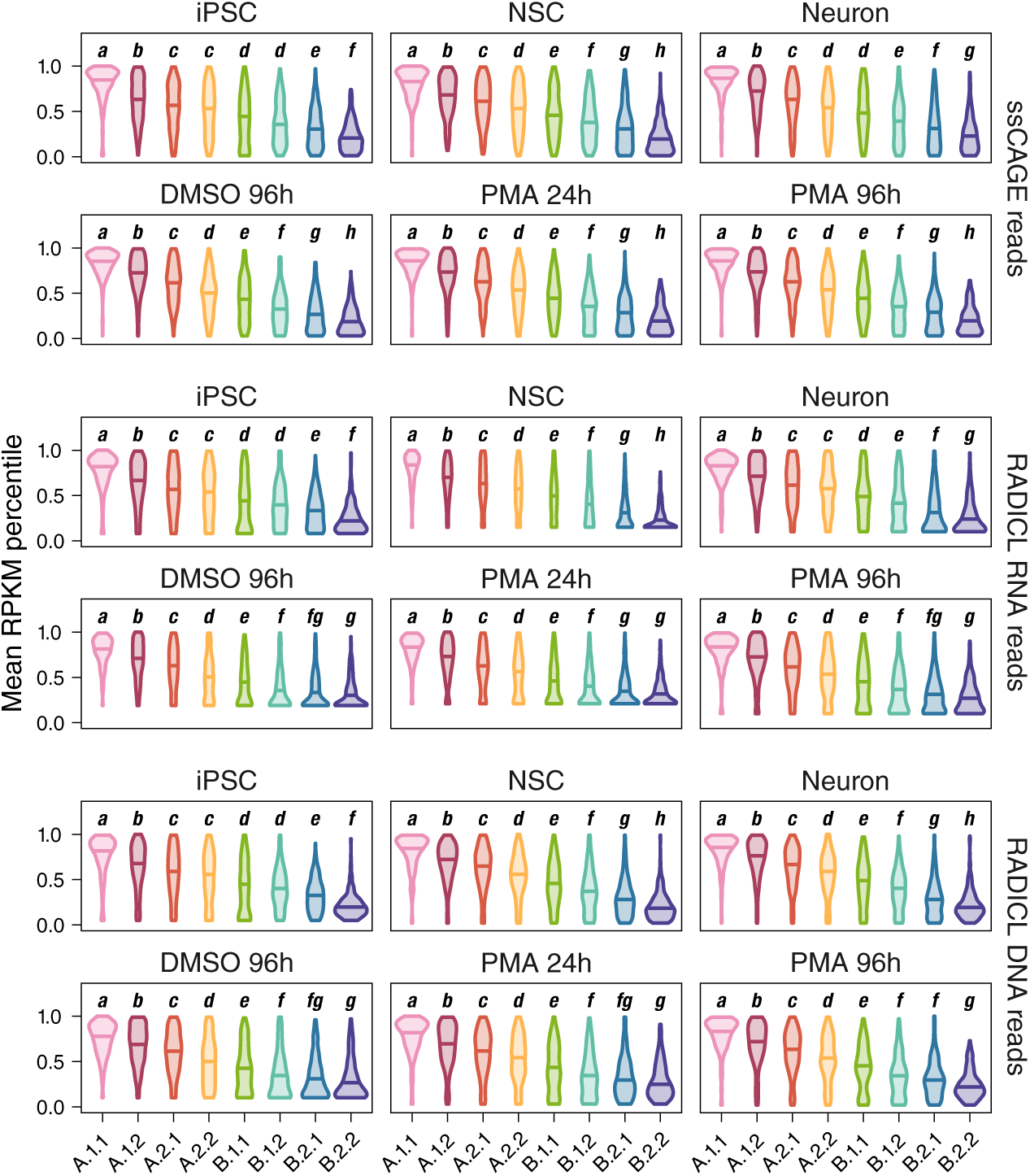
ssCAGE (top), RADICL-seq RNA (middle) and RADICL-seq DNA (bottom) read coverage (RPKM) over different types of A/B subcompartments, in each sample of the Neuron and THP-1 series. For better visualization, each individual subcompartment is assigned to the percentile to which its total read coverage corresponds, calculated across all subcompartments. The bold italic letters indicate significantly different groups of subcompartments, defined by one-way ANOVA with post-hoc Tukey test.

Considering that A/B subcompartments are associated with distinct epigenetic signatures (Figure 32; Liu *et al*., 2021; Rao *et al*., 2014), we next examined whether RNA-DNA interactions are significantly linked to specific features of the chromatin. To this end, we compared RADICL-seq data in the Neuron and THP-1 series with data obtained by CUT&Tag or ChIP-seq, CAGE, ATAC-seq, as well as Hi-C-detected loops. As a validation of our approach, we retrieved the expected enrichment in transcription start site (TSS) clusters and activating histone marks together with a depletion in repressive H3K27me3 at sources of RADICL-seq RNA fragments, which are necessarily expressed (Figure 33). On the other hand, loci corresponding to RADICL-seq DNA fragments are enriched relatively uniformly in all types of chromatin features except loops. These enrichment patterns do not exactly mirror those of source loci, demonstrating that DNA-interacting RNAs do not merely contact expressed regions. In particular, we found that up to ∼70% of peaks corresponding to accessible chromatin or to a given histone modification are not bound by any RNA, with variations between samples (Figure 34). This indicates that chromatin accessibility or the presence of an epigenetic mark alone is not sufficient to determine the targets of RNA-DNA interactions detected by RADICL-seq.

**Figure 32:**
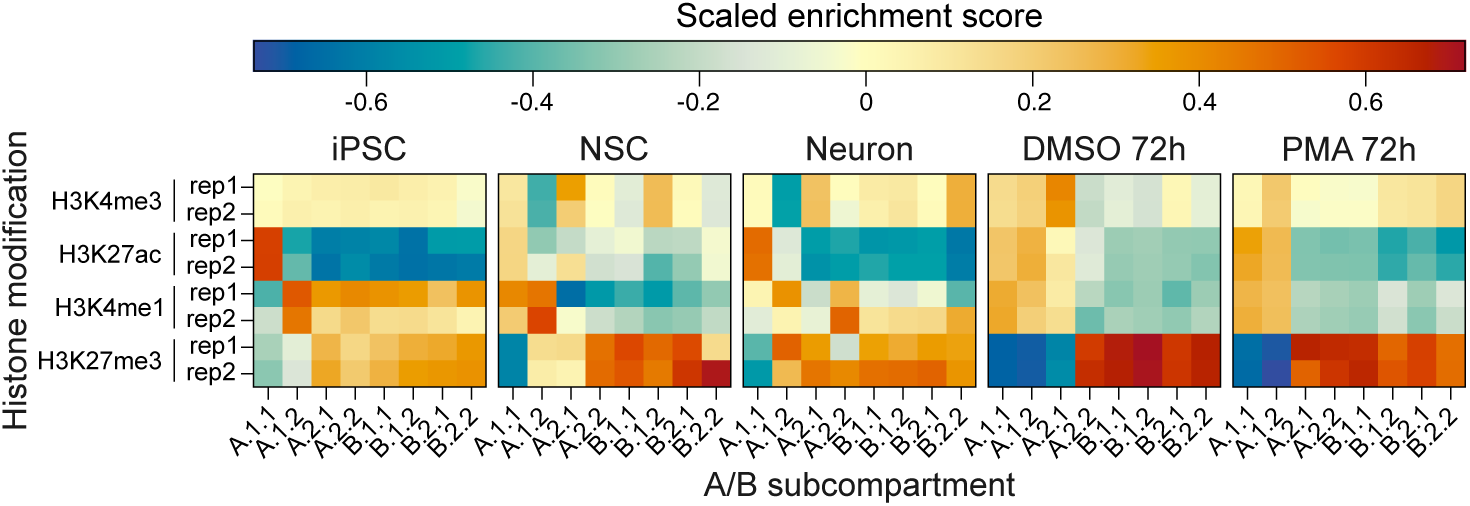
Enrichment in H3K27ac, H3K4me3, H3K4me1 or H3K27me3 for each type of A/B subcompartment compared to the whole genome, in each CUT&Tag or ChIP-seq replicate of the Neuron and THP-1 series. For better visualization, enrichment scores (calculated as described in “Methods”) were scaled by mean-centering within each subcompartment type in each sample.

**Figure 33:**
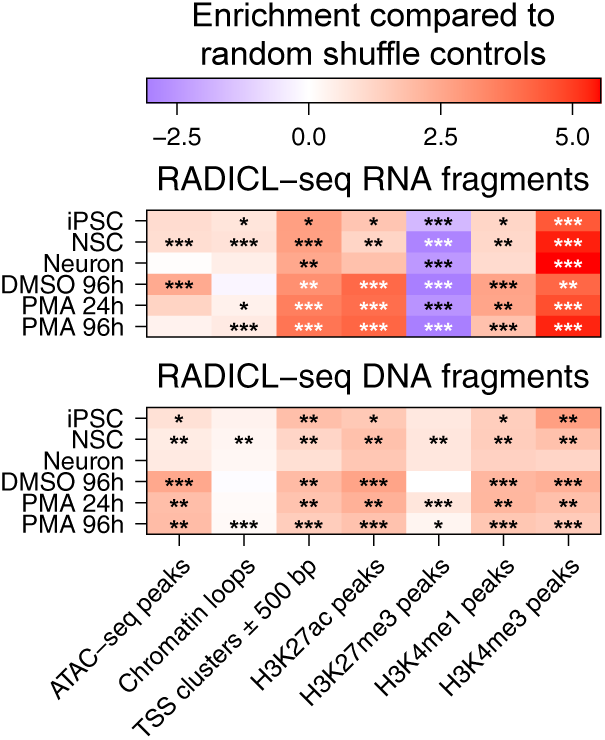
Enrichment of regions containing either an ATAC-seq peak, a chromatin loop, a TSS cluster or a histone mark peak among regions detected by RADICL-seq, either on the RNA side (top) or on the DNA side (bottom), in each sample of the Neuron and THP-1 series, compared to equivalent sets of randomly shuffled regions. A null enrichment value corresponds to no enrichment nor depletion compared to what would be expected by chance. Stars indicate the significance of the enrichment (Student’s t-test; *p value < 0.05; **p value < 0.01; ***p value < 0.001).

**Figure 34:**
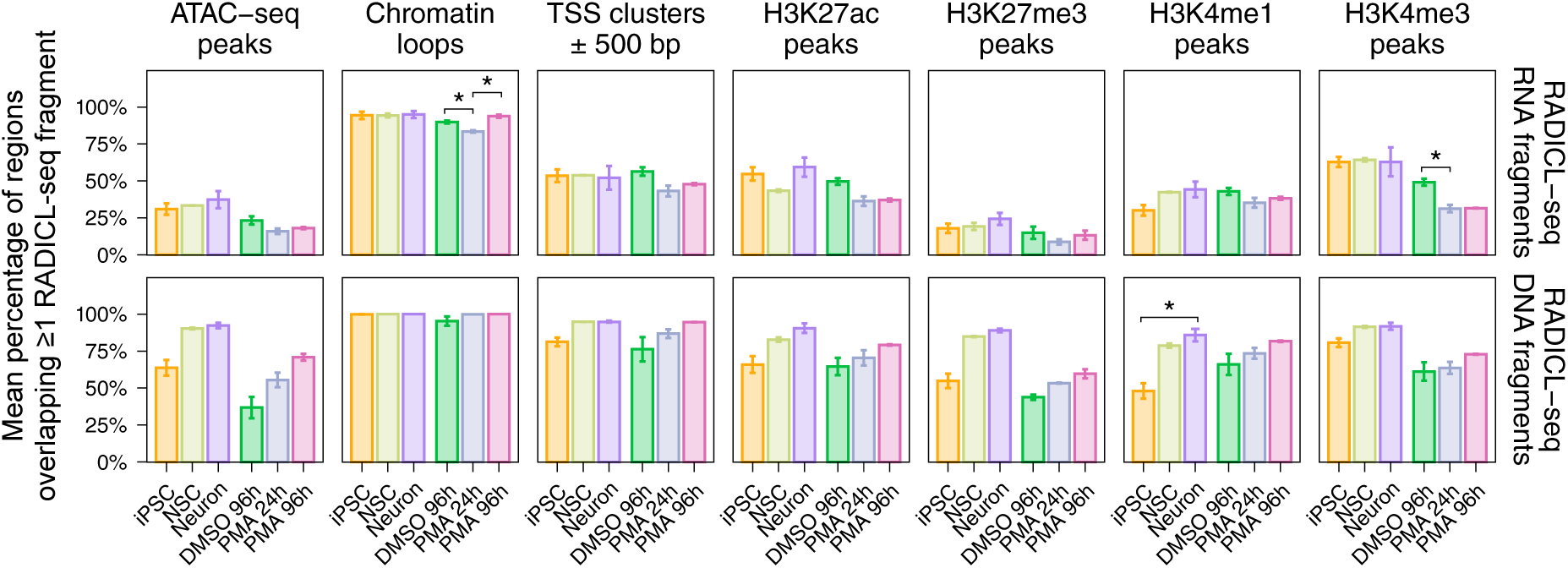
Percentage of ATAC-seq peaks, chromatin loops, extended TSS clusters (± 500bp) or histone mark peaks overlapping with at least one RADICL-seq RNA (top) or DNA read fragment (bottom), in each sample of the Neuron and THP-1 series. Values were averaged between RADICL-seq biological replicates. Stars indicate significant differences between samples (Student’s t-test; *p value < 0.05).

To assess whether different chromatin profiles may be associated with different types of RNA-DNA contacts, we thus adopted a more combinatorial approach. All RNA-DNA interaction target sites detected in at least one sample of the Neuron and THP-1 series (∼147 million sites in total, pooled across all samples) were grouped by *k*-means clustering relative to their A/B subcompartment, their enrichment in histone modifications, their distance to the nearest telomere, TAD border and chromatin loop, as well as their accessibility and transcriptional activity (Figures 35 and 36; Methods).

**Figure 35:**
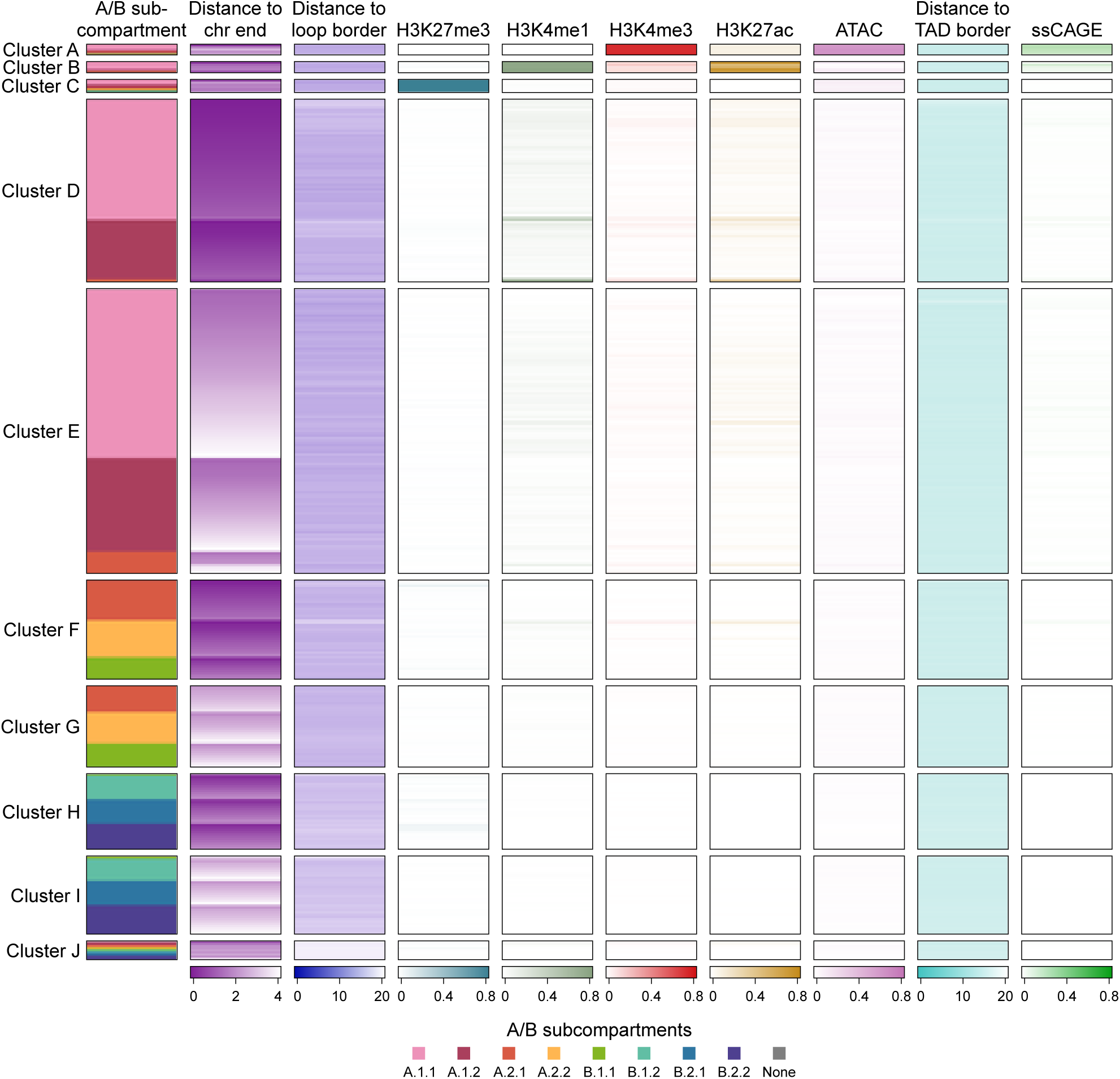
K-means clustering of all unique RADICL-seq DNA read fragments detected in any sample of the Neuron and THP-1 series, in function of the A/B subcompartment of the locus to which these fragments map, its distance to the nearest chromosome end, TAD border or loop border, the enrichment of its overlapping ATAC-seq, H3K27me3, H3K4me1, H3K4me3 and H3K27ac peaks, as well as the ssCAGE signal of its overlapping TSS clusters (± 500 bp), in each of the samples where these fragments were detected. Values for each chromatin parameter are shown normalized as described in Methods. Columns and rows of the heatmap are sorted in function of the results shown in Figure 37.

**Figure 36:**
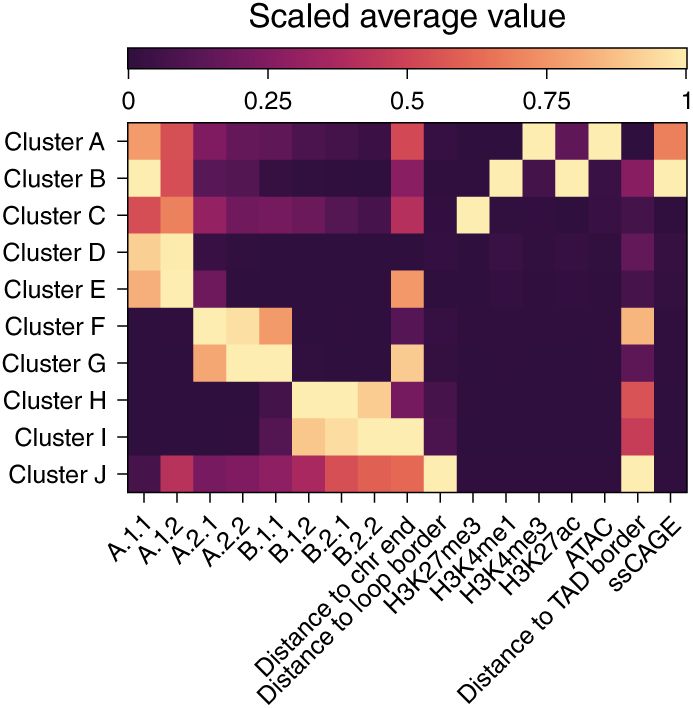
K-means clustering of all unique RADICL-seq DNA read fragments detected in any sample of the Neuron and THP-1 series, in function of the A/B subcompartment of the locus to which these fragments map, its distance to the nearest chromosome end, TAD border or loop border, the enrichment of its overlapping ATAC-seq, H3K27me3, H3K4me1, H3K4me3 and H3K27ac peaks, as well as the ssCAGE signal of its overlapping TSS clusters (± 500 bp), in each of the samples where these fragments were detected. For each chromatin parameter and each cluster, normalized values shown in Figure 35 were averaged over the whole cluster, then all resulting means were scaled to a 0-to-1 range.

By training a Random Forest model to predict the cluster assignments based on the target sites’ original chromatin parameters, we found that the most important factors to differentiate regions targeted by RNAs are their A/B subcompartments and their distance to the chromosome end (Figure 37). Meanwhile, the presence of histone marks, active TSS clusters or TAD/loop borders were much less impactful for the clustering, consistent with results from Tabe-Bordbar and Sinha (2023) who performed a similar analysis on lncRNA-mediated contacts from mouse embryonic stem cells.

**Figure 37:**
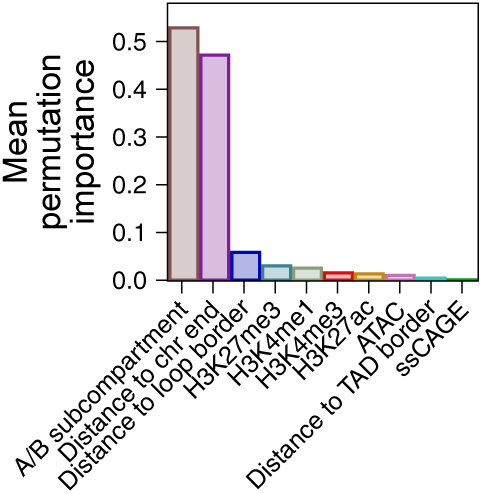
Mean permutation-based feature importance scores derived from ten Random Forest classifiers trained to predict k-means cluster assignments from the results shown in Figures 35 and 36, each computed on a different randomly selected 1% subset of the data.

Our clustering approach yielded ten different clusters of RNA-targeted sites, including three with an easily recognizable signature (Figures 35 and 36; V. W. Zhou *et al*., 2010): cluster A, characterized by an active promoter-like landscape rich in H3K4me3 and active TSS clusters, and mostly located in A.1 subcompartments; cluster B, composed of loci also primarily located in A.1 subcompartments and slightly less transcriptionally active, but enriched in H3K27ac and H3K4me1 like enhancers; and cluster C, consisted of loci present in any A/B subcompartment, but that all lack transcriptional activity and are heavily marked by H3K27me3, likely corresponding to repressed genes. Importantly, these three clusters represent only a very small fraction (4%) of all the target sites detected by RADICL-seq. The loci assigned to the remaining seven clusters are mainly differentiated by their A/B subcompartment and their distance to the nearest telomere, without any particular epigenetic or transcriptional signature. A notable exception is cluster J, which encompasses all regions far from loop borders, regardless of their A/B subcompartment.

Despite making up a minor portion of all RNA-targeted sites, we observed that loci belonging to clusters A, B or C are overall the most intensely targeted by RNAs in each sample of the Neuron and THP-1 series (Figure 38), both because they are targeted by the highest number of distinct transcripts (Figure 39) and because each of these transcript-to-cluster interactions tends to be more frequent than those targeting other clusters (Figure 40). This is in line with results obtained by other technologies in diverse cell types (Bell *et al*., 2018; Calandrelli *et al*., 2023; X. Li *et al*., 2017; Sridhar *et al*., 2017; Yun *et al*., 2024; B. Zhou *et al*., 2019) and, at least for clusters A and B, may be imputed to these loci being located in the most active A/B subcompartments (Figures 31, 35 and 36).

**Figure 38:**
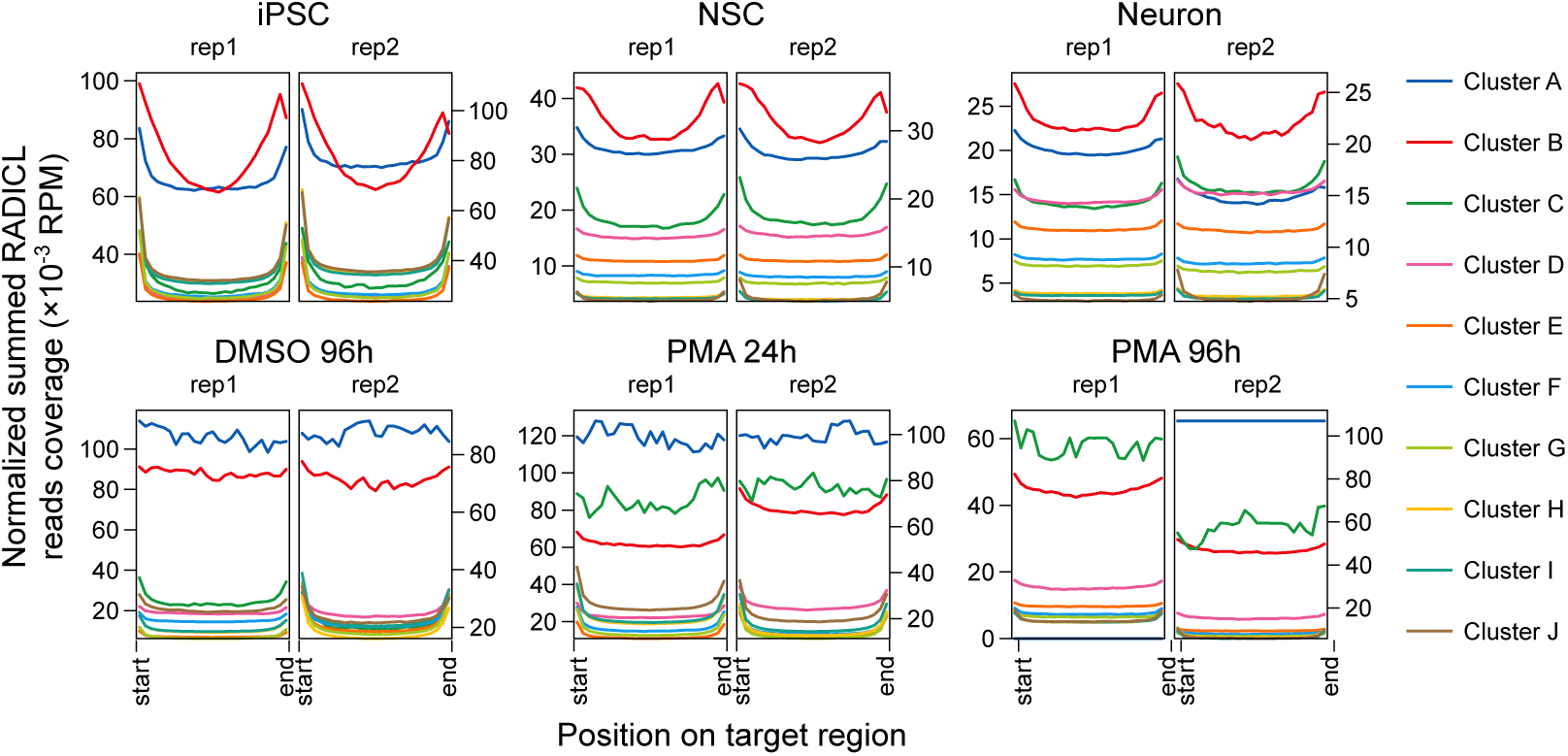
Distribution of DNA read fragments across all loci belonging to each cluster depicted in Figures 35 and 36, in each replicate of the Neuron and THP-1 series. Values for all loci belonging to the same cluster were summed together, after normalizing the position values to the total length of each locus.

**Figure 39:**
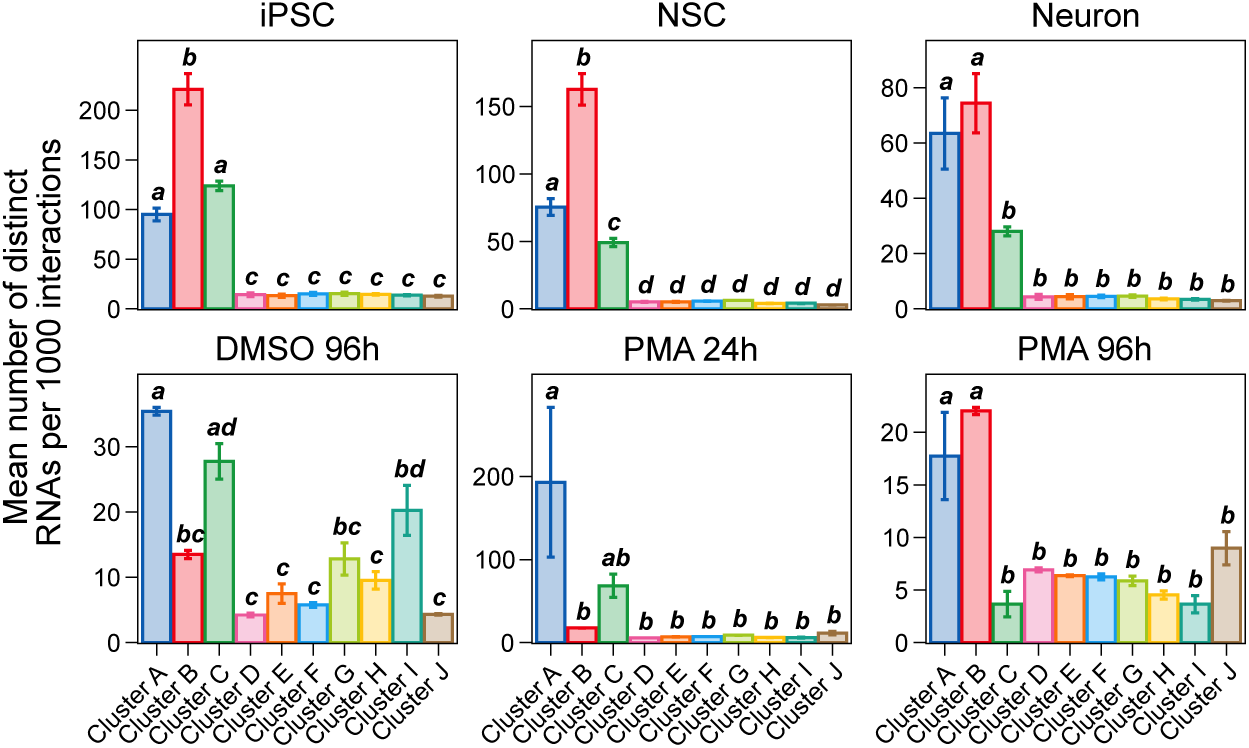
Number of distinct transcript isoforms targeting each cluster depicted in Figures 35 and 36, in each sample of the Neuron and THP-1 series. Values were averaged by RADICL-seq replicate and normalized by the total number of interactions targeting the same cluster. Error bars show the standard deviation between biological replicates; bold italic letters indicate significantly different groups of clusters, defined by one-way ANOVA with post-hoc Tukey test.

**Figure 40:**
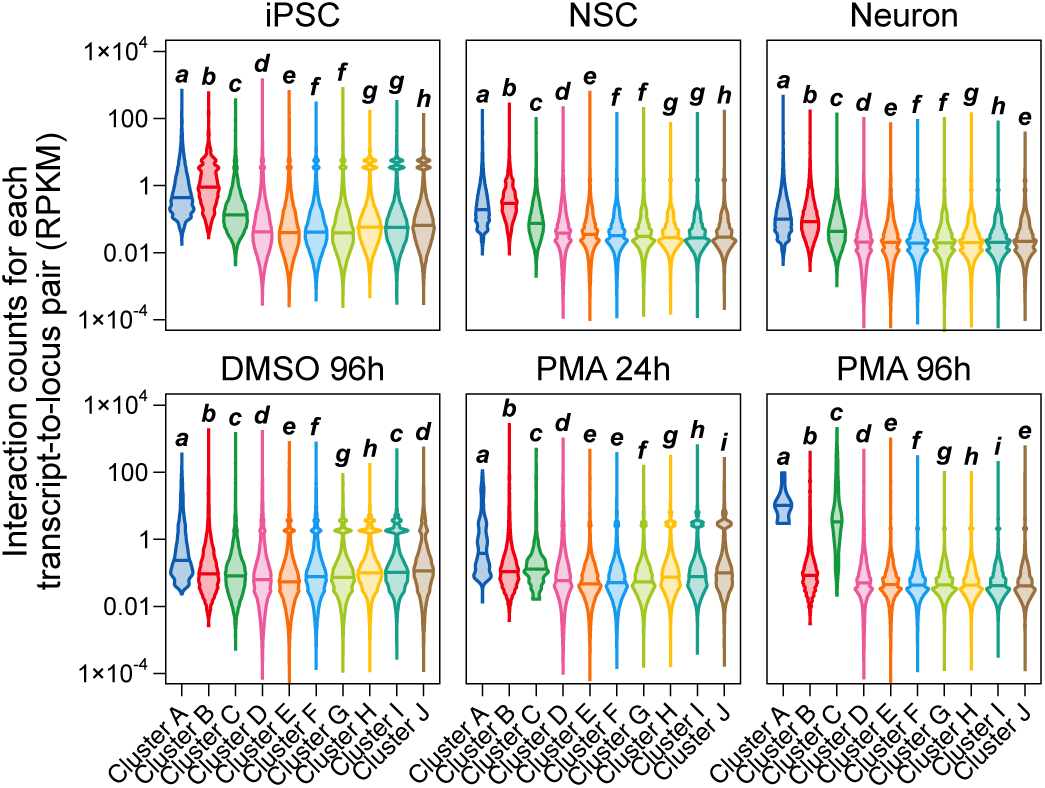
Frequency of interaction (RPKM) for each source-target pair in each sample of the Neuron and THP-1 series, in function of the cluster that the target belongs to, as defined in Figures 35 and 36. For each sample, the two RADICL-seq biological replicates were pooled together. Bold italic letters indicate significantly different groups of clusters, defined by one-way ANOVA with post-hoc Tukey test.

When evaluating the array of different clusters targeted by each individual RNA that contacts multiple loci, including at least one in *trans* to avoid proximity biases, we found that the vast majority of transcripts interact with several clusters (Figure 41). This suggests that RNAs as a whole have a relatively low specificity for different types of chromatin contexts, contrary to previous reports (Limouse *et al*., 2023; Zvezdin *et al*., 2025). Nonetheless, we observed that loci assigned to A subcompartments (clusters A to G) tend to be more often targeted in *cis* than in *trans* compared to clusters assigned to B subcompartments (clusters H and I), consistent with the global transcriptional trends of these subcompartments (Figure 31), but also reflecting some cell type-specificity. For instance, interactions targeting regions far from chromatin loops (cluster J) have the highest proportion of *cis* contacts of all clusters in DMSO 96h, but one of the lowest in PMA 96h, highlighting that both the chromatin and cellular contexts contribute to determining the interaction between a given transcript and a given target region.

**Figure 41:**
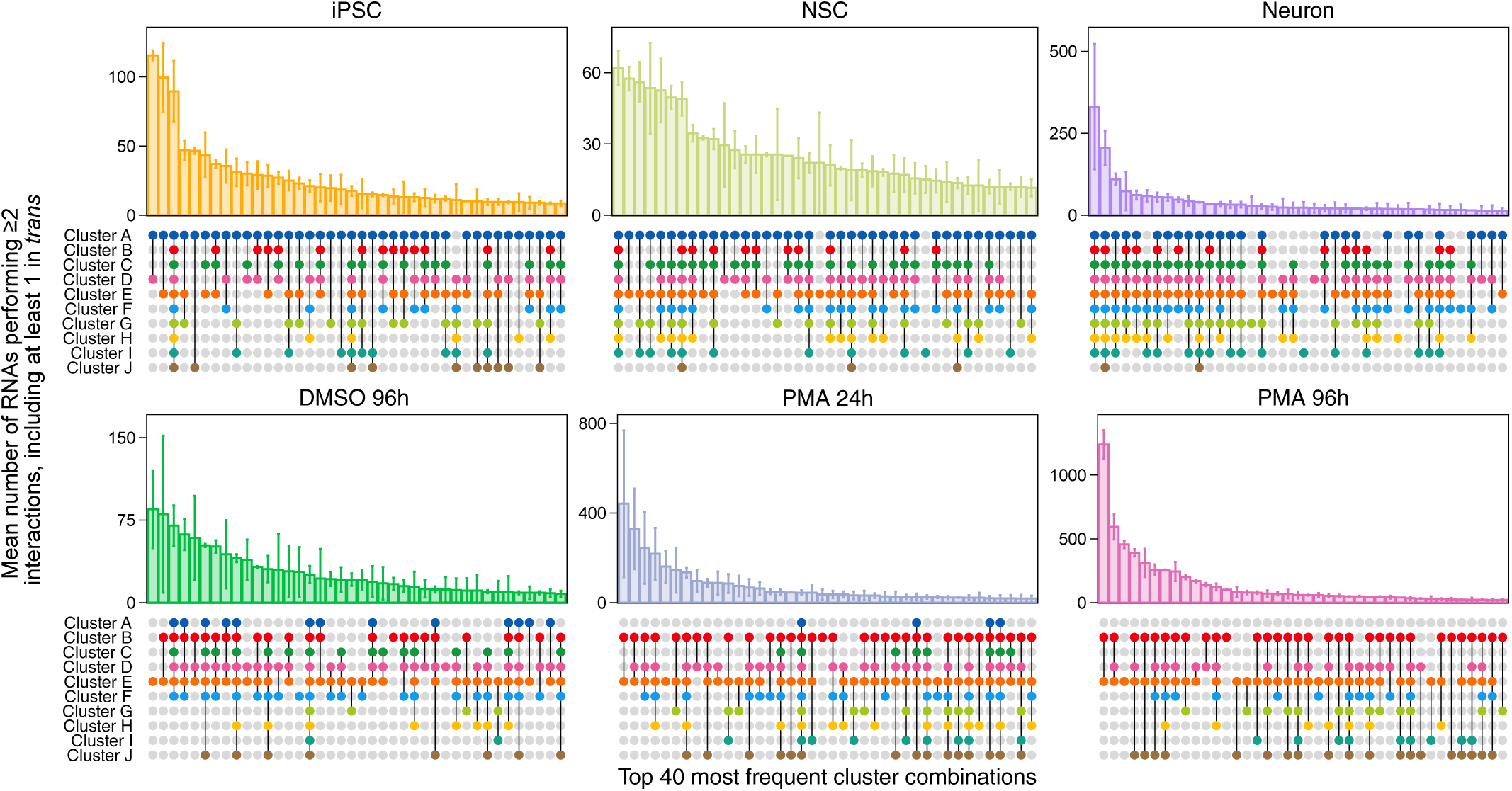
Number of DNA-interacting transcript isoforms in function of the combination of different clusters that they target, averaged over the two RADICL-seq biological replicates for each sample of the Neuron and THP-1 series. To avoid proximity and unique-interaction biases, only RNAs targeting at least two different loci, including at least one in trans, were considered for this figure. For better visibility, only the top 40 most frequent cluster combinations are shown. Error bars indicate the standard deviation between replicates.

Since our RADICL-seq data include two time series that represent different contexts of cell specialization, we could next explore how these dynamic processes affect the RNA-DNA interactome. For this purpose, we extracted all pairs of interacting transcript/clustered locus for which the frequency of interaction is significantly different between samples of the Neuron series on the one hand and of the THP-1 series on the other hand (Figure 43).

**Figure 42:**
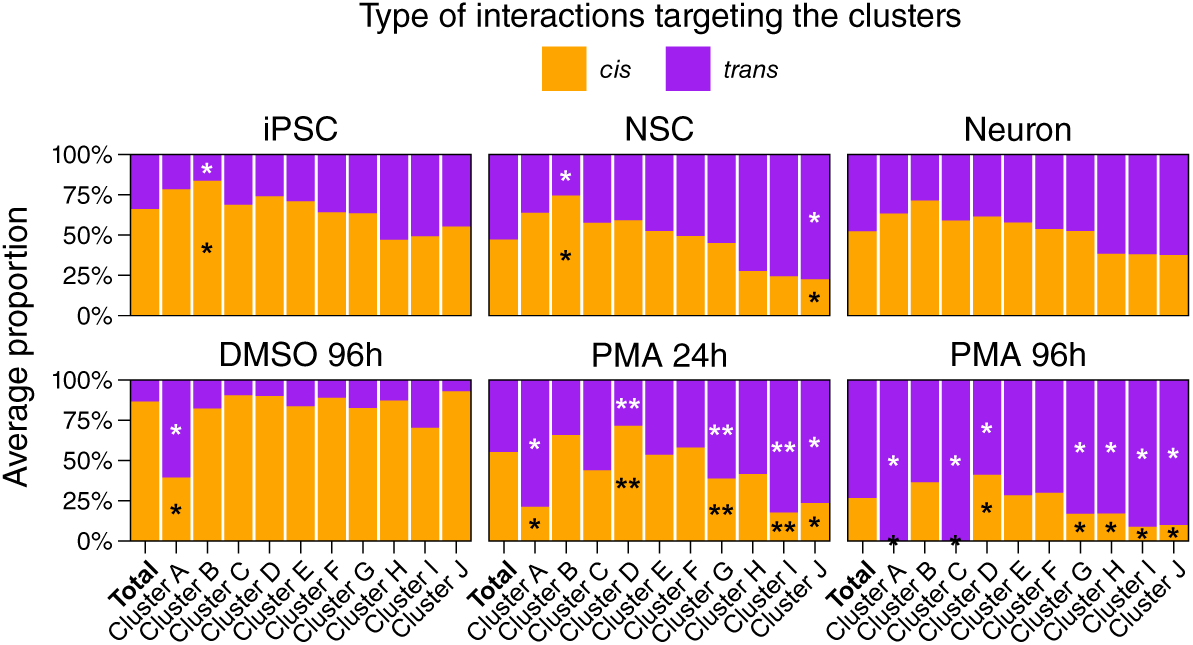
Proportion of cis and trans interactions among all interactions targeting each cluster depicted in Figures 35 and 36, averaged between RADICL-seq biological replicates in each sample of the Neuron and THP-1 series. The significance of the enrichment or depletion of a type of interaction among interactions targeting a given cluster was calculated by hypergeometric test against all interactions detected in that sample; * p value < 0.05; ** p value < 0.01.

**Figure 43:**
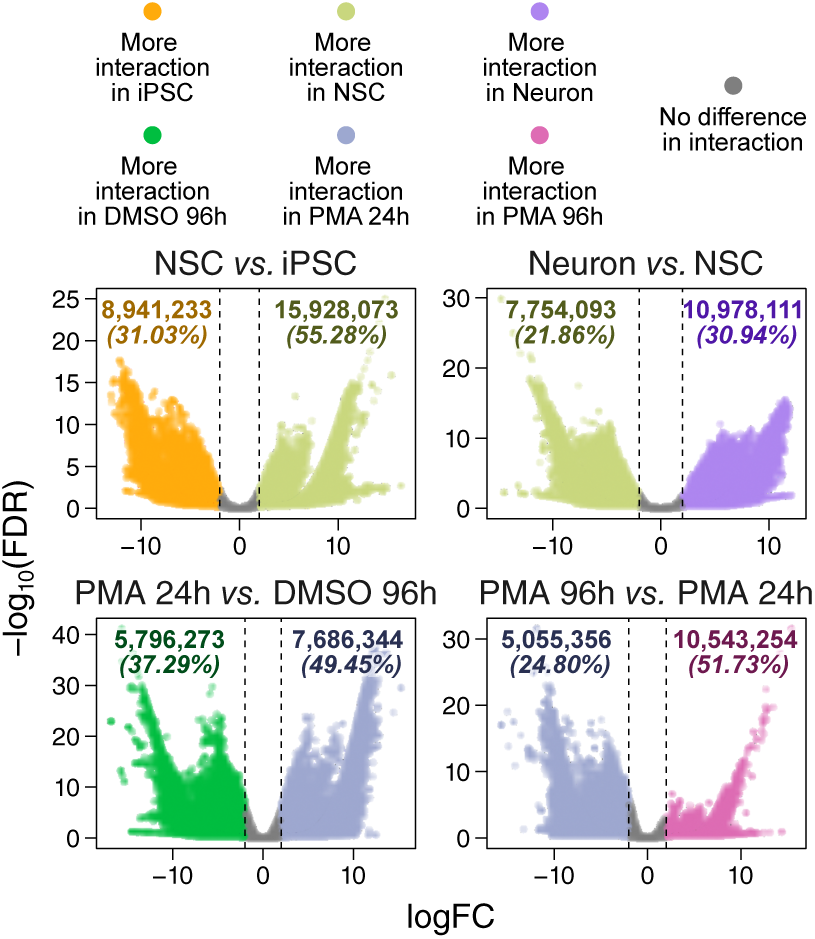
Changes in RNA-DNA interaction frequency between samples of the Neuron and THP-1 series, at the transcript-isoform-to-cluster-region level. Differential interactions (DIs) are defined as those having an absolute logFC > 2 (edgeR, Robinson et al., 2009) and their number is indicated at the top of each panel.

One mechanism underlying these differential interactions (DIs) can be the differential expression of their source transcripts, since RADICL-seq RNA signal correlates with expression measured by CFC-seq and RNA-seq, albeit less strongly than the correlation between CFC-seq and RNA-seq (Figure 44). We thus combined our RADICL-seq data with a differential expression analysis performed on CFC-seq data to separate DIs for which the change in frequency correlates with the change in source expression (hereinafter referred to as “DE-DIs”) from those for which the source transcript isoform is not differentially expressed (“nonDE-DIs”; Figure 45). We found that, on average, DE-DIs represent only ∼50% of all the differential RNA-DNA interactions detected in NSC *versus* iPSC, Neuron *versus* NSC, and PMA 24h *versus* DMSO 96h, with some sample comparison-specific variations in function of the cluster to which the targeted DNA locus belongs; in PMA 96h *versus* PMA 24h, this percentage even drops to less than 5%. Furthermore, the fold change and significance of DE-DIs are not substantially different from those of nonDE-DIs (Figure 46), revealing that a change in expression at the source locus is not the only parameter driving the evolution of the RNA-DNA interactome during cell specialization.

**Figure 44:**
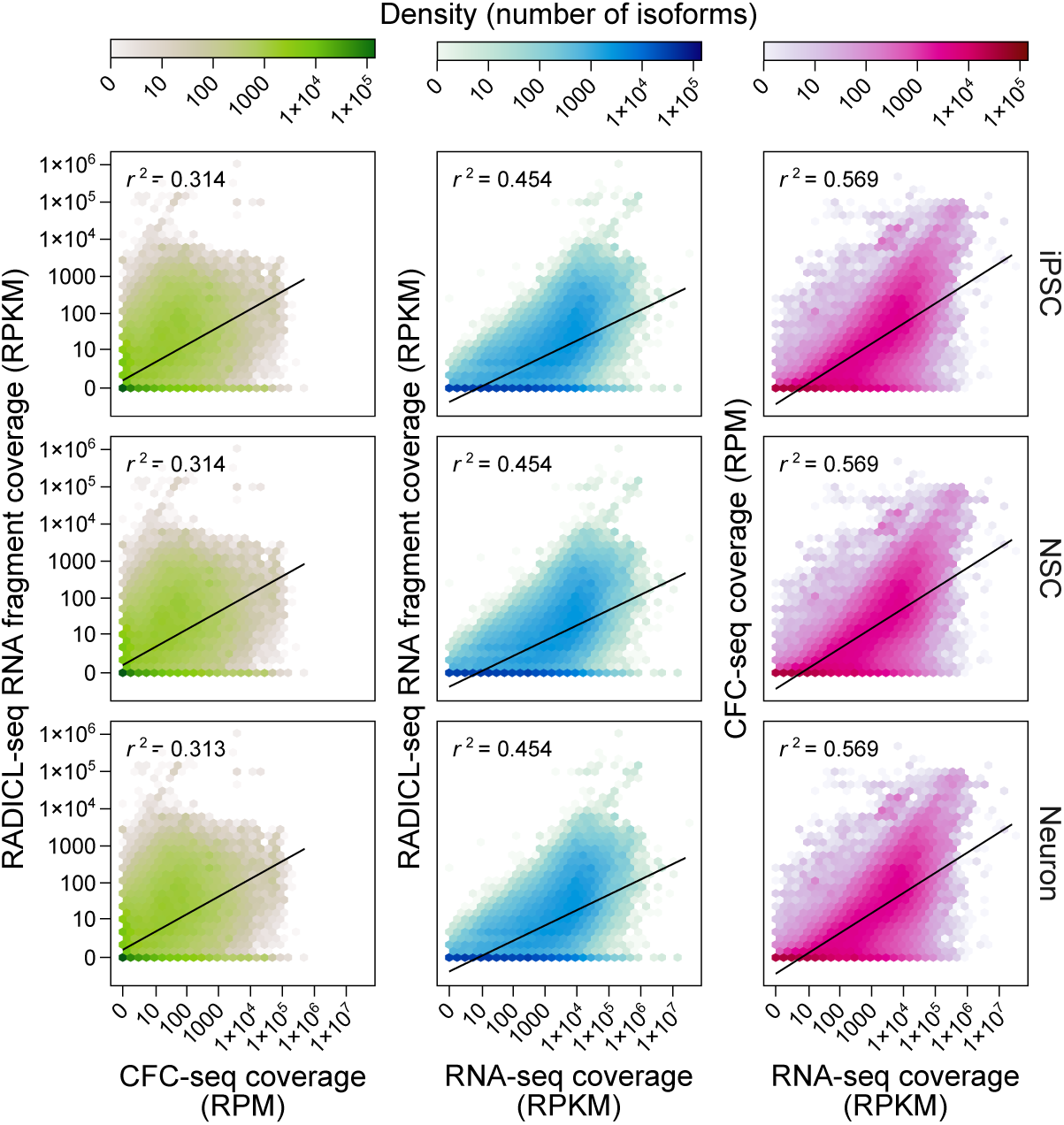
Correlation between read counts obtained from short-read RNA-seq, CFC-seq and RADICL-seq RNA fragments, for all transcript isoforms present in the SACAGE annotation and in each sample of the Neuron series. Black lines correspond to simple linear regressions between (x) and (y); the corresponding coefficients of determination (r^2^) are indicated in the top left corner of each panel.

**Figure 45:**
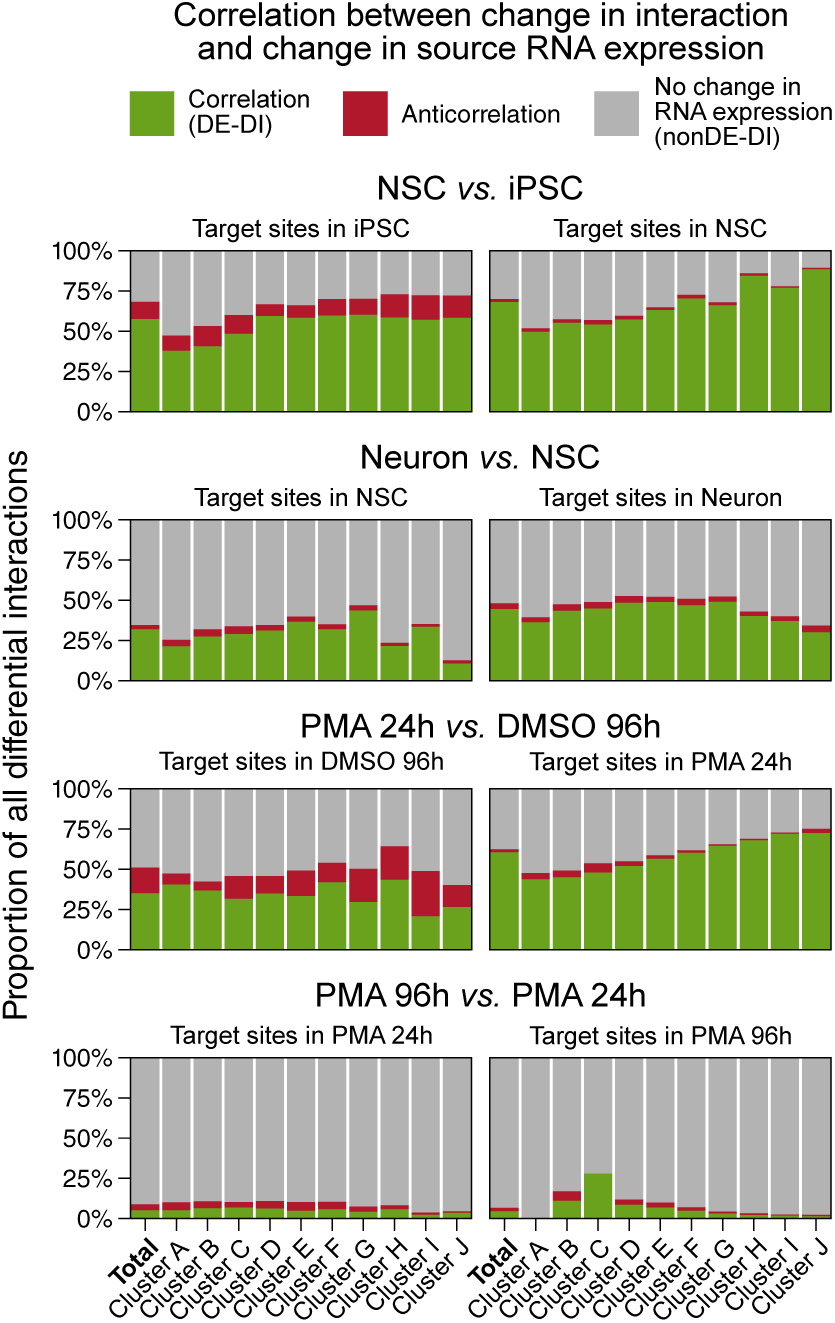
Percentage of DIs targeting each cluster for which the source RNA is differentially expressed (absolute logFC > 1.2, edgeR, Robinson et al., 2009) in the same direction (correlation) or in the opposite direction (anticorrelation) as the change in RNA-DNA interaction frequency. Interactions are separated in function of the cluster to which their target site belong, as defined in Figures 35 and 36.

**Figure 46:**
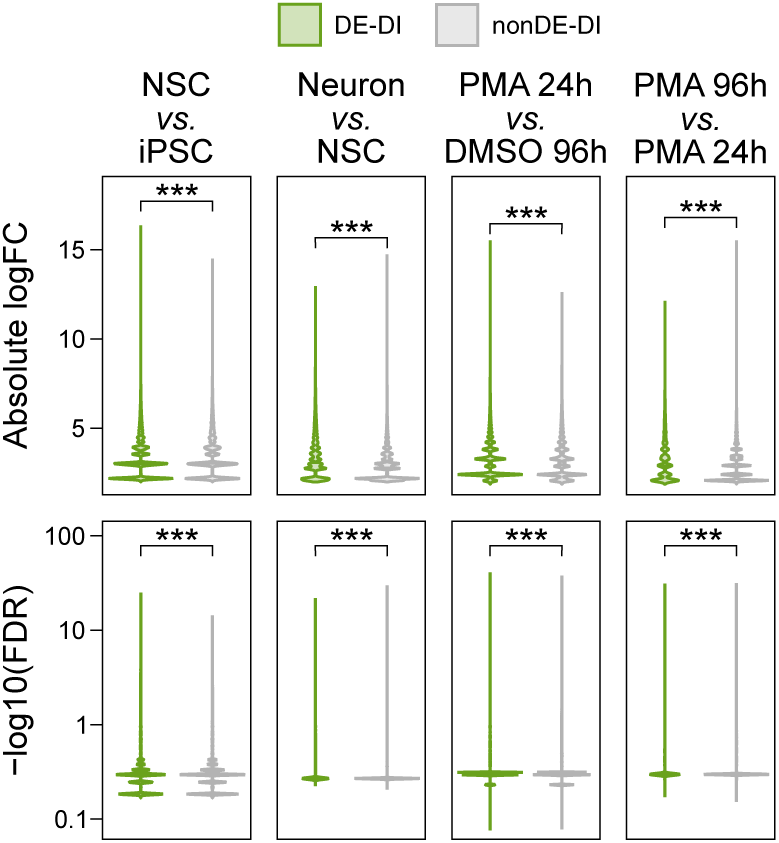
Distribution of the absolute logFC (top) and FDR (bottom) of DIs detected for each sample comparison in the Neuron and THP-1 (see Figure 43), in function of whether the source of these interactions is consistently differentially expressed (DE-DIs) or not differentially expressed (nonDE-DIs) between the same samples. Stars indicate significant differences between DE-DIs and nonDE-DIs (Wilcoxon signed-rank test; ***p value < 0.001).

Another influential parameter could be TAD reconfiguration: if TAD borders act as physical barriers to the spread of RNAs, as suggested by our results in Figures 14 and 30 and by previous reports (Bell *et al*., 2018; Bonetti *et al*., 2020; Calandrelli *et al*., 2023; Kuang & Pollard, 2024; Zvezdin *et al*., 2025), then changes in TAD organization between cellular states should, in principle, alter RNA-DNA interactions targeting TAD edges. However, only a very small fraction of nonDE-DIs are associated with the appearance or disappearance of a TAD border between their source and target loci during neuronal or macrophage specialization (Figure 47). In particular, interactions switching from being intra-TAD to inter-TAD tend to increase in frequency virtually as often and as significantly as those switching from inter-TAD to intra-TAD or as those that remain in the same TAD configuration (Figure 48), indicating that changes in TAD borders between two loci do not consistently affect RNA-DNA interaction frequency. Similar results were obtained at the chromatin loop level, with no enrichment of DIs among interactions affected by loop remodeling (Figures 49 and 50).

**Figure 47:**
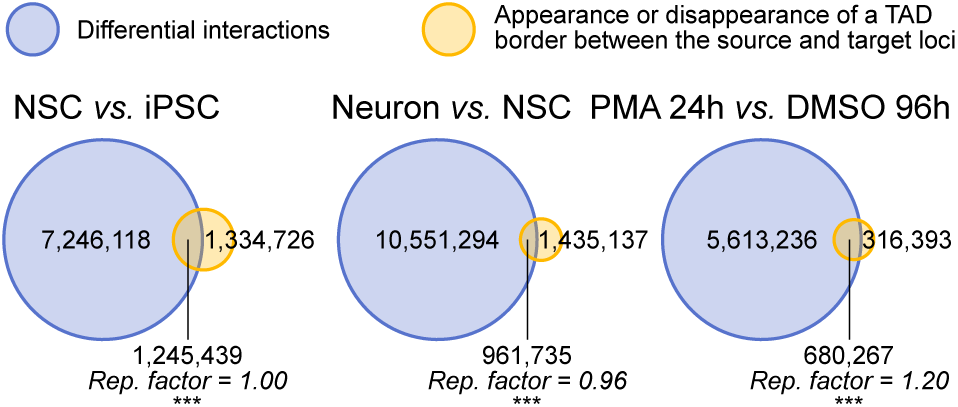
Overlap between nonDE-DIs and RNA-DNA interactions for which a TAD border appears or disappears between the source and the target loci, for each sample comparison in the Neuron and THP-1 series. The enrichment and significance of the overlap (representation factor, hypergeometric test) is indicated in italic; ***p value < 0.001.

**Figure 48:**
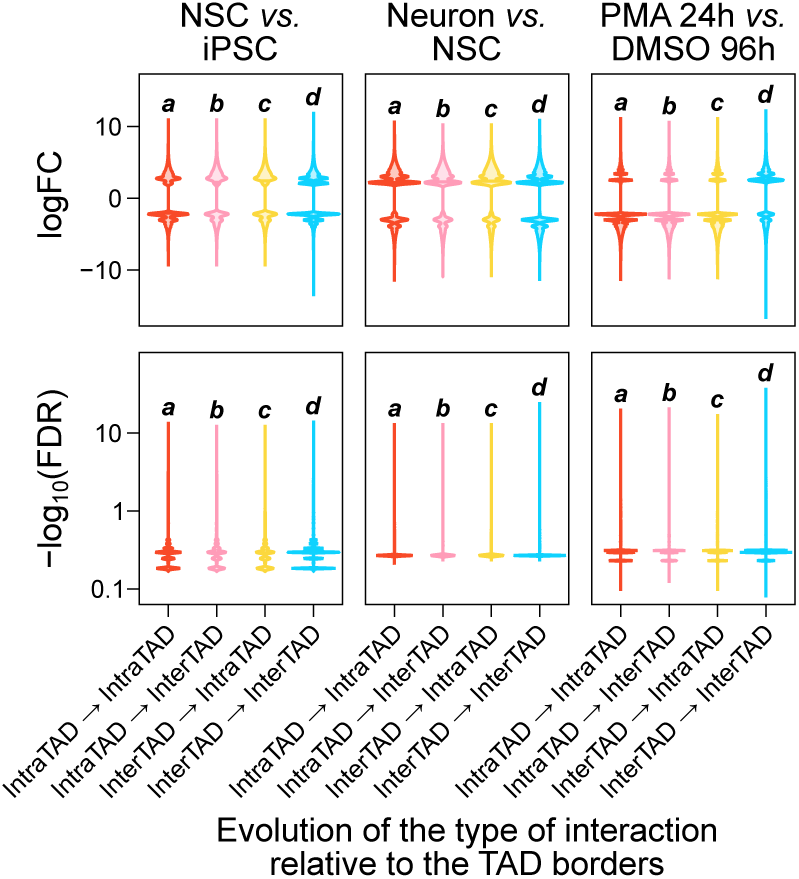
logFC and FDR (edgeR, Robinson et al., 2009) of the interaction frequency of all nonDE-DIs, separated in function of whether or not a TAD border appears or disappears between their source and target loci, for each sample comparison in the Neuron and the THP-1 series. The bold italic letters indicate significantly different groups of interaction changes, defined by one-way ANOVA with post-hoc Tukey test.

**Figure 49:**
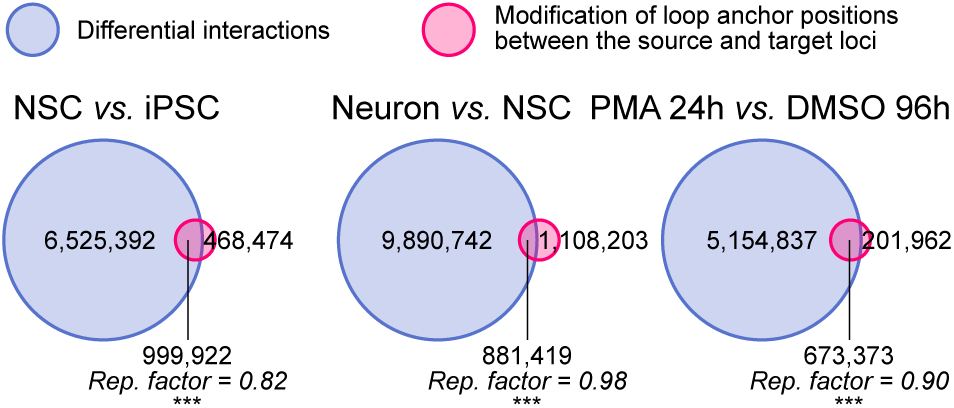
Overlap between nonDE-DIs and RNA-DNA interactions for which a chromatin loop anchor is modified between the source and the target loci, for each sample comparison in the Neuron and THP-1 series. The enrichment and significance of the overlap (representation factor, hypergeometric test) is indicated in italic; ***p value < 0.001.

**Figure 50:**
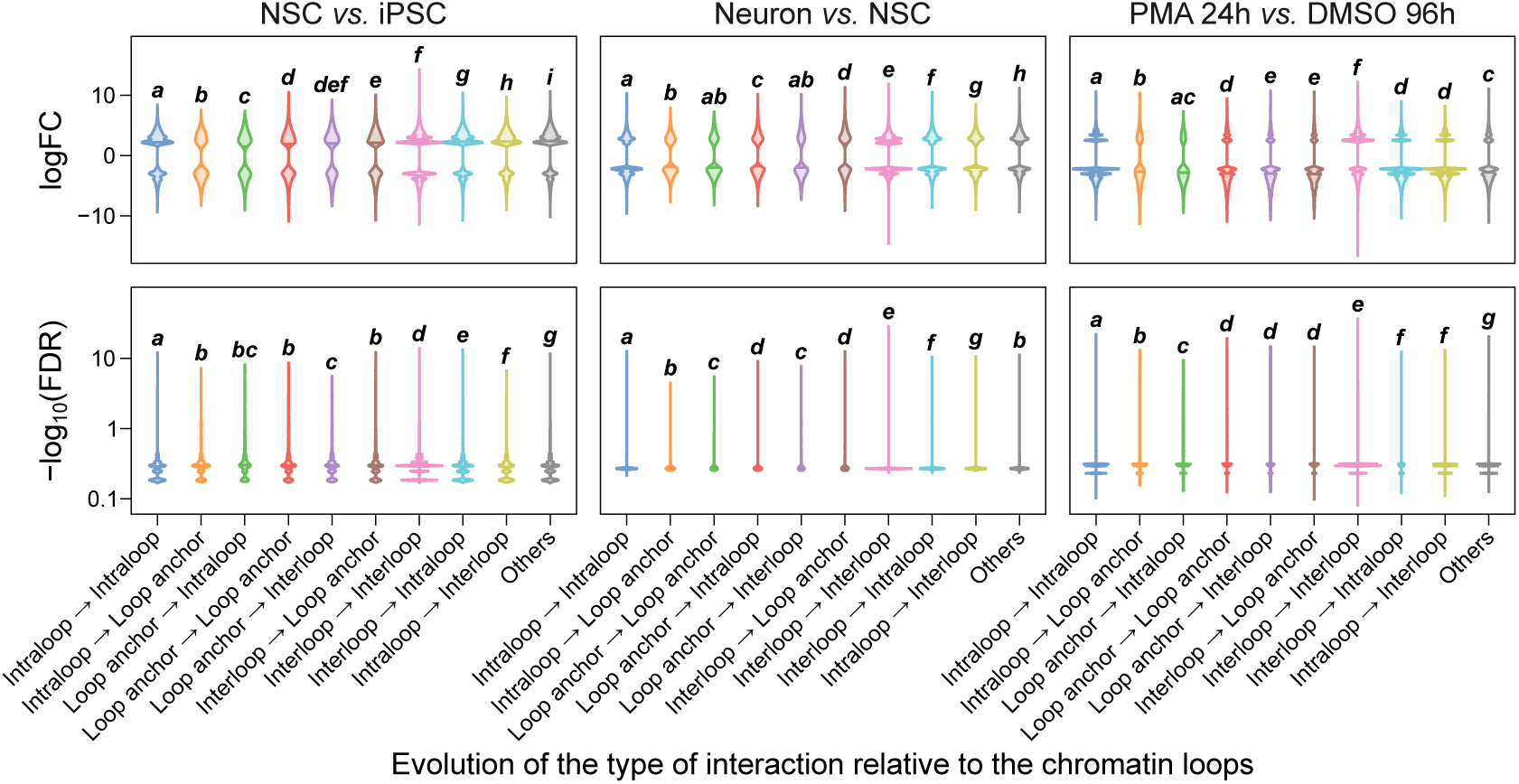
logFC and FDR (edgeR, Robinson et al., 2009) of the interaction frequency of all nonDE-DIs, separated in function of whether or not a chromatin loop anchor is modified between their source and target loci, for each sample comparison in the Neuron and the THP-1 series. The bold italic letters indicate significantly different groups of interaction changes, defined by one-way ANOVA with post-hoc Tukey test.

Instead of TADs and loops, the changes in 3D chromatin organization affecting RNA-DNA interactions could occur at the level of A/B subcompartments, especially given their correlation with global RNA-targeting frequency observed in Figure 31. We thus evaluated the proportion of nonDE-DIs the evolution of which correlates with a modification of the A/B subcompartment at their target site (Figures 51 and 52). While many target sites of nonDE-DIs, especially those that are not located in A.1 subcompartments (clusters F, G, H, I and J), are subjected to an A/B subcompartment change, we also found that these subcompartment changes correlate both positively and negatively at roughly equal rates with changes in interaction frequency (Figures 51 and 52). These findings suggest that an alteration of the A/B subcompartment at a given locus is indeed linked to variations in the RNA-DNA interactions targeting this locus, but this relationship seems to be case-dependent and likely relies on additional factors.

**Figure 51:**
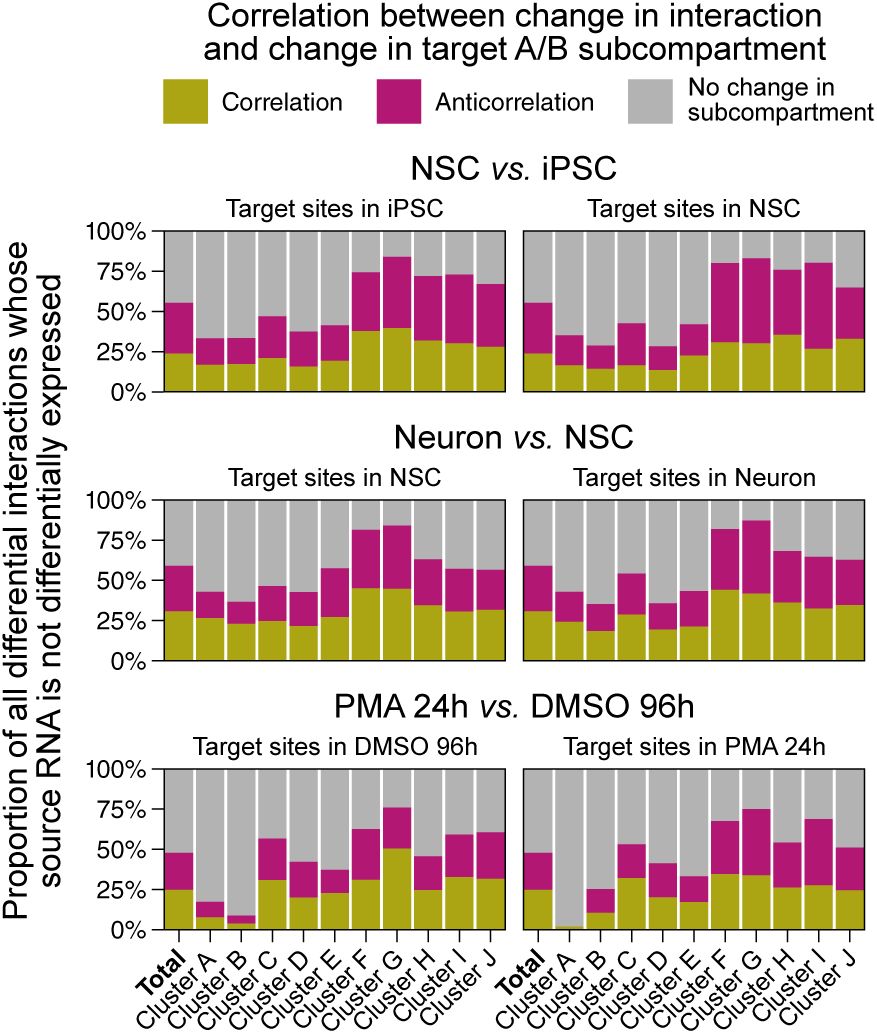
Percentage of the nonDE-DIs targeting an A/B subcompartment that is modified consistently (correlation) or inconsistently (anticorrelation) with the change in RNA-DNA interaction frequency, relative to the trend depicted in Figure 31, for each sample comparison in the Neuron and the THP-1 series. Interactions are separated in function of the cluster to which their target site belong, as defined in Figures 35 and 36.

**Figure 52:**
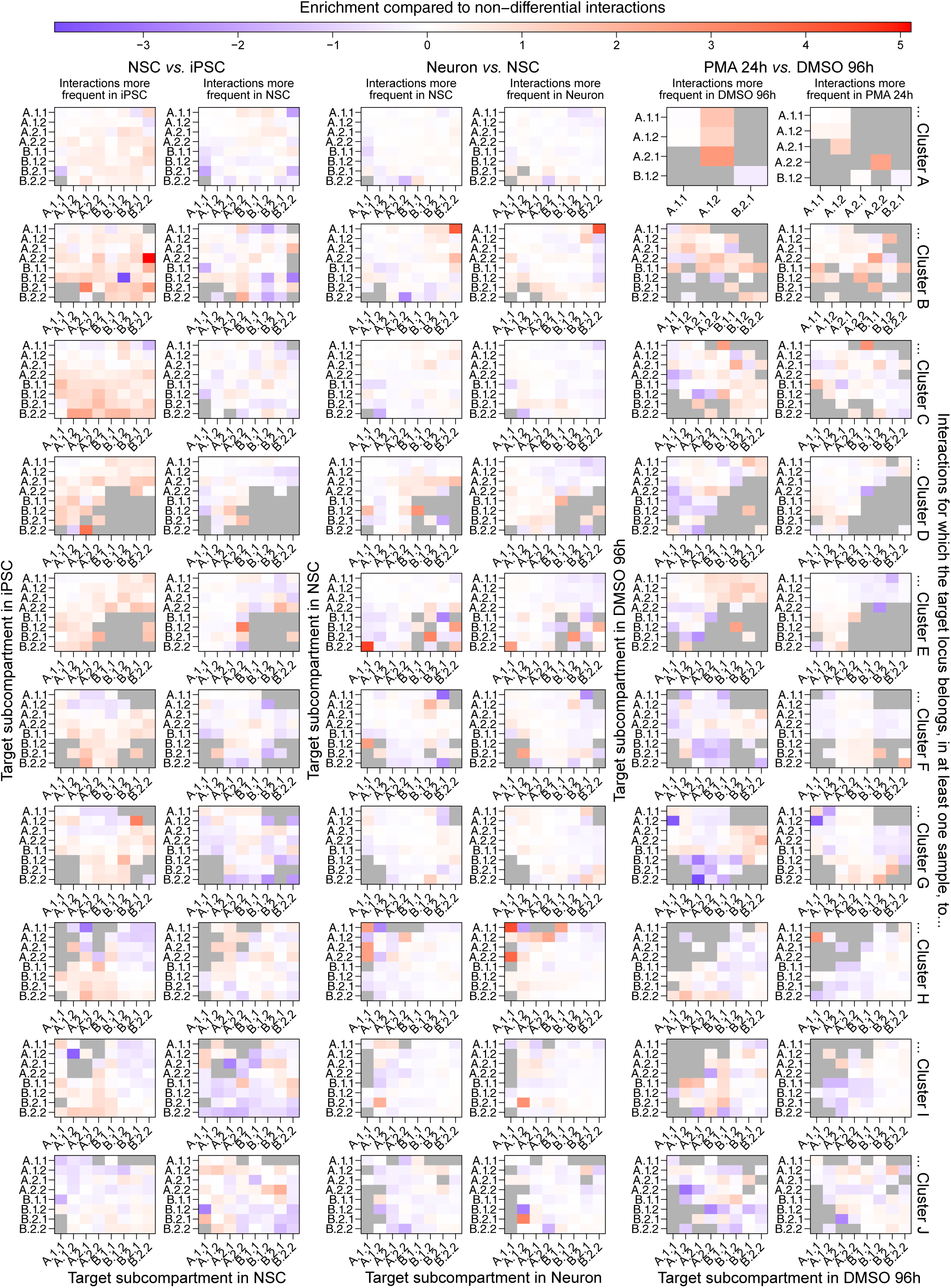
Enrichment (log of ratio) for each possible change in A/B subcompartment between samples of the Neuron and THP-1 series, when comparing the target sites of nonDE-DIs to those of non-DIs. Interactions are separated in function of the cluster to which their target site belong, as defined in Figures 35 and 36.

These additional factors do not appear to involve chromatin accessibility nor histone modifications, as promoters (cluster A), enhancers (cluster B) and repressed genes (cluster C) targeted by nonDE-DIs are not particularly enriched in regions showing differential accessibility or epigenetic mark enrichment compared to those targeted by non-DIs (Figure 53). Similarly, loci showing differential accessibility or histone modification enrichment are targeted by DIs at similar frequencies as other loci (Figure 54). Thus, changes in the epigenetic landscape at the target site are unlikely to be major drivers of RNA-DNA interaction dynamics.

**Figure 53:**
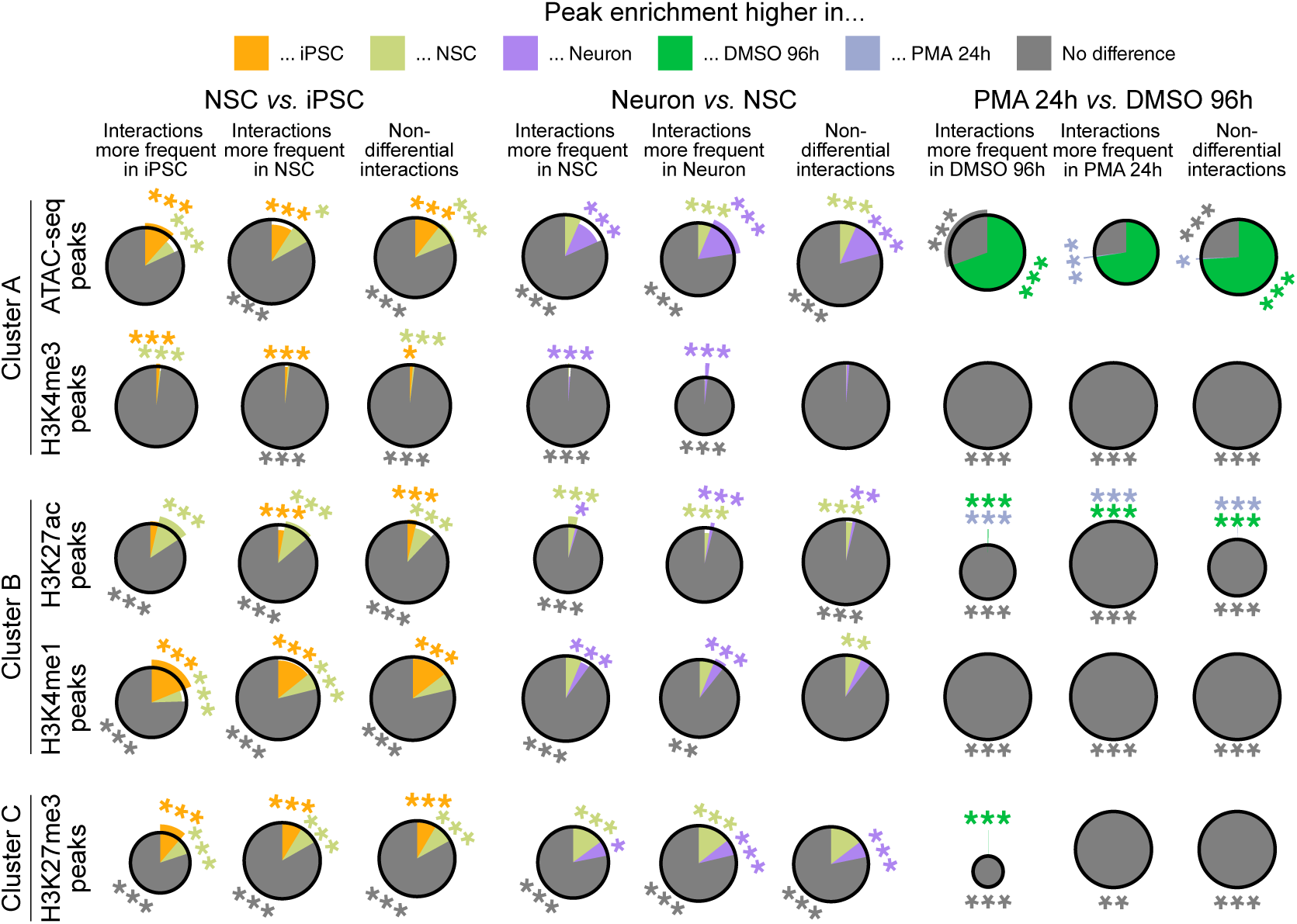
Distribution of nonDE-DIs targeting ATAC-seq, H3K4me3, H3K27ac, H3K4me1 and H3K27me3 peaks, in function of whether these peaks are differentially enriched between samples of the Neuron and THP-1 series. Only nonDE-DIs targeting loci assigned to cluster A (for ATAC-seq and H3K4me3), cluster B (for H3K27ac and H3K4me1) and cluster C (for H3K27me3) in at least one sample of the comparison were considered. nonDE-DIs are grouped based on their differential frequency, as defined in Figure 43, and non-DIs are shown for reference. The radius of each wedge represents the enrichment (representation factor) of a given DNA target category within an interaction group, relative to its representation among all interactions targeting cluster A, B or C loci. The black circles indicate a representation factor of 1, so that wedges encompassed within them depict an under-represented category and wedges that extend beyond them depict an overrepresented category. Significance was calculated by hypergeometric test; *p value < 0.05; **p value < 0.01; ***p value < 0.001.

**Figure 54:**
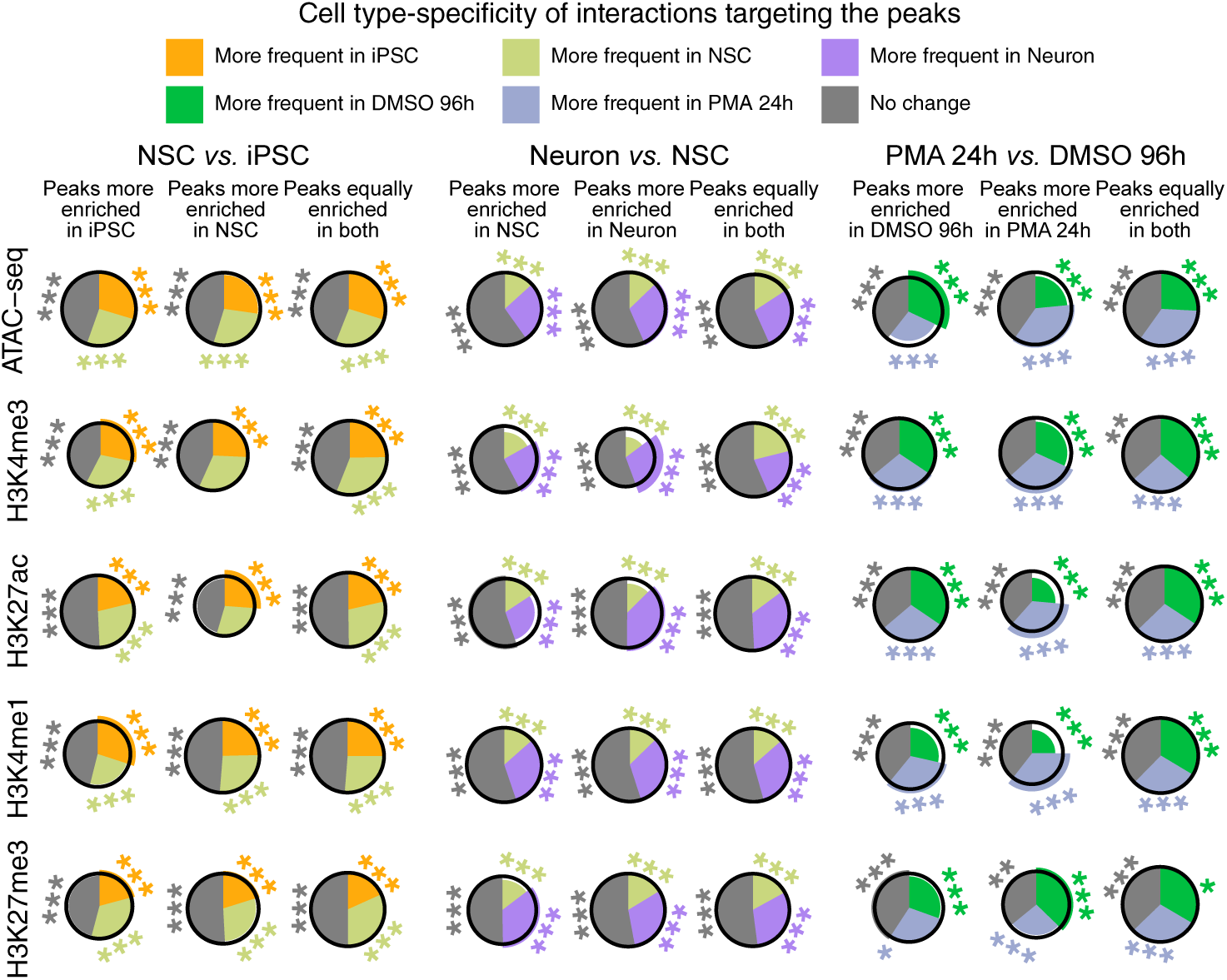
Distribution nonDE-DIs targeting ATAC-seq, H3K4me3, H3K27ac, H3K4me1 and H3K27me3 peaks, in function of the sample where these interactions are the most frequent in the Neuron and THP-1 series, as defined in Figure 43. Non-DIs are shown for reference. Targeted peaks are grouped based on their differential enrichment (FC < 0.5 or > 2 and FDR < 0.01, edgeR, Robinson et al., 2009). The radius of each wedge represents the enrichment (representation factor) of a given RNA-DNA interaction category within a group of targets, relative to its representation among all interactions targeting peaks of the same nature. The black circles indicate a representation factor of 1, so that wedges encompassed within them depict an under-represented category and wedges that extend beyond them depict an overrepresented category. Significance was calculated by hypergeometric test; *p value < 0.05; **p value < 0.01; ***p value < 0.001.

While histone modifications show no systematic association with the target side of the RNA-DNA interactome, we observed clear links between epigenetic state and transcription on the RNA side. Changes in H3K4me3, H3K4me1, H3K27ac and H3K27me3 enrichment correlate (or anticorrelate for H3K27me3) with expression changes at sources of DE-DIs, often more strongly than for other differentially expressed genes that do not produce chromatin-associated RNAs (Figure 55). Accordingly, loci producing RNAs that show increased transcription and interaction frequency during differentiation are more likely to concomitantly shift toward more active A/B subcompartments than sources of nonDE-DI, whereas those with decreased activity tend to shift toward less active compartments (Figure 56). Together, these observations indicate that RNA-DNA interactions are influenced by the evolution of the chromatin state at their source *via* the modulation of RNA abundance.

**Figure 55:**
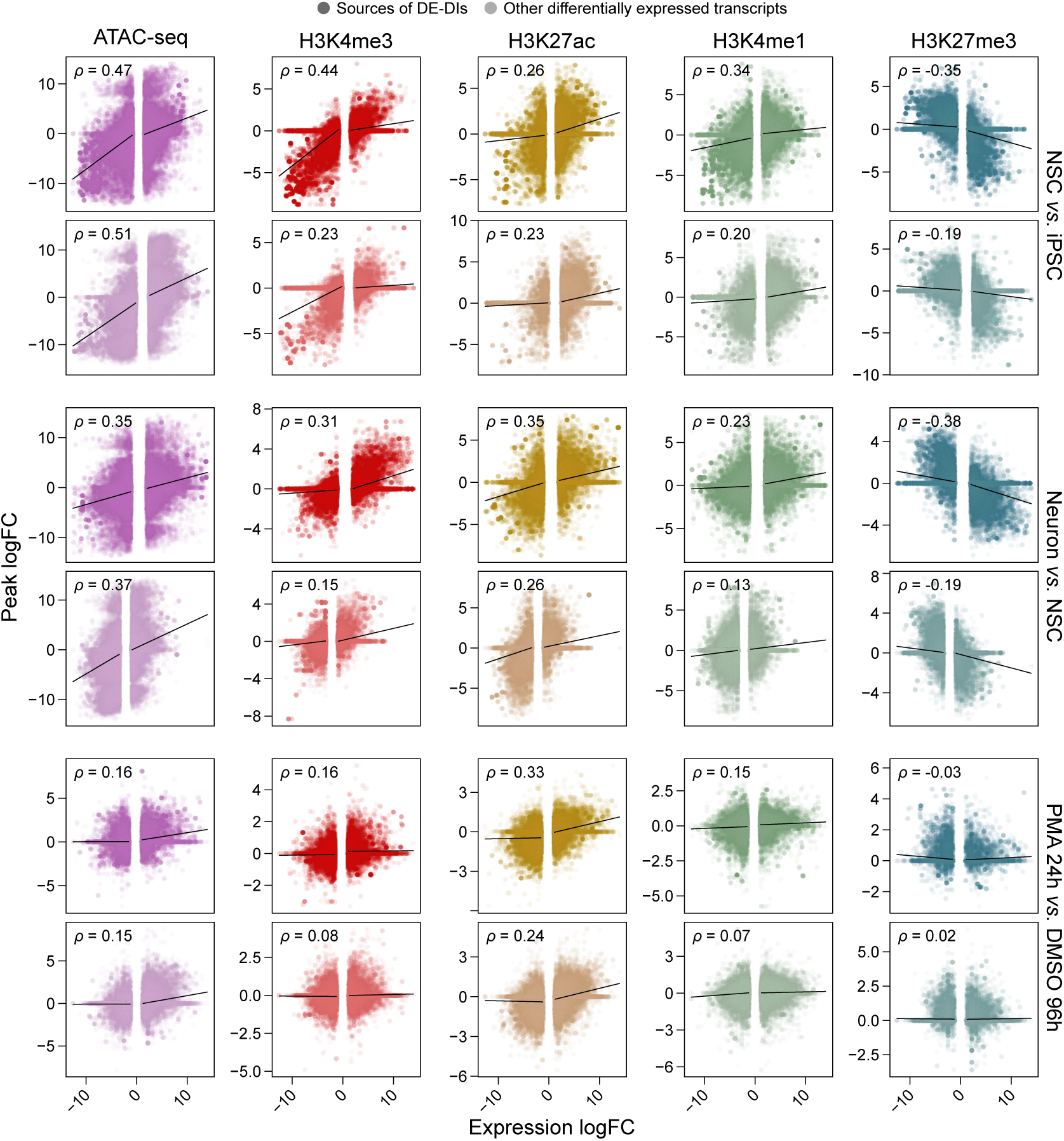
logFC (edgeR, Robinson et al., 2009) of the enrichment of ATAC-seq, H3K4me3, H3K27ac, H3K4me1 and H3K27me3 peaks at source loci of all differentially expressed SACAGE transcript isoforms (absolute logFC > 1.2, edgeR, Robinson et al., 2009) between samples of the Neuron and THP-1 series (y), in function of the logFC of the expression of these transcripts (x). Differentially expressed loci are separated in function of whether or not they are the source of DE-DIs. Black lines correspond to simple linear regressions between (x) and (y), performed independently for negative and positive values of (x). Spearman’s correlation coefficients (ρ) are indicated in the top left corner of each panel.

**Figure 56:**
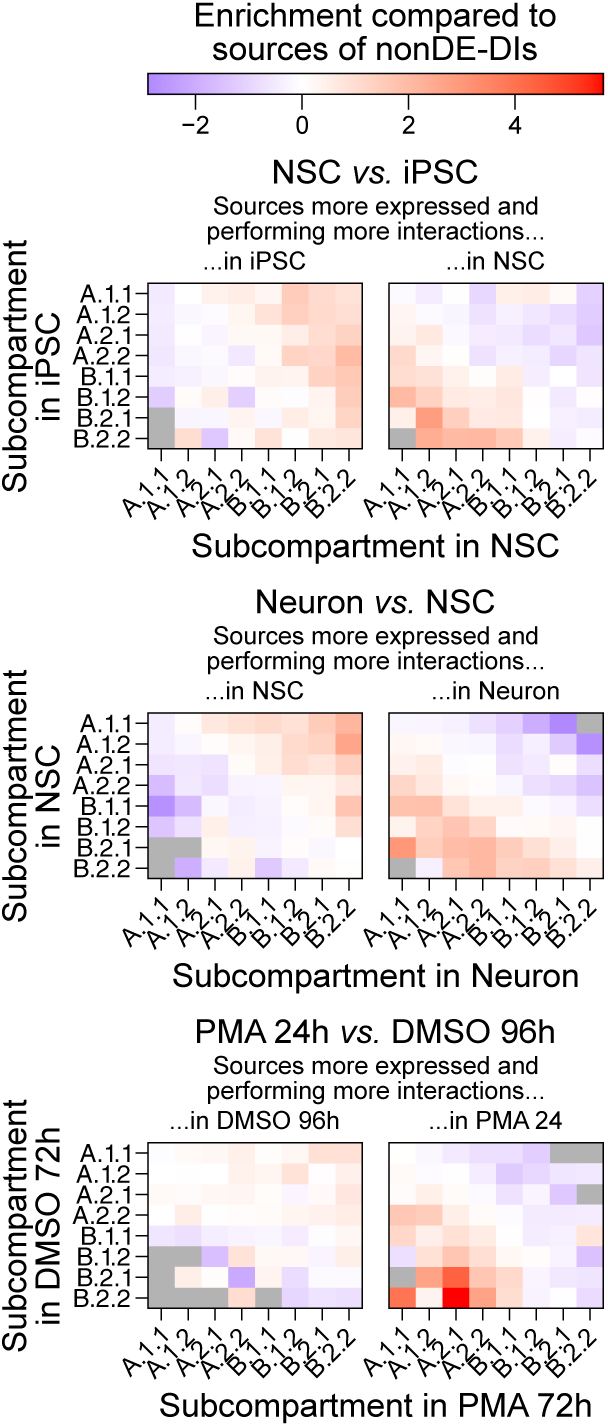
Enrichment (log of ratio) for each possible change in A/B subcompartment at the source loci of DE-DIs compared to those of nonDE-DI, for all sample comparisons in the Neuron and THP-1 series.

Interestingly, this abundance may also be modulate by other, “upstream” RNA-DNA contacts. Indeed, the TSS region of sources of DE-DIs are themselves significantly more often targeted by non-self DIs compared to the TSS region of sources of nonDE-DIs (Figure 57). Except for the Neuron *versus* NSC comparison, the dynamics of these “upstream” DIs tend to mirror those of their downstream DE-DIs: for instance, “upstream” DIs targeting the TSS of RNAs interacting more frequently in iPSC than NSC are enriched in interactions that are also more frequent in iPSC, and are depleted in interactions that are more frequent in NSC. This enrichment persists when restricting the analysis to “upstream” *trans* interactions, indicating that it cannot be solely imputed to increased local contacts within co-regulated regions. Consistent with several known instances of chromatin-associated RNAs regulating the expression of their target (Leisegang *et al*., 2024; Mangiavacchi *et al*., 2023; Oksuz *et al*., 2023; Statello *et al*., 2020), these results suggest that DNA-interacting RNAs can indirectly control the frequency of other, “downstream” RNA-DNA interactions.

**Figure 57:**
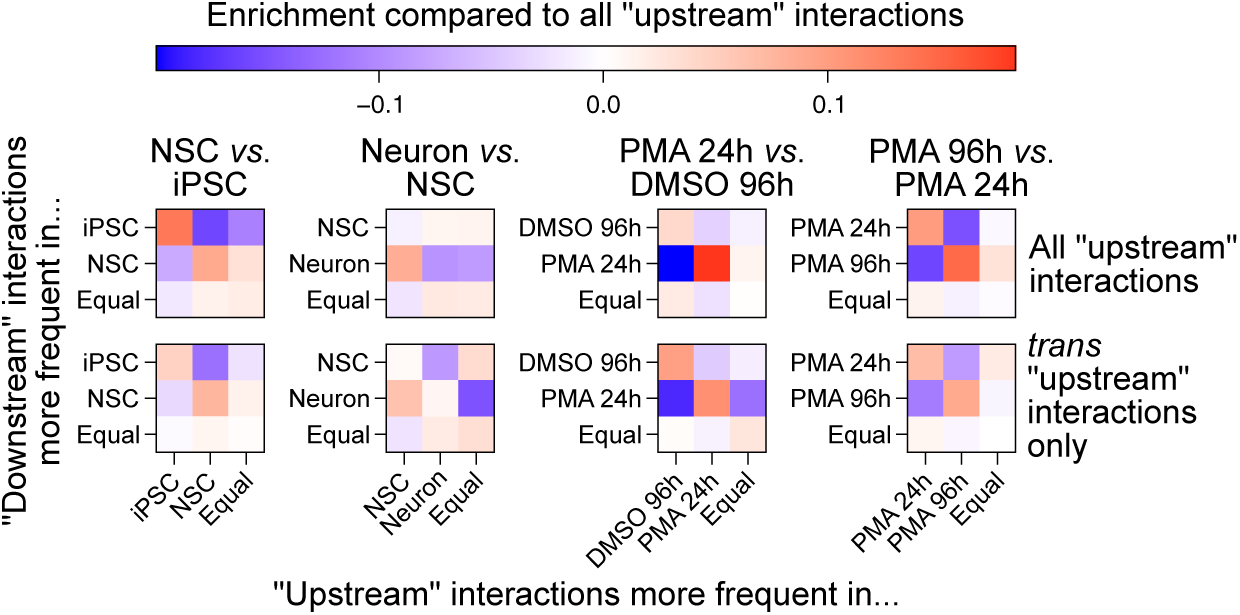
Enrichment (log of representation factor) of non-self RNA-DNA interactions (“upstream” interactions) targeting the TSS ±500 bp region of sources other DIs (“downstream” interactions), in function of whether these “upstream” interactions are themselves DIs, as defined in Figure 43. “Downstream” interactions are grouped based on whether or not they are DE-DIs, as depicted in Figure 45. All enrichment or depletion values are significant (p value < 0.001, hypergeometric test).

### RNA-DNA interactions can modulate the expression of their targets

The idea that RNA-DNA interactions can influence each other suggests an interactome architecture reminiscent of classical gene regulatory networks. However, since transcripts with high expression levels may exhibit increased chromatin contact simply because of their abundance (Figure 45), distinguishing whether enhanced targeting during differentiation reflects functional relevance or is merely driven by expression-related background interactions remains challenging. To overcome this obstacle, we once again employed a network-centered approach to investigate the potential transcriptional regulatory roles of RNA-DNA interactions, which was facilitated by the high reproducibility of the interactome networks obtained across samples in the Neuron and THP-1 series (Figure 58; Supplementary Table 6). Within this framework, we employed betweenness centrality, a topological metric that quantifies how often a node lies on the shortest paths between other nodes in the network (Potapov *et al*., 2005). Indeed, because RNA-DNA interactomes are ultra-small-world networks (Figure 7; Supplementary Table 4), source RNAs with high betweenness could coordinate signals across multiple regulatory environments, acting as hubs linking distant chromatin domains (Narang *et al*., 2015; Potapov *et al*., 2005). Importantly, the betweenness rank of a transcript only weakly correlates with its expression rank measured by CAGE (Figure 59) and shows little change when constructing the interactome networks using solely *trans* interactions (Figure 60), especially in the later stages of the two time courses where this type of contacts becomes more prevalent (Figures 18 and 20). These observations imply that the most pivotal transcripts in the RNA-DNA interaction networks are not necessarily the most expressed ones, that they perform a substantial number of their interactions in *trans*, and that *trans* interactions become increasingly important for the connectivity of the RNA-DNA interactome as cells specialize.

**Figure 58:**
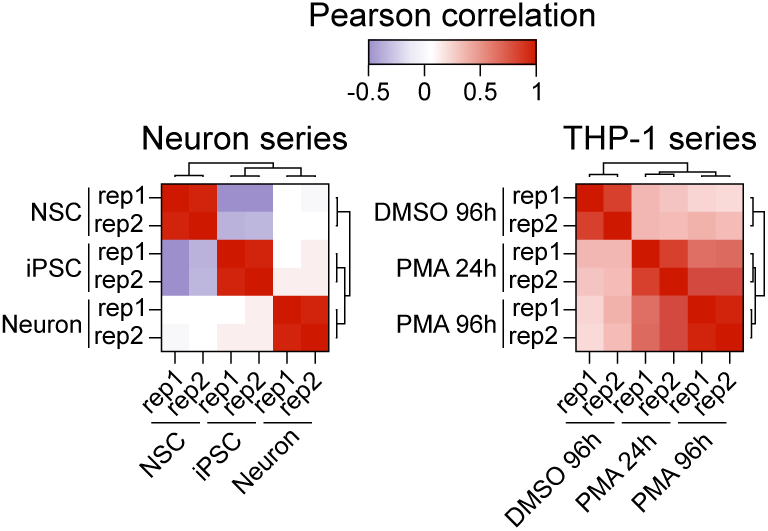
Pearson correlation of the degree of all common nodes between networks generated from all non-self RNA-DNA interactions detected in each RADICL-seq replicate of the Neuron and THP-1 series.

**Figure 59:**
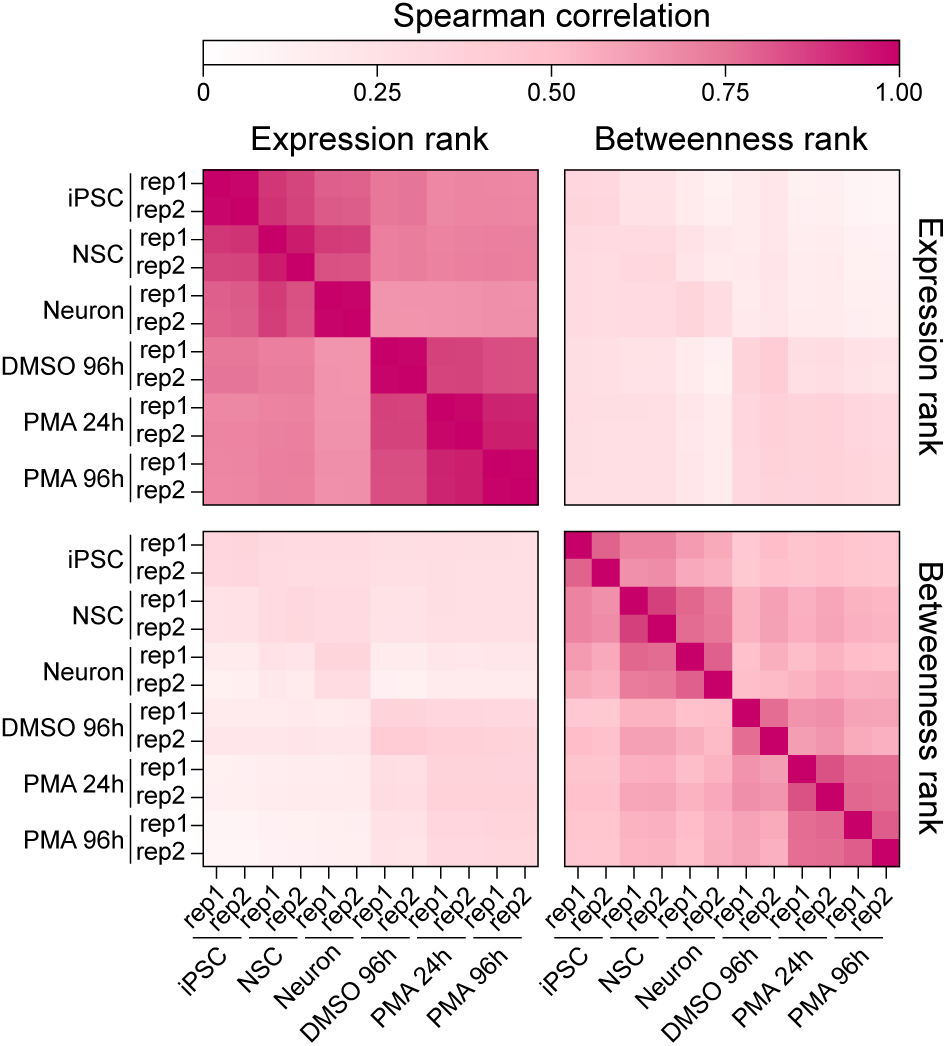
Spearman correlation of the betweenness rank and expression rank of all source RNA nodes between networks generated from all non-self RNA-DNA interactions detected in each RADICL-seq replicate of the Neuron and THP-1 series.

**Figure 60:**
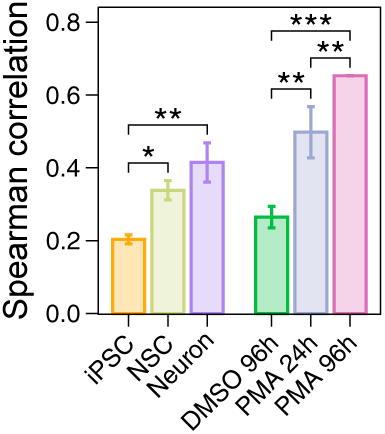
Spearman correlation of the betweenness rank for all nodes in networks generated from all non-self RNA-DNA interactions detected in each RADICL-seq replicate of the Neuron and THP-1 series, and networks generated from trans interactions only. Stars indicate significant differences between samples when comparing their Fisher z-transformed correlation coefficients (t-test with Benjamini-Hochberg correction; *p value < 0.05; **p value < 0.01; ***p value < 0.001).

To assess whether these central RNAs may have a role in regulating the expression of their targets, we focused on transcripts that have both a high betweenness rank in the first or last sample of the two time courses and a low betweenness rank in the opposite sample; these transcripts are hereinafter referred to as iPSC-highBW (26,020 transcript isoforms), Neuron-highBW (26,420 transcript isoforms), DMSO 96h-highBW (41,315 transcript isoforms) and PMA 96h-highBW sources (6,563 transcript isoforms; Figures 61 and 62). This choice was motivated by the significant enrichment in regions located within ±500 base pairs (bp) of a TSS cluster among the targets of each of these sets of highBW sources (Figure 63), suggesting their regulatory potential. On average and per cell type, a given highBW RNA targets a couple dozen TSS clusters that all tend to be located in the same region, as only ∼1.76% of RNAs have more than half of their targets further away than 10 Mb from each other, and this targeting occurs predominantly *via cis* interactions (Figures 64 to 66). TSS clusters showing significantly higher activity in a given sample are over-represented among TSSs targeted by RNAs that have high betweenness in that same sample (Figures 67 and 68). In particular, differentially active TSSs clusters that are subjected to highBW-RNA-mediated DIs show stronger modifications of their CAGE signal than those that are equivalently targeted between two samples (Figures 69 and 70), implying that their activity is modulated by the change in frequency of the RNA-DNA interaction targeting them. These results thus generalize the concept of regulatory RNA-DNA contacts, suggested by other authors (Calandrelli *et al*., 2020; Hu *et al*., 2025; L. Li *et al*., 2021; Limouse *et al*., 2023; Yip *et al*., 2022), to thousands of novel RNAs.

**Figure 61:**
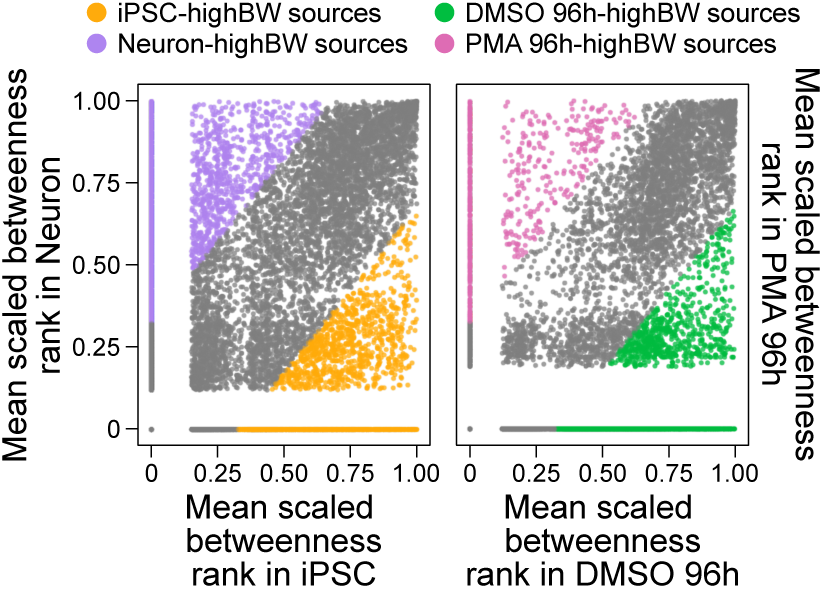
Comparison of the betweenness centrality of every source locus in networks generated from all non-self RNA-DNA interactions, in Neuron versus iPSC, and in PMA 96h versus DMSO 96h. Each source locus was ranked by its betweenness value in each replicate network, then ranks were scaled from 0 to 1 and averaged between replicates. RNAs with a mean scaled rank difference ≥0.325 are defined as “high betweenness” (highBW) sources.

**Figure 62:**
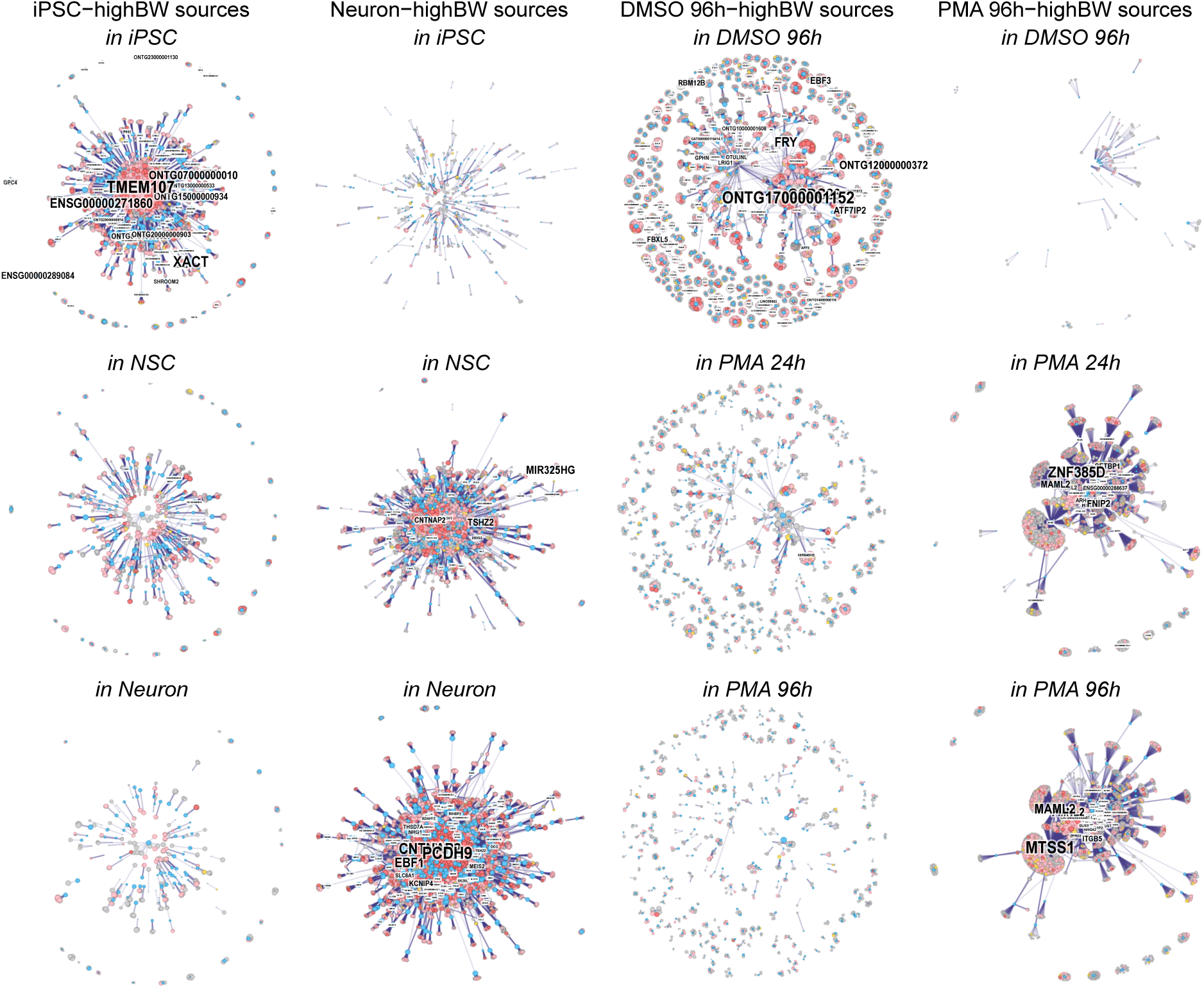
Undirected graphs depicting the interaction networks performed by a subset of iPSC-highBW and Neuron-highBW RNAs on the one hand, and of DMSO 96h-highBW and PMA 96h-highBW RNAs on the other hand, in each sample of the Neuron and THP-1 series, respectively. Only RNAs that belong to the top 25% betweenness rank in a given sample and have a 0 betweenness rank in the opposite sample are shown. Nodes corresponding to highBW RNAs are colored in blue, and those corresponding to 25-kb target windows are colored in pink, yellow or red if they contain at least one promoter, one enhancer or both, respectively. The size of nodes is proportional to their degree in the whole network. The names of sources belonging to the 35% top highBW RNAs depicted in each graph are indicated; the size of the label is proportional to their betweenness rank in the whole network. Node positions were computed as described in “Methods”.

**Figure 63:**
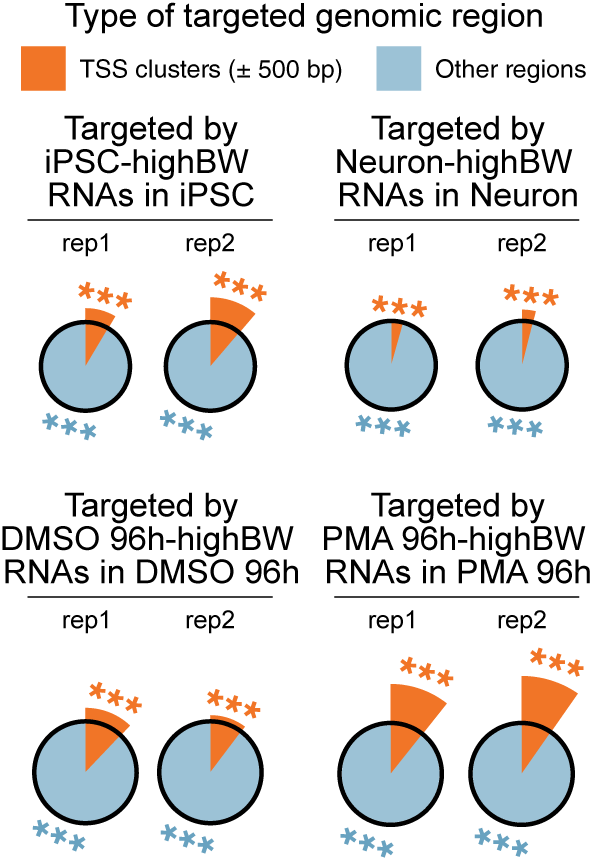
Distribution of RADICL-seq reads for which the RNA fragment corresponds to a iPSC-highBW, Neuron-highBW, DMSO 96h-highBW or PMA 96h-highBW source, in function of whether their DNA fragment maps within 500 bp of a TSS cluster, in each replicate of the Neuron and THP-1 series. The radius of each wedge represents the enrichment (representation factor) of a given target region category among all targets of highBW sources in a replicate, relative to its representation among all RADICL-seq DNA reads from that replicate. The black circles indicate a representation factor of 1, so that wedges encompassed within them depict an under-represented category and wedges that extend beyond them depict an overrepresented category. Significance was calculated by hypergeometric test; ***p value < 0.001.

**Figure 64:**
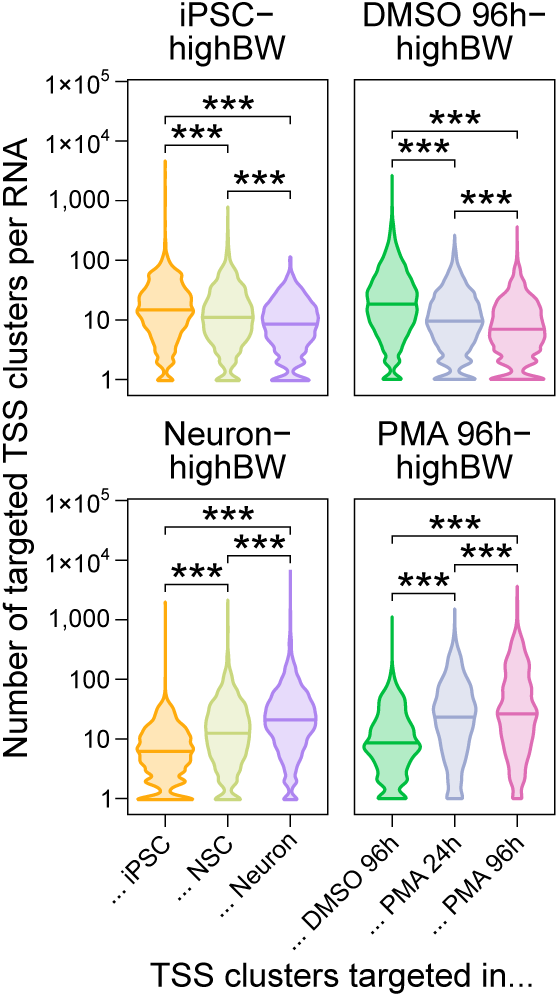
Distribution of iPSC-, Neuron-, DMSO 96h- and PMA 96h-highBW RNAs in function of the number of distinct TSS clusters that they target, in each sample of the Neuron and THP-1 series. Stars indicate significant differences between samples for a given set of highBW sources (Student’s t-test; ***p value < 0.001).

**Figure 65:**
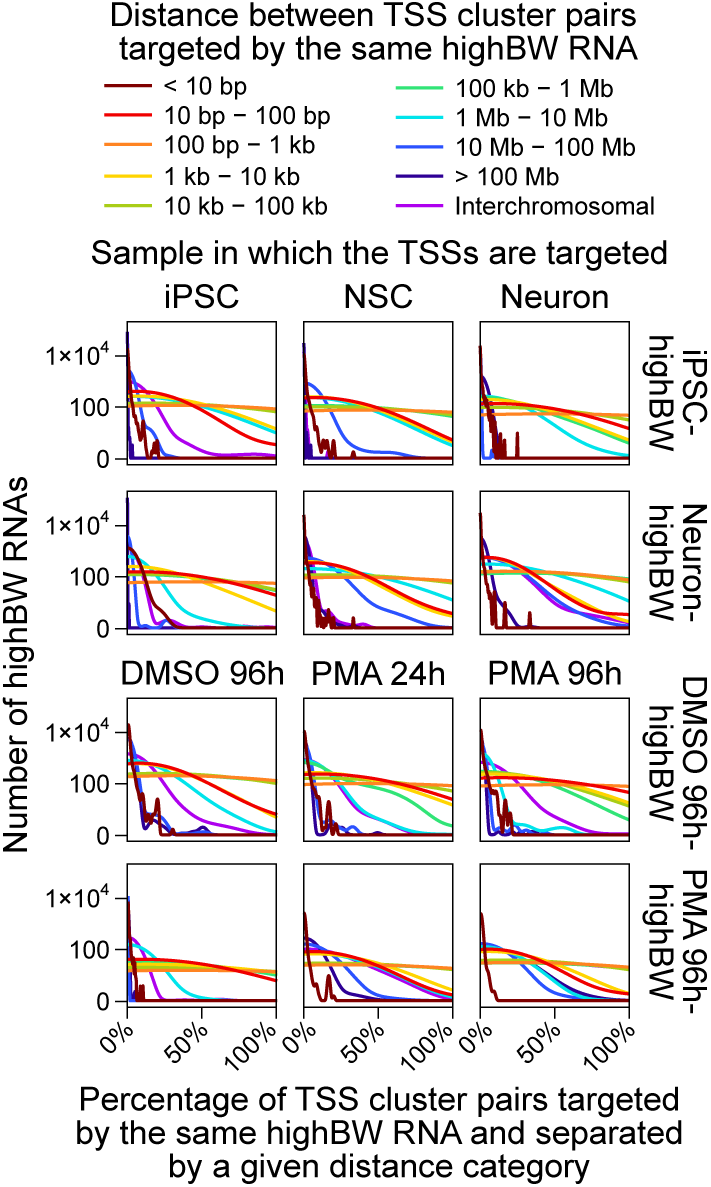
Distribution of iPSC-, Neuron-, DMSO 96h- and PMA 96h-highBW RNAs in function of the percentage of their targeted TSSs that are located at different distances from each other, in each sample of the Neuron and THP-1 series.

**Figure 66:**
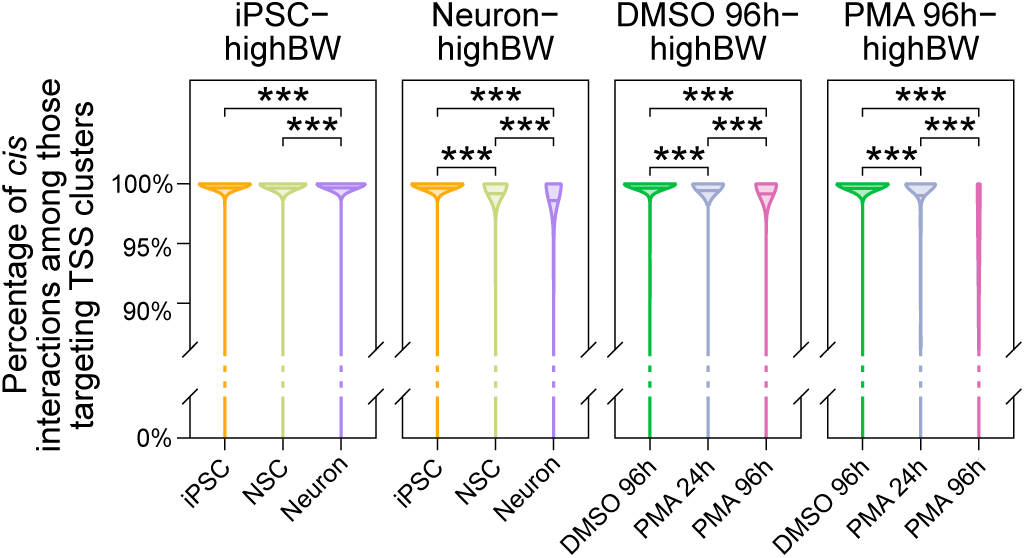
Distribution of iPSC-, Neuron-, DMSO 96h- and PMA 96h-highBW RNAs in function of the percentage of their TSS cluster-targeting interactions that are performed in cis, in each sample of the Neuron and THP-1 series. Stars indicate significant differences between samples for a given set of highBW sources (Student’s t-test; ***p value < 0.001).

**Figure 67:**
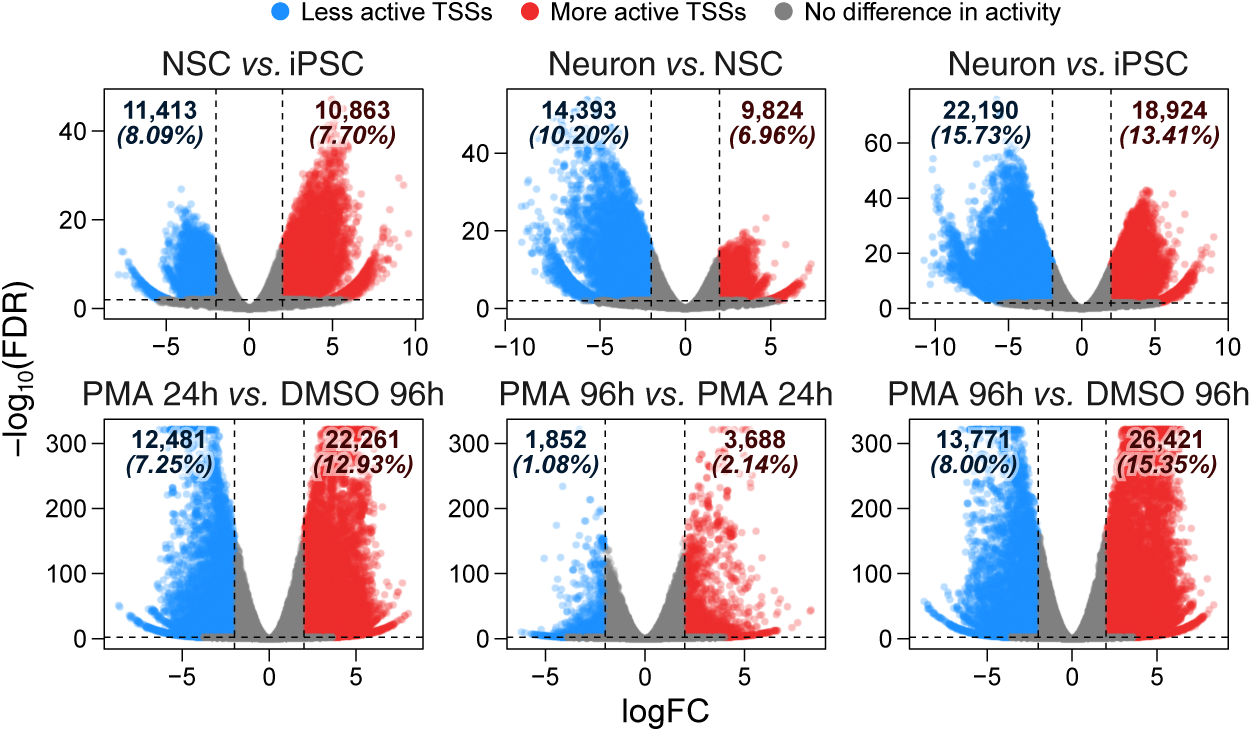
Differential expression of all TSS clusters detected by ssCAGE between each sample of the Neuron and THP-1 series. For each pairwise comparison, TSS clusters with a logFC < −2 or > 2 and an FDR < 0.01 (edgeR; Robinson et al., 2009) are considered as significantly less (blue) or more (red) active, respectively, and their number is indicated at the top of each panel.

**Figure 68:**
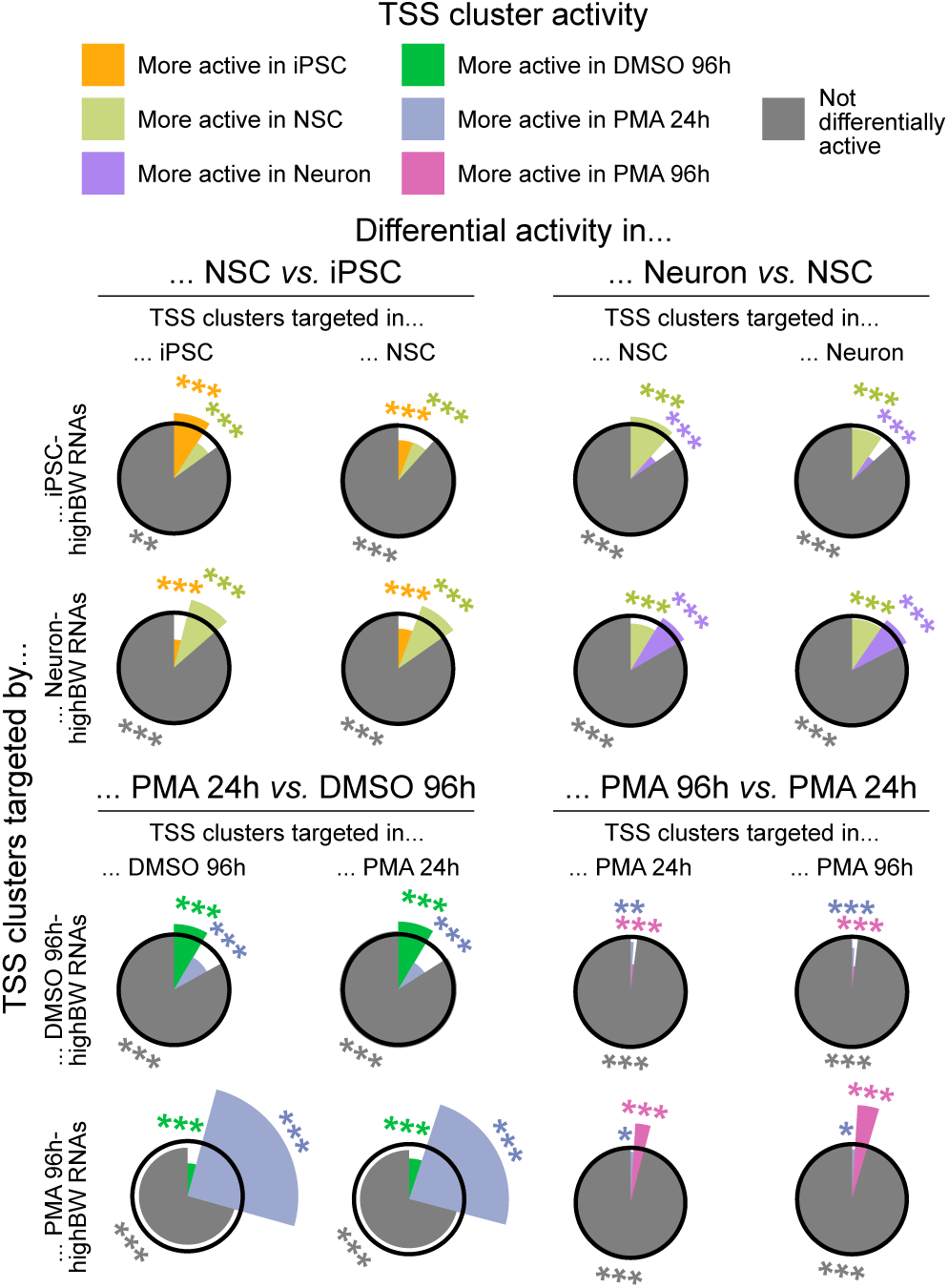
Distribution of TSS clusters targeted by iPSC-highBW, Neuron-highBW, DMSO 96h-highBW or PMA 96h-highBW sources, in function of whether they are more or less active in each sample of the Neuron and THP-1 series. Transcript-isoform-to-TSS-cluster interactions detected in each RADICL-seq replicate were pooled together for each sample. The radius of each wedge represents the enrichment (representation factor) of a given category of TSS clusters among those targeted by highBW RNAs in a sample, relative to its representation among all TSS clusters targeted by at least one RNA in that sample. The black circles indicate a representation factor of 1, so that wedges encompassed within them depict an under-represented category and wedges that extend beyond them depict an overrepresented category. Significance was calculated by hypergeometric test; *p value < 0.05; **p value < 0.01; ***p value < 0.001.

**Figure 69:**
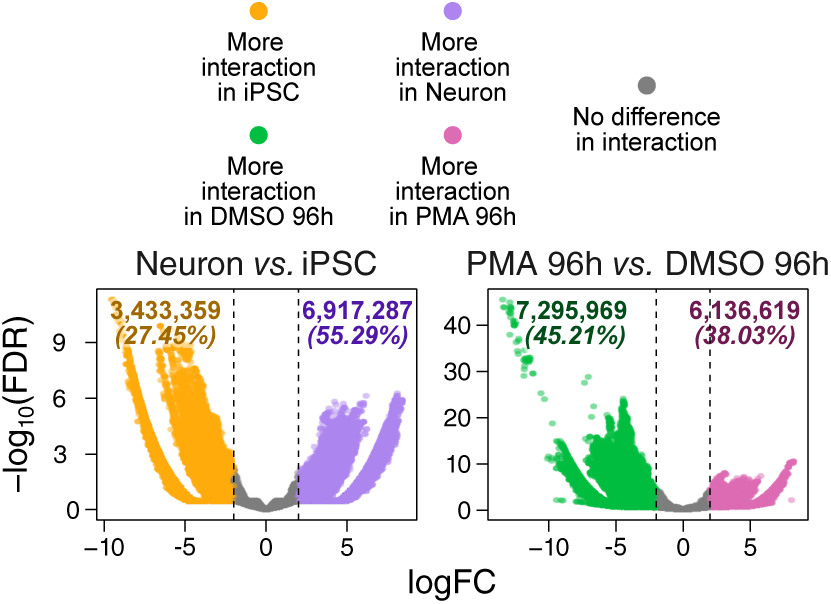
Changes in RNA-DNA interaction frequency between samples of the Neuron and THP-1 series, at the transcript-isoform-to-TSS-cluster level. DIs are defined as those having an absolute logFC > 2 (edgeR, Robinson et al., 2009) and their numbers are indicated at the top of each panel.

**Figure 70:**
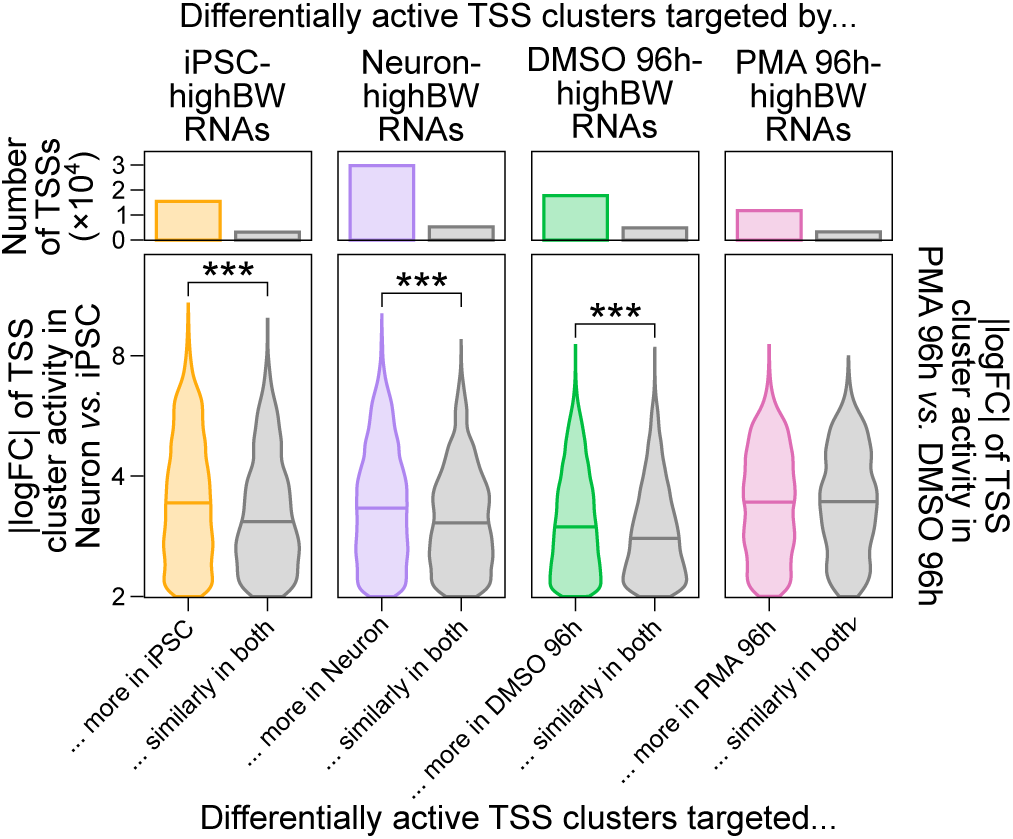
Distribution of the absolute logFC for the activity of all differentially active TSS clusters targeted by highBW RNAs, in Neuron versus iPSC (left) or in PMA 96h versus DMSO 96h (right). TSS clusters are separated in function of whether or not they are concomitantly more targeted (interaction logFC > 2; edgeR, Robinson et al., 2009) by these highBW RNAs. Stars indicate significant differences between TSS clusters that are more targeted in a given sample and those that are not (Wilcoxon signed-rank test; ***p value < 0.001). The total number of target TSS clusters in each category is indicated in the upper bar plots.

HighBW-RNA-targeted TSS clusters can be either significantly more or significantly less active in the sample where they are the most frequently targeted (Figure 71), indicating that our results are unlikely to merely reflect the presence of co-regulated chromatin domains encompassing both the source RNAs and their proximal targets. Furthermore, in most cases, all TSS clusters targeted by the same highBW RNA are differentially active in the same direction (Figure 72), meaning that highBW RNAs may function either as enhancers or repressors of transcription, but rarely both.

**Figure 71:**
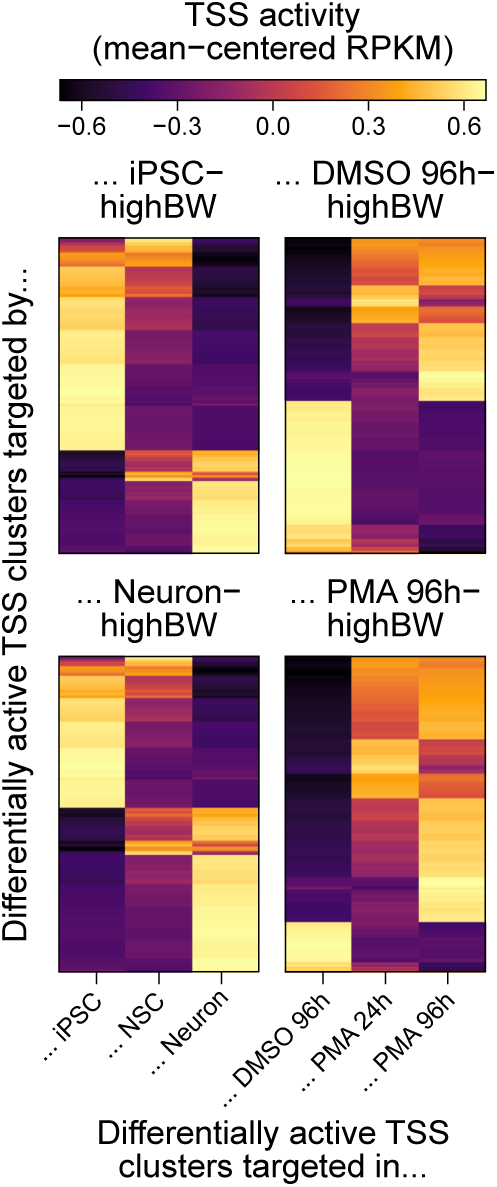
Mean expression (RPKM) of differentially active TSS clusters that are significantly more targeted by iPSC-, Neuron-, DMSO 96h- or PMA 96h-highBW RNAs, in the sample where these RNAs have the highest betweenness. RPKM values were averaged between ssCAGE replicates in each sample, then mean-centered for each individual TSS cluster across all samples in each series, before being hierarchically grouped using Ward’s method (Ward, 1963).

**Figure 72:**
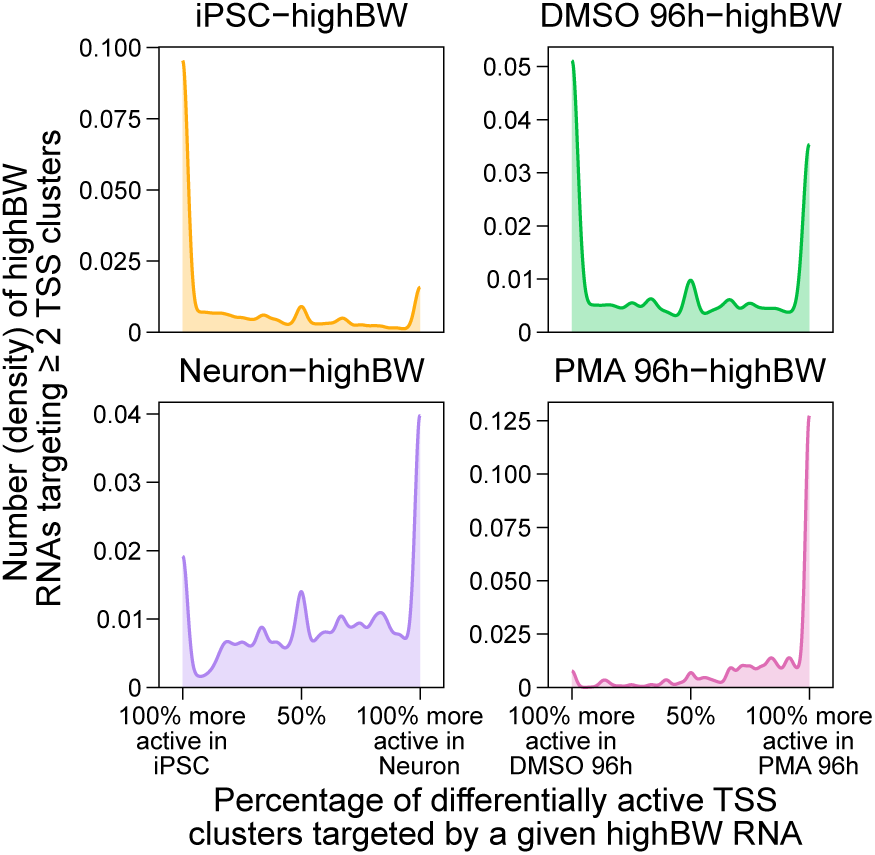
Distribution of iPSC-, Neuron-, DMSO 96h- and PMA 96h-highBW RNAs in function of the percentage of their targeted TSS clusters that are significantly more active in iPSC/DMSO 96h or in Neuron/PMA 96h. Only TSS clusters that are both significantly differentially active between two samples and that are significantly more targeted in the sample where the RNA has the highest betweenness were considered.

To investigate the functional role of the transcriptional regulation mediated by RNA-DNA interactions, we performed a Gene Ontology (GO) term enrichment analysis both on highBW RNAs and on differentially active TSS clusters that are more frequently targeted by these transcripts in the sample where they have the highest betweenness (Figure 73). We found that iPSC- and DMSO 96h-highBW RNAs as well as the TSSs that they appear to positively regulate are enriched in GO annotations related to cell cycle and chromatin organization. In contrast, the TSSs that they appear to negatively regulate are more associated with GO terms related to transcription (Figure 73). Conversely, PMA 96h-highBW sources and their targets are enriched in GO annotations related to blood cells and immunity (Figure 73). Even more strikingly, Neuron-highBW sources are enriched in genes annotated with neuron-specific terms and appear to promote the expression of genes involved in neuron development while repressing genes participating in other developmental pathways (Figure 73).

**Figure 73:**
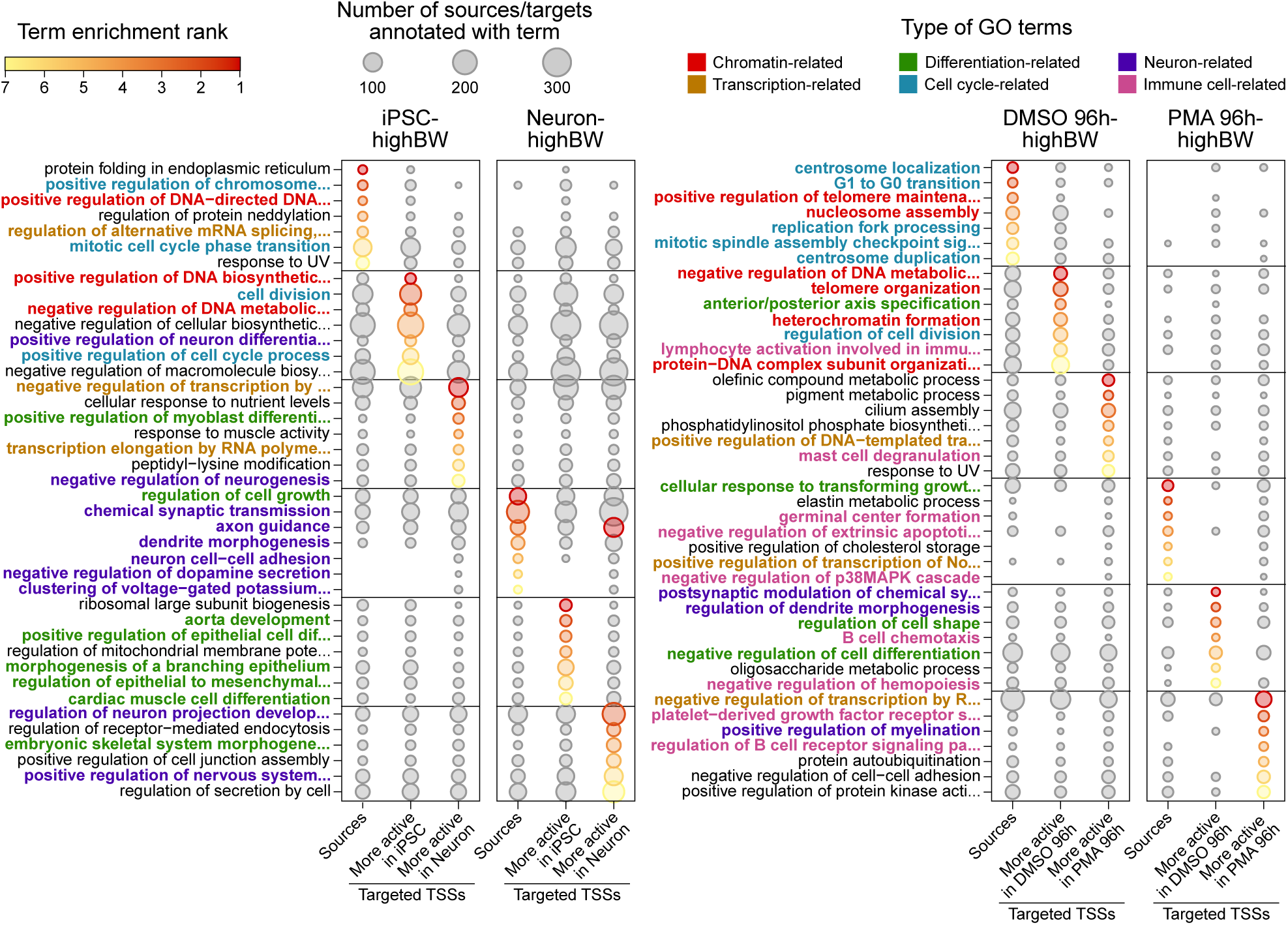
Top 7 most enriched (Fisher’s exact test, topGO, Alexa & Rahnenführer, 2006) gene ontology (GO) terms among the annotations of iPSC-, Neuron-, DMSO 96h- and PMA 96h-highBW RNAs, as well as among those of the genes associated with their target TSS clusters. Only TSS clusters that are both significantly differentially active between two samples and that are significantly more targeted in the sample where the RNA has the highest betweenness were considered. HighBW RNAs annotations were compared to those of all chromatin-associated RNAs detected in the same sample; highBW-RNA-targeted and differentially active TSS clusters were compared to all similarly differentially active TSS clusters in the same samples. The color gradient represents the significance rank of the enrichment of a given GO term and the size of the dots represent the number of genes annotated with each GO term within each group. GO terms were manually categorized and are colored accordingly.

Aiming to generalize these findings, we performed two types of enrichment analyses based on data from genome-wide association studies (GWAS), which we applied to all RNAs interacting with TSS clusters as well as their targeted regulatory regions. This assessment was performed across all cell types of our collection, except for the AML samples for which no CAGE data, and thus no TSS information, was generated. Our first approach leveraged GWAS results from European populations (European 1000 Genomes Project and UK Biobank) to evaluate the enrichment of polygenic trait loci using a linkage disequilibrium score regression applied to specifically expressed genes (LDSC-SEG; Finucane *et al*., 2018) for source loci, and a stratified linkage disequilibrium score regression (S-LDSC; Finucane *et al*., 2015) for target loci (Figure 74). For both sources and targets of RNA-DNA interactions, we detected a high enrichment for immune diseases in the T cell samples, with stronger values in the activated cells (Figure 74). Furthermore, the NSC and Neuron samples showed enrichment for psychological-related traits, such as Alzheimer’s disease and years of education, among RNA-targeted TSS clusters (Figure 74). Our second method consisted in calculating the odds ratios of genomic overlap between these same sources and target loci with phenotype-associated genetic variants, this time using results reported by the MRC IEU OpenGWAS project (Elsworth *et al*., 2020). After ensuring that these regions are enriched in variant hits relative to the genome-wide expectation (Figure 75), we found a significant over-representation of phenotypes related to immunity for either or both the RNA and DNA sides of interactions detected in each sample of our two immune cell series, which was notably absent in all other cell types (Figure 76). Results for brain-related traits were less conclusive, as they were either non-significant or depleted from source and target regions in most samples, but remained significantly enriched among the source loci detected in Neuron (Figure 76).

**Figure 74:**
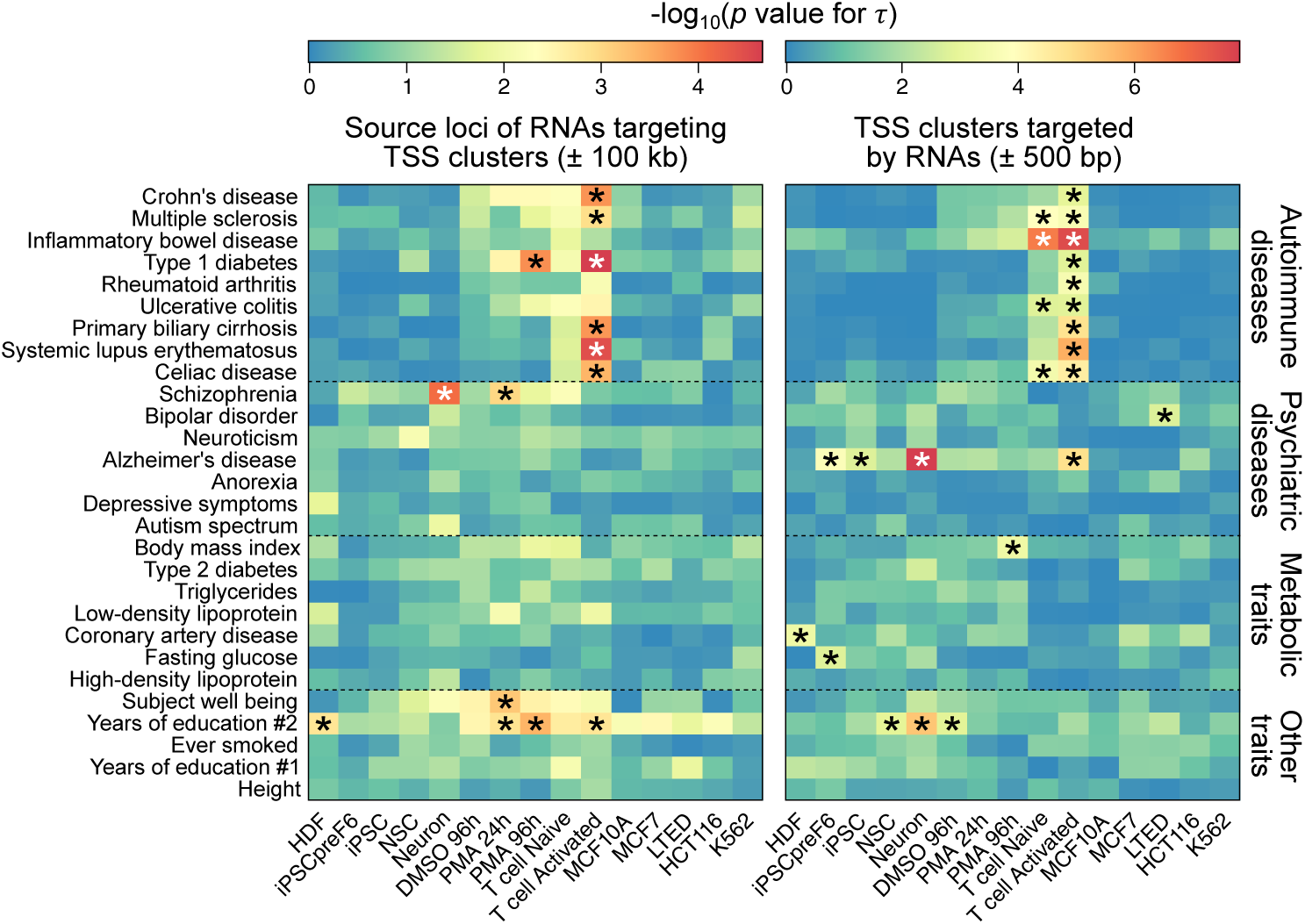
SNP heritability enrichment of complex traits from GWAS results in European populations determined in 15 samples of our RADICL-seq collection by: (left) an LDSC-SEG-like analysis (Finucane et al., 2018) on the source region of RNAs interacting with at least one TSS cluster, extended by ±100 kb; (right) an S-LDSC analysis (Finucane et al., 2015) on regions containing RNA-targeted TSS clusters ±500 bp. Stars indicates Benjamini-Hochberg’s FDR < 0.05 for LDSC-SEG or for the p value of S-LDSC’s coefficient (τ).

**Figure 75:**
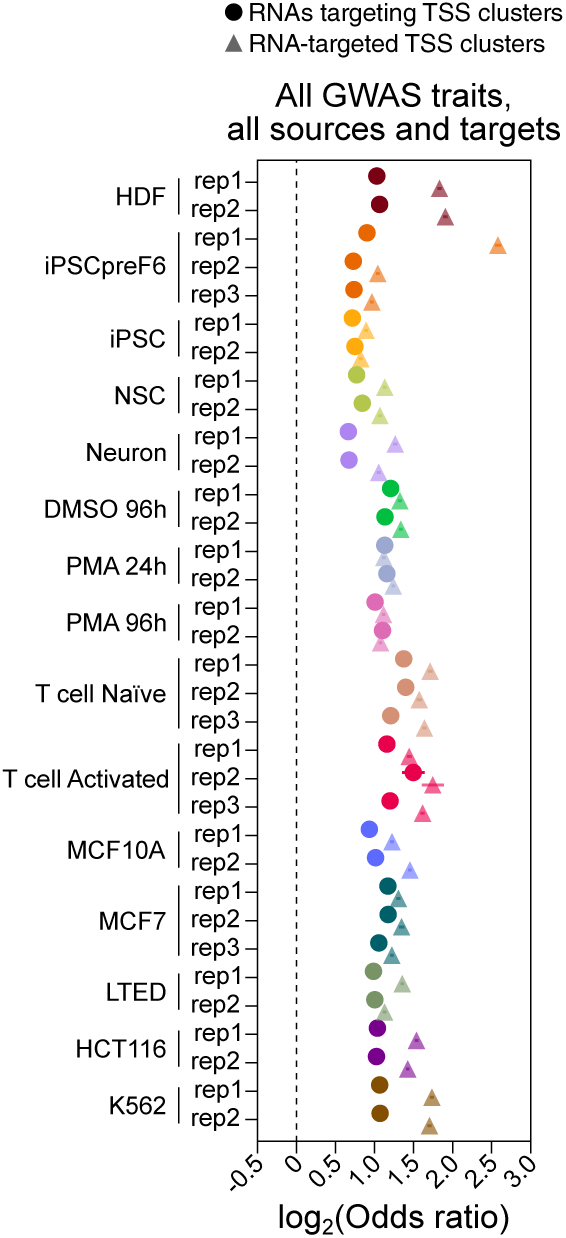
Odds ratios of genomic overlap between source (circles) or target (triangles) loci of all TSS cluster-targeting interactions detected in 15 samples of our RADICL-seq collection, and phenotype-associated genetic variants reported by the MRC IEU OpenGWAS project. Odds ratios were calculated relative to a permuted genome-wide expectation. Circles and triangles show the odd ratio point estimates while horizontal lines indicate the 95% confidence interval from a Fisher’s exact test.

**Figure 76:**
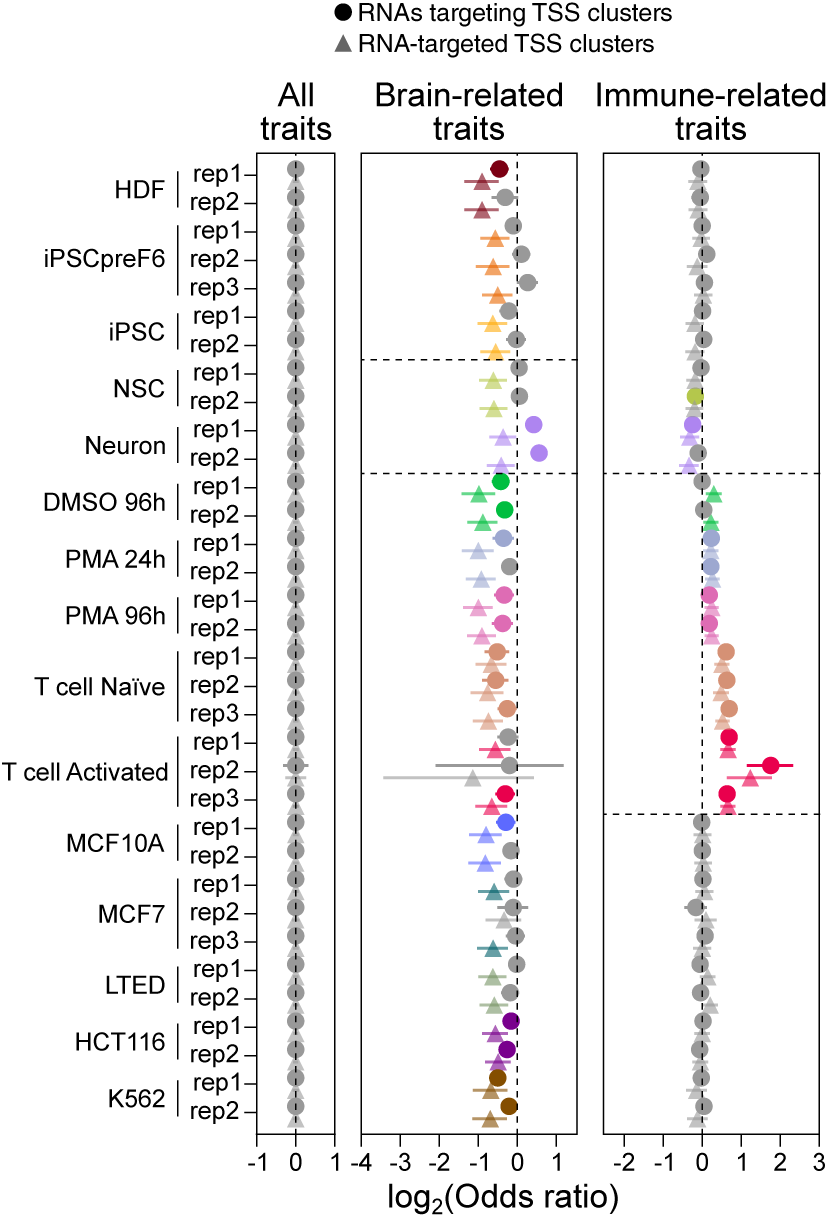
Odds ratios of genomic overlap between source (circles) or target (triangles) loci of all TSS cluster-targeting interactions detected in 15 samples of our RADICL-seq collection, and phenotype-associated genetic variants reported by the MRC IEU OpenGWAS project. Odds ratios are shown relative to the genomic overlap of all phenotype-associated variants. In the middle and right panels, only variants associated with brain-related or immune-related traits, respectively, were considered. Circles and triangles show the odd ratio point estimates while horizontal lines indicate the 95% confidence interval from a Fisher’s exact test. Non-significant results (i.e. those for which the log_2_-transformed confidence interval crosses 0) are shown in grey, while all other significant results are shown in color.

Combined with our GO term enrichment analysis and results from other studies (Calandrelli *et al*., 2020; Shu *et al*., 2024; W. Xu *et al*., 2021; Yip *et al*., 2022), these two integrations of our RADICL-seq data with GWAS data highlight that RNA-DNA contacts with potential regulatory roles are prone to be involved in cell type-specific processes. The disruption of these interactions may thus contribute to cellular dysfunction and the development of diseases.

### RNAs modulate expression by associating with regulatory proteins on the chromatin

Given the apparent importance of RNA-DNA interaction-mediated gene regulation for the biology of the cell, we next sought to understand the mechanisms by which this regulatory role is exerted. A preliminary answer can be obtained by examining the enrichment of known functional elements on the most prevalent classes of chromatin-associated RNAs detected in the Neuron and THP-1 series, *i.e.* transcripts derived from protein-coding genes and lncRNAs (Figure 77; Methods). The resulting enrichment profiles differ in function of the nature of the source RNA (lncRNA or protein-coding gene-derived, exonic or intronic) and the interaction distance (*cis* or *trans*; Figure 77). Chromatin-associated RNAs derived from exons of protein-coding genes notably show a robust enrichment in G-quadruplex-forming sequences and repeats across all samples, which is absent from their intronic counterparts (Figure 77). Exonic and intronic *cis*- and, to a lesser extent, *trans*-interacting lncRNA fragments are instead strongly enriched in elements associated with R-loops, RNA:DNA triplex and RBPs, similar to RNAs derived from introns of protein-coding genes (Figure 77). These results thus suggest fundamentally distinct structural bases for the chromatin contacts mediated by coding *versus* non-coding transcripts. Additionally, we observed a broad depletion in Phastcons-conserved elements across all samples and categories, further confirming that RNA-mediated chromatin contacts are dynamic and condition-responsive rather than constrained to ultra-conserved regulatory sequences (Figure 77).

**Figure 77:**
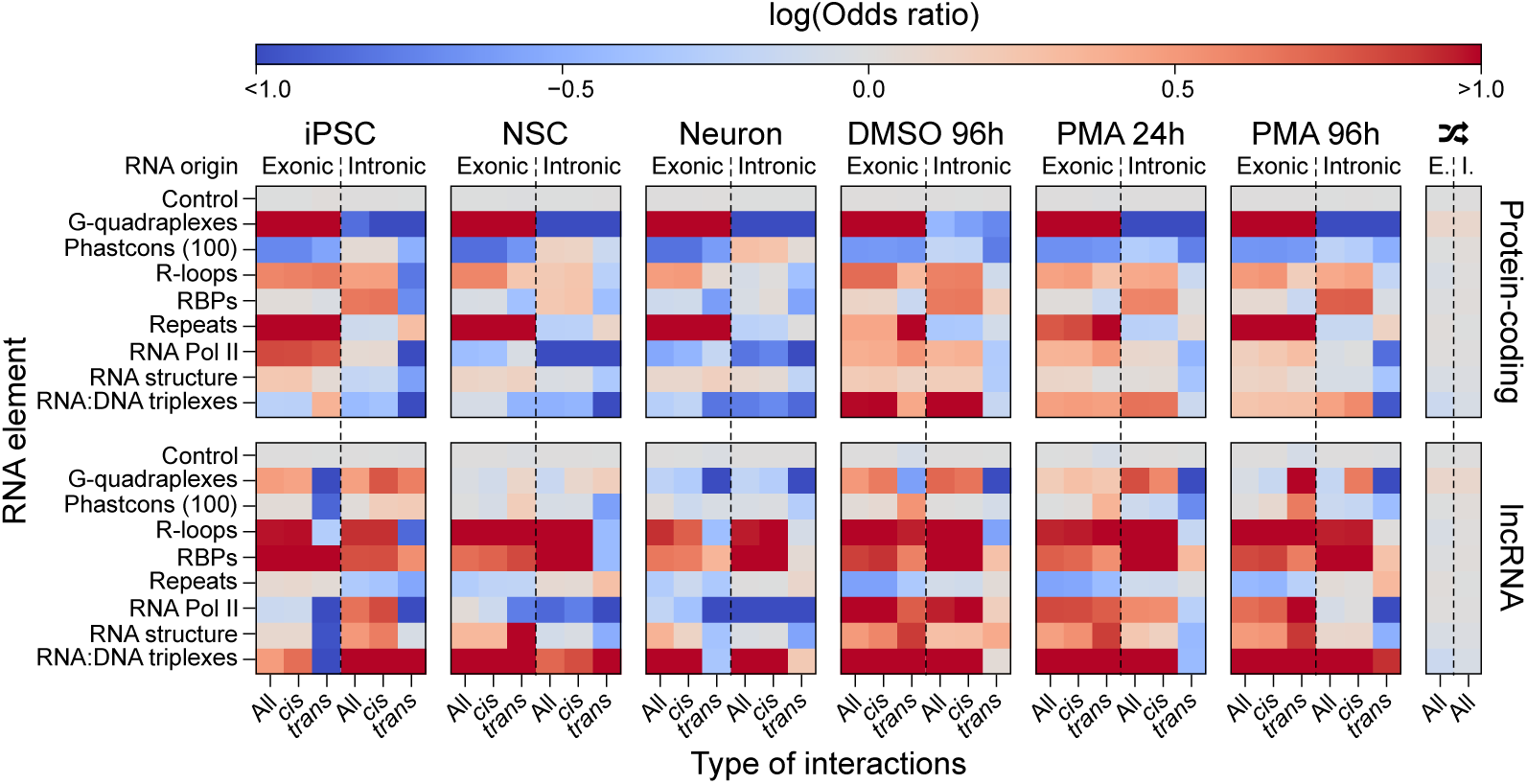
Enrichment (odds ratio over a matched background set; see “Methods”) of functional elements on all chromatin-interacting lncRNAs and RNAs derived from protein-coding genes detected by RADICL-seq in the Neuron and THP-1 series. For reference, a negative control for functional elements (consisting in randomly selected genomic regions) and a scrambled control for the lncRNAs (indicated by the twisted arrows symbol) are shown. DNA-interacting RNA fragments are separated in function of whether they originate from exonic or intronic regions of protein-coding genes or lncRNA-coding loci, and of whether they perform cis or trans interactions.

While we observed an enrichment for R-loop-associated elements among DNA-interacting lncRNAs and intronic RNAs (Figure 77), comparison of our RNA-DNA contacts with data obtained by DNA/RNA immunoprecipitation and high-throughput sequencing (DRIP-seq) in THP-1 cells (Bamezai *et al*., 2023) revealed that more than two-thirds of loci detected by both technologies correspond to targets of non-self interactions (Figure 78) that are not particularly predicted to hybridize as RNA:DNA duplexes (Figure 79). Given that the RADICL-seq protocol involves a treatment with RNAse H, which digests RNA:DNA duplexes (Methods; Bonetti *et al*., 2020; Huertas & Aguilera, 2003; Pracana *et al*., in preparation), we thus conjecture that most of the above-mentioned overlap between RADICL-seq and DRIP-seq data does not reflect *bona fide* R-loop-forming interactions.

**Figure 78:**
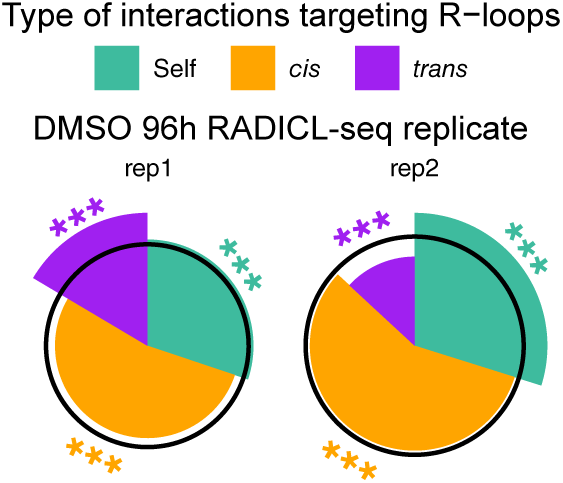
Distribution of RNA-DNA interactions targeting R-loops in function of whether they are self, cis (excluding self interactions) or trans interactions, in each replicate of the DMSO 96h sample. The radius of each wedge represents the enrichment (representation factor) of a given type of interaction among those targeting R-loops, relative to its representation among all interactions detected by RADICL-seq in that replicate. The black circles indicate a representation factor of 1, so that wedges encompassed within them depict an under-represented category and wedges that extend beyond them depict an overrepresented category. Significance was calculated by hypergeometric test; ***p value < 0.001.

**Figure 79:**
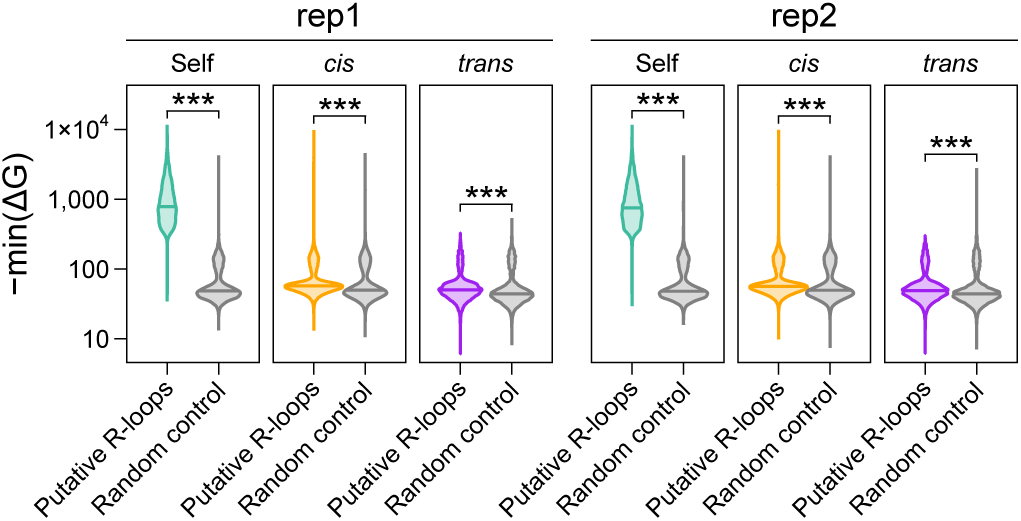
Distribution of the maximum amount of energy -ΔG predicted (RIsearch2; Alkan et al., 2017) to be released by a hybridization between RNAs and R-loop regions that are detected to interact by RADICL-seq (“putative R-loops”), in each replicate of the DMSO 96h sample. Interactions are separated in function of whether they are self interactions or are performed in cis (excluding self interactions) or in trans. For comparison, the same analysis was computed with the same RNA sequences and a set of randomly shuffled sequences equivalent to the R-loop regions targeted by these RNAs (“random control”). Stars indicate significant differences between the putative R-loops and their controls (Student’s t-test; ***p value < 0.001).

Our enrichment analysis in Figure 77 also indicated a strong enrichment for RBP-associated motifs on chromatin-associated lncRNAs and RNAs derived from introns of protein-coding genes. Performing a similar analysis at the individual RBP level revealed that DNA-interacting RNAs derived from introns of protein-coding genes or from exons of lncRNAs are highly enriched in motifs corresponding to proteins associated with the nuclear scaffold, such as SPLICING FACTOR PROLINE AND GLUTAMINE RICH (SFPQ; Knott *et al*., 2016), NON-POU DOMAIN CONTAINING OCTAMER BINDING (NONO; Knott *et al*., 2016), FUSED IN SARCOMA RNA BINDING PROTEIN (FUS; Yamaguchi & Takanashi, 2016), MATRIN 3 (MATR3; Yamaguchi & Takanashi, 2016) and SCAFFOLD ATTACHMENT FACTOR B2 (SAFB2; Hashimoto *et al*., 2012). This enrichment is more pronounced in the THP-1 series than in the Neuron series, consistent with the elevated number of interactions mediated by paraspeckle component *NEAT1* in these samples (Figure 19; Yamazaki *et al*., 2018). These RNAs are also abundant in motifs related to spliceosome- and other ribonucleoprotein particle-associated factors (Figure 80), including HETEROGENEOUS NUCLEAR RIBONUCLEOPROTEIN M and U (HNRNPM/U; Geuens *et al*., 2016), EWS RNA BINDING PROTEIN 1 (EWSR1; Lee *et al*., 2019), SPLICING FACTOR 3B SUBUNIT 1 (SF3B1; Sun, 2020), PRE-MRNA PROCESSING FACTOR 8 (PRPF8; Grainger & Beggs, 2005), U2 SMALL NUCLEAR RNA AUXILIARY FACTOR 1 and 2 (U2AF1/2; Wahl *et al*., 2009) and the SERINE/ARGININE SPLICING FACTOR (SRSF) family members (Zahler *et al*., 1992). These results indicate that interactions mediated by transcripts originating from introns of protein-coding genes and exons of lncRNAs are mostly involved in RNA processing and nuclear organization.

**Figure 80:**
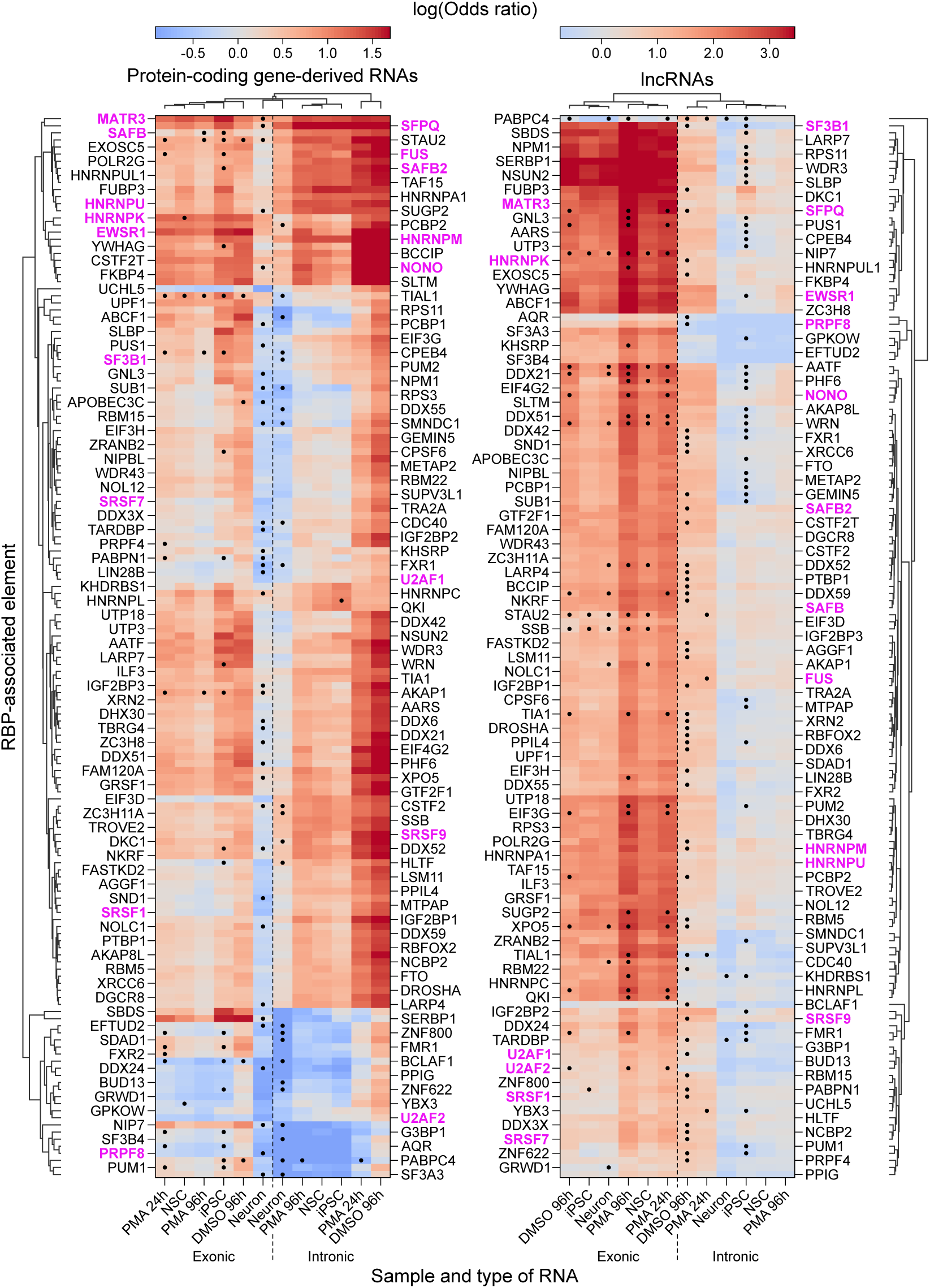
Enrichment (odds ratio of Fisher’s exact test) of RBP-associated elements on chromatin-interacting RNAs detected by RADICL-seq in the Neuron and THP-1 series. DNA-interacting RNA fragments are separated in function of whether they originate from exonic or intronic regions of protein-coding (left) or lncRNA genes (right). Dots in the heatmap indicate protein expression-driven enrichment or depletion (see “Methods”). For better visualization, the y-axes labels (corresponding to the RBP-associated elements) are split so that every other label is shown on each side of the plots. Protein names mentioned in-text are colored in magenta.

On the other hand, chromatin-associated RNAs derived from exons of protein-coding genes and introns of lncRNAs are also enriched for nuclear RBPs (e.g., EWSR1, MATR3, HNRNPK), but with consistently lower effect sizes and stronger expression-driven components (Figure 80), suggesting not only a partial coupling to transcript abundance, but also that they may perform their regulatory role *via* different mechanisms. Some of these mechanisms may involve chromatin modifiers as, in each sample of the Neuron and THP-1 series, we detected an over-representation of various known molecular quantitative trait loci (molQTLs; Y. I. Li *et al*., 2016; Young *et al*., 2022) on TSS clusters targeted by RNAs, particularly those associated with DNA methylation and nucleosome depletion (Figure 81).

**Figure 81:**
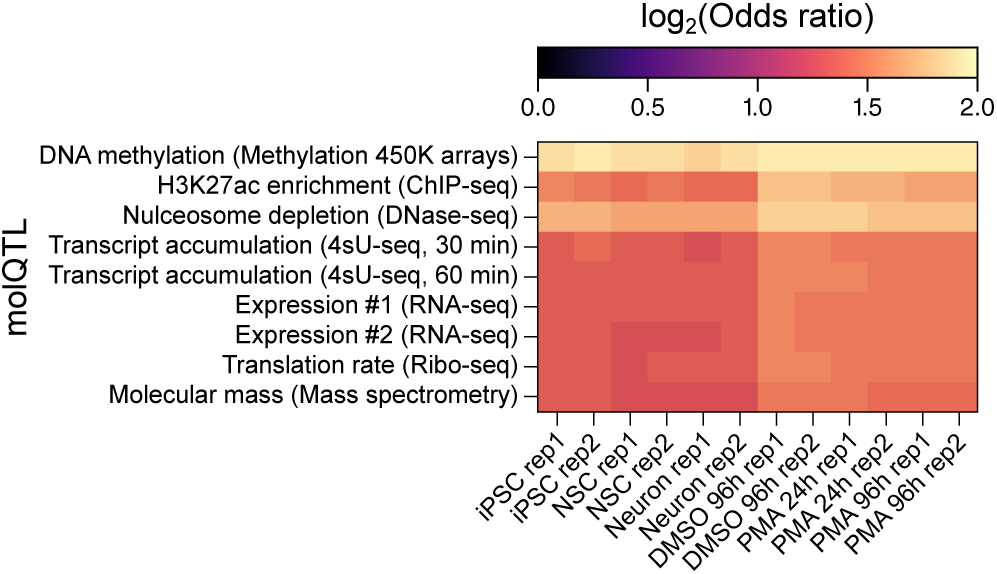
Enrichment (odds ratio) of genomic overlap between TSS clusters targeted by at least one RNA and a range of known molQTLs (Y. I. Li et al., 2016; Young et al., 2022), in each RADICL-seq replicate of the Neuron and THP-1 series. Odds ratios were calculated relative to a permuted genome-wide expectation. All overlaps are significantly enriched (log_2_-transformed odds ratio > 0 and Fisher’s exact test p value < 0.05)

It thus seems possible that many regulatory RNAs interact with chromatin modifiers and transcription factors to modulate the expression of their target TSSs, as previously reported for some specific transcripts such as *ZEB1-AS1* (W. Su *et al*., 2017) or *KCNQ1 OPPOSITE STRAND/ANTISENSE TRANSCRIPT 1* (*KCNQ1OT1*; Mohammad *et al*., 2012).

To further investigate this hypothesis, we selected a subset of iPSC-, Neuron-, DMSO 96h- and PMA 96h-highBW RNAs that show the greatest changes in betweenness but are not differentially expressed between iPSC and Neuron or between DMSO 96h and PMA 96h, respectively (Methods). We then estimated their binding propensity to a set of 1,163 known or predicted nuclear RBPs using the catRAPID algorithm (Armaos *et al*., 2021). To define a prediction score baseline, we also performed this analysis with an array of RNAs that were not detected by RADICL-seq, (Methods). 31% to 43% of each set of highBW RNAs were significantly predicted to interact with at least one tested RBP, while ∼32% of all RBPs were significantly predicted to interact with at least one RNA within each set, including all of the very few differentially expressed proteins included in our tested pool (Figures 82 and 83). On average, most RBPs are predicted to interact with 25 to 50 different RNAs per highBW set, while each RNA is predicted to interact with a variable number of different RBPs, with a peak at around 300 individual proteins. After pooling all interaction prediction results from each set of highBW RNAs, we grouped these RBPs in 3 different clusters A, B and C and, conversely, grouped our tested RNAs in 4 different clusters 1, 2, 3 and 4, as follows: RNAs from cluster 1 are predicted to interact with nearly all RBPs in clusters A, B and C; RNAs from cluster 2 are predicted to interact with nearly all RBPs in clusters A and B, but not C; RNAs from cluster 3 are predicted to interact with nearly all RBPs in clusters A, but not B nor C; and finally, RNAs from cluster 4 are predicted to interact with only a small number of RBPs, independently of their cluster (Figure 84). RNAs from each set of highBW RNAs from the THP-1 series are distributed relatively equivalently in all 4 RNA-clusters, while half of those from the Neuron series belong to RNA-cluster 1 and the rest are roughly equally spread between the three other RNA-clusters (Figure 85). We notably observed a positive correlation between the length of the RNA and the number of RBPs significantly predicted to interact with it, as the coding loci of RNAs belonging to cluster 1 tend to be longer than those of RNAs belonging to cluster 2, which are themselves longer than those of RNAs belonging to cluster 3 (Figure 86).

**Figure 82:**
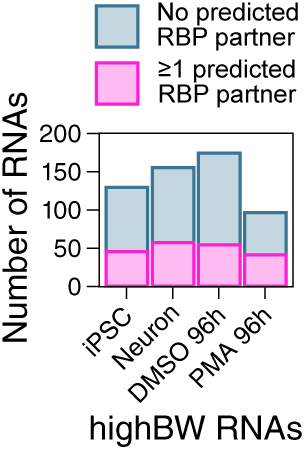
Number of tested iPSC-, Neuron-, DMSO 96h- and PMA 96h-highBW RNAs that are predicted to interact (catRAPID, Armaos et al., 2021) with at least one tested nuclear RBP.

**Figure 83:**
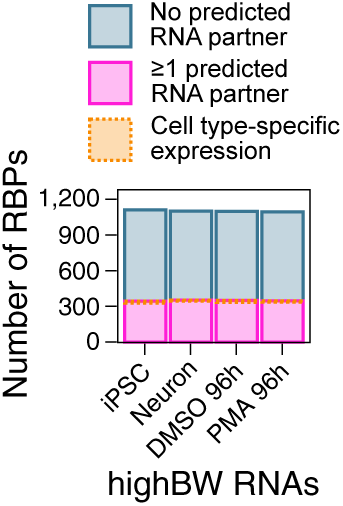
Number of tested nuclear RBPs predicted to interact (catRAPID, Armaos et al., 2021) with at least one tested iPSC-, Neuron-, DMSO 96h- or PMA 96h-highBW RNA. The proportion of RBPs the coding gene of which is differentially expressed (logFC > 2 and FDR < 0.01, edgeR, Robinson et al., 2009; applied to CFC-seq data) between iPSC and Neuron or between DMSO 96h and PMA 96h is highlighted in orange.

**Figure 84:**
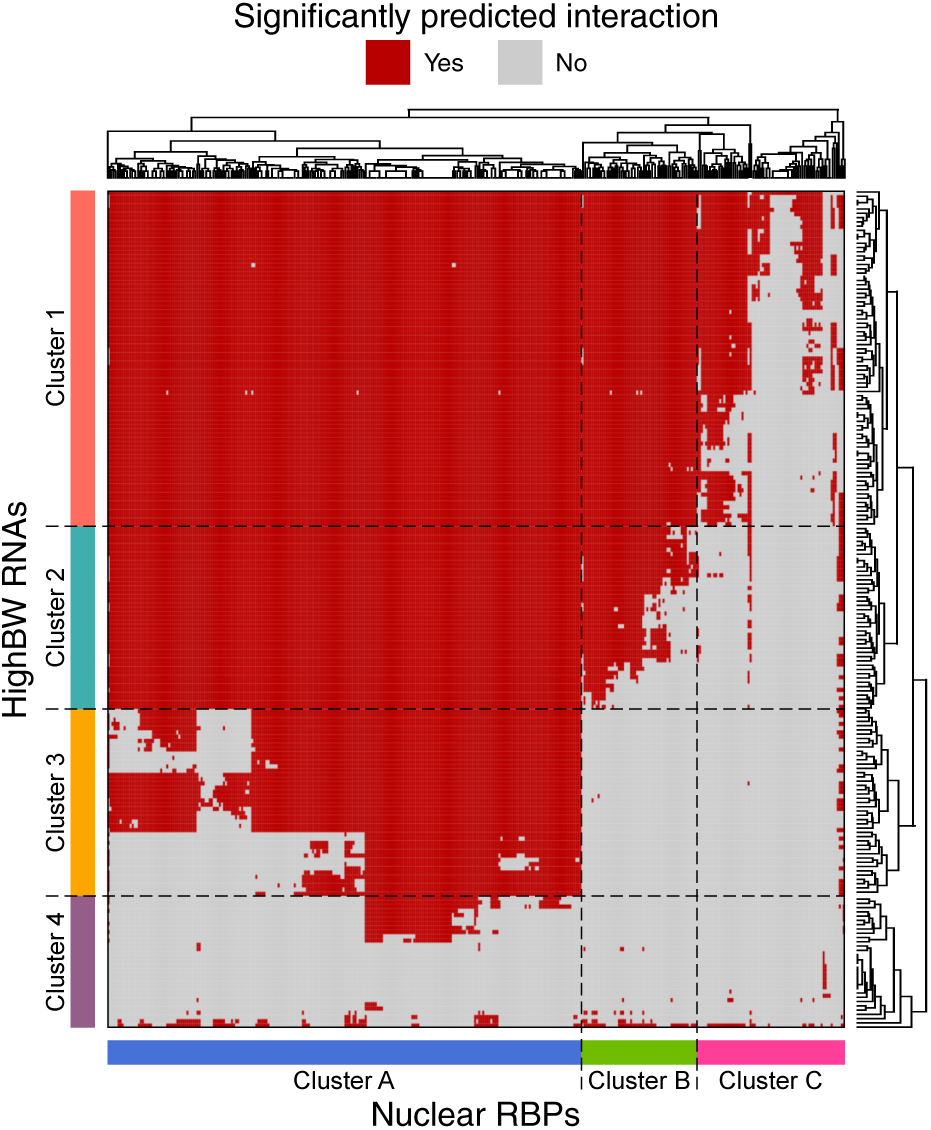
Heatmap depicting pairs of highBW RNAs and nuclear RBPs significantly predicted to interact with each other (red; catRAPID, Armaos et al., 2021). All transcripts from each set of tested highBW RNAs were pooled together. Tested RNAs and RBPs with no significantly predicted interactor were omitted from this figure. RNAs and RBPs are ordered and grouped based on hierarchical clustering by the weighted pair group method with arithmetic means (WPGMA).

**Figure 85:**
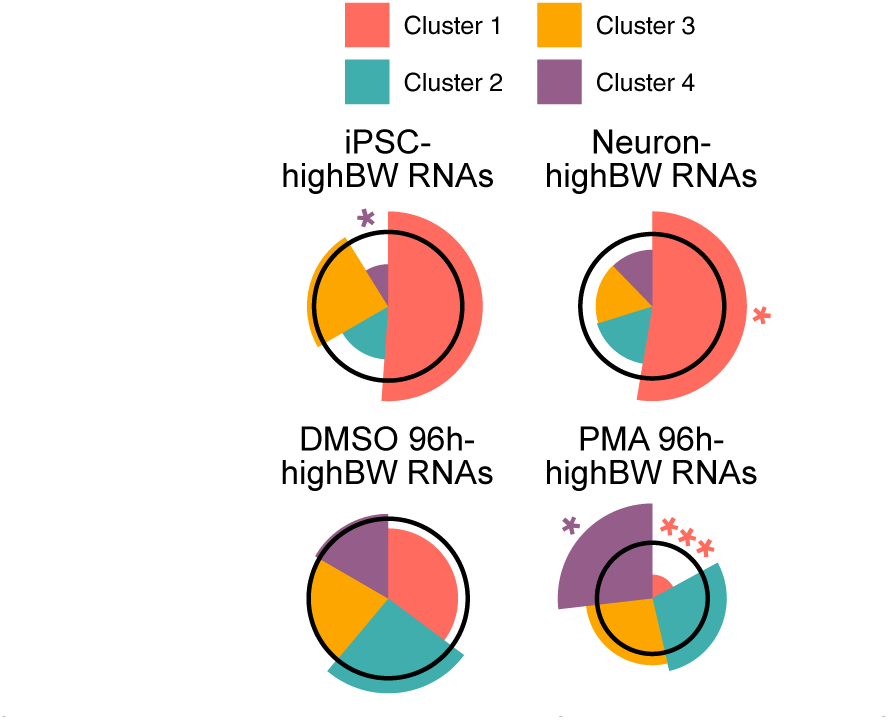
Distribution of RNAs belonging to each of the clusters defined in Figure 84 among all RNAs predicted to interact with at least one RBP, in each set of tested highBW transcripts. The radius of each wedge represents the enrichment (representation factor) of a given RNA-cluster among a specific set of highBW RNAs, relative to its representation among all tested RNAs pooled together. The black circles indicate a representation factor of 1, so that wedges encompassed within them depict an under-represented category and wedges that extend beyond them depict an overrepresented category. Significance was calculated by hypergeometric test; *p value < 0.05; ***p value < 0.001.

**Figure 86:**
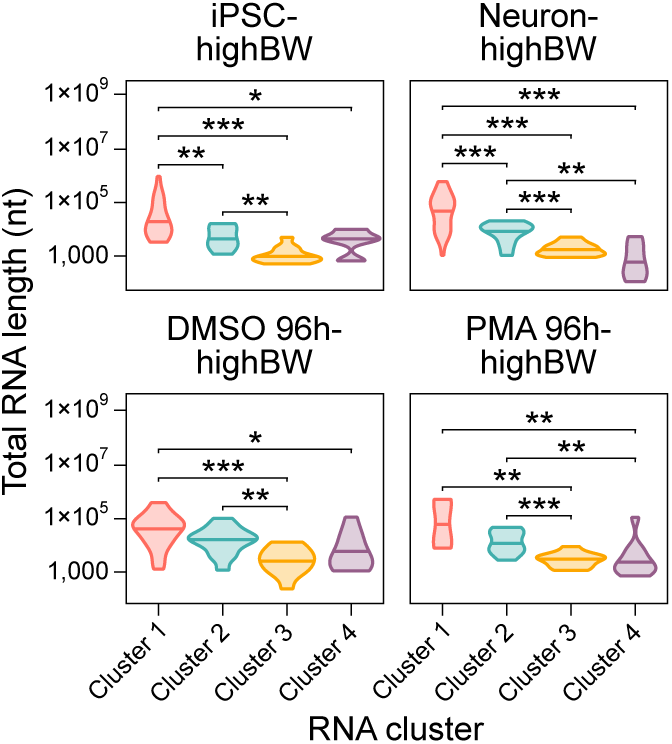
Distribution of coding locus length (including introns) for all highBW RNAs belonging to each cluster defined in Figure 84. Stars indicate significant differences between different clusters (Student’s t-test; *p value < 0.05; **p value < 0.01; ***p value < 0.001).

RNA-protein interaction predictions computed by catRAPID are generated per RNA fragment, after parsing each input transcript into 51-nt chunks. In our analysis, an average of approximately 0.5% of all 51-nt fragments from a given RNA were significantly predicted to interact with a given RBP (Figure 87). When pooled together, most fragments were associated with 1 to 30 different proteins, even for cluster 1 RNAs (Figure 88), highlighting that not all RBPs predicted to interact with the same RNA are expected to bind this transcript at the same position. To further understand the parameters that define the set of RBPs that a given RNA is likely to interact with, we performed a motif enrichment analysis on interacting fragments (*i.e.*, 51-nt fragments predicted to interact with at least one RBP) from cluster 3 RNA *versus* their non-interacting fragments (Figure 89). We found several motifs that are shared by up to 50% of these RNAs, suggesting that RNA-RBP pairing is strongly sequence-dependent. Further supporting this hypothesis, performing the same analysis on interacting fragments from cluster 2 RNAs *versus* interacting fragments from cluster 3 RNAs yields 6 different motifs that are highly shared between cluster 2 RNAs, which may explain why these transcripts are predicted to interact with cluster B RBPs while cluster 3 RNAs are not. Similar results were obtained when comparing interacting fragments from cluster 1 RNAs *versus* interacting fragments from cluster 2 RNAs. Importantly, most of these motifs are constituted of relatively repetitive sequences that are rich in adenines and uraciles. Consistently, we found that interacting fragments from nearly all RNA clusters are significantly enriched in repeat elements, specifically microsatellites and low-complexity regions (Figure 90). Combined with our results in Figures 26 and 27, it thus seems possible that RNA-DNA interactions involve the concomitant binding of RNAs to proteins *via* their repeated sequences, which is known to occur during RNA granule formation (Jain & Vale, 2017; Krzyzosiak *et al*., 2011).

**Figure 87:**
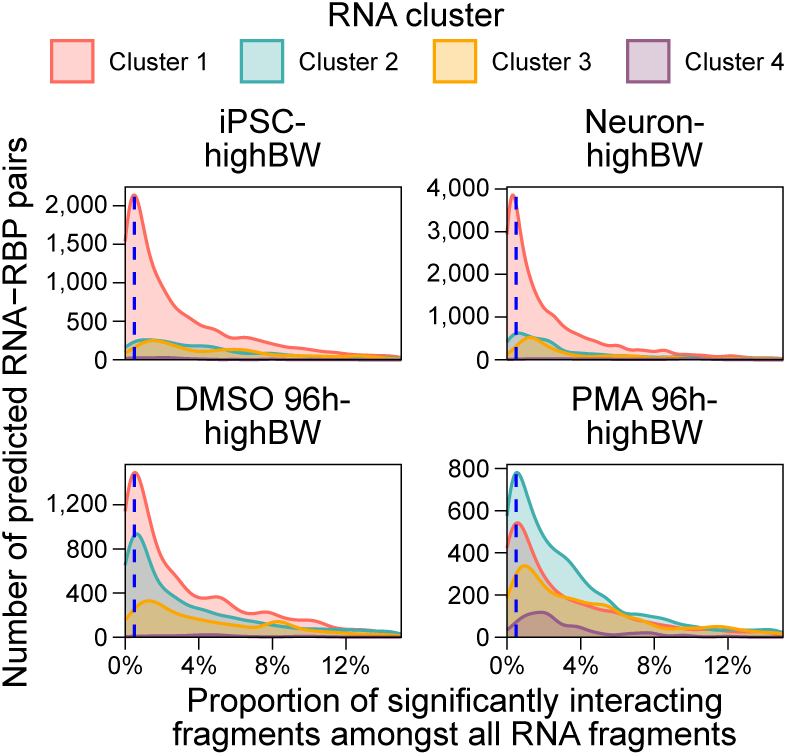
Distribution of RNA-RBP pairs in function of the percentage of all of the RNA’s 51-nt fragments (parsed by catRAPID, Armaos et al., 2021) that are significantly predicted (Z-score ≥ 1.5) to interact with that RBP. RNA-RBP pairs are separated in function of the sample where the RNA has the highest betweenness and in function of the cluster (defined in Figure 84) that this RNA belongs to. Dashed blue lines indicate a percentage of significant fragments equal to 0.5%. For better visibility, only RNA-RBP pairs with a percentage of significant fragments < 15% are shown, as pairs ranging beyond that threshold are extremely few.

**Figure 88:**
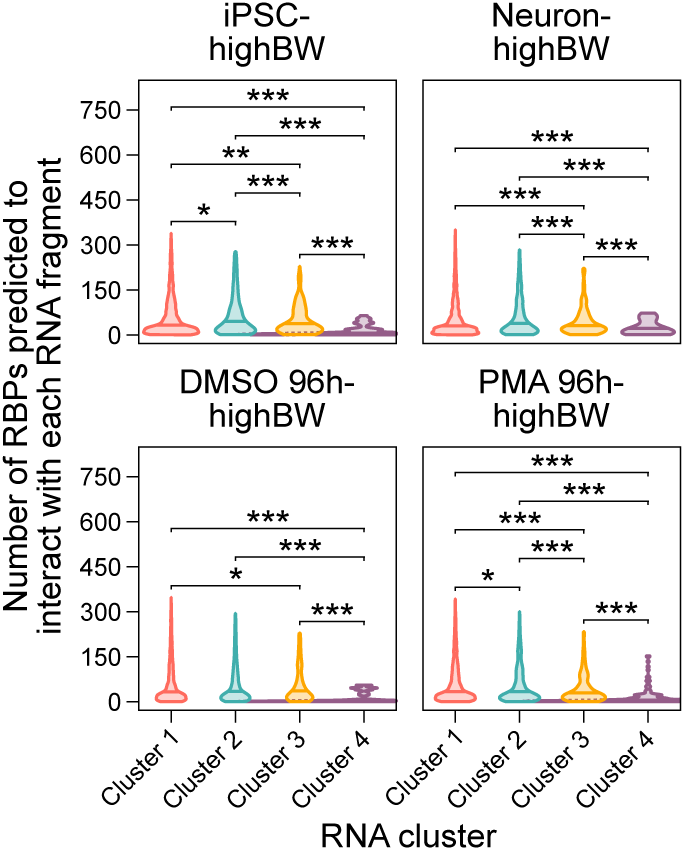
Distribution of the number of RBPs significantly predicted to interact with each 51-nt RNA fragment, in function of the sample where the RNA has the highest betweenness and in function of the cluster (defined in Figure 84) that this RNA belongs to. Only fragments predicted to interact with at least 1 RBP were considered. Stars indicate significant differences between different clusters (Student’s t-test; *p value < 0.05; **p value < 0.01; ***p value < 0.001). For better visibility, the violins corresponding to Cluster 4 are cropped on the right side.

**Figure 89:**
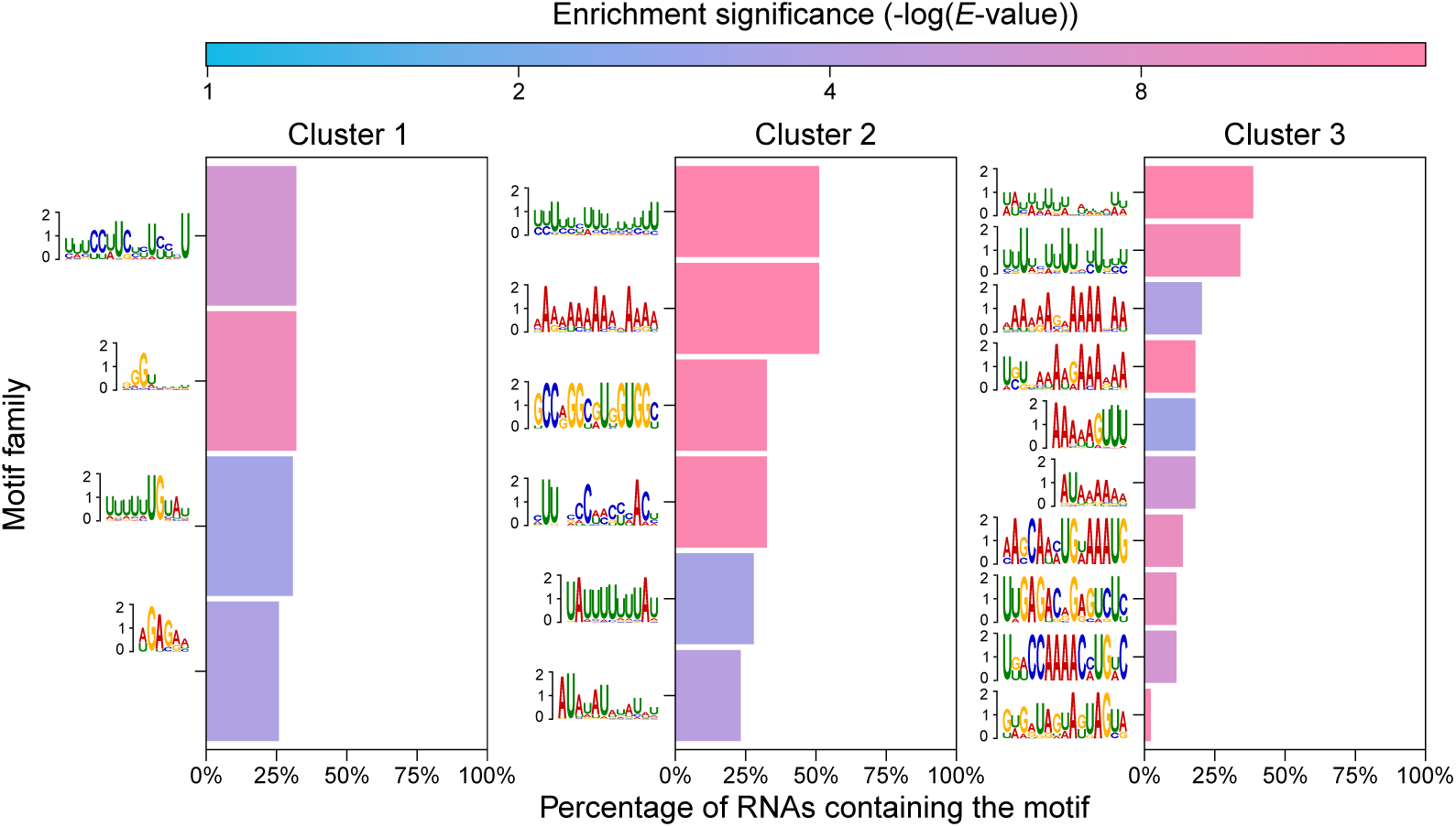
Proportion of RNAs in each of the clusters defined in Figure 84 that contain sequence motifs detected by a motif enrichment analysis (XSTREME; Grant & Bailey, 2021) performed on 51-nt fragments predicted to interact with at least one RBP. Fragments from cluster 3 RNAs for which no RBP partner was significantly predicted were used as control background for the significant RNA fragments of cluster 3. The sequences of these latter fragments were in turn used as control background for the significant fragments of cluster 2, which were themselves used as control background for the significant fragments of cluster 1. Only motif families with E-value < 0.05 are shown.

**Figure 90:**
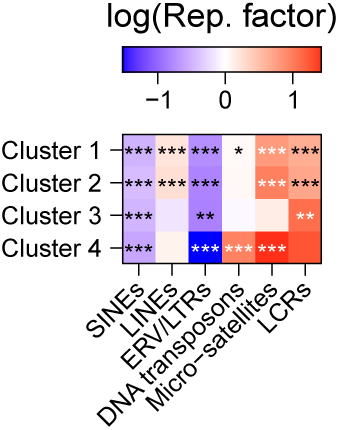
Enrichment (log-transformed representation factor) of various repeat elements on 51-nt RNA fragments predicted to interact with at least one RBP, compared to those that are not predicted to interact with any RBP. Fragments are separated in function of the cluster that their associated RNA belongs to, as defined in Figure 84. A null enrichment value corresponds to no enrichment nor depletion compared to what would be expected by chance. Stars indicate the significance of the enrichment (hypergeometric test; *p value < 0.05; **p value < 0.01; ***p value < 0.001).

In line with this conjecture, we detected a very significant enrichment for motifs containing repeats of glutamic acid (E) and/or aspartic acid (D) among the predicted interacting fragments of RBPs belonging to cluster A, *i.e.* RBPs that interact with the highest number of RNAs (Figure 91). In yeast, proteins containing D/E repeats were reported to regulate the binding of other proteins to the DNA by acting as competitors, and to participate in mRNA transcription and processing (Chou & Wang, 2015); in humans, these proteins appear to be involved in chromatin regulation, especially those containing longer repeats (Shukla *et al*., 2022). Consistently, while we found that the GO annotations of RBPs from all clusters are enriched in terms related to RNA transcription and processing, we also observed that those of RBPs in cluster A are specifically enriched in terms related to DNA maintenance and modification (Figure 92). Additionally, more than one fifth of all RBPs in each cluster are annotated with GO terms indicating that they interact with the chromatin (Figure 93).

**Figure 91:**
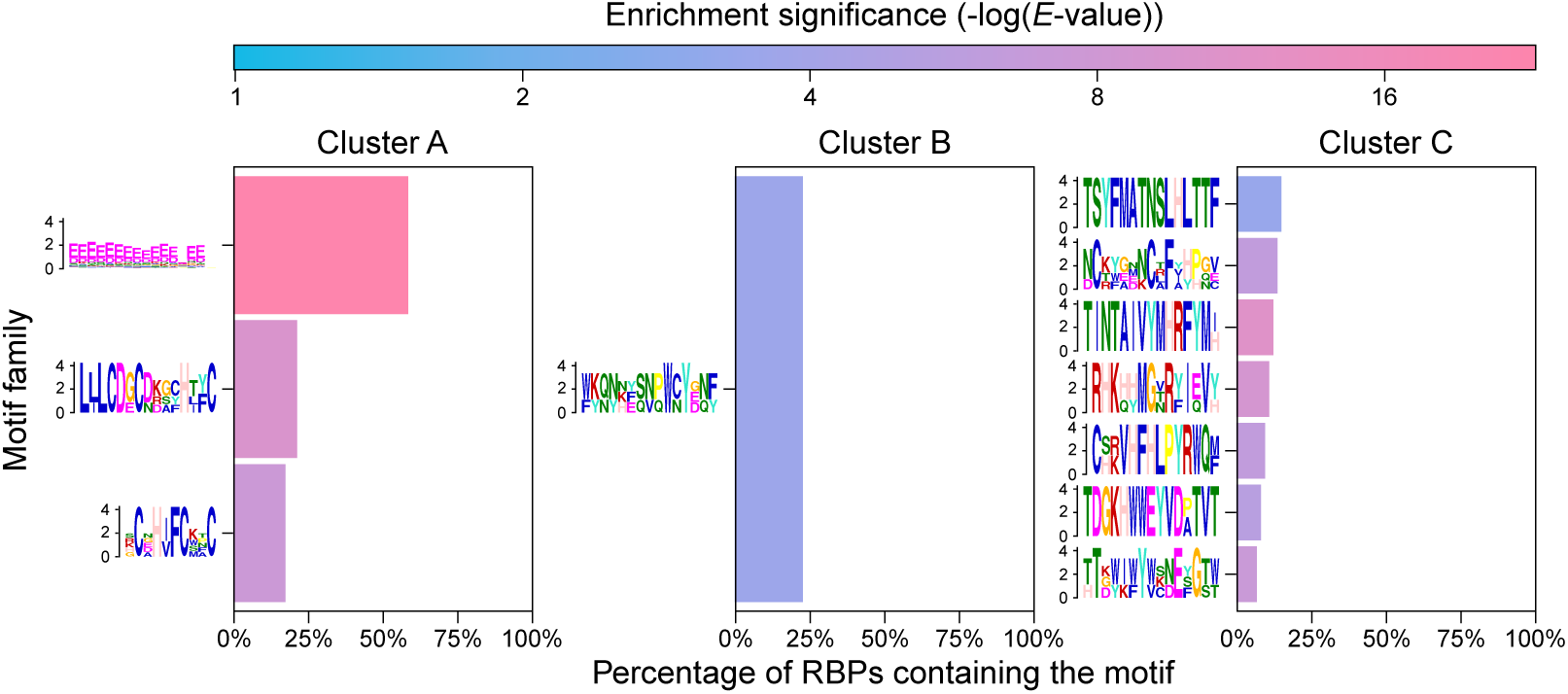
Proportion of RBPs in each of the clusters defined in Figure 84 that contain sequence motifs detected by motif enrichment analysis (XSTREME; Grant & Bailey, 2021). RBPs for which no RNA partner was significantly predicted were used as control background for RBPs belonging to cluster C. The sequences of these latter RBPs were in turn used as control background for the RBPs belonging to cluster B, which were themselves used as control background for the RBPs belonging to cluster A. Only motif families with E-value < 0.05 are shown.

**Figure 92:**
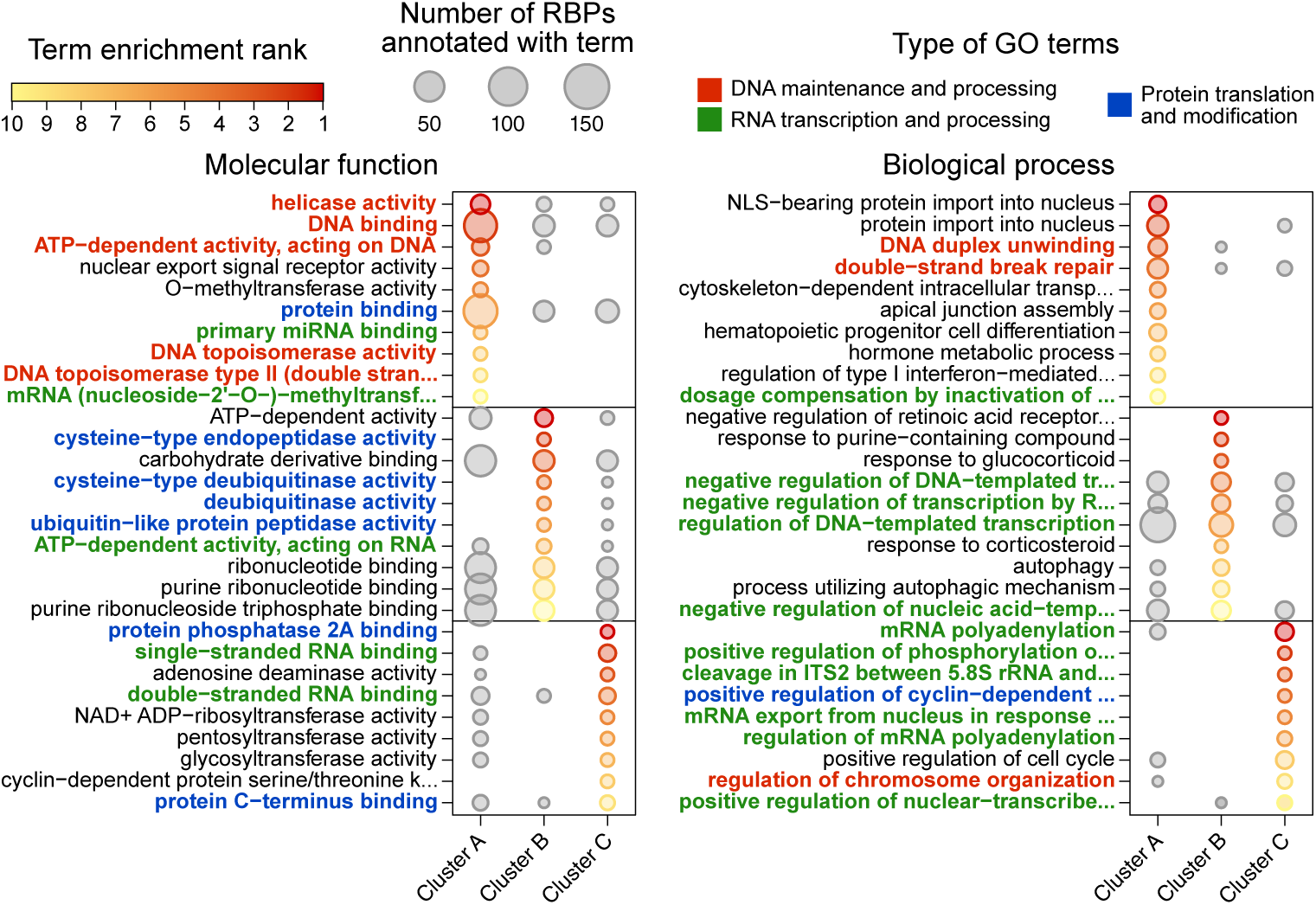
Top 10 most enriched (Fisher’s exact test, topGO, Alexa & Rahnenführer, 2006) gene ontology (GO) terms among the annotation of RBPs from each cluster defined in Figure 84, compared to the annotation of all nuclear RBPs tested in our catRAPID (Armaos et al., 2021) analysis. The color gradient represents the significance rank of the enrichment of a given GO term and the size of the dots represent the number of RBPs annotated with each GO term within each cluster. GO terms were manually categorized and are colored accordingly.

**Figure 93:**
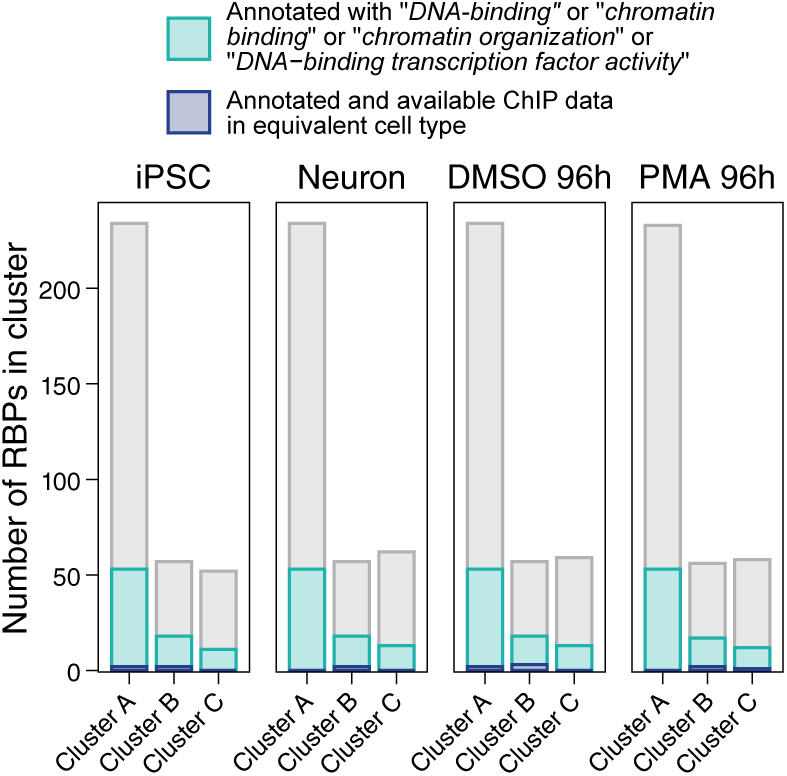
Number of proteins annotated with GO terms “DNA-binding”, “chromatin-binding”, “chromatin organization” and/or “DNA-binding transcription factor activity” among each RBP cluster defined in Figure 84. The number of these annotated proteins for which public ChIP-seq data in cell types equivalent to our iPSC, Neuron, DMSO 96h or PMA 96h samples were available at the time of this study are shown in dark blue.

To assess whether these chromatin-binding RBPs contact the chromatin at the same positions as the DNA-interacting RNAs that they are predicted to associate with, we collected 199 publicly available protein ChIP-seq datasets generated in cell types equivalent to our iPSC, Neuron, DMSO 96h and PMA 96h samples. Among these 199 datasets, 4, 2, 5 and 3 correspond to a ChIP of an RBP predicted to have at least one RNA partner in iPSC, Neuron, DMSO 96h and PMA 96h, respectively (Figures 93 and 94). When comparing the DNA-binding sites of these RBPs with those of their predicted RNA partners, we found that between ∼5% and ∼25% of our tested RBP-RNA pairs show a significantly stronger colocalization than any other protein (Figure 94). The 199 ChIP-seq datasets that we retrieved conveniently featured DNA-binding information for 3 RBPs, namely BROMODOMAIN-CONTAINING PROTEIN 2 (BRD2), INTEGRATOR COMPLEX SUBUNIT 10 (INT10) and ENHANCER OF ZESTE HOMOLOG 2 (EZH2), in more than one of our cell types of interest, which allowed us to evaluate the cell type-specificity of these colocalizations. Of the 25 RBP-RNA pairs that we could thus investigate (Figure 96), we identified 6 with a particularly striking whole-genome and cell type-specific colocalization, *i.e.* cases in which the RBP binds the DNA at the target sites of its predicted partner RNA in the cell type where this RNA has the highest betweenness, but not in the other(s) (Figure 95). An example of such cell type-specific colocalization is between BRD2 and *ADDITIONAL SEX COMBS-LIKE PROTEIN 2* (*ASXL2*), a DMSO 96h-highBW RNA, in the promoter region of FANTOM5 CAT gene *CATG00000042177* (Figure 97). BRD2 is an activating transcription factor that plays a role in the inflammatory response (Cheung *et al*., 2017; LeRoy *et al*., 2008) and is also known to associate with CTCF as well as to induce the formation of TADs (Hsu *et al*., 2017; Xie *et al*., 2022). While we detected a CTCF peak in the promoter of *CATG00000042177*, this peak exists in both DMSO 96h and PMA 96h, whereas BRD2 only binds this region in DMSO 96h when *ASLX2* transcripts are also present. Consistently, we observe higher *CATG00000042177* transcription in DMSO 96h, hinting that *ASXL2* and BRD2 may function together to activate its expression. Another example is that of TSS clusters overlapping the first exons of *ATPASE PHOSPHOLIPID TRANSPORTING 8B4* (*ATP8B4*) and *SOLUTE CARRIER FAMILY 27 MEMBER 2* (*SLC27A2*), which are more bound by both histone methyltransferase EZH2 and Neuron-highBW *RNA SRC HOMOLOGY 2 DOMAIN CONTAINING ADAPTOR PROTEIN 4* (*SHC4*) in Neuron than in iPSC, and concomitantly show an increase in H3K27me3 marking accompanied by a reduction in transcriptional activity (Figure 97). Despite this analysis being limited to a small subset of our data, these findings hint that a large number of DNA-interacting RNAs with a high regulatory potential may perform this function by associating with proteins at their target TSSs, echoing prior conclusions derived from different approaches in other cell types (Agrawal *et al*., 2024; Bonetti *et al*., 2020; Gavrilov *et al*., 2020; Khlebnikov *et al*., 2025; Shu *et al*., 2024).

**Figure 94:**
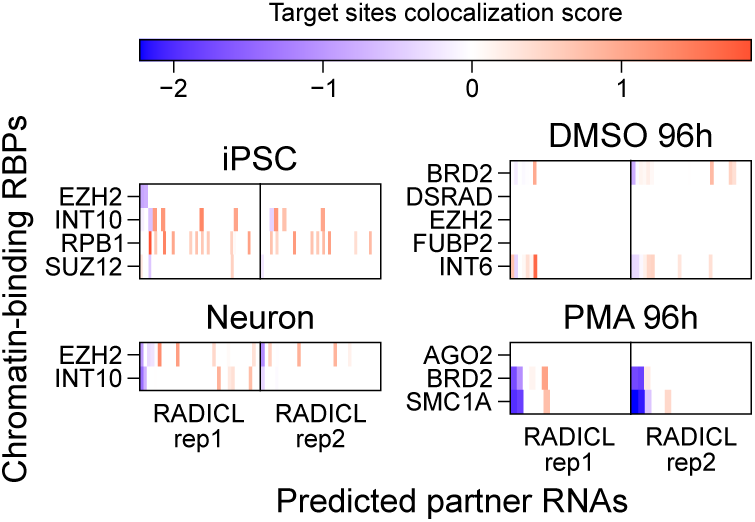
Colocalization score (see “Methods”) between the chromatin target sites of RBPs, as detected by ChIP-seq, and the target sites of each of their predicted partner RNAs, as detected by RADICL-seq. A colocalization score > 0 (red) indicates that the target sites of a given RBP (rows) significantly overlap those of a given RNA (columns). A colocalization score < 0 (blue) indicates that the target sites of a given RBP and a given RNA are significantly mutually exclusive. Non-significant colocalization scores are shown in white.

**Figure 95:**
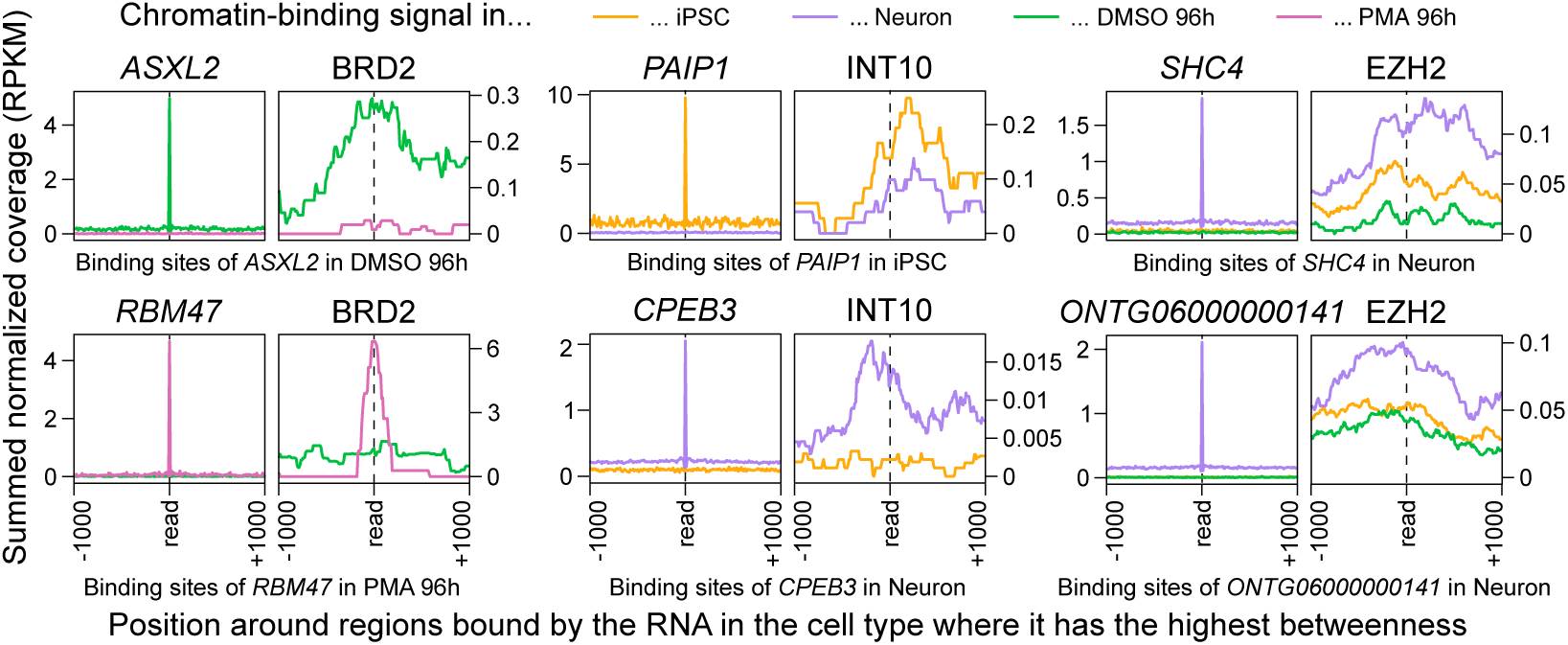
Distribution of RADICL-seq DNA read fragments (left) and BDR2, INT10 or EZH2 ChIP-seq reads (right) at loci targeted by transcripts from ASXL2, RBM47, PAIP1, CPEB3, SHC4 and ONTG06000000141, in the sample where these RNAs have the highest betweenness. Data is shown in all samples for which RBP ChIP-seq data was available, and for all RNA-RBP pairs the target sites of which significantly colocalize, as determined in Figure 94. Only RADICL-seq reads originating from the indicated transcript are pictured in the left panels. For each sample and each type of data, signals for all RNA-targeted loci (extended by ± 1 kb) were summed together, after normalizing coverage values to RPKM.

**Figure 96:**
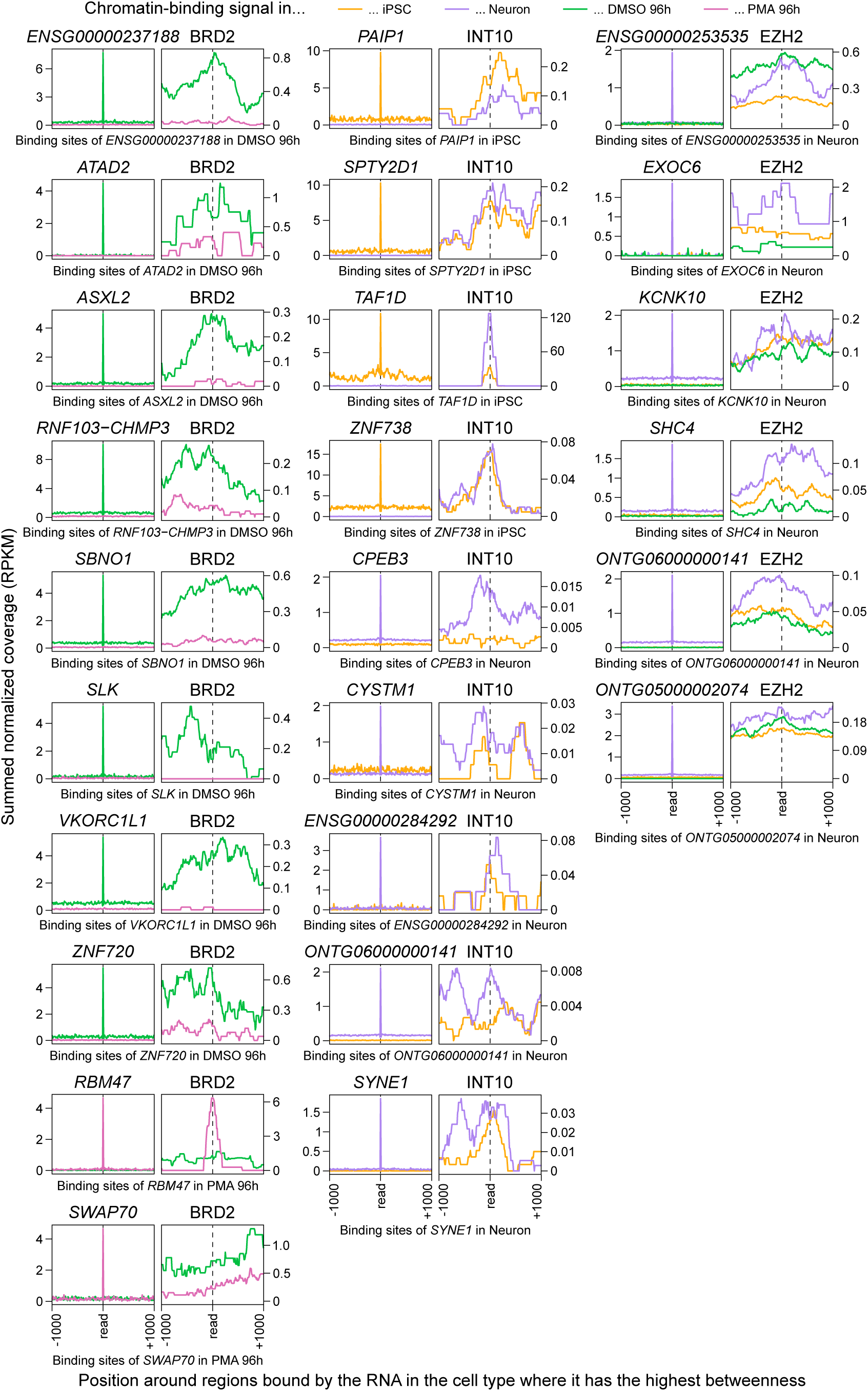
Distribution of RADICL-seq DNA read fragments (left) and BDR2, INT10 or EZH2 ChIP-seq reads (right) at loci targeted by a given RNA in the sample where this RNA has the highest betweenness. Data is shown in all samples for which RBP ChIP-seq data was available, and for all RNA-RBP pairs the target sites of which significantly colocalize, as determined in Figure 94. Only RADICL-seq reads originating from the indicated transcript are pictured in the left panels. For each sample and each type of data, signals for all RNA-targeted loci (extended by ± 1 kb) were summed together, after normalizing coverage values to RPKM.

**Figure 97:**
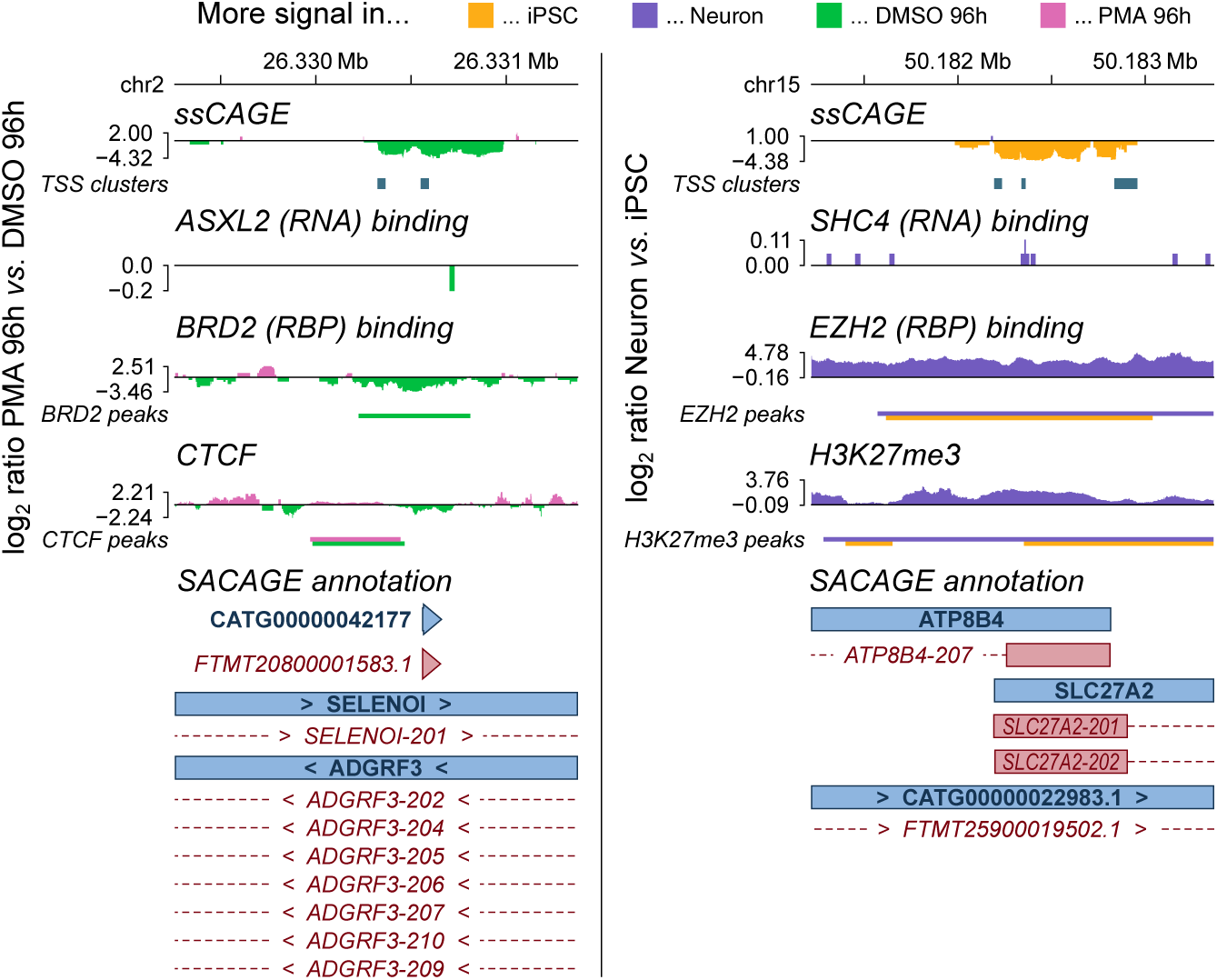
(Left) Genome browser-like view of ssCAGE signal, RADICL-seq DNA read fragments for which the RNA fragments originate from ASXL2, and ChIP-seq signal for BRD2 and CTCF at the promoter region of CATG00000042177, in the DMSO 96h and PMA 96h samples. (Right) Genome browser-like view of ssCAGE signal, RADICL-seq DNA read fragments for which the RNA fragments originate from SHC4, and CUT&Tag/ChIP-seq signal for EZH2 and H3K27me3 around the first exons of ATP8B4 and SLC27A2, in the iPSC and Neuron samples. On both sides, TSS clusters detected by ssCAGE in each series as well as ChIP-seq peaks detected in each sample are indicated as horizontal bars below the corresponding tracks. ssCAGE, RADICL-seq and ChIP-seq signals are shown as the log_2_ ratio of the coverage in PMA 96h vs. DMSO 96h or in Neuron vs. iPSC, and are colored respective to the sample in which they have the highest value. Gene and transcript models located in these regions according to the SACAGE annotation are drawn at the bottom.

## Discussion

By providing RADICL-seq datasets generated across 16 different cell types, the FANTOM6 data collection represents the most extensive survey of the human RNA-DNA interactome to date. Unlike previous reports, which often focused only on long-range contacts (Sridhar *et al*., 2017), on specific RNA classes (Agrawal *et al*., 2024; J. Li *et al*., 2022; Tenorio *et al*., 2023), or on the most abundant or most significant interactions (Bell *et al*., 2018; Calandrelli *et al*., 2023; X. Li *et al*., 2017; Sridhar *et al*., 2017; Zvezdin *et al*., 2025), the present study was applied to all significant interactions detected by RADICL-seq which, combined with the use of our expanded SACAGE annotation, permitted us to create an uniquely vast catalog of RNA-DNA contacts. This scope allowed us to confirm the cell type-specificity of this interactome, which had only been glimpsed by smaller-scale studies (Bonetti *et al*., 2020; X. Li *et al*., 2017; Wen *et al*., 2024; B. Zhou *et al*., 2019). Notably, the presence of multiple time series among our samples revealed a considerable reorganization of RNA-DNA contacts during cell differentiation, demonstrating that the RNA-DNA interactome is extremely and generally dynamic, as hinted by prior reports in mammals (Bell *et al*., 2018; Bonetti *et al*., 2020; Calandrelli *et al*., 2020; Limouse *et al*., 2023). Several results from our analyses, such as the dynamicity of RNA-DNA contacts and the major contribution of both lncRNAs and protein-coding gene-derived transcripts to the pool of chromatin-associated RNAs, were also previously obtained in reptiles (Tenorio *et al*., 2023) and/or in plants (L. Li *et al*., 2021). This suggests that the fundamental features of the RNA-DNA interactome are evolutionarily conserved across kingdoms.

We separated RNA-chromatin interactions into two classes: *cis* interactions, which occur largely within the same TAD as their source locus and likely rely more on passive RNA diffusion, and *trans* interactions, which target sites independently of genomic distance and would rely on specific guides or attractors. Distal contacts appear to be central to the specialization of the interactome, as their proportion and importance for the connectivity of the RNA-DNA interaction network increase during neuronal differentiation and monocyte-to-macrophage activation. While this *cis*/*trans* dichotomy has already been implemented in other publications (Bell *et al*., 2018; Engreitz *et al*., 2016; Limouse *et al*., 2023), our large number of samples reveals that some its previously reported characteristics are highly context-dependent and may not generalize across cell types. For instance, Sridhar *et al*. (2017) and Limouse *et al*. (2023) identified snoRNAs as the major *trans* interactors in human embryonic stem cells (hESCs) and human embryonic kidney cells. While we indeed observed a similarly elevated contribution from snoRNAs among *trans*-interacting transcripts in our iPSC samples, this was not the case in most of our more differentiated cells, suggesting that this feature may be limited to early development and highlighting the need for a broad sample selection to define global properties of the RNA-DNA interactome.

Integrating our RADICL-seq with complementary genome-wide sequencing data further enabled us to explore the parameters that influence RNA-DNA interaction patterns. For instance, the ability of a transcript to perform *trans* interactions does not appear to merely be the result of high expression levels that would allow its passive “spillover” beyond TAD boundaries, but rather relies other properties that are specific to that RNA, such as its length or the nature of its source locus. More generally, we found that expression levels are a relatively poor predictor of the interacting behavior of chromatin-associated transcripts. In particular, the most pivotal RNAs in our interactome networks are not necessarily the most transcribed and, similarly, we found that changes in RNA-DNA contact patterns are not solely dictated by changes in source expression, consistent with prior observations in mammals (Bell *et al*., 2018; Bonetti *et al*., 2020; Calandrelli *et al*., 2020) and in plants (L. Li *et al*., 2021). Moreover, unlike previously reported in Limouse *et al*. (2023) who compared only two different cell types, our large collection of samples revealed that the cell type-specificity of RNA-DNA interactomes resides more in their assortment of source/target combinations rather than in their overall sets of sources and targets. Altogether, these findings indicate that the RNA-DNA interactome cannot be inferred from transcriptional profiles alone, as attempted by other studies (Yun *et al*., 2024 or Hu *et al*., 2025, for instance).

Similar to how expression is not a good indicator of DNA-targeting propensity, we found that neither chromatin accessibility nor the histone modifications assessed by our global analysis can accurately predict RNA-targeted regions. In particular, changes in histone marks do not significantly correlate with changes in RNA-DNA interaction, even at target sites that do carry a recognizable epigenetic signature. Likewise, even if we observed that shifts in A/B subcompartment status appear to modulate the frequency of RNA interactions targeting a given locus, the output of this influence seems to be determined by additional parameters. Additionally, we found that most individual RNAs do not show a marked specificity for chromatin contexts solely defined by 3D compartmentalization and histone modifications. These results indicate that the putative relationships between RNA-mediated contacts and these chromatin properties are likely restricted to specific cases rather than general rules, which cautions against generalizing insights gained from studying individual RNAs to the interactome at large.

It also remains possible that other epigenetic markers that were not investigated here, such as DNA methylation (W. Huang *et al*., 2022), other histone modifications, or specific histone variants (Kohestani & Wereszczynski, 2023; Soboleva *et al*., 2017), play a more decisive part in shaping RNA-DNA interactions. In this regard, our recently developed RADIP-seq technology, which combines RADICL-seq with ChIP-seq to capture RNA-DNA contacts associated with specific chromatin features (Shu *et al*., 2024), will be a useful tool for further exploring the factors that attract RNAs to specific loci. An additional explanation may reside in nuclear bodies such as nuclear speckles and the nucleolus, which are known to be major hubs that attract diverse RNAs (Morf *et al*., 2019; Quinodoz *et al*., 2018) and which could thus shape the dispersed targeting patterns of *trans*-interacting transcripts (Limouse *et al*., 2023), possibly independently of the chromatin context. Post-transcriptional modifications of source RNAs could also be important in determining their patterns of association with the chromatin: for instance, the deposition of methylation marks on promoter-antisense RNAs was shown to increase the number and strength of their contacts, likely by promoting the stability of these transcripts (Hu *et al*., 2025).

The time series present in our sample collection allowed us to explore the regulatory potential of chromatin-associated RNAs at the genome-wide level for both sources and targets. We found that regions displaying attributes characteristic of promoters, enhancers or repressed genes are highly targeted by transcripts, in particular by those that are the most pivotal in the RNA-DNA interaction networks. Remarkably, variations in the frequency of interactions mediated by these key RNAs correlate with strong changes in activity at their targeted regulatory elements, implying that fluctuations in RNA-DNA contacts can modulate transcription. The expression of many chromatin-associated RNAs can notably be controlled by other upstream RNA-DNA interactions, unveiling a complex cascade of RNA-based transcriptional regulation.

Regulatory contacts are not necessarily proximal and may have either activating or repressive effects, depending on the source RNA. In line with prior studies (J. Li *et al*., 2022; X. Li *et al*., 2017), we also observed that regulatory RNAs can target dozens of TSS clusters, while individual promoters are contacted by numerous distinct RNAs. This multiplicity aligns with the “democratic model” proposed by Limouse *et al*. (2023), in which the combined influence of many weakly effective chromatin-associated RNAs determines the transcriptional activity of their shared target. Such redundancy may explain why the knock-down of individual chromatin-associated lncRNAs performed previously in our pilot studies (Agrawal *et al*., 2024; Yip *et al*., 2022) affected the expression of only a minority of their putative targets.

Many examples of RNAs controlling transcription by binding chromatin-remodeling complexes, and either guiding them to their targets or maintaining them in a poised state, have been described in the past (Cajigas *et al*., 2015; Davidovich *et al*., 2013; Kaneko *et al*., 2014; Rahnamoun *et al*., 2018; Ullah *et al*., 2022; Z. Zhao *et al*., 2018; among others). Consistently, we detected a high enrichment for RBP-associated motifs across chromatin-associated RNAs, including RBPs known to bind the chromatin themselves. Furthermore, we found an over-representation of adenines and uraciles on the fragments of putative regulatory RNAs predicted to interact with RBPs, similar to other studies investigating RNAs participating in condensate formation (Jain & Vale, 2017; Klobučar *et al*., 2025; Krzyzosiak *et al*., 2011) or binding H3K27me3-marked loci (Shu *et al*., 2024). A large number of RBPs could therefore recognize and interact with chromatin-associated RNAs *via* their repeated sequences, which may allow individual proteins to associate with multiple RNAs.

However, we also observed a drop of RADICL-seq RNA signal over most repeat elements, which possibly reflects our filtering for uniquely mapped RADICL-seq reads. One way to palliate this issue is to rescue the filtered-out multimappers, as described in Panariello *et al*. (in preparation), who thereby observed from our data that SINEs and LINEs are the most enriched among repeats involved in RNA-DNA interactions, particularly within sources of interchromosomal interactions for the latter. In parallel, the enrichment in RNA signal that we detected on microsatellites and LCRs might partially reflect an under-representation of these elements in the hg38 reference genome used in this study, as many of them have only recently been resolved in the Telomere-to-Telomere (T2T) assembly (Nurk *et al*., 2022). While the regulatory role of DNA-binding RNAs most likely involves their interaction with proteins, the mechanisms by which this interaction would affect transcription remain elusive and are presumably case-specific. For instance, our results suggest that the association between *SHC4* transcripts and EZH2 at the *ATP8B4*/*SLC27A2* promoters in neurons triggers H3K27me3 deposition and gene repression. However, prior reports indicate that the binding of EZH2 to RNAs prevents its catalytic activity (Beltran *et al*., 2016; Gail *et al*., 2024; X. Wang *et al*., 2017), suggesting that *SHC4* may actually dampen and thereby finetune EZH2 activity, or transiently guide it to its targets, or even be recruited after H3K27me3 deposition, implying no direct regulatory role. A similar ambiguity arises when considering the monocyte-specific colocalization of *ASXL2* RNAs and BRD2 at the *CATG00000042177* promoter, where *ASXL2* may either recruit or instead be recruited by BRD2.

RNA-DNA interactions may also directly influence transcription through R-loop formation (Niehrs & Luke, 2020). As reported in mice (Bonetti *et al*., 2020), we found that a large fraction of RADICL-seq DNA reads map to R-loop regions; however, our analyses indicate that the RNAs involved in these contacts are unlikely to be actual transcripts forming the RNA:DNA hybrids. Chromatin-associated RNAs may still target R-loops, notably *via* proteins: for example, lncRNA *TAURINE-UPREGULATED GENE 1* (*TUG1*) is known to bind REPLICATION PROTEIN A (RPA) heterotrimers, which stabilize R-loops (M. M. Suzuki *et al*., 2023). Furthermore, iMARGI data generated without RNase H (W. Xu *et al*., 2021) suggest that ERV and LINE-1 repeats facilitate R-loop formation; the removal of R-loops during the RADICL-seq procedure may thus explain the depletion in LINEs and ERV/LTRs among our RNA reads. Developing variants of the RADICL-seq protocol that preserve R-loops will therefore be decisive to further understand how these structures contribute to the regulatory role of RNA-DNA interactions.

Moreover, it should be noted that direct regulatory RNA-DNA interactions in general, whether they involve proteins, R-loops, or any other immediate mechanism, represent only a small fraction of all RNA-DNA contacts. Indeed, most RNA targets do not overlap regulatory elements, and most TSS clusters contacted by RNAs do not show changes in activity during cell specialization. Thus, while RNA-DNA interactions can directly influence transcription at select loci, this role represents a specific rather than a predominant function of the interactome.

Instead, RNA-DNA interactions may more often affect the transcriptional landscape indirectly, by modulating the 3D organization of the genome. Notably, by combining our data in the Neuron series with Promoter Capture Hi-C data, Sahlén *et al*. (2025) found that a significant fraction of enhancer-promoter contacts colocalize with RNA-targeted regions. Like in the present study, this report shows that promoters differentially bound by RNAs are also differentially active during cell specialization, suggesting that RNA-DNA contacts structurally contribute to enhancer-mediated regulation.

From a wider perspective, the RNA-DNA interactome may be involved in similar structural roles at all scales of the chromatin. At the chromosomal scale, we detected a high number of RNA-DNA interactions involving telomeres, especially in cancer cells known to rely on telomeric restoration for replicative immortality (Jafri *et al*., 2016). Considering past evidence that chromatin-associated RNAs like *TERRA* contribute to telomere stability (Cusanelli & Chartrand, 2015) as well as a recent report suggesting that DNA-targeting RNAs regulate centromeric chromatin (Fryer *et al*., 2025), it is plausible that RNA-DNA interactions participate in maintaining the integrity of large chromosomal regions. RNA-DNA contacts may also define the positioning of these regions within the nucleus: integrating our data with GPSeq allowed Kang *et al*. (2025) to identify *trans*-contacting intronic RNAs (TIRs) that form nuclear “clouds” on the chromatin at a subset of DNA loci spread across the genome.

Within chromosomes, we found that TAD borders and loop anchors tend to attract DNA-interacting RNAs, supporting the hypothesis that RNA-DNA interactions are associated with these chromatin structures (Bell *et al*., 2018; Bonetti *et al*., 2020; Calandrelli *et al*., 2023; A. S. Hansen *et al*., 2019; Kuang & Pollard, 2024; Lai *et al*., 2013; W. Li *et al*., 2013; Lucero *et al*., 2025; Saldaña-Meyer *et al*., 2019; Shu *et al*., 2024; Zvezdin *et al*., 2025). In particular, the enrichment in RNA-targeted sites at the edges of TADs may reflect an accumulation of RNAs that are blocked by TAD borders acting as barriers for diffusion, as previously proposed (Bonetti *et al*., 2020; Calandrelli *et al*., 2023). However, our analyses in the dynamic context of cellular specialization showed little correspondence between changes in RNA-DNA interactions and modifications of chromatin loops or TADs, casting doubts on these hypotheses. As the formation of TAD borders has been suggested to involve R-loops (Luo *et al*., 2022), it is also possible that we did not detect the relevant interactions mediating these changes by RADICL-seq.

At the sub-megabase scale, RNAs may act as chromatin scaffolds, independently of their target sites. Digesting all nuclear RNAs leads to the collapse of the chromatin (Hall *et al*., 2014), while its insoluble fraction was reported to contain a pool of transcripts the composition of which is very similar to those detected by RADICL-seq (Creamer *et al*., 2021). These putative scaffold RNAs could perform their structural role locally *via* passive diffusion, either by recruiting architectural proteins to the chromatin (Creamer *et al*., 2021) or by competitively weakening the electrostatic bond between histones and DNA. In light of this hypothesis, the reduction of the *cis*/*trans* interactions ratio that we noticed during neuronal and macrophage specialization may explain the condensation of the chromatin that accompanies cell differentiation (Meshorer *et al*., 2006; Ricci *et al*., 2015; Ugarte *et al*., 2015).

Finally, our results indicate that the multiple roles of the RNA-DNA interactome are embedded in processes key to cell identity and function. Our work thus extends the growing body of evidence linking specific RNA-DNA contacts to cellular dysfunction and human disease (Calandrelli *et al*., 2020; Wen *et al*., 2024; W. Xu *et al*., 2021; Yan *et al*., 2019) by providing a genome-wide view across multiple cellular contexts. A past study by Calandrelli *et al*. (2020) remarkably reported that stress-induced alterations in RNA-DNA interactions persist even after withdrawal of a diabetes-like treatment in endothelial cells. This observation is reminiscent of the phenomenon of epigenetic memory (Thiagalingam, 2020) and could explain the chronic nature of diabetes (Calandrelli *et al*., 2020). Similarly, analyses of our MCF7 and LTED samples by Kato *et al*. (in preparation) and Ichikawa *et al*. (in preparation) revealed that breast cancer cells resistant to estrogen depletion acquire multiple intronic RNA-chromatin condensates, including a “cloud” of *ESR1 LOCUS ENHANCING AND ACTIVATING NONCODING RNA*s (*ELEANOR*s) at the *ESTROGEN RECEPTOR 1* (*ESR1*) super-enhancer, which reinforces *ESR1* activation and may contribute to breast cancer resurgence after endocrine therapy. Furthermore, in our Neuron sample, the aforementioned TIRs identified by Kang *et al*. (2025) are enriched at neurodevelopmental risk loci and may therefore be involved in disorders such as the autism spectrum. Together, these last two examples illustrate the power of the FANTOM6 data collection as a resource for understanding RNA-mediated mechanisms linked to human diseases. In particular, current strategies for RNA-based therapies, such as antisense oligonucleotides (ASO), RNA interference (RNAi) or CRISPR-based genome editing, primarily focus on altering the expression of given target RNAs (Singh *et al*., 2026; Zhu *et al*., 2022) and may therefore lead to various adverse effects due to the potentially diverse functions of their targets.

Our data greatly facilitates the development of safer approaches in which only nefarious RNA-chromatin interactions would be targeted, for instance by precisely altering RNA-contacted regions on the DNA *via* CRISPR-editing therapy (Frangoul *et al*., 2021; D. Wang *et al*., 2020) or by leveraging prime editing technologies (Anzalone *et al*., 2019; Murray *et al*., 2024) to selectively disrupt specific interactions within few base pairs.

Aside from the medical field, our datasets can bring new insights into the mechanisms underlying cell differentiation. For instance, the *ASXL2*/BRD2-mediated regulation suggested by our study may play an important role in the monocyte-to-macrophage transition, given the known immune functions of BRD2 (Cheung *et al*., 2017). Likewise, the RNA-DNA interactions derived from repeat elements investigated by Panariello *et al*. (in preparation) appear to contribute to neuronal and immune lineage specification, while our single-cell datasets enabled Yip *et al*. (in preparation) to identify bimodal *cis*-regulatory elements that cooperate with transcription factors during neuronal differentiation. By integrating the FANTOM6 data with fetal single-cell RNA-seq from human prefrontal cortex and variants acquired by modern humans after divergence from Neanderthals, Vitriolo *et al*. (2026) further uncovered transcription factors that have a evolutionarily-specific activity, as well as the RNAs that seem to operate as their co-regulators.

The FANTOM6 collection can also be used as a database to develop machine-learning algorithms aimed at better understanding the regulation of the genome. Cassan *et al*. (2025) notably propose a clustering method to investigate the changes in *cis* regulatory element activity during differentiation. Similarly, Grapotte *et al*. (2025) produced a fully interpretable model that identifies regulatory elements driving short tandem repeat (STR)-initiated transcription and computes corresponding molecular QTLs, which allowed them to uncover a regulatory interplay between STR-derived RNAs and Alu repeats.

Given the rising interest for research on chromatin-associated RNAs, there is a crucial need for datasets that specifically assess RNA-DNA contacts across a wide array of cell types, similar to the resources already available for transcriptomes, epigenomes, and 3D genome organizations (Abugessaisa *et al*., 2020; Dekker *et al*., 2025; Kawaji *et al*., 2017; The ENCODE Project Consortium, 2012, 2020). While this study lays the foundation for this effort, profiling the RNA-DNA interactome across an even broader range of samples will be essential for systematically integrating RNA-DNA interaction data into future inter-omics analyses. Unfortunately, the current protocols for such profiling remain labor-intensive and costly, which hinders this endeavor. Furthermore, the short reads obtained by RADICL-seq limit the identification of the specific transcript isoform that binds a given DNA region. The expansion of our RNA-DNA contact collection would therefore greatly benefit from combining RADICL-seq with long-read technologies as well as optimizing its protocol to lower its financial and technical burdens. Additionally, developing a single-cell version of RADICL-seq, analogous to the multinucleic acid interaction mapping in single cells (MUSIC) described in Wen *et al*. (2024), would not only facilitate the profiling of hundreds of different cell types, but might also reveal new features of the RNA-DNA interactome that remain hidden at the bulk level.

Altogether, analyses of the FANTOM6 data collection highlight RNA-DNA interactions as a critical layer of chromatin and transcriptional regulation. By providing an unprecedentedly comprehensive atlas of these interactions across diverse human samples, the present study provides the resources and establishes the ground principles for future research on RNA-mediated control of the genome, opening the door to numerous advances in genomics and human medicine.

## Methods

### Human sample collection and cell culture

All human samples used in this study were either commercially available, accessible in public collections, or were collected from patients under informed consent and in accordance with the recommendations from the Declaration of Helsinki. The use of all non-exempt human materials for research in this project has been approved by the RIKEN Yokohama Branch Ethics Committee (approval number RIKEN-Y-2024-098).

For the T cell samples, human peripheral blood from anonymous healthy donors was provided by the Department of Transfusion Medicine and Hematology at Fondazione Istituto di Ricovero e Cura a Carattere Scientifico (IRCCS) Cà Granda Ospedale Maggiore Policlinico in Milan, Italy. Their use for research in this project has been approved by the foundation’s ethics committee (approval number 708_2020).

#### Human dermal fibroblasts (HDF sample) and induced pluripotent stem cells (iPSCpreF6 sample)

The human dermal fibroblasts (HDF) and induced pluripotent stem cells referred to as “iPSCpreF6” in this study are the same cell lines as those described in Yip *et al*. (2022).

The HDF sample corresponds to primary cells derived from the neonatal foreskin of a healthy individual (Lonza, catalog no. CC-2509) cultured at 37°C in a 5% CO_2_ incubator in Dulbecco’s Modified Eagle Medium (DMEM, high glucose with L-glutamine; Gibco^®^, catalog no. 11965092) supplemented with 10% fetal bovine serum (FBS; Gibco^®^, catalog no. 26140079).

The iPSCpreF6 cells were generated from human fetal dermal fibroblasts (HDF-f, Cell Applications™, catalog no. 106-05n) using a non-integrating Sendai-based viral vector (SeV) coding for *OCTAMER-BINDING TRANSCRIPTION FACTOR 3/4* (*OCT3/4*), *SRY-BOX TRANSCRIPTION FACTOR 2* (*SOX2*), *KRÜPPEL-LIKE FACTOR 4* (*KLF4*) and *C-MYC* (Fort *et al*., 2014). These cells were cultured in StemFit^®^ medium (Takara^®^, catalog no. AJ100) under feeder-free conditions at 37°C in a 5% CO_2_ incubator. The cells were seeded on dishes coated with iMatrix™-511 (Nippi^®^, catalog no. 892012) and in presence of 10 μM Rho-associated kinase (ROCK) inhibitor (CultureSure^®^ Y-27632, Wako^®^, FUJIFILM^®^, catalog no. 036-24023). 24 hours after seeding, the ROCK inhibitor was removed by refreshing the StemFit^®^ medium, which was from then on replaced every day until collection. Cell viability was monitored by trypan blue exclusion assay. For sample preparation, cells were dissociated by incubation with 3 mL per 10-cm dish of TrypLE™ Select enzyme (Gibco^®^, catalog no. 12563011), then scraped in StemFit^®^ medium.

#### Induced pluripotent stem cells (iPSC sample), neuron stem cells (NSC sample) and cortical neurons (Neuron sample)

For all experiments, the cells referred to as the “iPSC” sample in this study correspond to human i3N iPSCs in the male WTC11 background, which harbor a doxycycline-inducible *NEUROGENIN 2* (*NEUROG2*) transgene integrated at the adeno-associated virus integration site 1 (AAVS1) locus (C. Wang *et al*., 2017). These cells were kindly gifted by Dr. Michael Ward (National Institutes of Health, United States).

All cells were cultured in StemFit^®^ medium (Takara^®^, catalog no. AJ100) under feeder-free conditions at 37°C in a 5% CO_2_ incubator. The cells were seeded dishes coated with iMatrix™-511 (Nippi^®^, catalog no. 892012) in presence of 10 μM ROCK inhibitor (CultureSure^®^ Y-27632, Wako^®^, FUJIFILM^®^, catalog no. 036-24023). 24 hours after seeding, the ROCK inhibitor was removed by refreshing the StemFit^®^ medium, which was from then on replaced every other day. When the cells reached 70–80% confluency, they were dissociated and detached by incubation with Accutase^®^ (Innovative Cell Technologies^®^, catalog no. AT104) for 10 min at 37°C in a 5% CO_2_ incubator. Cell viability was monitored by trypan blue exclusion assay. For long-term storage, the cells were first washed in Dulbecco’s Phosphate Buffered Saline (DPBS, Wako^®^, FUJIFILM^®^, catalog no. 049-29793) and centrifuged at 200 *g* for 5 min, then resuspended in STEM-CELLBANKER^®^ (Takara^®^, catalog no. CB047) freezing solution and slowly frozen in a freezing container (CoolCell^®^, Corning^®^, catalog no. 432000) at −80°C, before transfer at −150°C.

NSCs were generated from the i3N iPSCs described above following the protocol provided by Gibco^®^ (Publication Number MAN0008031), with slight modifications. Briefly, the iPSCs were seeded on iMatrix™-511-coated 6-well plates (Falcon^®^, catalog no. 353046) at a density of 2.5×10^5^ cells per well in StemFit^®^ medium supplemented with 10 μM ROCK inhibitor. On the following day, the ROCK inhibitor was removed by replacing the medium with PSC Neural Induction Medium (NIM; Gibco^®^, catalog no. A1647801). From then on, the NIM was refreshed every other day.

On the seventh day of neural induction, these P0 NSCs were dissociated and detached by incubation with Accutase^®^ followed by scraping. Cells were pelleted at 300 *g* for 4 min. The pellet was resuspended in Neural Expansion Medium (NEM, Gibco^®^, catalog no. A1647801) with 10 μM ROCK inhibitor and seeded on iMatrix™-511-coated 10-cm dishes (TPP^®^, catalog no. 93100) at a density of 1.0×10^6^ cells per dish. On the following day, the NEM was refreshed to remove the ROCK inhibitor. From then on, the NEM was replaced every other day until the NSCs reached 70-80% confluency (P1 NSCs). The expanded P1 NSCs were harvested and resuspended in STEM-CELLBANKER^®^ freezing solution, then slowly frozen in a freezing container at −80°C, before transfer at −150°C.

Cryopreserved P1 NSCs were passaged once after thawing to ensure that the cells were in a homogeneously healthy condition, as follows: a vial of P1 NSCs was quickly thawed at 37°C in a water bath, then the cells were gently transferred into a conical tube containing pre-warmed NEM medium; after centrifugation at 300 *g* for 4 min, the cell pellets were resuspended in NEM medium with 10 μM ROCK inhibitor and seeded on 10-cm dishes at a density of 1.0×10^6^ cells per dish; following overnight incubation, the NEM was refreshed to remove the ROCK inhibitor and, from then on, the NEM was replaced every other day until the NSCs reached 70-80% confluency (P2 NSCs).

The induction of cortical neurons from these NSCs was performed by doxycycline-inducible *NEUROG2* following the protocol developed by Dr. Peter Heutink’s laboratory (Deutsches Zentrum für Neurodegenerative Erkrankungen, Germany; Rosa *et al*., 2020), with slight modifications. P2 NSCs were harvested and replated at a density of 5.0×10^6^ cells per 10-cm dish coated with poly-L-ornithine (PLO; Sigma-Aldrich^®^, catalog no. P3655) and in Differentiation Medium I, which contains half DMEM/Nutrient Mixture F-12 (DMEM/F-12) supplemented with GlutaMAX™ (Gibco^®^, catalog no. 10565018) and half Neurobasal Medium (Gibco^®^, catalog no. 21103049) as the base, mixed with 0.5× N-2 Supplement (Gibco^®^, catalog no. 17502048), 0.5× B-27 Supplement (Gibco^®^, catalog no. 12587010), 2.5 μg×mL^-1^ insulin (Sigma-Aldrich^®^, catalog no. 19278), 0.5× Minimum Essential Medium (MEM) Non-Essential Amino Acids (Gibco^®^, catalog no. 11140-050), 50 μM β-mercaptoethanol (Gibco^®^, catalog no. 21985023), 2 μM DAPT (Cayman Chemical^®^, catalog no. 3197), 5 μg×mL^-1^ laminin (Gibco^®^, catalog no. 23-17015), 2 μg×mL^-1^ doxycycline (Sigma-Aldrich^®^, catalog no. D-9891) to induce the expression of *NEUROG2*, and 10 μM ROCK inhibitor. On the following day, the ROCK inhibitor was removed by replacing the differentiation medium with an inhibitor-free batch.

On the third day of neural induction, the medium was changed to Differentiation Medium II containing half DMEM/F-12-GlutaMAX™ and half Neurobasal Medium as the base, mixed with 0.5× N-2 Supplement, 0.5× B-27 Supplement, 2.5 μg×mL^-1^ insulin, 0.5× MEM Non-Essential Amino Acids, 50 μM β-mercaptoethanol, 10 μM DAPT, 0.5 μg×mL^-1^ laminin, 2 μg×mL^-1^ doxycycline, 10 ng×mL^-1^ brain-derived neurotrophic factor (BDNF, PeproTech^®^, catalog no. 450-02), 10 ng×mL^-1^ glial cell-derived neurotrophic factor (GDNF, PeproTech^®^, catalog no. 450-10), and 10 ng×mL^-1^ neurotrophin-3 (NT-3, PeproTech^®^, catalog no. 450-03). From the sixth day onwards, half of the medium was replaced every 3 days with a batch that did not contain doxycycline. The cells started exhibiting a neuron-like morphology after only a few days, and displayed a mature neuron morphology within 1 week.

#### THP-1 monocytes (DMSO 96h sample) and PMA-activated macrophages (PMA 24h and PMA 96h samples)

The THP-1 monocytes (originally collected from a 1-year-old patient with acute monocytic leukemia; Tsuchiya *et al*., 1980) used in this study are the same as those used by the FANTOM4 project (H. Suzuki *et al*., 2009). Because THP-1 cells exhibit heterogeneous responses to stimuli such as exposure to phorbol 12-myristate 13-acetate (PMA; Aldo *et al*., 2013; Eperon *et al*., 1997), we sub-cloned this cell line by limiting dilution, in order to enhance the signal-to-noise ratio in our analyses. A single clone, 1-E9, was selected based on its ability to differentiate in a relatively homogeneous manner in response to PMA, as demonstrated by alterations in cell morphology. 1-E9-derived cells were cultured in Roswell Park Memorial Institute 1640 Medium supplemented with GlutaMAX™ (RPMI1640-GlutaMAX™, Gibco^®^, catalog no. 61870036) mixed with 10% FBS (Gibco^®^, catalog no. 10437028) and 1% penicillin-streptomycin (Wako^®^, FUJIFILM^®^, catalog no. 168-23191). To induce macrophage differentiation, cells were seeded on 10-cm dishes at a density of 2.0×10^6^ cells per dish and treated with either 30 ng×mL^-1^ of PMA (Sigma-Aldrich^®^, catalog no. P8139) or, for the negative control, 0.1% (v/v) dimethyl sulfoxide (DMSO, Wako^®^, FUJIFILM^®^, catalog no. 043-07216) in the culturing medium for 24 hours and 96 hours.

#### Naïve and activated T cells (T cell Naïve and T cell Activated samples)

The CD4^+^ T cell samples used in this study and the data generated from them are fully described in Panariello *et al*. (in preparation). Human peripheral blood mononuclear cells (PBMCs) were purified from blood samples obtained from donors by density gradient centrifugation in Ficoll-Paque™ PLUS medium (Cytiva^®^, catalog no. 17144002). Naïve CD4^+^ T cells were initially negatively selected from PBMCs with a magnetic separator (autoMACS^®^ Pro Separator, Miltenyi Biotec^®^, catalog no. 130-092-545) with the Naïve CD4^+^ T Cell Isolation kit II (Miltenyi Biotec^®^, catalog no. 130-094-131). The purified cells were then further sorted by flow cytometry (BD FACSAria™ SORP, BD Biosciences^®^) as CD4^+^/CD45RA^+^ and CD4^+^/CD45RO^-^ cells, using the following antibodies: αCD4-VioGreen (Miltenyi Biotec^®^, catalog no. 130-113-221) or αCD4-APC-Cy7 (BD Bioscience, catalog no. 557871); αCD45RA-PECy5 (BD Bioscience^®^, catalog no. 552888); αCD45RO-APC (Miltenyi Biotec^®^, catalog no. 130-113-546) or αCD45RO-BV605 (BioLegend^®^, catalog no. 304237).

Naïve CD4^+^ T cells were activated using Dynabeads™ Human T-activator CD3/CD28 (Gibco^®^, catalog no. 1131D) *via* a 16-hour culture at a concentration of 1.5×10^6^ cells×mL^-1^ in T helper medium. The T helper medium is composed of RPMI1640-GlutaMAX™ (Gibco^®^, catalog no. 61870036), 10% FBS (Gibco^®^, catalog no. A5256701), 1% non-essential amino acids (Euroclone^®^, catalog no. ECB3054D), 1 mM sodium pyruvate (Euroclone^®^, catalog no. ECM0542D), 50 U×mL^-1^ penicillin/50 µg×mL^-1^ streptomycin (Euroclone^®^, catalog no. ECB3001D) and supplemented with cytokines (Marasca *et al*., 2022), namely 20 IU×mL^-1^ recombinant IL-2 (Miltenyi Biotec^®^, catalog no. 130-097-744), 10 ng×mL^-1^ recombinant IL-12 (Miltenyi Biotec^®^, catalog no. 130-096-704) and 2 µg×mL^-1^ neutralizing anti-IL-4 (Miltenyi Biotec^®^, catalog no. 130-095-753). Naïve and activated T cells were cultured at 37°C in a 5% CO_2_ incubator.

#### Healthy breast epithelial cells (MCF10A sample)

MCF10A cells, originally derived from the fibrocystic mammary tissue of a 36-year-old patient (Soule *et al*., 1990), were acquired from the American Tissue Culture Collection (ATCC^®^; CRL-10317™) and cultured in DMEM/F-12 with 4-(2-hydroxyethyl)-1-piperazineethanesulfonic acid (HEPES, Gibco^®^, catalog no. 31330038) supplemented with 5% horse serum (Gibco^®^, catalog no. 16050122), 20 ng×mL^-1^ human epidermal growth factor (PeproTech^®^, catalog no. AF-100-15-500UG), 500 ng×mL^-1^ hydrocortisone (Sigma-Aldrich^®^, catalog no. H0135), 100 ng×mL^-1^ cholera toxin (Bio Academia^®^, catalog no. BAM-01-511) and 10 µg×mL^-1^ insulin (Sigma-Aldrich^®^, catalog no. I1882).

#### Breast cancer cells (MCF7 sample) and estrogen deprivation-resistant breast cancer cells (LTED sample)

MCF7 breast cancer cells, originally collected from the pleural effusion of a 69-year-old patient (Soule *et al*., 1973), were acquired from ATCC^®^ (HTB-22™) and cultured in RPMI1640-GlutaMAX™ (Gibco^®^, catalog no. 61870036) supplemented with 10% FBS (Gibco^®^, catalog no. 10270-106) and 1% penicillin-streptomycin (Wako^®^, FUJIFILM^®^, catalog no. 168-23191).

MCF7 cells resistant to long-term estrogen deprivation (LTED; Masamura *et al*., 1995) were isolated and maintained by culturing MCF7 cells in phenol red-free Roswell Park Memorial Institute 1640 Medium (RPMI1640; Nacalai Tesque^®^, catalog no. 06261-65) containing 4% charcoal-stripped FBS (Thermo Fisher Scientific^®^, catalog no. 12676029) for 3 to 8 months at 37°C in a humidified 5% CO_2_ atmosphere.

#### Colon cancer cells (HCT116 sample)

HCT116 cells, originally collected from the colorectal carcinoma of a 48-year-old patient (Brattain *et al*., 1981), were acquired from ATCC^®^ (CCL-247™) and cultured in either DMEM, low glucose, pyruvate (Gibco^®^, catalog no. 11885084; for CAGE experiments) or DMEM (Gibco^®^, catalog no. 10437028) supplemented with 1% sodium pyruvate (Gibco^®^, catalog no. 11360070; for RADICL-seq). In both cases, the culture medium also included 10% FBS (Gibco^®^, catalog no. 10437028 or 10500056), and 1% penicillin-streptomycin (Wako^®^, FUJIFILM^®^, catalog no. 168-23191 or Gibco^®^, catalog no. 15140122).

#### Chronic myelogenous leukemia cells (K562 sample)

K562 lymphoblast cells, originally collected from the bone marrow of a 53-year-old chronic myelogenous leukemia patient (Klein *et al*., 1976), were acquired from ATCC^®^ (CCL-243™). Cells were cultured in Iscove’s Modified Dulbecco’s Medium (IMDM, SAFC^®^, catalog no. FG0465) mixed with 10% FBS (Gibco^®^, catalog no. A5670701).

#### Acute myeloid leukemia cells (AML sample)

Bone marrow samples from two AML patients with normal karyotypes were obtained at diagnosis. Mononuclear cells (BM-MNC) were isolated by density gradient centrifugation in Ficoll^®^ medium (Cytiva^®^, catalog no. 17-1440-03) and frozen in liquid nitrogen. The samples were thawed in prewarmed RPMI1640 (Gibco^®^, catalog no. 72400047) mixed with 20% FBS (Gibco^®^, catalog no. 10500064) containing DNase I (Sigma-Aldrich^®^, catalog no. D4513). Cells were gradually diluted, then centrifuged at 300 *g* for 10 min at room temperature, before being resuspended in cold RPMI1640 mixed with 10% FBS and filtered to remove aggregates.

For staining and fluorescence-activated cell sorting (FACS), cells were divided into fractions for unstained controls, fluorescence-minus-one (FMO) controls, and full staining. Staining was performed on ice and included incubation with the LIVE/DEAD™ Fixable Aqua Dead Cell Stain (Invitrogen™, catalog no. L34957) and the antibodies Brilliant Violet 421™ anti-human CD45 (BioLegend^®^, catalog no. 304031), PE anti-human CD19 (BioLegend^®^, catalog no. 302207), BD™ CD3 APC (BD Biosciences^®^, catalog no. 340661) and FITC anti-human CD335 (NKp46) (BioLegend^®^, catalog no. 331921). Following staining, cells were washed with FACS buffer constituted of DPBS (Gibco^®^, catalog no. 14190144) containing 2% FBS and 1 mM ethylenediaminetetraacetic acid (EDTA; Invitrogen™, catalog no. AM9260G). FACS was conducted on a gated CD45^+^ population, followed by negative gating to exclude CD3^+^ T lymphocytes, CD19^+^ B lymphocytes, and CD335^+^ natural killer cells. Sorted cells were collected in RPMI1640 mixed with 50% FBS, then pelleted, counted, and crosslinked in freshly prepared 2% formaldehyde (Thermo Fisher Scientific^®^, catalog no. 410731000). The reaction was quenched with 2.5 M glycine (Sigma-Aldrich^®^, catalog no. G7126), and cells were finally washed, centrifuged, and snap-frozen in liquid nitrogen for downstream processing.

### CFC-seq library preparation

Sample preparation for cap-trap full-length cDNA sequencing (CFC-seq) in the Neuron and THP-1 series is fully detailed in Yip *et al*. (2024). The same protocol, with some minor modifications, was applied to the T cell samples, as described in Panariello *et al*. (in preparation).

Briefly, RNAs were extracted and purified from cells using the RNAeasy^®^ Kit (QIAGEN^®^, catalog no. 74104) with on-column DNase I treatment, following the manufacturer’s instructions (RNAeasy^®^ Mini Handbook). After quantification by NanoDrop™ spectrophotometry (Thermo Fisher Scientific^®^), 5–10 µg of RNAs from each sample of the Neuron series, 80–100 µg of RNAs from each sample of the THP-1 series, and 3.8–5 µg of RNAs from each samples of the T cell series were polyadenylated by incubation with *E-coli* poly(A) Polymerase (PAP; New England Biolabs^®^, catalog no. M0276). These RNAs were then reverse-transcribed using the PrimeScript™ II Reverse Transcriptase (Takara^®^, catalog no. 2690) for the Neuron and THP-1 series, and a mix of SuperScript^®^ IV Reverse Transcriptase (Invitrogen™, catalog no. 18090050) and Induro^®^ Reverse Transcriptase (New England Biolabs^®^, catalog no. M0681) for the T cell samples. For this step, oligo dT primers containing unique molecular identifier (UMI) “GAGATGTCTCGTGGGCTCGGN_15_CTACGT_16_VN” were used in the Neuron series, and UMI “GAGATGTCTCGTGGGCTCGG[NB01-NB08]CTACGT_16_VN” in the THP-1 series and T cell samples.

Cap-trapping of the RNA/cDNA hybrids was performed as described in Delobel *et al*. (2025), followed by RNase H (Takara^®^, catalog no. 2150) digestion. Double stranded 5’ linkers of N6 and GN5 were ligated to the product using the Mighty Mix DNA ligation kit (Takara^®^, catalog no. 6023) and, for the Neuron and THP-1 series, removed of their phosphates by a shrimp alkaline phosphatase (SAP; Takara^®^, catalog no. 2660) treatment. Second strand synthesis of the 5’ linker-ligated cDNAs was performed with the KAPA HiFi™ DNA polymerase (Kapa Biosystems^®^, catalog no. 07958838001) followed by Exonuclease I (Takara^®^, catalog no. 2650) treatment to eliminate the excess primers.

The resulting cDNA/DNA hybrids were amplified for 10 cycles with the PrimerSTAR^®^ GXL DNA polymerase (Takara^®^, catalog no. R050) for the samples of the Neuron series and with the LongAmp^®^ Taq DNA polymerase (New England Biolabs^®^, catalog no. M0323) for the T cell samples; no amplification was performed in the THP-1 series. In the T cell samples, the amplified products were digested by Exonuclease I and purified by a 2:1 ratio of SPRIselect™ beads (Beckman Coulter^®^, catalog no. B23318) to remove fragments shorter than 600 base pairs (bp).

Libraries were prepared for both the Neuron and THP-1 series using the SQK-LSK110 DNA ligation kit (Oxford Nanopore Technologies^®^) following the manufacturer’s instructions (Document GDE_9108_v110_revV_10Nov2020), then sequenced on R9.4 flow cells (Oxford Nanopore Technologies^®^, catalog no. FLO-MIN-106) on a MinION device (Oxford Nanopore Technologies^®^). For the T cell samples, libraries were prepared using the SQK-LSK114 DNA ligation kit (Oxford Nanopore Technologies^®^) following the manufacturer’s instructions (Document GDE_9161_v114_revAA_30May2025), then sequenced on R10.4.1 flow cells (Oxford Nanopore Technologies^®^, catalog no. FLO-PRO114M) on a PromethION device (Oxford Nanopore Technologies^®^).

### CFC-seq data processing

#### Read alignment

CFC-seq data were processed as described in Yip *et al*. (2024).

Briefly, after basecalling by Dorado version 0.2.4 (Oxford Nanopore Technologies^®^) with the --minqscore 10 option, linkers were trimmed and reads were orientated using primer-chop version 1418 (M. Frith, 2022). Poly(A) tails were then removed to only keep up to a maximum of 5 consecutive As, thanks to our in-house tool tail_trimmer version 1.4 (Yip *et al*., 2024).

Reads were aligned to the hg38 genome using Minimap2 version 2.26 (H. Li, 2018) with the splice preset and the -u f option, keeping only primary alignments and using the GENCODE annotation version 39 for splice junction information. The resulting .sam files were converted to .bam format and sorted using functions view and sort from SAMtools version 1.11 (H. Li *et al*., 2009).

To construct the SACAGE annotation (see below), demultiplexed and trimmed reads were also aligned to the GENCODE transcriptome version 39 with the map-ont preset of Minimap2 and options -p 1, -N 100 and -u f. These raw alignments were then corrected by TranscriptClean version 2.0.3 (Wyman & Mortazavi, 2019), using the junction information from GENCODE version 39 combined with confident junctions extracted from our short-read RNA-seq data in the Neuron series described below.

#### Construction of the SACAGE annotation

The identification of new transcript models derived from CFC-seq data in the Neuron and THP-1 series is fully described in Yip *et al*. (2024).

Briefly, high-confidence transcription start site (TSS) clusters were first detected from the genome-aligned reads of each individual sample in both series, using the Single Cell Analysis of Five-prime Ends (SCAFE) pipeline version 1.01 (Moody *et al*., 2022). Strand-specific TSS clusters within 75 nucleotides (nt) from each other were next merged across samples with the aggregate workflow of SCAFE, then split at the midpoint of the two summits of their RLE-normalized signal (edgeR; Robinson *et al*., 2009) in case of bi-modal distribution. In parallel, 3’ end clusters were defined using Paraclu (M. C. Frith *et al*., 2008) on the 3’ end tags from each CFC-seq sample of both series, keeping only clusters with ≥3 assigned tags.

TSS clusters defined by SCAFE, 3’ end clusters defined by Paraclu, transcript models from GENCODE version 39, and splicing junctions extracted both from the transcriptome-aligned CFC-seq reads and from our short-read RNA-seq data described below were all used as input for the Start-site Aware Long-read Assembler (SALA; Yip *et al*., 2024) to predict confident 5’ end clusters, 3’ end clusters, and splicing junctions. Like above, confident 5’ clusters located within 75 nt from each other or from a GENCODE 5’ end were merged, then split at the midpoint of the two summits of their signal in case of bi-modal distribution. Confident 3’ clusters located within 150 nt from each other or from a GENCODE 3’ end were also merged together.

Next, all CFC-seq reads for which both the 5’ and 3’ ends overlap the same confident 5’ and 3’ clusters, respectively, and for which all observed splicing junctions match were combined together as a single transcript model. The definitive start and end coordinates of a given transcript model were assigned to the most common 5’ and 3’ ends of all the reads that contributed to its determination. CFC-seq reads for which either or both end(s) do not correspond to a confident end cluster were deemed incomplete and were solely used to support the aforementioned transcript models that they intersect, if any.

GENCODE version 39 was used as a reference to classify all transcript models, leading to the recovery of known ENST transcripts as well as the identification of both novel transcripts derived from ENSG genes and novel transcripts derived from yet-unknown gene regions. The lncRNA transcript models, including the 115,275 novel ones identified with our method, were classified as divergent lncRNAs, sense overlap lncRNAs, sense intronic lncRNAs, antisense lncRNAs and intergenic lncRNAs, in this order of priority and relative to their overlap with GENCODE-annotated transcript models. To annotate regulatory regions, the TSS clusters identified previously by SCAFE were first extended into transcribed *cis*-regulatory elements (tCREs) and overlaps were merged, then classified as “promoter-like”, “enhancer-like”, “CTCF-alone” or “unclassed” by intersection with SCREEN data (The ENCODE Project Consortium, 2020). This information was transferred onto the transcript models obtained from SALA, and only tCREs located in open chromatin regions, as indicated by single-nucleus Assay for Transposase-Accessible Chromatin using sequencing (snATAC-seq; see below), or corresponding to regulatory regions already annotated in GENCODE version 39 were kept in the final annotation.

Finally, unmatched permissive gene models from the FANTOM5 CAGE-associated transcriptome (CAT) annotation (Hon *et al*., 2017) were added to these SALA-identified novel transcript models, then combined with GENCODE version 39. All together, these models form the FANTOM6 SACAGE (<u>SA</u>LA+<u>CA</u>T+<u>GE</u>NCODE) annotation, which was used as the reference hg38 annotation for all analyses in this study.

#### Differential expression analysis

For differential expression analysis, read counts for all transcript isoforms in the SACAGE annotation were calculated from the genome-aligned .bam files of each CFC-seq replicate, using the multicov function from BEDTools version 2.30.0 (Quinlan & Hall, 2010). Differential expression between samples was assessed using “edgeR” version 3.40.2 (Robinson *et al*., 2009) in R version 4.3.3 (R Core Team, 2023). Transcripts with a significantly different expression between two samples were defined as those with a *log*(fold change) (logFC) < −1.2 or > 1.2.

### CAGE library preparation

#### nAnT-iCAGE in the HDF sample

No-amplification non-tagging CAGE library preparation for Illumina (nAnT-iCAGE; Murata *et al*., 2014) in HDF cells was performed as described in Yip *et al*. (2022) and Agrawal *et al*. (2024).

Briefly, RNAs were extracted and purified from cells using the RNAeasy^®^ Kit (QIAGEN^®^, catalog no. 74104) with on-column DNase I treatment, following the manufacturer’s instructions (RNAeasy^®^ Mini Handbook). After quantification by NanoDrop™ spectrophotometry (Thermo Fisher Scientific^®^), 50 ng of RNA were used to synthesize cDNAs with a random N_6_+TCT primer and the SuperScript™ III Reverse Transcriptase (Invitrogen™, catalog no. 18080085) in presence of 0.14 M D-(+)- trehalose dihydrate (Life Sciences Advanced Technologies^®^, catalog no. TDH033) and 0.65 M D-sorbitol (Fluka™, catalog no. 85529-250g). RNA/cDNA hybrids were purified with Agencourt^®^ RNAClean™ XP beads (Beckman Coulter^®^, catalog no. A66514) and eluted by resuspending the beads for 5 min in water pre-heated at 37°C. Diol residues were oxidized by adding 250 mM sodium periodate (NaIO_4_; MP Biomedicals^®^, catalog no. 152577) on ice and away from light. The capped RNAs were then biotinylated using 10 mM Biotin (Long Arm) Hydrazide (Vector Laboratories^®^, catalog no. SP-1100) in DMSO (Wako^®^, FUJIFILM^®^, catalog no. 043-07216).

cDNAs hybridized to the biotinylated capped RNAs were next treated with RNase ONE™ ribonuclease (Promega^®^, catalog no. M4265), before selection with the CAP trapper method (Carninci *et al*., 1997) in 3.15 M NaCl (Promega^®^, catalog no. V4221) buffer supplemented with 0.1% Tween^®^ 20 (Sigma-Aldrich^®^, catalog no. P1379) and using 15 µL of Dynabeads™ M-270 Streptavidin beads (Invitrogen™, catalog no. 65306) that were coated beforehand with 3.75 µg of tRNAs for 30 min. The cDNAs were then dissociated from the beads with 65 µl RNase ONE™ buffer at 95°C for 5 min, then treated once with RNase H (Takara^®^, catalog no. 2150) and twice with RNase ONE™, each time with a purification step using AMPure™ XP beads (Beckman Coulter^®^, catalog no. A63882) then an elution in 42 µL water pre-heated at 37°C.

The resulting single-strand cDNAs were first ligated to a sample-specific 5’ linker (Murata *et al*., 2014) containing a recognition site for barcodes “ACC” or “CAC” with the Mighty Mix DNA ligation kit (Takara^®^, catalog no. 6023) and concentrated with a SpeedVac™ vacuum concentrator (Thermo Fisher Scientific^®^). Ligated cDNAs were purified twice with AMPure™ XP beads to eliminate the unligated linkers, then re-concentrated by SpeedVac™. cDNAs were then ligated to a second 3’ linker (Murata *et al*., 2014) and once again purified as described above. Following this second ligation, the upper strands of the 5′ and 3′ linkers were degraded by treatment with SAP (Applied Biosystems™, catalog no 783905000UN) and USER™ enzyme (New England Biolabs^®^, catalog no. M5505). The cDNAs were then once again subjected to AMPure™ XP beads purification and SpeedVac™ concentration.

The second strands of the cDNAs were synthesized by the Deep Vent^®^ (exo-) DNA Polymerase (New England Biolabs^®^, catalog no. M0259) with the nAnT-iCAGE second primer (Murata *et al*., 2014). The resulting double-stranded cDNAs were subjected to an Exonuclease I (New England Biolabs^®^, catalog no. M0293) treatment in order to eliminate the extra primers, followed by two AMPure™ XP beads purification and SpeedVac™ concentration.

Libraries were sequenced in single-end on a HiSeq™ 2000 system with the TruSeq^®^ SBS Kit v3-HS (200 cycles, Illumina^®^, catalog no. FC-401-3001).

#### ssCAGE in the Neuron and THP-1 series

Samples from the Neuron series and the THP-1 series were processed in duplicates as follows. RNAs were extracted and purified from cells using the RNAeasy^®^ Kit (QIAGEN^®^, catalog no. 74104) with on-column DNase I treatment, following the manufacturer’s instructions (RNAeasy^®^ Mini Handbook). After quantification by NanoDrop™ spectrophotometry (Thermo Fisher Scientific^®^), 5 μg of RNA were used from each sample to construct a single-stranded CAGE library using a modified version of the CAGE protocol, which combines the nAnT-iCAGE protocol described above (Murata *et al*., 2014) and low quantity single strand CAGE (LQ-ssCAGE; Takahashi *et al*., 2021). Reverse transcription of RNAs was carried out as in nAnT-iCAGE, then Cap-trap, 5’ linker and 3’ indexed linker ligation steps were performed following the LQ-ssCAGE protocol, changing the second round of purification from 1.2× Agencourt^®^ AMPure™ XP beads (Beckman Coulter^®^) to 0.8× SPRIselect™ beads (Beckman Coulter^®^, catalog no. B23318) for a more stringent removal of free linkers and linker dimers.

Libraries were sequenced in paired-end for 100 cycles on a NextSeq™ 1000/2000 system with the corresponding P2 Reagents Kit v3 (200 cycles, Illumina^®^, catalog no. 20046812) for the Neuron series, and on a HiSeq™ 2500 system with the corresponding Rapid SBS Kit v2 (200 cycles, Illumina^®^, catalog no. FC-402-4021) for the THP-1 series.

#### SLIC-CAGE in HCT116 cells

HCT116 cells were cultured to 75-80% confluence before trypsinization with 0.25% Trypsin-EDTA (Gibco^®^, catalog no. 25200056) and splitting 1:5-1:8. Sample were harvested at passage 5, washed with phosphate-buffered saline (PBS; Gibco^®^, catalog no. 10010031) and counted. 3 replicates of 2.5×10^5^ cells each were collected and immediately resuspended in 300 µL of RNA lysis buffer from the PureLink™ RNA Mini Kit (Invitrogen™, catalog no. 12183025) including 1% *β*-mercaptoethanol (Sigma-Aldrich^®^, catalog no. M6250). Total RNA from all 3 replicates was extracted with the PureLink™ RNA Mini Kit according to the manufacturer’s instruction. On-column DNA digestion was performed using the corresponding PureLink™ DNase Set (Invitrogen™, catalog no. 12185010) and the quality of the extracted total RNA in each replicate was assessed using the automated Bioanalyzer 2100 gel electrophoresis system (Agilent^®^) with the corresponding RNA 6000 Pico kit (Agilent^®^, catalog no. 5067-1513).

2 µg of RNA per replicate were used for the Super-Low Input Carrier-CAGE (SLIC-CAGE) protocol published in Cvetesic *et al*. (2018). Briefly, 3 µg of carrier RNA were added prior to first-strand cDNA synthesis. Subsequently, cDNA samples were oxidized and biotinylated, and non-hybridized, overhanging RNA stretches were removed by treatment with RNase ONE™ ribonuclease (Promega^®^, catalog no. M4265) followed by a pull-down with Dynabeads™ M-270 Streptavidin beads (Invitrogen™, catalog no. 65305) to separate fully-reverse transcribed and capped RNAs from non-capped RNA and/or partially transcribed RNA species. Complementary DNA was released from the beads and separated from the RNA by denaturing the cDNA/RNA hybrids bound to the paramagnetic Streptavidin beads *via* the capped RNA and by incubating the samples with RNase H (Takara^®^, catalog no. 2150A) and RNase ONE™. Libraries were prepared by 5’ and 3’ nAnT-iCAGE linker ligation, second-strand DNA synthesis, carrier digestion, size selection and PCR amplification. The fragment distribution of the resulting libraries was quantified using the automated Bioanalyzer 2100 gel electrophoresis system with the corresponding High Sensitivity DNA kit (Agilent^®^, catalog no. 2067-4626). Precise concentrations were then measured with the Quant-iT™ PicoGreen™ dsDNA Assay kit (Invitrogen™, catalog no. P7589). All 3 replicates were finally sequenced together in single-end on a NextSeq™ 550 system with the corresponding High Output Kit v2.5 (75 cycles, Illumina^®^, catalog no. 20024906).

### CAGE data processing

nAnT-iCAGE data in the iPSCpreF6 sample were originally generated by Agrawal *et al*. (2024) and are accessible on the DNA Data Bank of Japan (accession DRA013848). CAGE data in MCF10A cells were originally generated by Watanabe *et al*. (2019) and were retrieved from the Gene Expression Omnibus database (accession GSE124843). ssCAGE data in MCF7 and LTED cells were generated by Shu *et al*. (in preparation) and are accessible on the Gene Expression Omnibus database (accession GSE299910). CAGE data in K562 cells were downloaded from the ENCODE database (accession ENCSR000CJN; The ENCODE Project Consortium, 2012).

#### Read alignment

nAnT-iCAGE reads in the iPSCpreF6 and HDF samples were demultiplexed, removed of primer dimers and rRNAs, and finally aligned to the hg38 genome assembly using TopHat version 2.0.12 (D. Kim *et al*., 2013) with a minimum anchor length of 20 within the MOIRAI pipeline (Hasegawa *et al*., 2014). The same processes were performed with the MOIRAI pipeline on CAGE and ssCAGE reads in the Neuron series, the THP-1 series, and the MCF10A, MCF7, LTED and K562 samples, using instead STAR (Dobin *et al*., 2012) version 2.5.3a (for the MCF7 and LTED samples) or 2.5.0a (for all other samples) with options --outFilterMultimapNmax 20 (for all), --alignIntronMin 20 and --alignIntronMax 1000000 (for the Neuron and THP-1 series and the MCF7 and LTED samples) and --twopassMode None (for the Neuron and THP-1 series and the MCF10A and K562 samples).

SLIC-CAGE reads in the HCT116 sample were first demultiplexed using bcl2fastq version 1.8.4 (Illumina^®^). The quality of the sequencing data was validated with FastQC version 0.11.9 (Andrews, 2010) and compared with MultiQC version 1.11 (Ewels *et al*., 2016). The first 5 bases from each read corresponding to the SLIC-CAGE adapter as well as low-quality bases at the 3’ end were trimmed with cutadapt version 4.4 (Martin, 2011). Reads were then filtered to keep only those longer than 30 bp, with a minimum base quality of 30 and > 50% passing percentage using FASTX-Toolkit version 0.0.14 (Gordon, 2014). rRNA-derived reads were removed with rRNAdust version 1.06 (http://fantom.gsc.riken.jp/5/suppl/rRNAdust/). These filtered reads were aligned to the hg38 genome assembly using BWA version 0.7.17 (H. Li & Durbin, 2009) with default parameters, before being converted to .bam format, sorted and indexed with the view, sort and index functions of SAMtools version 1.17 (H. Li *et al*., 2009).

#### Transcription start site identification

For each sample, the CAGE .bam files of all replicates were submitted together to the SCAFE pipeline version 1.0.0 (Moody *et al*., 2022) with the aggregate workflow and GENCODE version 39 for the annotation step. For the K652 data only, the min_total_match parameter in the scafe.tool.bk.bam_to_ctss script was changed from 30 to 20, to palliate for the fact that these previously published data were generated using an older CAGE protocol, with an expected read length of 27 bp (Kodzius *et al*., 2006; The ENCODE Project Consortium, 2012; Valen *et al*., 2009).

TSS identification was also performed in the T cell samples, using the CFC-seq data described above instead. For these samples, we used SCAFE version 1.0.1, which is better suited to the CFC-seq technology and is fully described in Yip *et al*. (2024). This version of SCAFE was additionally leveraged to generate a unique list of TSS clusters in each of the Neuron and THP-1 series, using the SCAFE aggregate workflow with all samples of each series.

TSS clusters identified by SCAFE were labeled as “promoter” if they overlap the ±500 bp region surrounding the start coordinate of any coding gene or pseudogene in the SACAGE annotation, “exonic” if they overlap any exon of a coding gene or pseudogene (excluding the first 500 bp of the first exon), “intronic” if they overlap any intron of a coding gene or pseudogene, or “intergenic” if they do not overlap any coding gene or pseudogene, in this order of priority.

#### Differential TSS activity analysis

To assess for differential TSS activity in the Neuron and THP-1 series, we used the TSS clusters obtained from SCAFE with the aggregate workflow, both directly and after extending these regions by ±500 bp. The coverage of these TSS clusters was calculated from the aligned reads in each replicate for each sample of the relevant series, using the multicov function from BEDTools version 2.30.0 (Quinlan & Hall, 2010). Differential expression between samples was assessed using “edgeR” version 3.40.2 (Robinson *et al*., 2009) in R version 4.3.3 (R Core Team, 2023). TSS clusters with a significantly different activity between two samples were defined as those with logFC < −2 or > 2 and FDR < 0.01.

### RNA-seq library preparation

Total RNAs were extracted and purified from each sample of the Neuron series using the RNAeasy^®^ Kit (QIAGEN^®^, catalog no. 74104) with on-column DNase I treatment, following the manufacturer’s instructions (RNAeasy^®^ Mini Handbook). After quantification by NanoDrop™ spectrophotometry (Thermo Fisher Scientific^®^), 0.5 µg of RNAs were used to prepare libraries with the TruSeq^®^ Stranded Total RNA Sample kit with Ribo-Zero™ Human/Mouse/Rat (Illumina^®^, catalog no. 20020596) and the TruSeq^®^ RNA Single Indexes Set A (12 indexes, 24 samples; Illumina^®^, catalog no. 20020492). Sequencing was performed in paired-end with a Novaseq™ 6000 system on a single lane of an S4 flow cell (200 cycles, Illumina^®^, catalog no. 20028313).

### RNA-seq data processing

Paired-end reads were aligned to the hg38 reference genome using the hisat2-align-s function from HISAT2 version 2.2.1 (D. Kim *et al*., 2019) with options --sensitive and --read-lengths 150. Output files were further processed with SAMtools version 1.11 (H. Li *et al*., 2009): mapped reads were filtered to keep only those that were primarily aligned with MAPQ ≥ 10, then converted to .bam format with the view function before being sorted and indexed with the sort and index functions.

RNA-seq read counts on all SACAGE transcript isoforms were calculated using the multicov function from BEDTools version 2.30.0 (Quinlan & Hall, 2010).

### RADICL-seq library preparation

RNA And DNA Interacting Complexes Ligated and sequenced (RADICL-seq) was performed on all samples as described in Bonetti *et al*. (2020), Shu *et al*. (2024) and Pracana *et al*. (in preparation), with some modifications and with two to three biological replicates per sample.

Briefly, pellets containing approximately 2×10^6^ cross-linked cells were resuspended in cold lysis buffer (Pracana *et al*., in preparation; Shu *et al*., 2024) with 1 mM phenylmethylsulfonyl fluoride (PMSF; Roche^®^, catalog no. 10837091011), 1× cOmplete™ Protease Inhibitor Cocktail (Roche^®^, catalog no.

4693116001), 0.8 U×μL^-1^ RNasin^®^ Plus (Promega^®^, catalog no. N2611) and incubated on ice for 10 min. Nuclei were pelleted at 2,500 *g* for 60 seconds, resuspended in DNase I digestion buffer I (Pracana *et al*., in preparation; Shu *et al*., 2024) for permeabilization and then subjected to DNase I digestion buffer II (Pracana *et al*., in preparation; Shu *et al*., 2024) to quench the sodium dodecyl sulfate (SDS) present in buffer I. Next, gDNA digestion was carried out with 1.5 U DNase I (Thermo Scientific™, catalog no. EN0525) at room temperature under rotation, and the reaction was subsequently terminated by adding DNase I stop buffer (Pracana *et al*., in preparation; Shu *et al*., 2024) to the samples. The resulting nuclei were resuspended in nuclease-free water (Invitrogen™, catalog no. 10977015) and purified with RNAClean™ XP beads (Beckman Coulter^®^, catalog no. A63987).

Chromatin ends were repaired by incubating the bead-attached nuclei with T4 DNA ligase (Thermo Scientific™, catalog no. EP0061) and Klenow Fragment (Thermo Scientific™, catalog no. EP0051), stopping the reaction with 10% SDS (Invitrogen™, catalog no. 15553-027), and finally dA-tailing with Klenow Fragment exo- (Thermo Scientific™, catalog no. EP0421). RNA-DNA interactions coming from nascent transcripts were next removed by treatment with RNase H (New England Biolabs^®^, catalog no. M0297S), then the bead-nuclei mixture was pelleted at 2,500 *g* for 60 seconds and resuspended in 200 μL water. To remove all soluble RNAs, two steps of polyethylene glycol (PEG) purification were carried out by adding 165 µL of 20% PEG (Promega^®^, catalog no. V3011) in 2.5 M NaCl (Invitrogen™, catalog no. AM9759) to the bead-attached nuclei, purifying them with RNAClean™ XP, and resuspending them in 200 μL water.

The RNA and DNA fragments were then bound by proximity ligation with pre-adenylated and biotinylated bridge adaptors. First, these adaptors were ligated to the 3’-OH end of the RNA molecules with T4 RNA ligase 2, truncated KQ (New England Biolabs^®^, catalog no. M0373S), before stopping the reaction with Triton™ X-100 (Sigma-Aldrich^®^, catalog no. T8787-50ML), pelleting the nuclei at 2,500 *g* for 60 seconds and resuspending them in 200 µL water. Excess unligated adaptors were removed by two rounds of RNAClean™ XP purification. Second, the RNA-bound adaptors were ligated *in situ* to proximal DNA fragments by incubation with T4 DNA ligase (New England Biolabs^®^, catalog no. M0202S), then nuclei were once more pelleted and purified as described above and finally resuspended in 200 µL water.

Cross-linking was reversed by treating the bead-attached nuclei with Recombinant Proteinase K (Invitrogen™, catalog no. AM2546). The RNA-DNA complexes were then precipitated with isopropanol (Wako^®^, FUJIFILM^®^, catalog no. 166-04836), purified again with RNAClean™ XP beads, and concentrated with a SpeedVac™ vacuum concentrator (Thermo Fisher Scientific^®^).

Reverse transcription of the adaptor-bound RNAs was carried out by incubating the samples with SuperScript^®^ IV Reverse Transcriptase (Invitrogen™, catalog no. 18090010), and second strand synthesis was performed using *E. coli* DNA Polymerase I (New England Biolabs^®^, catalog no. M0209S), RNase H and *E. coli* DNA Ligase (New England Biolabs, catalog no. M0205S). The resulting products were purified using the QIAquick™ Nucleotide Removal Kit (QIAGEN^®^, catalog no. 28306) and re-concentrated by SpeedVac™.

To ensure that only cDNA-bridge adaptor-gDNA complexes would be sequenced, hairpin linker “5’-/5Phos/GGCCCTCCAAAAGGAGGGCA-3’” (Integrated DNA Technologies^®^) was ligated specifically to cDNA-bound adaptors missing the gDNA side with the T4 DNA ligase (Promega^®^, catalog no. M8221). The samples were next purified with the QIAquick™ PCR Purification kit (QIAGEN^®^, catalog no. 28104) following the manufacturer’s instructions (Document 1114321 07/2018 HB-0900-003). After re-concentration by SpeedVac™, the samples’ concentration was measured with the Qubit™ 1X dsDNA High Sensitivity (HS) Assay Kit (Invitrogen™, catalog no. Q33230).

10 U×1.5 µg DNA^-1^ of EcoP15I (New England Biolabs^®^, catalog no. R0646) were used to digest the double-stranded cDNA-adaptor-gDNA molecules and cleaned with the QIAquick™ Nucleotide Removal Kit. The digested samples were prepared for sequencing linker ligation with the NEBNext^®^ Ultra™ II End Prep Reaction buffer and Enzyme Mix (New England Biolabs^®^, catalog no. E7645S) and ligated to Y-shaped sequencing linkers with the NEBNext^®^ Ultra™ II Ligation Master Mix and Ligation Enhancer (New England Biolabs^®^, catalog no. E7645S).

The cDNA-adaptor-gDNA complexes were next pulled-down with Dynabeads™ MyOne™ Streptavidin C1 beads (Invitrogen™, catalog no. 65001) thanks to their biotinylated bridge adaptor and amplified by Phusion™ High-Fidelity DNA Polymerases (Thermo Scientific™, catalog no. F530S) in four independent PCR reactions using a different barcoded primer for each. The amplified libraries were purified first using the QIAquick™ PCR Purification kit, then using Novex™ TBE-Urea Gels, 6% (Invitrogen™ catalog no. EC6865BOX) and excising only the targeted 225 bp band. The resulting materials were cleaned by ethanol precipitation after overnight elution.

Library size was assessed using the Agilent^®^ High Sensitivity DNA Kit (Agilent^®^, catalog no. 5067-4626) and measured by quantitative PCR using the Library Quantification Kit for Illumina^®^ sequencing platforms (Kapa Biosystems^®^, catalog no. KK4835) together with a StepOne™ Real-Time PCR System (Applied Biosystems^®^). Libraries were finally sequenced in single-end on a Novaseq™ 6000 system with the corresponding S2 Reagent Kit v1.5 (200 cycles, Illumina^®^, catalog no. 20028315).

### RADICL-seq data processing

RADICL-seq data was processed as fully described in Pracana *et al*. (in preparation).

#### Demultiplexing, deduplication, trimming, filtering and splitting

Briefly, raw reads were first demultiplexed using cutadapt version 4.1 (Martin, 2011) with the --discard-untrimmed and --action retain options, then deduplicated using the clumpify.sh script from BBMap version 39.01 (Bushnell, 2022) with the dedupe=f and subs=2 parameters. Sequencing linkers were trimmed using cutadapt with the --discard-untrimmed and --action trim options. Reads that did not contain the bridge adapter were discarded by running cutadapt with the --discard-untrimmed, --revcomp and --action none options.

Next, read sequences were split into two files, one containing the RNA side and the other the DNA side. To extract the RNA sequences, cutadapt was run with the --discard-untrimmed and --action trim options, treating the bridge adapter like a 3’ adapter to remove it and everything after it, and keeping only trimmed reads ranging from 25 bp to 29 bp. For the DNA sequences, the same strategy was applied, this time treating the bridge adapter like a 5’ adapter to remove it and everything before it, and keeping only trimmed reads ranging from 24 bp to 28 bp. The trimmed and filtered DNA read fragments were then reverse complemented using the fastx_reverse_complement function from FASTX-Toolkit version 0.0.14 (Gordon, 2014). Only reads for which both an RNA and a DNA side could be extracted and passed the size filtering were kept for downstream processing.

#### Read alignment

The DNA read fragments were aligned to the hg38 reference genome using Bowtie 2 version 2.5.0 (Langmead & Salzberg, 2012) with the --fast preset from the --end-to-end mode and the -L 9 and -k 20 options. On the RNA side, read fragments corresponding to rRNAs were first removed using Bowtie 2 with the --sensitive-local parameter, using a rRNA-specific index created from the complete human rRNA-coding region catalog provided by the European Nucleotide Archive (accession U13369). These filtered RNA read fragments were then aligned to the hg38 reference genome with respect to the SACAGE annotation using STAR version 2.7.4a (Dobin *et al*., 2012) with the following parameters: --outFilterMultimapNmax 20, --outFilterIntronMotifs RemoveNoncanonicalUnannotated, --outFilterMismatchNmax 999, --alignSJoverhangMin 6, --alignSJDBoverhangMin 1, --twopassMode None, --alignEndsType EndToEnd, --outSAMstrandField intronMotif, --outSAMattributes All, --outFilterType BySJout and --outFilterMismatchNoverReadLmax 0.04.

Aligned read fragments from both sides were converted to .bam format, sorted and indexed with functions view, sort and index from SAMtools version 1.16.1 (H. Li *et al*., 2009). RNA fragments and DNA fragments originating from the same initial reads were then paired back together into a .bedpe file using packages “rtracklayer” version 1.58.0 (Lawrence *et al*., 2009), “GenomicFeatures” version 1.50.2 (Lawrence *et al*., 2013) and “Rsamtools” version 2.14.0 (Morgan *et al*., 2017) in R version 4.2.3 (R Core Team, 2023). Only RNA-DNA pairs for which both read fragments uniquely mapped to the genome were kept, and .bedpe files from all technical replicates of the same biological replicate were merged together. Furthermore, to compensate for low sequencing depth, two biological replicates of DMSO 96h (rep1 and rep3) were merged to create the final DMSO 96h rep1 analyzed in this study.

#### CHiCANE filtering

RNA read fragments were annotated using the SACAGE annotation at the gene level with R packages “rtracklayer” and “GenomicFeatures”, considering only fragments the coordinates of which were fully enclosed within those of their source gene locus. In parallel, each DNA read fragment was assigned to a 25-kb genomic window. Read counts for each pair of source gene and 25-kb target window were then calculated in R and used to filter significant interactions with Capture Hi-C Analysis Engine (CHiCANE) version 0.1.8 (Holgersen *et al*., 2021). The .bedpe files created previously were filtered to keep only reads corresponding to gene/window pairs with a CHiCANE false discovery rate (FDR) < 0.01. These filtered reads were used for all further analyses.

Unless otherwise specified, interaction counts from the CHiCANE-filtered reads were calculated at the transcript isoform resolution on the source side, by overlapping the RNA fragments of these reads with all transcript isoforms listed in the SACAGE annotation thanks to the intersect function from BEDTools version 2.30.0 (Quinlan & Hall, 2010). This same function was used to overlap the DNA read fragments with the genomic regions indicated for each analysis (25-kb windows, target site clusters, TSS clusters, and so on). The number of fragments pairs corresponding to each transcript/target region combination were then counted the help of the “data.table” package version 1.14.8 (Dowle & Srinivasan, 2023) in R version 4.2.3 (R Core Team, 2023). Self interactions were defined as all interactions for which the RNA and DNA fragments of the read overlap the same transcript locus.

#### Multidimensional scaling and adjusted Rand index calculation

Similarity between biological replicates was quantified based on the above-mentioned interaction counts. To account for differences in sequencing depth, counts were normalized by the total number of interactions per replicate. The resulting values were scaled to counts per million and log-transformed. Pairwise similarity between samples was assessed using the Pearson correlation coefficient computed across all interactions. Correlation values (*r*) were converted into Euclidean distance metrics (*d*) with:

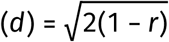

Multidimensional scaling (MDS) was then performed on the resulting distance matrix using the cmdscale function in R version 4.5.1 (R Core Team, 2023).

Based on the MDS results, samples were visually grouped in five biological categories: blood cells (DMSO 96h, PMA 24h, PMA 96h, T cell Naïve, T cell Activated, AML and K562), epithelial cells (MCF10A, MCF7, LTED and HCT116), neural cells (NSC and Neuron), iPSCs (iPSC and iPSCpreF6) and fibroblasts (HDF). To quantify the extent to which this visual grouping accurately reflects RADICL-seq-based sample similarity, hierarchical clustering was performed on the same distance matrix using Ward’s method (Ward, 1963). The resulting dendrogram was cut into *k* = 5 clusters, then agreement between this clustering and our visual grouping was assessed by the adjusted Rand index (ARI) implemented in R package “mclust” version 6.1.1 (Scrucca *et al*., 2023). To evaluate statistical significance, a permutation-based null distribution was generated by randomly reassigning biological group labels while preserving replicate structure, and recomputing the ARI across 1,000 iterations. The empirical *p* value was calculated as the proportion of permuted ARI values greater than or equal to the observed ARI.

### snATAC-seq library preparation

For single nucleus ATAC-seq (snATAC-seq), iPSC, NSC and Neuron cells were harvested by incubation in a buffer containing 50% Accutase^®^ (Innovative Cell Technologies^®^, catalog no. AT104), 50% DPBS (Wako^®^, FUJIFILM^®^, catalog no. 049-29793) and ≥50 units papain enzyme (Worthington Bio-chemical Corporation^®^, catalog no. LK003178) at 37°C for 20 min. After washing with DMEM/F-12 supplemented with GlutaMAX™ (Gibco^®^, catalog no. 10565018), 10 nM ROCK inhibitor (CultureSure^®^ Y-27632, Wako^®^, FUJIFILM^®^, catalog no. 036-24023), and ≥500 Kunitz units DNase I (Worthington Biochemical Corporation^®^, catalog no. LK003172), the cells were collected and washed 3 times with DPBS, then centrifuged at 150 *g* for 10 min to remove the debris.

Nuclei isolation was performed as instructed in the “Nuclei Isolation for Single Cell ATAC Sequencing” protocol from 10x Genomics^®^ (Demonstrated Protocol CG000169 Rev D), with one modification. In brief, centrifuged cells were resuspended in phosphate-buffered saline (PBS; Wako^®^, FUJIFILM^®^, catalog no. 166-23555) with 0.1% (instead of 0.04%) bovine serum albumin (BSA; Invitrogen™, catalog no. AM2616), then once again centrifuged at 300 *g* for 5 min. The supernatant was replaced by a lysis buffer containing 10 mM Tris-HCl (Sigma-Aldrich^®^, catalog no. 10812846001), 10 mM NaCl (Sigma-Aldrich^®^, catalog no. S9888), 3 mM MgCl_2_ (Sigma-Aldrich^®^, catalog no. M1028), 0.1% Tween^®^ 20 (Sigma-Aldrich^®^, catalog no. P1379), 0.1% Nonidet™ P40 substitute (Thermo Fisher Scientific^®^, catalog no. J19628.K2), 0.01% digitonin (Invitrogen™, catalog no. BN2006) and 1% BSA for 5 min on ice. Lysed cells were then washed with a buffer containing 10 mM Tris-HCl and 10 mM Na, once more centrifuged at 500 *g* for 5 min, then resuspended in 1× Nuclei Buffer (10x Genomics^®^, catalog no. 2000153/2000207). snATAC-seq libraries were prepared using the Chromium™ Next GEM™ Single Cell ATAC Reagent Kits v1.1 (10x Genomics^®^, catalog no. 1000175) following the manufacturer’s instructions (Document CG000209 Rev G) and sequenced on a HiSeq™ X Ten system (Illumina^®^), with the iPSC, NSC and Neuron-derived materials all pooled into one lane.

### ATAC-seq data processing

#### Read alignment and peak calling in the Neuron series

snATAC-seq reads in the Neuron series were processed with the Cell Ranger™ ATAC pipeline version 2.0.0 (10x Genomics^®^) on the hg38 genome assembly to generate peak-to-cell count matrices.

These ATAC-seq data in the Neuron series were originally generated at a single-nucleus level for the study detailed in Yip *et al*. (in preparation). For the purpose of the present study, this data was converted into a bulk-like format comparable to the publicly available ATAC-seq data from Lin *et al*. (2022) used for the THP-1 series (see below). This was achieved by randomly splitting the individual cells defined in the count matrices generated by Cell Ranger™ in two artificial bulk replicates for each sample, using R version 4.3.3 (R Core Team, 2023) with packages “Seurat” version 5.1.0 (Hao *et al*., 2023) and “hdf5r” version 1.3.11 (Hoefling & Annau, 2017). Overlapping peaks detected in several samples were merged together, keeping the minimum and maximum coordinates of all merged peaks as the new coordinates, and counts in each artificial replicate were summed using the merge and intersect functions from BEDTools version 2.30.0 (Quinlan & Hall, 2010), in order to create a set of unique, non-overlapping peak regions for the whole series.

#### Read alignment and peak calling in the THP-1 series

Raw data corresponding to ATAC-seq in THP-1 monocytes treated or not with PMA for 48h, then washed and cultured for 24 extra hours, were originally generated by Lin *et al*. (2022) and retrieved from the Gene Expression Omnibus database (accession GSE208046). Paired-end reads were trimmed and filtered by Trimmomatic version 0.39 (Bolger *et al*., 2014) with a Phred quality score threshold of 31 and a minimum read length of 36 bp. Reads were aligned to the hg38 reference genome using Bowtie 2 version 2.4.2 (Langmead & Salzberg, 2012) with the --no-1mm-upfront option. Output files were further processed with SAMtools version 1.11 (H. Li *et al*., 2009): mapped reads were converted to .bam format and sorted with the sort function, then mate information was updated using the fixmate function with the -m option, duplicates were removed thanks to the markdup function with the -r option, and finally indexes were generated using the index function.

Peaks were called using MACS2 version 2.2.7.1 (Y. Zhang *et al*., 2008) without a control. Overlapping peaks detected in several samples were merged together, keeping the minimum and maximum coordinates of all merged peaks as the new coordinates, using the merge function from BEDTools version 2.30.0 (Quinlan & Hall, 2010), in order to create a set of unique, non-overlapping peak regions for the whole series. Counts per merged peak region in each replicate of each sample were calculated using the multicov function from BEDTools.

#### Differential peak enrichment analysis

For both series, differential ATAC-seq peak enrichment between samples was assessed using “edgeR” version 3.40.2 (Robinson *et al*., 2009) in R version 4.3.3 (R Core Team, 2023) on the aforementioned read count tables. Peaks with a significantly different enrichment between two samples were defined as those with FC < 0.5 or > 2 and FDR < 0.01.

### CUT&Tag library preparation

Cells from the Neuron series were harvested as described above in two biological replicates per sample. Nuclei isolation was performed as instructed in the “Nuclei Isolation for Single Cell ATAC Sequencing” protocol from 10x Genomics^®^ (Demonstrated Protocol CG000169 Rev D) after optimizing the concentration of BSA in PBS to 0.1% instead of 0.04%, as described above.

The isolated nuclei were used for library preparation with the CUT&Tag-IT™ Assay Kit (Active motif^®^, catalog no. 53160), following the manufacturer’s instructions (Document 2200 version B6). In brief, nuclei were immobilized on concanavalin A-coated beads and magnetically separated. The nuclei were then resuspended in a buffer containing a protease inhibitor cocktail and digitonin, and 1 µg of either of the following antibodies was added to each sample: CTCF antibody (pAb) AB_2614975 (Active Motif^®^, catalog no. 61312); histone H3K4me3 antibody (pAb) AB_2615077 (Active Motif^®^, catalog no. 39060); histone H3K27me3 antibody (pAb) AB_2561020 (Active Motif^®^, catalog no. 39157); histone H3K27ac antibody (pAb) AB_2561016 (Active Motif^®^, catalog no. 39034); anti-histone H3 (mono methyl K4) antibody (Abcam^®^, catalog no. ab8895); CUTANA™ IgG Negative Control Antibody (Epicypher^®^, catalog no. 13-0042). Nuclei were left to incubate with the antibodies overnight, then once again magnetically separated, incubated for 1 hour with 1:100 guinea pig anti-rabbit secondary antibody in Dig-Wash buffer at room temperature, and finally washed several times with Dig-Wash buffer. The antibody-bound chromatin was next incubated for 1 hour with 1:100 diluted CUT&Tag-IT™ Assembled pA-Tn5 transposomes in Dig-300 buffer at room temperature, then stringently washed several times with Dig-300 buffer. The chromatin was sheared and sequence adaptors were added by tagmentation for 1 hour at 37°C. The samples were de-crosslinked for 1 hour at 55°C with buffers containing 0.1% SDS, 16 mM EDTA, and 1 µg proteinase K, which was followed by DNA extraction. Libraries were amplified by PCR with the index primers, and size selection was performed using solid phase reversible immobilization (SPRI) beads to remove primer dimers. Sequencing was performed with a HiSeq X™ Ten sequencing system (Illumina^®^) in the following conditions: R1, 150 cycles; R2, 150 cycles; Index1, 8 cycles; Index2, 8 cycles.

### CUT&Tag and ChIP-seq data processing

#### Read alignment in the Neuron series

Raw sequencing data from CUT&Tag were trimmed, aligned to the hg38 genome, sorted and deduplicated using the ENCODE ATAC-seq pipeline version 1.7.0 (D. S. Kim, 2023), with default parameters.

#### Read alignment in the THP-1 series

Raw data corresponding to ChIP-seq against mono- and trimethylation of the 4^th^ lysine of histone 3 (H3K4me3 and H3K4me1) as well as acetylation and trimethylation of the 27^th^ lysine of histone 3 (H3K27ac and H3K27me3) in THP-1 monocytes treated or not with PMA for 48h, then washed and cultured for 24 extra hours, were originally generated by Lin *et al*. (2022) and retrieved from the Gene Expression Omnibus database (accession GSE208046). Raw data corresponding to ChIP-seq against CCCTC-binding factor (CTCF) in THP-1 monocytes treated or not with PMA for 72h were originally generated by Phanstiel *et al*. (2017) and retrieved from the Gene Expression Omnibus database (accession GSE96800).

Paired-end reads were trimmed and filtered by Trimmomatic version 0.39 (Bolger *et al*., 2014) with a Phred quality score threshold of 31 and a minimum read length of 36 bp. Reads were aligned to the hg38 reference genome using Bowtie 2 version 2.4.2 (Langmead & Salzberg, 2012) with the --no-1mmupfront option. Output files were further processed with SAMtools version 1.11 (H. Li *et al*., 2009): mapped reads were converted to .bam format and sorted with the sort function, then mate information was updated using the fixmate function with the -m option, duplicates were removed thanks to the markdup function with the -r option, and finally indexes were generated using the index function.

#### Peak calling and differential peak enrichment analysis

For all CUT&Tag and ChIP-seq data, peaks were called using MACS2 version 2.2.7.1 (Y. Zhang *et al*., 2008), with the IgG data as a control for the CUT&Tag data in the Neuron series, the input as a control for the H3K4me3, H3K4me1, H3K27ac and H3K27me3 ChIP-seq data in the THP-1 series, and without a control for the CTCF ChIP-seq data in the THP-1 series; the --broad option was additionally used for H3K27me3 and H3K4me1 in both series. Overlapping peaks detected in several samples were merged together, keeping the minimum and maximum coordinates of all merged peaks as the new coordinates, using the merge function from BEDTools version 2.30.0 (Quinlan & Hall, 2010), in order to create a set of unique, non-overlapping peak regions for the whole series for downstream differential enrichment analysis. Counts per merged peak region in each replicate of each sample were calculated using the multicov function from BEDTools.

For all data, differential peak enrichment between samples was assessed using “edgeR” version 3.40.2 (Robinson *et al*., 2009) in R version 4.3.3 (R Core Team, 2023) on the read per kilobase per million (RPKM) values of each peak in each replicate of each sample, normalized to their RPKM value in the IgG or input control (except for the CTCF data in the THP-1 series, which did not come with a control). Peaks with a significantly different enrichment between two samples were defined as those with FC < 0.5 or > 2 and FDR < 0.01.

### Hi-C library preparation

iPSC, NSC and Neuron samples were cross-linked by exposure to 1× PBS (Wako^®^, FUJIFILM^®^, catalog no. 166-23555) solution supplemented with 2% formaldehyde (Thermo Fisher Scientific^®^, catalog no. 28906) for 10 min at room temperature under rotation. The formaldehyde was then quenched and washed, and cells were pelleted at 200 *g* for the iPSC sample, 300 *g* for the NSC sample and 150-200 *g* for the Neuron sample, before storage at −80°C.

*In situ* proximity ligation and Hi-C library preparation were performed with the Arima-HiC+ kit (Arima Genomics^®^, catalog no. PN A510008) and the Arima Library Prep Module (Arima Genomics^®^, catalog no. PN A303011) following the manufacturer’s instructions (Document A160134 v01). Libraries were sequenced on an Illumina^®^ NovaSeq™6000 platform using the two lanes of the S4 Reagent Kit v1.5 (300 cycles, Illumina^®^, catalog no. 20028312) in paired-end 150 sequencing mode.

### Hi-C data processing

#### Contact matrix generation in the Neuron series

Paired-end reads were trimmed and filtered by Trimmomatic version 0.39 (Bolger *et al*., 2014) with a Phred quality score threshold of 31 and a minimum read length of 60 bp. Read alignment to the hg38 reference genome, chimeric read handling and read deduplication were performed using Juicer version 2.0 (Durand, Shamim, *et al*., 2016) with the Arima option for ligation and with minor modifications to the pipeline’s script to adapt it to our high performance computing environment. After merging biological and technical replicates and keeping only read pairs with MAPQ >30, contact matrices in the .hic file format were generated with Juicer Tools version 1.22.01 (Durand, Shamim, *et al*., 2016) at 2.5 Mb, 1 Mb, 500 kb, 250 kb, 100 kb, 50 kb, 25 kb, 10 kb, 5 kb, 2 kb and 1 kb resolution.

#### Contact matrix generation in the THP-1 series

Raw data corresponding to *in situ* DLO Hi-C in THP-1 monocytes treated or not with PMA for 48h, then washed and cultured for 24 extra hours, were originally generated by Lin *et al*. (2022) and retrieved from the Gene Expression Omnibus database (accession GSE208046). The DLO Hi-C tool pipeline version 0.3.9 (Hong *et al*., 2020) was used to extract paired-end tag sequences (extract_PET function with options --cut-adapter “auto”, --linker-A “GTCGGAGAACCAGTAGCT”, --mismatch 0.3 and --rest “T^TAA”), to align reads to the hg38 genome, to build .bedpe files and to reduce noise with default options and the MseI restriction fragment map for hg38. .bedpe files were then converted to .pairs format thanks to the bedpe2pairs function from DLO Hi-C tool, and contact matrices in the .hic file format were generated from these .pairs file with Juicer Tools version 1.11.09 (Durand, Shamim, *et al*., 2016) at 2.5 Mb, 1 Mb, 500 kb, 250 kb, 100 kb, 50 kb, 25 kb, 10 kb, 5 kb, 2 kb and 1 kb resolution.

#### A/B subcompartment identification

For A/B subcompartments detection in each sample of the Neuron and THP-1 series, intrachromosomal contacts were extracted for each chromosome from the Knight-Ruiz-balanced (Knight & Ruiz, 2012) contact matrices at 100 kb resolution contained in the aforementioned .hic files, using the dump function from Juicer Tools version 2.20.00 (Durand, Shamim, *et al*., 2016). These intrachromosomal contacts were then each submitted to CALDER version 2.0 (Liu *et al*., 2021) in R version 4.3.3 (R Core Team, 2023) to identify A/B subcompartments, and the output files were combined back together for each sample thanks to the merge function from BEDTools version 2.30.0 (Quinlan & Hall, 2010).

To calculate the enrichment of histone marks in the different A/B subcompartments, the coordinates of each subcompartment were crossed with those of CUT&Tag/ChIP-seq peaks detected in the same sample using the intersect function from BEDTools. For a given type of A/B subcompartment (*s*), the lengths of all overlaps between (*s*) and peaks corresponding to a given histone modification (*h*) were summed as (*x*_*h*,*s*_), and compared to (*k_s_*) the summed length of all (*s*) subcompartments in the genome, (*N*) the summed length of all subcompartments in the genome irrespective of their type, and (*m*_*h*_) the summed lengths of all (*h*) peaks, to define a representation factor-like enrichment score (*r*_*h*,*s*_) as:

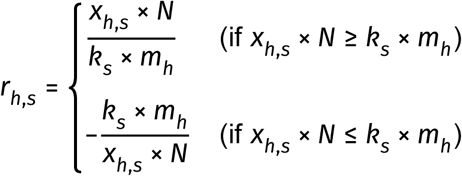

#### TAD calling

To identify TADs in each sample of the Neuron and THP-1 series, the aforementioned .hic files were first converted to .h5 format at 50 kb resolution, then normalized to a 0-to-1 range and corrected by Knight-Ruiz matrix balancing (Knight & Ruiz, 2012) thanks to the hicConvertFormat, hicNormalize and hicCorrectMatrix functions from HiCExplorer version 3.7.6 (Ramírez *et al*., 2018). TADs were then called with the hicFindTADs function from HiCExplorer and the following parameters: --minDepth 150000, --maxDepth 500000, --step 50000, --thresholdComparisons 0.5, --delta 0.25, --correctForMultipleTesting fdr.

#### Chromatin loop detection

In each sample of the Neuron and THP-1 series, chromatin loops were called using Peakachu version 2.3 (Salameh *et al*., 2020) on contact matrices at 10kb resolution, which had first been converted to .cool format from the aforementioned .hic files using the hicConvertFormat function from HiCExplorer version 3.7.6 (Ramírez *et al*., 2018), then balanced by iterative correction and eigenvector decomposition (ICE, Imakaev *et al*., 2012) with the balance function from cooler version 0.10.2 (Abdennur & Mirny, 2019) in order to be as close as possible to the data format on which Peakachu has been trained (Salameh *et al*., 2020). The training data with the closest depth value to that of each sample’s contact matrix was used to call loop anchors using the score_genome function with the --clr-weight-name weight option and the pool function with a probability threshold of 0.95. The resulting .bedpe files were crossed on both sides using the intersect function from BEDTools version 2.30.0 (Quinlan & Hall, 2010) with the CTCF peaks generated previously; based on past results (de Wit *et al*., 2015; Guo *et al*., 2015; Pugacheva *et al*., 2020; Rao *et al*., 2014; among others), only loops for which both anchors overlap a CTCF peak were kept for further analyses.

### Network embedding in 2D hyperbolic disk spaces

Networks were generated from the FDR value computed by CHiCANE for every pair of source gene/25-kb target DNA window, in each RADICL-seq replicate of the Neuron and THP-1 series.

#### Coalescent embedding

Coalescent embedding (Muscoloni *et al*., 2017; Y. Zhao *et al*., 2023) is a class of topological-based machine learning algorithms for non-linear, unsupervised dimensionality reduction and embedding of networks in a geometric space, such as a hyperbolic disk. In this study, coalescent embedding was performed using the RA-ncMCE-EA described in the original publication by Muscoloni *et al*. (2017). First, links in a given network were pre-weighted by a repulsion-attraction (RA) rule that approximates this network’s underlying geometry, defined as:

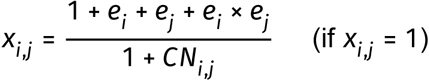

where (*x_i_*_,*j*_) is the approximated pre-weight of a link between nodes (*i*) and (*j*), (*e_i_*) is the external degree of node (*i*) and (*CN_i_*_,*j*_) is the number of common neighbors between nodes (*i*) and (*j*).

Next, the non-centered minimum curvilinear embedding (ncMCE) method (Cannistraci *et al*., 2013, 2010) was used for non-linear dimension reduction.

Angular coordinates were then calculated by equidistant adjustment (EA), while radial coordinates (*r*) were calculated by the following log-based popularity fading formula, given (*N*) the total number of nodes in the network, a node number (*i*)∈ [1, 2,…, *N*], (*K*) the curvature of the hyperbolic space (equal to −1) and (*γ*) the exponent of the power law degree distribution of the network:

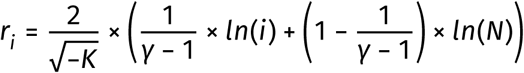

To minimize computational time and memory requirement, coalescent embedding was first conducted on networks from which nodes with degree = 1 were excluded. These nodes were added back to the embedding by post-hoc calculation of the midpoint between the angular coordinates of their parent and that of their closest neighbor. In parallel, their radial coordinates were calculated with the log-based popularity fading formula described above.

#### Calculation of power law exponent (γ) and minimum scale-free significance value (*ξ̂*)

Unlike uniform random networks, the degrees of which follow a Binomial distribution, the degrees of scale-free networks follow a power law distribution *P*(*k*) ∼ *k*^−*γ*^, where (*γ*) ≤ 3 (Barabási & Albert, 1999). Power law exponents (*γ*) and scale-free significance values (*ξ̄*) were computed following Voitalov *et al*. (2019)’s method, according to which networks are significantly power law if min(*ξ̂*) > 0.25, hardly power law if 0 < min(*ξ̂*) ≤ 0.25, and not power law if min(*ξ̂*) ≤ 0.

#### Calculation of the characteristic path length (CPL)

The characteristic path length (CPL) describes the average of all shortest path lengths (*sp_i_*_,*j*_) between all (*i*, *j*) pairs of vertices in a given network, and is calculated as follows, given (*n*) the total number of vertices in the network:

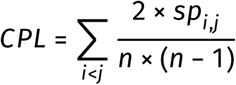

#### Calculation of the angular separability index (ASI)

The angular separability index (ASI; Muscoloni & Cannistraci, 2019) assesses the extent of separation between different node communities with regards to a specific label. In this study, the label was the chromosome to which loci belong. Practically, ASI measures the extent of angular misplacement by counting the number (*w_i_*) of nodes from different communities that intrude within the angular boundaries of a given community (*i*), then aggregating these values across all communities and comparing them to the maximum number of such misplacements observed in (*r*) null models (1,000 in this study) constructed by random shuffling of the nodes’ angular coordinates:

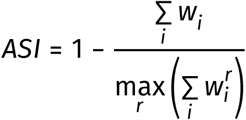

An ASI value close to 1 signifies that the communities are distinctly separated with no intermixing in their angular positions, while an ASI value near 0 indicates that the angular distribution of nodes across different communities is essentially random.

#### Permutation test for comparison of RNA and DNA radial coordinates

To assess the significance of the difference between the radial coordinates of RNA and DNA nodes, all nodes in each network replicate were pooled together and ranked in increasing order in function of their radial coordinates. Then, the 5,000^th^ first nodes were extracted and the ratio of RNA *versus* DNA nodes among those was calculated. The same process was applied after randomly permuting the radial coordinates from both types of nodes (*R*) independent times (*R* = 1,000 in this study). The RNA/DNA node ratio among the mock 5,000 most central nodes (*D_r_*) was once again calculated in each of the random control (*r*), then compared back to the real ratio (*D_obs_*) to calculate a *p* value as follows:

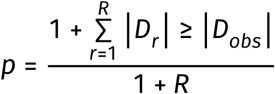

### Comparison of RADICL-seq data with replication timing and copy number variations

#### Replication timing data

High-quality replication timing data for K562 cells were originally generated by Y. Wang *et al*. (2021) (GSE148362) and directly retrieved as processed bigWig files in the hg38 genome build. Additional replication timing datasets for K562 (ENCSR591OXO) and MCF7 cells (ENCSR188HJH) were obtained from the ENCODE project (R. S. Hansen *et al*., 2009; The ENCODE Project Consortium, 2020) in .fastq format and processed based on the methodology documented in the ENCODE Repli-seq protocol. Briefly, reads were aligned to the hg38 reference genome using Bowtie version 1.3.1 (Langmead *et al*., 2009) with options -m 1, -v 2 and --no-unal. Output files were further processed with SAMtools version 1.22.1 (H. Li *et al*., 2009): mapped reads were converted to .bam format and sorted with the view and sort functions, then duplicates were removed thanks to the markdup function with the -r option, and finally indexes were generated using the index function. Reads mapping to black-listed regions contained in the ENCODE DAC Exclusion List Regions (ENCFF356LFX) were removed using BEDTools version 2.31.0 (Quinlan & Hall, 2010), then the density of the Repli-seq signal within 50 kb windows with a 1 kb step were calculated and normalized to 4,000,000 tag counts per Repli-seq phase using our custom script densityNormalization.py (accessible at https://gite.lirmm.fr/christophe.vroland/repliseq-pipeline-fastq2bw/-/blob/main/src/densityNormalization.py) in Python version 3.12.11. For each window, the percentage of Repliseq signal corresponding to each phase (G1, S1, S2, S3 and S4) was computed and used to generate bigWig files with the Python library “pyBigWig” version 0.3.24 (accessible at https://github.com/deeptools/pyBigWig).

#### Copy number variation calling

Whole genome sequencing reads in K562 and MCF7 samples were first processed using the nfcore/sarek workflow version 3.2.0, running on Nextflow version 23.10.1 with default parameters. CNVs were then identified by Control-FREEC version 11.6b (Boeva *et al*., 2011), and the output .txt files were converted to bigWig format using our custom script fantomFreecCnvToBw.sh (accessible at https://gite.lirmm.fr/christophe.vroland/fantom6-rt-and-cnv-as-confounding-factor-of-radiclseq/-/blob/main/tools/misc/fantomFreecCnvToBw.sh), with a default signal value of 2 assigned to genomic regions outside CNV calls.

#### Data preprocessing

Raw and CHiCANE-filtered RADICL-seq reads in .bedpe format from each replicate in K562 and MCF7 were merged per sample and per data type. Interactions overlapping the ENCODE DAC Exclusion List Regions (ENCFF356LFX) as well self interactions, defined as RADICL-seq reads for which the DNA fragment maps within ± 500 bp of any transcript region of the RNA fragment, were both excluded. The number of 5’ ends from RADICL-seq DNA read fragments within each 1-kb bin of the genome were then counted to generate signal tracks for downstream analysis.

For each dataset combinations (K562 RADICL-seq with GSE148362 Repli-seq, K562 RADICL-seq with ENCSR591OXO Repli-seq and MCF7 RADICL-seq with ENCSR188HJH Repli-seq), a unified per-1k-bin signal table was generated from RADICL-seq as follows: for each 1-kb bin for which replication timing data was available, the average CNV signal and the summed RADICL-seq signal (either raw or filtered) were calculated using the bigWigAverageOverBed script from UCSC (acessible at https://hgdownload.soe.ucsc.edu/admin/exe/linux.x86_64/bigWigAverageOverBed); then, RADICL-seq counts were log-transformed (natural logarithm) with a pseudocount of 0.5 to account for zero values.

#### RADICL-seq DNA read fragment count prediction

A support vector regression (SVR) implemented by the StandardScaler function from the “scikit-learn” library version 1.7.1 (Pedregosa *et al*., 2011) with Intel Extension (“scikit-learn-intelex” version 2025.8.0) in Python version 3.12.11 was employed to predict the log-transformed CHiCANE-filtered RADICL-seq signal on the DNA side, using standardized replication timing and CNVs as predictive features. Hyperparameters were optimized with the “Ray Tune” library version 2.49.0 (Liaw *et al*., 2018) using the Tree-structured Parzen Estimator implemented in the HyperOptSearch function over 100 trials. The hyperparameter search space was defined with regularization parameter (*C*) and kernel coefficient (*γ*) being sampled from a log-uniform distribution spanning 10^−3^ to 10^3^, and loss function tolerance parameter (*ε*) being sampled from a log-uniform distribution spanning 10^−4^ to 1. Models were trained on 1% of the data and validated on 4%. Pearson correlation was used as the optimization criterion, and the model with the highest validation performance was selected.

Permutation feature importance was applied in 50 iterations using the permutation_importance function from “scikit-learn” to assess the contribution of each feature to the model’s Pearson correlation coefficient.

### *K*-means clustering of chromatin-associated RNAs in function of their interaction distance distribution

CHiCANE-filtered reads in each RADICL-seq replicate were sorted into different categories based on the distance (*d*) between the start or the RNA fragment and the start of the DNA fragment. For intrachromosomal interactions, reads were assigned to a distance category (*i*) as follows:

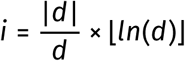

Reads corresponding to interchromosomal interactions were all assigned to distance category *ln*(10^12^), as all human chromosomes are shorter than 10^12^ bp. The number of interactions per distance category was counted for all transcript isoforms described in the SACAGE annotation with the help of the “data.table” package version 1.14.8 (Dowle & Srinivasan, 2023) in R version 4.2.3 (R Core Team, 2023). Data from all replicates were next pooled together into a single interaction counts matrix, in which each row corresponds to a transcript-in-replicate and each column corresponds to a distance category. Interchromosomal interaction counts were first divided by the total number of interaction counts for each transcript-in-replicate, then intrachromosomal interaction counts were normalized to the target window size of the corresponding distance category. To make all transcripts-in-replicate comparable with each other, values (*x_t_*_,*i*_) on each row (*t*) of the matrix were mean-centered and scaled as follows:

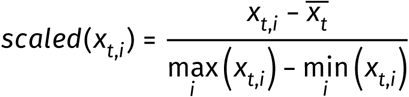

These scaled values were finally used to group all transcripts-in-replicate in function of the shape of their interaction landscape by *k*-means clustering using the kmeans function from R package “stats” version 4.2.3. The optimal number of clusters was determined by the elbow method on the within-cluster sum of squares (WCSS) plotted for *k* ranging from 2 to 15, revealing an inflection of the curve at *k* = 5.

### RADICL-seq RNA and DNA read enrichment on repeat elements

#### Presence of repeat elements within SACAGE transcripts

Regions containing repeat elements in hg38 were retrieved from the UCSC Genome Browser (Repeat-Masker track, https://genome.ucsc.edu/cgi-bin/hgTrackUi?g=rmsk; Smit *et al*., 2013) and converted to .bed format. All transcripts contained in the SACAGE annotation were next crossed with these repeat regions with respect to strandness using the intersect function from BEDTools version 2.30.0 (Quinlan & Hall, 2010); transcripts were defined as containing a specific repeat element if they overlap it by at least 6 bp.

#### Enrichment in RADICL-seq reads at repeat elements

Summed coverage in RNA read fragments was computed for every RADICL-seq replicate on all repeat elements that overlap a SACAGE transcript, extended by ± 50% of the average length of each given type of repeats, using the “EnrichedHeatmap” package version 1.28.1 (Gu *et al*., 2018) in R with parameters mean_mode=“w0”, and keep=c(0, 1) in the normalizeToMatrix function. This analysis was performed with respect to strandness, after assigning a mock RPM value of 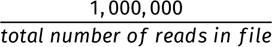 to each read. To assess whether this coverage was higher or lower than expected by chance, a random control for each type of repeat element was created with the shuffle function from BEDTools, using the merged coordinates of all transcript regions in the SACAGE annotation as input for the -incl parameter and conserving their chromosomal distribution with the -chrom option. These random controls were then subjected to the same coverage calculation as the real repeat regions.

To assess the significance of the differences in RNA read fragment coverage between real repeats and their random counterparts, both types of regions were first parsed into chunks equal to 25% of their length, including the ±50% region flanking their start and end site. All RADICL-seq RNA read fragments from all replicates were pooled for each sample, then those overlapping the parsed repeats and controls were counted with respect to strandness using BEDTools intersect. The resulting counts were summed for each type of repeat and for each 25% chunk, both across the real repeats and across their respective controls, and the difference (*D_obs_*) between the real and random regions was calculated. The assignment of each region as a “real repeat” or a “random control” was then randomly permuted (*R*) independent times (*R* = 1,000 in this study). Counts were once again summed to calculate a mock “real *versus* random” difference (*D_r_*) for each permutation (*r*), which was eventually compared back to the real (*D_obs_*) to calculate *p* values as follows:

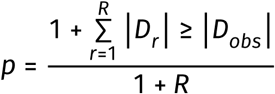

The same process was also performed with RADICL-seq DNA read fragments, this time using all repeat regions in the genome and without taking strandness into account.

cis*/* trans *ratio among interactions performed by RNAs or targeting regions containing repeat elements* To evaluate the percentage of *trans* interactions in function of the class of repeat elements present in the source RNA, RADICL-seq reads were filtered with BEDTools intersect to keep only those for which the RNA side overlap by at least 3 nt with a ± 50%-extended repeat element located within a SACAGE transcript, as well as reads originating from transcripts that do not contain any kind of repeats.

### Comparison of RADICL-seq data with CAGE, ATAC-seq, CUT&Tag/ChIP-seq and Hi-C data

#### Visualization of RADICL-seq data in Juicebox

In order to visually compare RADICL-seq data with Hi-C data, CHiCANE-filtered RADICL-seq reads in the .bedpe format for all replicates in each sample were merged and converted into tab-delimited files with the following eight columns: RNA read strand (coded as 0 for -, 1 for +); RNA read chromosome; RNA read start; 0 value; DNA read strand (coded as 0 for -, 1 for +); DNA read chromosome; DNA read start; 1 value. Next, for every read pair for which the chromosome in column 2 had a higher number than that of column 6, columns 1, 2 and 3 were switched with columns 5, 6 and 7. These files were then sorted by second, sixth, third and seventh column, before being processed by the pre function from Juicer Tools version 1.22.01 (Durand, Shamim, *et al*., 2016) with the -n (no matrix normalization) option, in order to generate RADICL-seq .hic files at 2.5 Mb, 1 Mb, 500 kb, 250 kb, 100 kb, 50 kb, 25 kb, 10 kb, 5 kb, 2 kb and 1 kb resolution, which could then be loaded in the Juicebox viewer (Durand, Robinson, *et al*., 2016).

#### Percentage of intra-TAD and inter-TAD interactions per transcript

To calculate the percentage of intra-*versus* inter-TAD RNA-DNA interactions in the Neuron and THP-1 series, TAD information in each sample was overlapped with both the RNA and DNA side of RADICL-seq reads, using the intersect function from BEDTools version 2.30.0 (Quinlan & Hall, 2010) with the -loj parameter. The same function was used to annotate the RNA read fragments with the SACAGE annotation at the transcript isoform level. With the help of the “data.table” package version 1.14.8 (Dowle & Srinivasan, 2023) in R version 4.2.3 (R Core Team, 2023), RNA-DNA read counts for each transcript were calculated in function of whether the DNA target locus is present in the same or in a different TAD as the coding locus of the source RNA. These counts were converted to percentages after division by the total number of RNA-DNA reads coming for the corresponding transcript, then all transcripts were sorted into three different categories, in function of whether <1%, >99%, or an intermediate percentage of all RNA-DNA reads originating from them correspond to intra-TAD interactions.

#### Coverage in RADICL-seq RNA and DNA reads across specific regions

The coverage in CHiCANE-filtered RADICL-seq RNA and DNA reads across chromosomes, TADs and chromatin loop regions in each replicate of the Neuron and THP-1 series were computed from the .bedpe files with the “EnrichedHeatmap” package version 1.28.1 (Gu *et al*., 2018) in R, after assigning a mock RPM value of 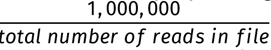 to each read and using parameters mean_mode=“w0”, and keep=c(0, 1) in the normalizeToMatrix function.

The number of CHiCANE-filtered RADICL-seq RNA reads originating from, or RADICL-seq DNA reads targeting each A/B subcompartment was calculated in each biological replicate of the Neuron and THP-1 series using the coverage function from BEDTools with the -counts option. These counts were then converted to RPKM and averaged between the biological replicates of each sample. For expression, the same strategy was applied with the multicov function from BEDTools, using .bam files from ssCAGE data as input. Significant differences in coverage for each of the three types of reads (RNA, DNA and CAGE) between different A/B subcompartments was assessed by one-way ANOVA with post-hoc Tukey test in R.

To evaluate whether source RNA loci and target DNA loci detected by RADICL-seq in the Neuron and THP-1 series are enriched in specific chromatin features (histone marks, open chromatin and TSSs), the coordinates of all RADICL-seq RNA and DNA reads in each replicate were first randomly shuffled (*R*) times (*R* = 3 in this study) using BEDTools shuffle with the -chrom option and excluding blacklisted regions (hg38 blacklist version 2, Amemiya *et al*., 2019), in order to create three “random shuffle” controls per replicate. The number of RADICL-seq RNA and DNA reads in each replicate (*n_f_*_,*obs*_) and their respective controls (*n_f_*_,1−*R*_) that map to a chromatin feature (*f*), which can be an ATAC-seq peak, CUT&Tag/ChIP-seq peak or ssCAGE TSS cluster extended by ± 500 bp, was calculated using BEDTools intersect with the -u option. The enrichment score (*e_f_*) of RNA or DNA reads among (*f*) regions in a given replicate was defined as:

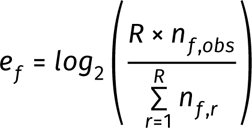

The significance of the enrichment (mean *e_f_* >0) or depletion (mean *e_f_* <0) of a feature (*f*) among source or target loci of RNA-DNA interactions in a given sample was calculated by Student’s *t*-test in R (t.test function) after pooling all (*n_f_*_,*obs*_) and (*n_f_*_,1−*R*_) values of the two biological replicates together.

In parallel, the percentage of ATAC-seq and CUT&Tag/ChIP-seq peaks as well as extended TSS clusters overlapping with at least one RADICL-seq RNA read or DNA read in each replicate of the corresponding sample was calculated using BEDTools intersect with the -u option, then averaged between the two biological replicates.

#### K-means clustering of RADICL-seq target sites

ATAC-seq, CUT&Tag/ChIP-seq and ssCAGE read counts for each peak and TSS cluster (extended by ± 500 bp), calculated with the multicov function from BEDTools, were first converted to RPKM, then averaged between the different replicates.

The coordinates of all unique DNA reads in each RADICL-seq replicate in the Neuron and THP-1 series were extracted from their CHiCANE-filtered .bedpe files, then overlapped with A/B subcompartments, ATAC-seq peaks, CUT&Tag/ChIP-seq peaks and extended TSS clusters using BEDTools intersect. For the latter three, the RPKM value associated with the whole peak or extended TSS cluster that each unique RADICL-seq DNA read intersects was assigned to that read. For A/B subcompartments, the type was converted to a numerical value ranging from 1 (A.1.1) to 8 (B.2.2), with non-assigned regions being labeled as 0. Next, distances between the center of each unique RADICL-seq DNA read and the nearest chromosome end, nearest TAD boundary and nearest loop border (start coordinate of the first anchor or end coordinate of the second anchor) were calculated using the closest function from BEDTools.

All unique DNA reads from each RADICL-seq replicate in both series were concatenated together as the rows of a single matrix, containing in each column the numerical values for each chromatin parameter described above. These values were standardized using Z-score normalization with the StandardScaler function from the “scikit-learn” library version 1.6.1 (Pedregosa *et al*., 2011) in Python version 3.11.12. *K*-means clustering was performed on rows of the scaled matrix with the MiniBatchKMeans algorithm from the “scikit-learn” library and a batch size of 100,000. WCSS were computed for *k* ranging from 2 to 20, then plotted to identify the optimal number of clusters with the elbow method, leading to *k* = 10. The resulting *k*-means clusters and their scaled values were plotted in R with the “ComplexHeatmap” package version 2.14.0 (Gu *et al*., 2016). To create summarized heatmaps, all numerical values shown in the full heatmaps created by “ComplexHeatmap”, except the A/B subcompartments, were averaged and scaled from 0 to 1 across all unique RADICL-seq DNA reads in each cluster. Each A/B subcompartment was treated as an individual feature, and RADICL-seq DNA reads were given “1” value for the subcompartment in which they were located and “0” for the others; these values were similarly averaged across the whole clusters.

To quantify the contribution of each chromatin feature to the clustering structure, 10 independent RandomForestClassifier (100 estimators) from the “scikit-learn” library were trained to predict cluster assignments, each based on a different random 1% subsample of the rescaled feature values used for clustering. Feature importance was then assessed using the permutation_importance function with 5 permutations.

In each sample, DNA reads belonging to the same cluster and located within 1.5 kb from each other were merged with the BEDTools merge function. Overlapping regions belonging to different clusters were split at their midpoint to ensure that each genomic interval was uniquely assigned to a single cluster. Directly adjacent regions assigned to the same cluster were merged into contiguous, non-overlapping segments. To enable sample comparisons within the Neuron and THP-1 series, the genome was split into a consistent set of non-overlapping intervals for each series using the BEDTools unionbedg function. If no DNA read fragments had been detected for a given interval in a given sample, meaning that this interval-in-sample had been omitted from the *k*-means clustering, the interval was labeled “None” for this sample.

The coverage in CHiCANE-filtered RADICL-seq DNA reads across each cluster interval was computed from the .bedpe files with the “EnrichedHeatmap” package in R, after assigning a mock RPM value of 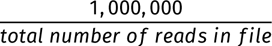 to each read and using parameters mean_mode=“w0”, and keep=c(0, 1) in the normalizeToMatrix function.

BEDTools intersect and R were used on the CHiCANE-filtered RADICL-seq .bedpe files in each replicate, in conjunction with the SACAGE annotation at the transcript level for the RNA side and the cluster intervals annotation described above for the DNA side, to explore the landscape of RNA-DNA interactions targeting each type of cluster.

The number of CHiCANE-filtered RADICL-seq linking each SACAGE transcript isoform to each target cluster interval were counted in each replicate of each series with the help of the “data.table” package in R. The resulting transcript-isoform-to-cluster count tables were used for differential analysis between samples of the Neuron and THP-1 series with the “edgeR” package version 3.40.2 (Robinson *et al*., 2009) in R. Differential interactions (DIs) were defined as those with a logFC < −2 or > 2. The coordinates of the source RNAs or of the targeted cluster intervals of these DIs were overlapped *via* BEDTools intersect with the processed data generated from CFC-seq, ssCAGE, Hi-C, ATAC-seq and/or CUT&Tag/ChIP-seq that are described in the previous sections.

Sources of DIs were defined as all SACAGE transcript isoforms performing at least one differential RNA-DNA interaction with a cluster interval. CFC-seq data in the corresponding samples was used to determine whether these sources are differentially expressed, as described above.

“Upstream” RNA-DNA interactions targeting sources of other RNA-DNA interactions were defined as all pairs of transcript/locus for which at least one non-self RADICL-seq read map, on the DNA side, to the origin locus of a source of DIs, extended by ±500 bp from its TSS, and on the RNA side, to the origin locus of another transcript. The number of such reads were counted for each “upstream” RNA-DNA interaction using BEDTools intersect and the “data.table” package in R, in each replicate of the two sample series. Like before, the resulting count tables were submitted to the “edgeR” pipeline, and differential “upstream” interactions between samples were defined as those with a logFC < −2 or > 2.

#### Identification of transcript/target locus pairs subjected to chromatin loop reorganization

All SACAGE transcripts and all target cluster intervals in the Neuron and THP-1 series were overlapped with loop anchors detected by Hi-C in each sample, as well as with the corresponding “intraloop” regions (*i.e.*, regions starting from the end coordinate of the upstream anchor and ending at the start coordinate of the downstream anchor of a given loop), using BEDTools intersect.

The resulting loop annotations where then applied to each transcript/target locus pair detected by RADICL-seq in the corresponding sample, as follows: if the coding and target locus of a DNA-interacting transcript both overlap the same or each anchor of a loop, the interaction was labeled “Loop anchor”; if they both overlap the same intraloop region, excluding the anchors, the interaction was labeled “Intraloop”; if one locus overlaps an anchor and the other overlaps the corresponding intraloop region, the interaction was labeled “Complex”; all other cases were labeled “Interloop”. Interactions for which a chromatin loop anchor is modified between the source and target loci include all interactions with a different label between two samples.

### Analysis of sources with high betweenness centrality in the RNA-DNA interaction networks

#### Network generation

Interactions lists were generated in each RADICL-seq replicate of the Neuron and THP-1 series at the source-gene-to-25-kb-target-window level, using CHiCANE-filtered data and excluding all reads corresponding to self interactions or with MAPQ < 37. From these lists, the C++ implementation of NetworkKit version 11.0 (Staudt *et al*., 2015) was used to construct individual networks for each replicate, with each DNA-interacting RNA source gene and each 25-kb target window being represented as nodes; edges indicate all significant interactions between them. The total numbers of nodes and edges for each network are detailed in Supplementary Table 6.

For each network, the degree of each node (*i.e.* the number of edges that connect to it) was calculated in R version 4.4.2 (R Core Team, 2023), while NetworkKit computed their betweenness, which reflects how often a node appears on the shortest paths between other nodes, and thereby highlights the importance of a node in connecting different parts of the network, effectively identifying nodes that act as “bridges” (Iacono *et al*., 2019; Potapov *et al*., 2005).

#### Identification of RNAs with high differential betweenness

Each source locus was ranked in terms of its betweenness value in each replicate network, then ranks were scaled from 0 to 1 and averaged between replicates. RNAs with an absolute mean scaled rank difference ≥0.325 between the first (iPSC or DMSO 96h) and the last sample (Neuron or PMA 96h) of each time series were defined as “high betweenness” (highBW) sources.

#### Subsetted network visualization

For network visualization, target windows were overlapped thanks to the intersect function from BED-Tools version 2.30.0 (Quinlan & Hall, 2010) with annotated promoters and enhancers derived from CFC-seq, as described before and in Yip *et al*. (2024). For each first *versus* last cell type and last *versus* first cell type comparison in each series, nodes corresponding to highBW sources that belong to the top 25% mean betweenness rank in one sample and have a mean betweenness rank = 0 in the opposite sample were extracted, along with all the nodes that they are connected to in each network of the corresponding series. For each sample, the node and edge information extracted from both replicate networks were pooled together, and the average weight for each replicated edge was calculated while that of un-replicated edges was divided by two. Plots were built using packages “igraph” version 2.1.3 (Csárdi & Nepusz, 2006) and “ggraph” version 2.2.1 (Pedersen, 2017) in R. To fix the position of the nodes between all three plots for a given set of highBW RNAs, their coordinates were first calculated by the Fruchterman Reingold algorithm with the “qgraph” package version 1.9.8 (Epskamp *et al*., 2012) in R, on a mock network that included all interactions performed by the selected highBW RNAs across all 3 samples of the relevant series.

#### Identification of TSS clusters targeted by highBW RNAs

CHiCANE-filtered RADICL-seq reads for which the RNA fragment map to the coding locus of a highBW RNA were extracted from each replicate of the relevant time series. The DNA fragments of these reads were then subjected to the intersect function from BEDTools with the -loj option, to be overlapped against the coordinates of TSS clusters detected by the SCAFE pipeline (as described above; Moody *et al*., 2022), extended by ± 500 bp.

#### Differential frequency analysis for interactions targeting TSS clusters

The number of CHiCANE-filtered RADICL-seq linking each SACAGE transcript isoform to each TSS cluster detected in the Neuron and THP-1 series were counted in each replicate of each series with the help of the “data.table” package in R. The resulting transcript-isoform-to-cluster count tables were used for differential analysis between samples of the Neuron and THP-1 series with the “edgeR” package version 3.40.2 (Robinson *et al*., 2009) in R. DIs were defined as those with a logFC < −2 or > 2.

#### Gene Ontology term enrichment analysis

TSS clusters were annotated with the genes that they overlap using the intersect function from BED-Tools. Gene Ontology (GO) term enrichment analyses were performed on source and target genes bearing at least one GO annotation in R using packages “biomaRt” version 2.54.1 (Durinck *et al*., 2005, 2009), “GO.db” version 3.16.0 (Carlson, 2002) and “topGO” version 2.50.0 (Alexa & Rahnenführer, 2006). The GO annotations of highBW RNAs were compared to those of all chromatin-associated RNAs detected in the same sample, while highBW-RNA-targeted and differentially active TSS clusters were compared to all similarly differentially active TSS clusters in the same samples. For each source or target category in Figure 73, the 7 most significantly enriched terms (Fisher’s exact test) were selected.

### Trait enrichment analyses from GWAS data

#### Enrichment assessment by S-LDSC

In each cell type of our collection except the AML sample, all TSS clusters detected by SCAFE (Moody *et al*., 2022) from CAGE data and which overlap with at least one RADICL-seq DNA read fragment were isolated. These regions were extended by ± 500 bp and subjected to stratified linkage disequilibrium score regression (S-LDSC; Finucane *et al*., 2015). To this end, the ldsc software version 1.0.1 (Bulik-Sullivan *et al*., 2015; Finucane *et al*., 2015) was used to partition the heritability of common single nucleotide polymorphisms (SNP), defined as those with a minor allele frequency > 5% in the European 1000 Genomes Project Phase 3 data when conditioning on the baseline model version 1.2 (Gazal, 2017) and when including enhancers present in the FANTOM5 CAT annotation (Hon *et al*., 2017). Analyses were performed on 28 highly polygenic traits, *i.e.* traits with a *Z* score for the estimation of heritability (*h*^2^*g*) > 5, for which linkage disequilibrium scores results were made publicly available by the Alkes Price group (Harvard T.H. Chan School of Public Health; data accessible at https://alkesgroup.broadinstitute.org/sumstats_formatted/). Significance of trait-annotation pairs was assessed by the Benjamini-Hochberg procedure applied to *p* values of S-LDSC’s coefficient (*τ*), with a cut-off at FDR < 0.05.

An LDSC-SEG-like (Finucane *et al*., 2018) analysis, which can be summarized as an S-LDSC analysis performed on expressed regions extended by ±100 kb in order to include regulatory variants for each transcript following suggestions from de Goede *et al*. (2021) and Võsa *et al*. (2021), was performed on all source regions of RNAs interacting with at least one TSS cluster in each cell type of our collection except the AML sample.

#### Odds ratio of overlap with genetic variants

A total of 106,736 unique phenotype-associated genetic variants with a standard significant *p* value < 5×10^-8^ were obtained from the MRC IEU OpenGWAS project (Elsworth *et al*., 2020; database accessed on September 1^st^, 2023). These GWAS variants were considered “brain-related” if their trait annotation contained one of the following terms: “*neuro*”, “*brain*”, “*intel*”, “*schizo*”, “*bipolar* ”, “*autism*”; “immune-related” variants were defined as those with a trait annotation containing at least one of the following terms: “*lymph*”, “*arthrit*”, “*inflam*”, “*immun*”, “*COVID*”, “*Treg*”, “*T cell*”, “*B cell*”, “*white blood*”, “*neutrophil*”, “*basophil*”. SNPs were also included if they display a significant linkage disequilibrium, defined as r^2^ ≥ 0.8, and are located within 25 kb of the lead SNPs identified above, based on data from the 1000 Genomes Project (The 1000 Genomes Project Consortium, 2015; data accessible at http://ftp.1000genomes.ebi.ac.uk/vol1/ftp/release/20130502/).

Our RADICL-seq data was lifted over to the hg19 genome assembly with the UCSC LiftOver program (accessible at https://genome-store.ucsc.edu/), then integrated with these GWAS results using BED-tools version 2.27.1 (Quinlan & Hall, 2010). Phenotype-associated variants were considered only if they were located either within 700 nt before and 200 nt after the RNA fragment, or within ±200 nt of the DNA fragment of a RADICL-seq read corresponding to an interaction targeting a TSS cluster. The odds of a variant overlapping an interaction were compared to a genome-wide shuffled permutation of the variant set in order to construct a 2×2 contingency table. Fisher’s exact tests were then performed in R version 4.3.2 (R Core Team, 2023) to determine the 95% confidence intervals of the odds ratios. For the focus on brain- and immune-related variants, the numbers of hits between TSS-targeting RNAs and RNA-targeted TSSs with these specific variants were compared to those obtained when overlapping these same interactions with all GWAS variants.

### Enrichment in functional elements on DNA-interacting RNAs

#### Data selection

On one hand, CHiCANE-filtered RNA-DNA interactions from each replicate of the Neuron and THP-1 series were pooled per sample. Interactions were stratified according to RNA type, and analyses were performed separately for lncRNAs and RNAs originating from protein-coding genes, based on transcript annotations. Self-interacting events (*i.e.*, RNA fragments interacting with their own genomic locus) were removed, then the remaining interactions were categorized in function of whether the RNA read fragment originate from an intronic or exonic region, and whether the interaction is performed in *cis* or in *trans*. The DNA-interacting fragments of these RNAs were represented as genomic coordinates in BED-like format for further analyses.

On the other hand, coordinates for various functional RNA elements were collected, namely conserved RNA structures (Seemann *et al*., 2017), R-loops (Sanz *et al*., 2016), RNA:DNA triplexes (Sentürk Cetin *et al*., 2019), repeat elements (Smit *et al*., 2013), evolutionary conserved sequences (Phastcons (100); Siepel *et al*., 2005), RNA-binding proteins (RBPs; Van Nostrand *et al*., 2020), sample-specific binding data for RNA Pol II from ChIP-Atlas (Zou *et al*., 2022), and combined empirical RNA G-quadruplex data from G4Atlas (Yu *et al*., 2022).

#### Background selection

For each sample and each category of interactions (protein-coding RNA or lncRNA, exonic- or intronic-derived RNA fragment, and all, *cis*-only or *trans*-only interactions), a background set was created by sampling RNAs of the same nature that are not detected by RADICL-seq, but still exhibit a similar distribution of expression levels as the DNA-interacting RNAs (“*test set*”). These background RNAs were further filtered to keep only those with transcript per million (TPM) values > 0.1 based on expression data in K562 and HepG2 cells obtained from the Cancer Cell Line Encyclopedia (Barretina *et al*., 2012), as the above-mentioned RBP eCLIP data were generated in these two cell lines.

Simulated interactions originating from these RNAs were then generated by random sampling, with respect to the distributions of tag size, relative position and overlap within intron or exon bodies found in the corresponding test set.

#### Enrichment analysis

Enrichment calculations for each functional element among each category of DNA-interacting RNAs were performed by comparing each test RNA set (*t*) with its corresponding background set (*b*), in each sample of the Neuron and THP-1 series. We first defined the interacting areas (*i*) of each set as all fragments of the RNAs that are detected by RADICL-seq (either from real or simulated data), and the non-interacting areas (*n*) as those that are not detected. Given (*c_i_*) and (*c_n_*) the fraction of all the bases forming the (*i*) and (*n*) areas that overlap with a given functional element, the enrichment ratio (*e*) in this element for the test set compared to the background was calculated as follows:

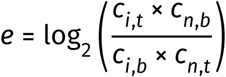

To create a negative control for the functional elements, regions with a size distribution range matching that of all combined elements were randomly selected from the whole genome. To create negative controls for the DNA-interacting RNAs at the sample and exonic/intronic levels, 10% of the entries in each test and background sets were randomly selected with respect to the original chromosomal distribution and then pooled together.

#### Enrichment of individual RBP-associated motifs

Besides evaluating the collective presence of RBP-associated elements on DNA-interacting RNAs, we also calculated these enrichment values for each RBP individually, using the same test and background RNA sets described above.

For this second analysis, the enrichment scores were calculated as contingency tables comparing the number of unique RNA interaction intervals overlapping a given element in the test *versus* background sets. A pseudocount of 1 was added to each cell of the contingency table to stabilize estimates for sparse overlaps, then a one-sided Fisher’s exact test was applied to estimate their odds ratios and associated *p* values.

To control for potential confounding effects on the association between RBPs and highly interacting RNA regions caused by the RNAs’ difference in expression, we fitted a linear regression model for each RBP. In this model, RADICL-seq read counts per interacting region were used as the dependent variable, and RNA expression (log_2_ (*TPM* + 1)) together with a binary indicator of RBP overlap were used as predictors. The regression coefficient associated with RBP presence was used to assess expression-adjusted enrichment. RBPs that did not show a significant association after expression adjustment were excluded from downstream interpretation.

### Comparison of RADICL-seq data with DRIP-seq data

#### DRIP-seq data processing, identification of R-loop regions and overlap with RADICL-seq DNA read fragments

DNA/RNA immunoprecipitation and high-throughput sequencing (DRIP-seq) in THP-1 cells published in Bamezai *et al*. (2023) data were provided in .bam format after direct inquiry to the authors.

Because this .bam file was aligned to the hg19 genome assembly but still contained unaligned reads, we converted it back into .fastq format using the bamtofastq function from BEDTools version 2.30.0 (Quinlan & Hall, 2010), then re-mapped the reads to the hg38 genome assembly using Bowtie 2 version 2.4.2 (Langmead & Salzberg, 2012) with the --no-1mm-upfront option. Output files were further processed with SAMtools version 1.11 (H. Li *et al*., 2009): mapped reads were converted to .bam format and sorted with the sort function, then duplicates were removed thanks to the markdup function with the -r option, and finally indexes were generated using the index function.

R-loop regions were then called with MACS2 version 2.2.7.1 (Y. Zhang *et al*., 2008) with the –broad option and without a control, leading to the identification of 42,545 peaks.

These peaks were next overlapped with the DNA read fragment coordinates of CHiCANE-filtered RADICL-seq data in each replicate of the DMSO 96h sample, thanks to the intersect function from BEDTools. R-loop targeting interactions were separated in function of whether the DNA read fragment overlap the coding locus of the source RNA (self interactions), or whether they correspond to *cis* (excluding self interactions) or *trans* contacts.

#### Prediction of R-loop hybridization strength

To predict the RNA:DNA hybridization strength between each transcript and R-loop region detected to interact by RADICL-seq, we subjected the sequences of the whole source RNA isoform (including its introns) and the whole R-loop region to RIsearch2 version 2.1rc1 (Alkan *et al*., 2017) with parameters -e 0 and -d 30. RNA:DNA hybridization strength for a given source-target pair was defined as the minimum value of (Δ*G*) for all RNA/R-loop fragment combinations tested by RIsearch2 for this pair.

As a negative control, the same analysis was performed using the same set of R-loop-interacting RNAs for the sources and, for the targets, a set of randomly shuffled genomic regions equivalent to the original R-loop peaks targeted by each of these RNAs, which was generated with the shuffle function of BEDTools while excluding blacklisted regions (hg38 blacklist version 2, Amemiya *et al*., 2019).

### Enrichment in molecular quantitative trait loci

Molecular quantitative trait loci (molQTLs) corresponding to DNA methylation assessed by methylation 450K arrays, H3K27ac enrichment assessed by ChIP-seq, nucleosome depletion assessed by DNAseI treatment followed by sequencing (DNase-seq), transcript accumulation assessed by 4sU sequencing (4sU-seq), gene expression assessed by RNA-seq, translation rate assessed by ribosome profiling (Ribo-seq) and molecular mass assessed by mass spectrometry were retrieved from Y. I. Li *et al*. (2016) and further processed as previously described (Young *et al*., 2022). These genomic locations were lifted over to the hg38 genome assembly with the UCSC LiftOver program (accessible at https://genome-store.ucsc.edu/), then integrated with our RADICL-seq data using BEDtools version 2.27.1 (Quinlan & Hall, 2010). In each sample of the Neuron and the THP-1 series, only molQTLs directly intersecting a TSS cluster detected by SCAFE (Moody *et al*., 2022) from CAGE data and which overlap with at least one RADICL-seq DNA read fragment were considered. The odds of a variant overlapping an RNA-targeted TSS cluster were compared to a genome-wide shuffled permutation of the variant set in order to construct a 2×2 contingency table. Fisher’s exact tests were then performed in R version 4.3.2 (R Core Team, 2023) to assess the significance of the corresponding enrichment.

### Prediction of RNA-RBP interaction and comparison with RADICL-seq data

#### RBP selection

1,163 RNA-binding proteins (RBPs) originating from the catRAPID Omics version 2 dataset (Armaos *et al*., 2021) and which were also annotated as having a subcellular localization in the nucleus in the Uniprot database (https://www.uniprot.org/) were selected as potential binding partners for DNA-interacting RNAs detected by RADICL-seq. These proteins were then further filtered to keep only those for which at least one read mapping to their coding gene could be detected by CFC-seq in either the iPSC, Neuron, DMSO 96h or PMA 96h sample.

#### RNA selection

To test against these proteins and to keep the analysis at a computationally manageable scale, we first extracted a subset of iPSC-, Neuron-, DMSO 96h- and PMA 96-highBW RNAs, as defined previously, which showed the highest change in betweenness between their two relevant samples. In practice, we first filtered out any source gene (*g*) for which

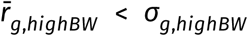

with (*r̄_g,highBW_*) the mean betweenness rank of (*g*) between the two replicate networks for the sample where (*g*) has the highest bewteenness, and (*σ_g_*_,*highBW*_) the standard deviation between these two rank values. Then, we excluded all RNAs for which

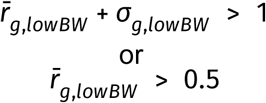

with (*r̄_g,lowBW_*) and (*σ_g_*_,*lowBW*_) the mean and standard deviation of the betweenness ranks of (*g*) in the sample where (*g*) does not have the highest betweenness. To simplify the analysis, we next removed all source genes showing a significant change in expression between the two samples of the corresponding series as determined by CFC-seq. Finally, we selected the top 100 genes with the highest 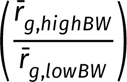 ratio.

All possible transcripts isoforms from these source genes were then retrieved from the SACAGE annotation and manually filtered to keep only those with the strongest transcriptional activity in our CFC-seq data (“*candidate RNAs*”).

In parallel and for each sample, we isolated a twice-equivalent number of transcripts that were not detected by RADICL-seq in any replicates of the iPSC and Neuron samples or of the DMSO 96h and PMA 96h samples, that are not differentially expressed between these samples, and that all have an expression level above the average of the candidate highBW RNAs in the corresponding sample (“*control RNAs*”). These control RNAs were randomly split into two control sets.

To avoid biases and redundancy due to overlapping transcript models, the coordinates of all candidate RNAs on the one hand and each and of each set of control RNAs on the other hand were merged using the merge function from BEDTools version 2.30.0 (Quinlan & Hall, 2010).

#### Prediction of RNA-RBP interaction

The nucleotide sequences of all RBPs and all highBW and control RNAs described above were extracted from the hg38 genome using the getfasta function from BEDTools, then submitted to catRAPID version 2.0 (Armaos *et al*., 2021). The raw scores (*r*) of interaction propensity yielded by catRAPID for each 51-nt RNA fragment/RBP pair were next converted to *Z*-scores (*z*) as follows:

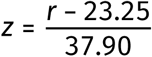

For each RBP (*p*) in each sample (*s*), we then calculated a threshold value (*t_p_*_,*s*_) equal to the 75% quantile of all *Z*-scores between (*p*) and the control transcripts for (*s*). Interactions between (*p*) and a highBW transcript (*h*) were considered significant if the mean Z-score (*z̄_p,h_*) across all of (*h*)’s 51-nt fragments is greater than (*t_p_*_,*s*_), and if at least one 51-nt fragment (*f*) of (*h*) has a *Z*-score (*z_p,h_f__*) ≥ 1.5.

RNAs and RBPs were clustered in function of their set of predicted partners using the weighted pair group method with arithmetic means (WPGMA) in R version 4.3.2 (R Core Team, 2023).

### Comparison of RADICL-seq data and ChIP-seq data from RBPs predicted to bind DNA-interacting RNAs

#### Protein ChIP-seq data selection

Using the ChIP-Atlas database (Zou *et al*., 2022) as a starting point, we listed all publicly available ChIP-seq datasets assessing the chromatin-binding of proteins in human cell types equivalent to our iPSC, Neuron, DMSO 96h and PMA 96h samples. For iPSC, we considered any data produced in either wild-type (WT) or healthy iPSCs or hESCs. For Neuron, we accepted all data generated in neurons differentiated *in vitro* from WT or healthy iPSCs or hESCs, including bipolar spindle neurons, motor neurons and other, non-specified neuron samples. For DMSO 96h, we deemed suitable any data originating from untreated THP-1 cells or from any kind of WT or healthy monocytes. For PMA 96h, we kept any data obtained from macrophages differentiated *in vitro* from monocytes as well as from any kind of WT or healthy macrophage sample. This selection yielded us 650 individual sequencing runs.

#### Protein ChIP-seq data processing

Raw data in .fastq format was retrieved for all the public ChIP-seq data described above. Reads from these files were trimmed using Trimmomatic version 0.39 (Bolger *et al*., 2014) with a Phred quality score threshold of 3 and a minimum read length of 5 bp; when applicable, only paired-reads where kept. These reads were then aligned to the hg38 reference genome using Bowtie 2 version 2.4.2 (Langmead & Salzberg, 2012) with the --no-1mm-upfront option and the paired-end-specific algorithm when applicable. Output files were further processed with SAMtools version 1.11 (H. Li *et al*., 2009): mapped reads were converted to .bam format and sorted with the sort function and, for paired-end data, mate information was updated using the fixmate function with the -m option. Duplicates were removed thanks to SAMtools markdup with the -r option, and indexes were generated using SAMtools index. Peaks were called using MACS2 version 2.2.7.1 (Y. Zhang *et al*., 2008) against the control (IgG or input) when available, and without any control in other cases.

For data series originating from the same study and in which several ChIP-seq replicates for a given pulled-down protein in the same sample were available, only peaks that were detected in at least 2 replicates were kept, then merged using the merge function from BEDtools. This merging step was then repeated for all proteins for which several ChIP-seq datasets in equivalent cell types but originating from different studies were available. In total, this process yielded 199 different peaks files, each one corresponding to a single protein in a single cell type.

#### Evaluation of DNA-binding site overlap between RBPs and their predicted RNA partners

In this analysis, we considered only highBW RNAs predicted to interact with at least one RBP by catRAPID, as described above. For each catRAPID-predicted RNA-RBP pair (*p*, *R*) for which ChIP-seq data for (*p*) was available, we defined an overlap score (*o_interacting_p,R__*) as the number (*n_p,R_real__*) of (*p*) peaks (extended by ± 1 kb) overlapping (*R*)-targeted region (also extended by ± 1 kb) compared to the number (*n_p,R_control__*) of those (*p*) peaks that overlap a set of randomly shuffled regions equivalent to the (*R*)-targeted sites, as follows:

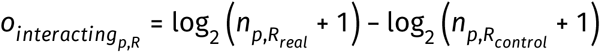

The significance of the difference between (*n_p,R_real__*) and (*n_p,R_control__*) was assessed by Fischer’s exact test, with a *p* value threshold of 0.05.

This calculation was then performed for RNA (*R*) with all chromatin-binding proteins (*c*) that were not predicted to interact with any RNA in the relevant samples, resulting in an array of “negative control” overlap scores (*o_negative_c,R__*).

By combining these overlap scores, we determined the target site colocalization score (*t_p,R_*) for RNA (*R*) and RBP (*p*) as:

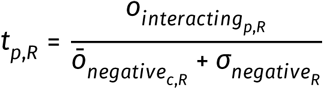

where (*σ_negative_R__*) is the standard deviation between all (*o_negative_c,R__*) values.

For further analyses, we considered that the target sites of an RBP (*p*) and its predicted RNA partner (*R*) significantly colocalize if (*o_interacting_p,R__*) > 0, if (*t_p_*_,*R*_) > 1 and if the associated Fischer’s exact test *p* value < 0.05.

## Data and code accessibility

Raw sequence files for all sequencing data generated by the FANTOM6 consortium will be uploaded to the DNA DataBank of Japan or the Gene Expression Omnibus database; corresponding accession numbers for data that has already been submitted are listed in Supplementary Tables 1 and 2.

Raw sequence files for DLO Hi-C, ATAC-seq and ChIP-seq against H3K4me3, H3K4me1, H3K27ac and H3K27me3 in PMA- or DMSO-treated THP-1 monocytes generated by Lin *et al*. (2022) are available on the Gene Expression Omnibus database (accession GSE208046). Raw sequence files for ChIP-seq against CTCF in PMA- or DMSO-treated THP-1 monocytes generated by Phanstiel *et al*. (2017) are available on the Gene Expression Omnibus database (accession GSE96800). Raw sequence files for nAnT-iCAGE data in iPSCpreF6 cells generated by Agrawal *et al*. (2024) are available on the DNA Data Bank of Japan (accession DRA013848). Raw sequence files for CAGE in MCF10A cells generated by Watanabe *et al*. (2019) are available on the Gene Expression Omnibus database (accession GSE124843). Raw sequence files for ssCAGE in MCF7 and LTED cells generated by Shu *et al*. (in preparation) are available on the Gene Expression Omnibus database (accession GSE299910). Raw sequence files for CAGE in K562 cells are available on the ENCODE database (accession ENCSR000CJN). Repli-seq data for K562 cells generated by Y. Wang *et al*. (2021) are available on the Gene Expression Omnibus database (accession GSE148362). Additional Repli-seq datasets for K562 and MCF7 are available on the ENCODE database (accessions ENCSR591OXO and ENCSR188HJH). Aligned reads for DRIP-seq in THP-1 cells generated by Bamezai *et al*. (2023) are available upon request to the authors of that study.

The processed files generated in this study from these raw sequences will be made available on the FANTOM6 website (https://fantom.gsc.riken.jp/6/). An interactive webpage to visualize the data will be available in our Zenbu viewer (Severin *et al*., 2014). Code created for this study will be accessible on the FANTOM GitHub repository (https://github.com/fantom-prj) and at https://gite.lirmm.fr/christophe.vroland/fantom6-rt-and-cnv-as-confounding-factor-of-radiclseq for the comparison of RADICL-seq data with replication timing and CNVs.

## Authors contribution

H. T., T. Kasukawa, M. Kato, C. W. Y. and J. W. S. managed the FANTOM6 project under the supervision of P. C.. P. C., H. T. and A. Lambolez conceived the present study with inputs from all authors.

K. Y., S. T., Y. Ichikawa, F. M., R. V., V. D. G., Y. Inaba, E. C., S. L., A. Lennartsson, M. Á. B. R., O. H., F. N., E. M. and S. Giussani isolated and cultured the samples.

M. Murata, S. T., M. Kato, W. H. Y., I. N., M. J. M. and S. Giussani prepared the RADICL-seq librairies, R. Pracana and X. S. processed the sequencing files with help from A. H., and A. Lambolez, W. K., R. Pracana, B. P., V. R., V. K. and A. K. analyzed the data. C. Parr, K. K., M. Murata, D. D., S. K., R. V., V. D. G., M. J. M. and S. Giussani prepared the CFC-seq libraries, C. W. Y. processed the sequencing files with help from C. Parr, and A. Lambolez and H. N.-S. analyzed the data. D. D., Y. Inaba, T. Kawashima, R. K., C. Vaagensø, I. N. and S. Giussani prepared the CAGE libraries, M. V., X. S. and H. E. processed the sequencing files and A. Lambolez and H. N.-S. analyzed the data with help from A. H.. T. Kawashima, K. O. and M. T. prepared the RNA-seq libraries and A. Lambolez, J. M. and C. C. H. processed the sequencing files. W. H. Y., W. K., I. C., S. Giussani and X. L. prepared the Hi-C and micro-C libraries and A. Lambolez processed and analyzed the Hi-C data. T. Kouno prepared the ATAC-seq libraries, A. Lambolez and J.-C. C. processed the sequencing files and A. Lambolez analyzed the data. K. Y. prepared the CUT&Tag libraries, A. Lambolez and J.-C. C. processed the sequencing files and A. Lambolez analyzed the data. C. W. Y. created the SACAGE annotation.

T. N., T. Kasukawa and A. Lambolez managed the FANTOM6 data and T. N., T. Kasukawa, H. T. and M. Kato created the FANTOM6 website. J. S. created the ZENBU visualization with inputs from A. Lambolez and H. T.

I. Abdelhamid, C. V. C. and Z. L. generated the network embeddings of RADICL-seq interactions in 2D hyperbolic disk spaces while P. S. generated the RNA-DNA interaction networks used for the study on high-betweenness sources, which was conducted by A. Lambolez. C. Vroland retrieved and processed the Repli-seq data, A. Z. and P. S. generated the CNV data and C. Vroland and C.-H. L. integrated them both with RADICL-seq data. A. Lambolez, W. K., R. Pracana, V. K. and A. K. integrated the CAGE, CFC-seq, Hi-C, ATAC-seq, CUT&Tag/ChIP-seq and RNA-seq data with the RADICL-seq data. M. R. M. and R. S. Y. performed the molQTL enrichment analysis. R. S. Y., M. Koido, C. T. and K. T. performed polygenic trait enrichment analyses from GWAS data. B. D. assessed the enrichment of functional elements on DNA-interacting lncRNAs. A. Lambolez and X. Z. retrieved, re-processed and analyzed the DRIP-seq data and A. Lambolez, X. Z., A. N. and A. Lennartsson integrated them with the RADICL-seq data. A. Lambolez and P. S. selected the highBW RNAs to test against nuclear RBPs, A. Vandelli performed the catRAPID RNA-RBP interaction prediction and A. Lambolez analyzed the catRAPID results with help from A. Vandelli. A. Lambolez selected, re-processed, analyzed and integrated the public protein ChIP-seq data. H. T., P. C., M. Kato, P. S., W. K., R. Pracana, C. W. Y., A. Vitriolo, V. K., M. Bienko, A. K., C. A. W., G. P., X. S., B. B., A. M., L. P. T., R. J., L. Carpen, A. Lennartsson, A. N., X. Z., L. B., J. N. T., M. J. L. d. H., Y. Ishikawa, P. R., G. G. T., O. H., H. K., T. H., T. Y., Y. Ciani, N. C., S. Gustincich, Y. M., N. P., N. Soranzo, C. A. S., D. G.-C., R. L., V. L., B. S., I. L., G. T., I. F., L. Calviello, C. Parr and I. Abugessaisa provided useful insights on the project design, the analyses and the results.

A. Lambolez wrote the manuscript for this paper with help from H. T. and inputs from P. C., C. V. C., B. D., C.-H. L., C. Vroland, P. S., K. Y., D. D., H. N.-S., M. Murata, C. W. Y., F. M., R. A., N. Saitoh, J. S. and M. G.. A. Lambolez, W. K., P. S., I. Abdelhamid, B. D., C. Vroland, B. P., V. R., C. W. Y., R. S. Y., M. R. M., M. Koido, C. T. and J. S. made the figures. H. T., P. C., G. P., J. N. T., D. G.-C., V. L., R. L., I. L., E. M., I. F., C. A. S., A. S., C. A. W., M. Bienko, W. K., P. S., Y. Ciani, A. M., M. J. M., L. P. T., L. Calviello, V. R., C. W. Y., C.-H. L., R. S., M. Kato, R. S. Y., M. Koido, C. V. C., B. B., N. Saitoh, R. Pracana, K. Y., F. M., I. Abdelhamid and C. Vroland internally reviewed the paper.

## Acknowledgments

Sequencing was performed by the Laboratory for Genotyping Development at RIKEN IMS and by the National Facility for Genomics and the Centre for Genomics at Human Technopole.

The authors thank Nobuyuki Takeda and Teruaki Kitakura from RIKEN IMS for their support of the IT infrastructure used for the FANTOM6 collaboration, Dr. Michael E. Ward from NIH for providing the cells constituting the “iPSC” sample, and Emi Ito from RIKEN IMS for her administrative support.

For their help providing computing resources, W. K. and M. Bienko acknowledge the National Academic Infrastructure for Supercomputing in Sweden (NAISS), R. Pracana and P. C. thank the scientific computing support team at Human Technopole, and C. V. C. and I. A. are grateful to the Center for Information Services and High-Performance Computing (ZIH) of the TU Dresden and ScaDS.AI.

P. S. and A. Z. acknowledge support from the National Genomics Infrastructure in Stockholm funded by the Science for Life Laboratory, the Knut and Alice Wallenberg Foundation and the Swedish Research Council, as well as the SNIC/Uppsala Multidisciplinary Center for Advanced Computational Science for assistance with massive parallel sequencing and access to the UPPMAX computational infrastructure.

This work was produced as part of the FANTOM6 project; we thank all members of the FANTOM6 consortium for their insights and suggestions.

## Funding

FANTOM is supported by a research grant to the RIKEN Center for Integrative Medical Sciences (IMS) from the Japanese Ministry of Education, Culture, Sports, Science and Technology (MEXT). FANTOM6 was also supported by the Human Technopole Foundation, a research foundation funded by the Italian Government under the Ministries of Economy & Finance, of Health, and of Education, University and Research.

This work additionally received financial support from the following programs and organizations: the European Union - Next Generation EU, MISSION 4, COMPONENT 2, ‘From Research to Business’, INVESTMENT 1.4, ‘Strengthening research infrastructures and creation of national R&D champions’, in relation to the project identified by code CN00000041, titled ‘National Center for Gene Therapy and Drugs based on RNA Technology and CUP CNR B83C22002860006’, for P.C.; the EU Horizon 2020 Marie-Skłodowska-Curie ENHPATHY project (grant no. 860002) for P. S. and A. Z.; the Swedish Brain Foundation (Hjärnfonden; grant no. PS2023-0023) for W. K.; the Zhou Yahui Chair Professorship award of Tsinghua University and the National High-Level Talent Program of the Ministry of Science and Technology of China (grant no. 20241710001) for C. V. C.; the Shuimu postdoctoral fellowship of Tsinghua University for I. Abdelhamid; the French National Research Agency (no. ANR-22-CE45-0031-01) for C.-H. L. and C. Vroland; the JSPS KAKENHI program (grants no. 23K14164 and no. 20H00462) for M. Koido; the Villum Foundation (grant no. 1624-00072111) and the Lundbeck Foundation (grant no. R370-2021-924) for R. K.; the Novo Nordisk Foundation (grants no. NNF20OC0059796 and no. NNF21SA0072102) for R. A.; the JST FOREST program (grant no. JPMJFR2314) for T. Y.; the EU Horizon 2020 Marie-Skłodowska-Curie DevelopMed COFUND Action (grant no. 945425) for Z. A.; the Giovanni Armenise-Harvard Foundation (Career Development Award, 2022) for I. F.; the Stewart J. Rahr Foundation - Prostate Cancer Foundation (PCF; Challenge Award 2024) for Y. Ciani; the Fondazione AIRC per la Ricerca sul Cancro (Start Up grant no. 29106), the FPRC “5xmille” 2021 Ministry of Health project (EMAGEN-LongMYND) and the Italian Ministry of Health (Ricerca corrente 2025) for E. M.; the MRC Programme (grant no. MC_UU_00035/1) for C. A. S.; the Marie Skłodowska-Curie postdoctoral fellowship (no. UNDERPIN_101063903) for L. B.; the Swedish Childhood Cancer fund (grant no. PR2022-0134) and the Swedish Cancer fund (grant no. 23 2892 Pj 01 H) for A. Lennartsson; the Swedish Research Council (grant no. 2024-06358) for S. L.; the World Premier International Research Center Initiative (WPI) from the Japanese Ministry of Education, Culture, Sports, Science and Technology (MEXT) for I. Abugessaisa.

## Competing interest

C. is a co-founder of Harness Therapeutics, a start-up company working on synthetic antisense lncRNAs that stimulate the translation of sense mRNAs. M. Koido is a consultant for Takeda Pharmaceutical Co., Ltd. None of the other authors have any conflict of interest to declare.

## Supplementary materials

**Supplementary Table 1:**
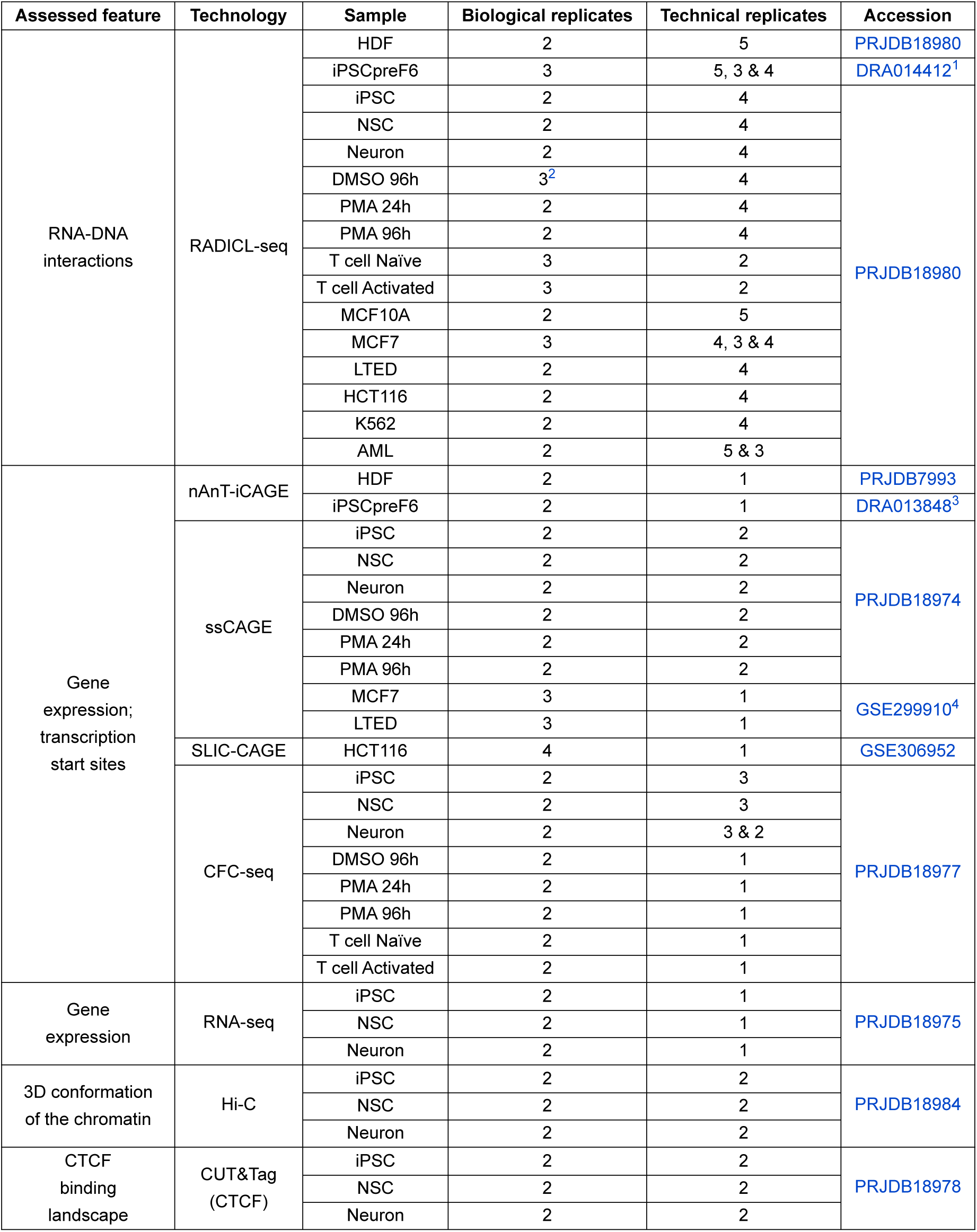
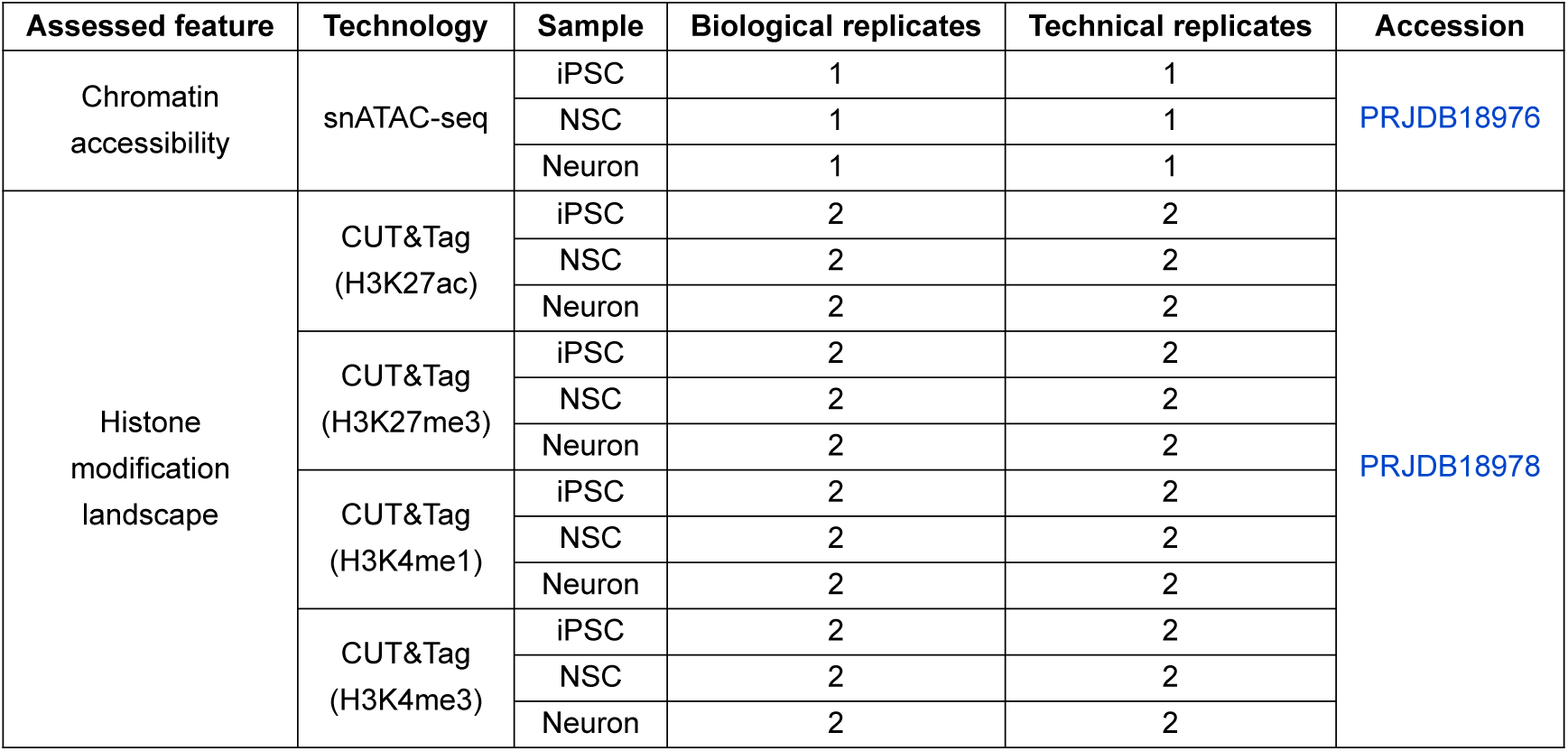
List of all datasets included in the FANTOM6 collection that were analyzed in this study.

**Supplementary Table 2:** List of all datasets included in the FANTOM6 collection that were not analyzed in this study.

| Assessed feature | Technology | Sample | Biological replicates | Technical replicates | Analyzed in... | Accession |
| --- | --- | --- | --- | --- | --- | --- |
| RNA-DNA interactions | RADICL-seq | iPSC | 2 | 3 & 4 | *In preparation* | PRJDB18980 |
|  |  | MPC | 2 | 4 |  |  |
|  |  | iMac | 2 | 4 |  |  |
| Gene expression; transcription start sites | Direct cDNA CAGE | iPSC | 3 | 1 | *Berrocal-Rubio et al., 2026* | PRJDB20614 |
|  |  | MPC | 3 | 1 |  |  |
|  |  | iMac | 3 | 1 |  |  |
|  | CFC-seq | iPSC | 2 | 1 |  | PRJDB18977 |
|  |  | MPC | 2 | 1 |  |  |
|  |  | iMac | 2 | 1 |  |  |
| 3D conformation of the chromatin | MicroC | iPSC | 1 | 3 | *In preparation* | PRJDB20615 |
|  |  | MPC | 1 | 4 |  |  |
|  |  | iMac | 1 | 4 |  |  |
| RNA-RNA interactions | PARIS | iPSC | 2 | 2 | *In preparation* | *To be determined* |
|  |  | NSC | 2 | 2 |  |  |
|  |  | Neuron | 2 | 2 |  |  |
| Enhancer-promoter DNA-DNA interactions | Promoter Capture Hi-C | iPSC | 2 | 1 | *Sahlén et al., 2025* | PRJDB18979 |
|  |  | NSC | 2 | 1 |  |  |
|  |  | Neuron | 2 | 1 |  |  |
| Radial organization of the chromatin | GPSeq | iPSC | 1 | 1 | *Kang et al., 2025* | PRJDB40303 |
|  |  | NSC | 1 | 1 |  |  |
|  |  | Neuron | 1 | 1 |  |  |
| Gene expression | scRNA-seq | iPSC | 2 | 1 | *Yip et al., in preparation* | PRJDB18975 |
|  |  | NSC | 2 | 1 |  |  |
|  |  | Neuron | 2 | 1 |  |  |
| Genetic screening | gDNA-seq | iPSC | 2 | 1 | *In preparation* | PRJDB18982 |
|  |  | NSC | 2 | 1 |  |  |
|  |  | Neuron | 2 | 1 |  |  |

**Supplementary Table 3:** List of all other, publicly available datasets that were analyzed in this study, excluding the protein ChIP-seq datasets used to compare the DNA binding sites of RNAs and their predicted RBP partners.

| Assessed feature | Technology | Cell type | Source | Accession |
| --- | --- | --- | --- | --- |
| Gene expression;<br>transcription start sites | CAGE | K562 | ENCODE | ENCSR000CJN |
|  |  | MCF10A | Watanabe *et al.*, 2019 | GSE124843 |
| 3D conformation<br>of the chromatin | DLO-Hi-C | DMSO 72h<sup>5</sup> | Lin *et al.*, 2022 | GSE208046 |
|  |  | PMA 72h<sup>6</sup> |  |  |
| CTCF binding<br>landscape | ChIP-seq (CTCF) | DMSO 72h<sup>5</sup> | Phanstiel *et al.*, 2017 | GSE96800 |
|  |  | PMA 72h<sup>6</sup> |  |  |
| Chromatin<br>accessibility | ATAC-seq | DMSO 72h<sup>5</sup> | Lin *et al.*, 2022 | GSE208046 |
|  |  | PMA 72h<sup>6</sup> |  |  |
| Histone<br>modification<br>landscape | ChIP-seq (H3K27ac) | DMSO 72h<sup>5</sup> |  |  |
|  |  | PMA 72h<sup>6</sup> |  |  |
|  | ChIP-seq (H3K27me3) | DMSO 72h<sup>5</sup> |  |  |
|  |  | PMA 72h<sup>6</sup> |  |  |
|  | ChIP-seq (H3K4me1) | DMSO 72h<sup>5</sup> |  |  |
|  |  | PMA 72h<sup>6</sup> |  |  |
|  | ChIP-seq (H3K4me3) | DMSO 72h<sup>5</sup> |  |  |
|  |  | PMA 72h<sup>6</sup> |  |  |
| R-loops | DRIP-seq | THP-1<sup>5</sup> | Bamezai *et al.*, 2023 | N/A<sup>7</sup> |
| Replication<br>timing | Repli-seq | K562 | Y. Wang *et al.*, 2021 | GSE148362 |
|  |  |  | ENCODE | ENCSR591OXO |
|  |  | MCF7 | ENCODE | ENCSR188HJH |

**Supplementary Table 4:**
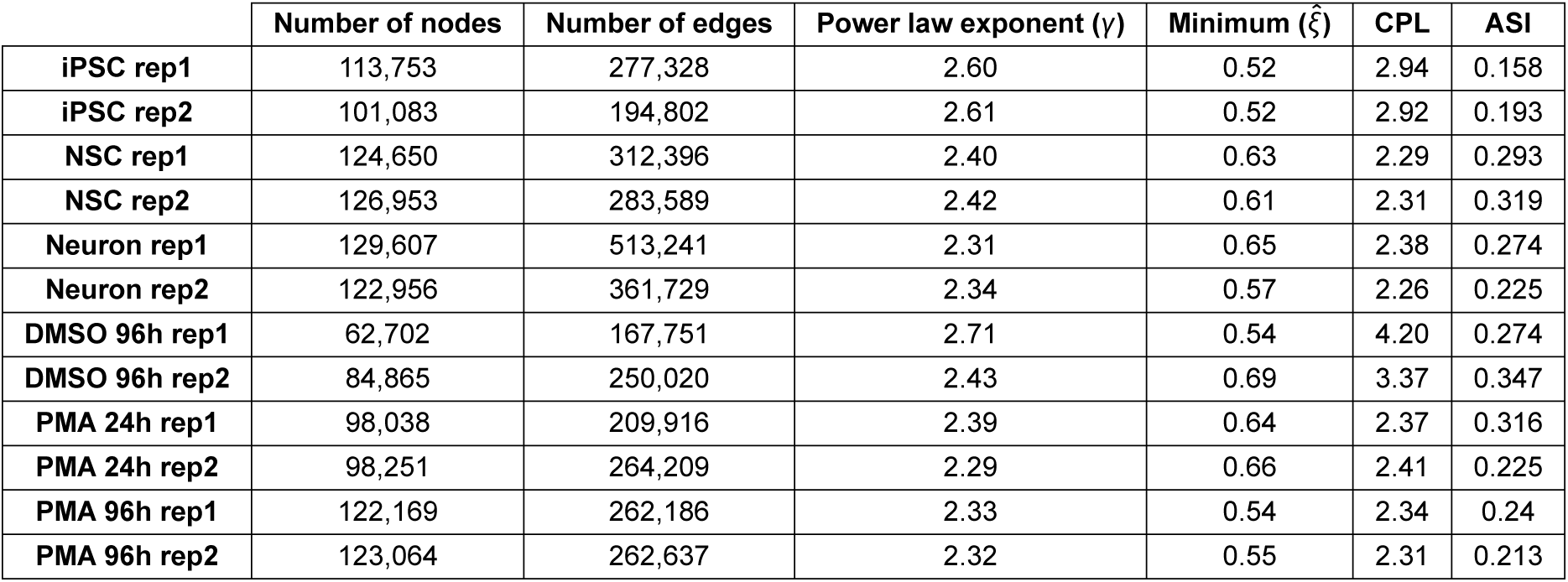
Number of nodes and edges, power law exponent (γ), minimum (*ξ̂*) (computed following the method of Voitalov et al., 2019), characteristic path length (CPL) and angular separability index (ASI) of RADICL-seq-derived networks in each replicate of the Neuron and THP-1 series.

**Supplementary Table 5:** Number of genomic intervals belonging to each of the cluster defined in Figures 35 and 36, obtained from merging RADICL-seq read fragments that were located close to each other (<1.5 kb) and assigned to the same cluster (see “Methods”), in each sample of the Neuron and THP-1 series.

|  | iPSC | NSC | Neuron | DMSO 96h | PMA 24h | PMA 96h |
| --- | --- | --- | --- | --- | --- | --- |
| **Cluster A** | 19,784 | 27,242 | 33,913 | 69 | 23 | 2 |
| **Cluster B** | 5,328 | 3,113 | 7,699 | 418 | 7,876 | 7,276 |
| **Cluster C** | 10,144 | 21,916 | 36,158 | 4,654 | 96 | 18 |
| **Cluster D** | 57,473 | 66,046 | 47,909 | 26,030 | 38,346 | 43,135 |
| **Cluster E** | 96,048 | 152,249 | 90,095 | 103,149 | 115,453 | 154,621 |
| **Cluster F** | 34,910 | 71,382 | 72,290 | 37,731 | 51,941 | 74,543 |
| **Cluster G** | 36,955 | 64,293 | 74,008 | 42,786 | 66,937 | 100,476 |
| **Cluster H** | 54,942 | 142,192 | 163,984 | 54,414 | 126,032 | 191,290 |
| **Cluster I** | 70,748 | 166,989 | 203,982 | 34,568 | 128,179 | 209,804 |
| **Cluster J** | 60,749 | 48,672 | 39,370 | 7,550 | 22,416 | 35,868 |

**Supplementary Table 6:** Total number of nodes and edges in networks generated from all non-self RNA-DNA interactions detected in each RADICL-seq replicate of the Neuron and THP-1 series.

|  | All nodes | Target nodes | Source nodes | Edges |
| --- | --- | --- | --- | --- |
| **iPSC rep1** | 103,884 | 87,599 | 16,285 | 228,880 |
| **iPSC rep2** | 93,127 | 80,524 | 12,603 | 159,790 |
| **NSC rep1** | 121,617 | 106,231 | 15,386 | 277,363 |
| **NSC rep2** | 122,453 | 107,153 | 15,300 | 245,656 |
| **Neuron rep1** | 125,044 | 105,412 | 19,632 | 468,550 |
| **Neuron rep2** | 119,073 | 106,949 | 12,124 | 328,682 |
| **DMSO 96h rep1** | 53,668 | 36,603 | 15,065 | 142,135 |
| **DMSO 96h rep2** | 77,007 | 61,616 | 15,391 | 220,269 |
| **PMA 24h rep1** | 92,702 | 84,526 | 8,176 | 189,180 |
| **PMA 24h rep2** | 94,086 | 83,861 | 10,225 | 243,486 |
| **PMA 96h rep1** | 111,368 | 101,392 | 9,976 | 225,197 |
| **PMA 96h rep2** | 115,051 | 103,709 | 11,342 | 232,804 |

## Footnotes

1 Originally published in Yip et al. (2022)

2 Biological replicates 1 and 3 were merged for downstream analyses

3 Originally published in Agrawal et al. (2024)

4 Originally published in Shu et al. (in preparation)

5 Used as a proxy for DMSO 96h

6 Used as a proxy for PMA 24h and PMA 96h

7 Obtained upon request to the original authors

## Notes

### Competing Interest Statement

P. C. is a co-founder of Harness Therapeutics, a start-up company working on synthetic antisense lncRNAs that stimulate the translation of sense mRNAs. M. Koido is a consultant for Takeda Pharmaceutical Co., Ltd. None of the other authors have any conflict of interest to declare.

